# Distinguishing Pseudotransduction and True Transduction Enables Characterization and Bioengineering of Extracellular Vesicle-Adeno-Associated Virus Vectors

**DOI:** 10.1101/2025.07.02.662894

**Authors:** Jonathan D. Boucher, Devin M. Stranford, Hailey I. Edelstein, Danielle Tullman-Ercek, Neha P. Kamat, Joshua N. Leonard

## Abstract

Adeno-associated virus (AAV) gene therapies have achieved some clinical success, with multiple products reaching regulatory approval. Encapsulation of AAV vectors within engineered extracellular vesicles (EVs) is an emerging strategy which could help overcome challenges including pre-existing anti-capsid immunity and the need for controlling targeting and tropism. To guide the development of EV-AAV technologies, we developed an assay for quantifying and controlling for the contribution of pseudotransduction to evaluations of EV-AAV-mediated transduction. We developed an AAV vector that switches its transgene output from one reporter to another when acted upon by Cre recombinase expressed in a recipient cell. Using this platform, we investigated EV-AAV transduction as a function of various engineered EV surface modifications. For actively endocytic cells (HEK293FTs), modifications that enhance EV uptake and membrane fusion influence pseudotransduction but not true transduction. Conversely, in less endocytic Jurkat T cells, modifications enhancing EV uptake enhanced both pseudotransduction and true transduction. These conclusions held across two AAV serotypes. Our results provide new insight into prior reports and suggest that effects of enhancing uptake and membrane fusion of EV-AAV vectors are recipient cell type-specific. The methods developed here unambiguously dissect EV-AAV transduction mechanisms and can guide future bioengineering of EV-AAV vectors.

## INTRODUCTION

Adeno-associated virus (AAV) gene therapies have substantial promise for the treatment of genetic diseases. Over the last decade, at least seven AAV-based gene therapies have been approved by the US Food and Drug Administration, with hundreds of clinical trials ongoing.^1^ Current efforts focus on challenges such as evading preexisting humoral immunity^2^, and reducing off-target tissue transduction^3^, which both necessitate higher vector doses that are associated with adverse effects^4,5^. One potential strategy for addressing these needs is encapsulation of AAV within engineered extracellular vesicles (EVs).

EVs are nanoscale, membrane-bound particles secreted by all cells and include exosomes, microvesicles, and particles such as apoptotic bodies.^6^ Exosomes are formed through the invagination of endosomal compartments, resulting in multivesicular bodies that fuse with the plasma membrane to release exosomes into the extracellular space. These vesicles protect their cargo—cytoplasmic RNA, proteins, and other biomolecules—from degradation by nucleases and proteases.^7^ Microvesicles bud from the cell surface and encapsulate a distinct set of biomolecules.^6^ When recipient cells take up EVs, some cargo molecules can be released into the cytoplasm to impact cell state.^8,9^ Because EVs mediate intercellular communication, exhibit lower toxicity and immunogenicity than synthetic delivery vehicles, and have longer circulation times *in vivo*, they are attractive candidates for therapeutic cargo delivery.^10–12^

An interesting observation is that when AAV particles are assembled in producer cells, secreted EVs can incorporate AAV particles capable of conferring gene expression in recipient cells, and such EV-AAV vehicles have distinct properties that are useful for gene delivery.^13^ EV-AAVs can mediate transduction of the mouse brain, even when using serotypes not generally able to cross the blood-brain barrier.^14^ EV-AAV1 and EV-AAV9 outperformed their free AAV (standard, non-enveloped AAV particle) counterparts in transduction of cochlear hair cells in explant cultures.^15^ EV-AAV1 encoding *Lhfpl5,* a protein required for mechanotransduction in outer and inner hair cells and for which loss leads to profound deafness, improved hearing and movement abnormalities in *Lhfpl5^−^*^/*−*^ mice.^15^ Upon intravitreal injection, EV-AAV2 outperformed free AAV2 in retinal transduction via delivery of GFP, increasing transduction of cell types that are transduced by free AAV2 as well as increasing the extent of transduction across different regions of the retina.^16^ EV-AAV8 encoding hF.IX enhanced correction of hemophilia B phenotypes compared to standard AAV8 vectors in C57BL/6 *F9−/−* mice, demonstrating enhanced liver targeting after intravenous injection.^17^ Both standard AAV8 and EV-AAV8 can transduce a variety of immune cell populations, including CD4^+^ T cells, CD8^+^ T cells, B cells, macrophages, and dendritic cells after systemic delivery in mice.^18^ EV-AAV8 can mediate the expression of human interleukin-2 receptor alpha chain (IL-2Rα) in multiple lymphocyte populations, suggesting that EV-AAVs could be utilized for *in vivo* cell engineering.^18^ In a lung cancer xenograft mouse model, local injection of EV-AAV6 was much more potent in transducing tumor cells compared to free AAV6.^19^ Upon intratracheal injection in C57BL/6 mice, EV-AAV6 enhanced transduction of the lung compared to either free AAV6 or a mixture of free AAV6 and EVs, demonstrating that the enhanced lung transduction by EV-AAV6 was attributable to the direct association of EVs and AAV6.^20^ EV-AAV9 delivering *SERCA2a* significantly improved cardiac function in a mouse model of myocardial infarction compared to free AAV9.^21^ In addition to improving transduction, EV-AAVs resist neutralization by AAV specific antibodies, suggesting that encapsulation can help address needs for AAV vector delivery.^14,17,19–25^ Moreover, since EVs have been engineered to target specific cell types^26–29^, it is possible that such methods could target EV-AAVs as well.

Although EV-AAV vectors are promising, they are still in the early stages of development as a therapeutic platform. EV-AAVs have been analyzed in pre-clinical models but have yet to be evaluated in human clinical trials. Several key mechanistic questions about EV-AAVs are incompletely understood. First, the functional consequences of surface-functionalizing EV-AAV vectors on recipient cell binding and particle uptake are unclear. Second, reports conflict as to how increasing EV-endosomal fusion modulates transduction by EV-AAV. An early study suggested that functionalizing EV-AAVs by incorporating the low pH-activated fusogen vesicular stomatitis virus glycoprotein (VSV-G) on the EV surface can diminish transduction of HEK293T and U87 glioma cells by AAV2, compared to EV-AAV2 lacking VSV-G.^13^ This is surprising as increasing uptake (via VSV-G binding to its receptor, low density lipoprotein receptor, LDL-R^30^) and fusion-mediated entry of the AAV capsid into the cytosol may be expected to increase transduction. While a possible explanation is that AAV2 requires processing in the endosome—capsid cleavage by the endosomal cysteine proteases cathepsins B and L^31^, autoproteolytic processing of the capsid at low pH^32^, and exposure of the VP1/2 N-terminal region exposing a phospholipase A2 domain that helps the virion escape the endosome and traffic to the nucleus^33,34^—that mechanism is untested in EV-AAV delivery. VSV-G display on EV-AAV2 showed no effect on transduction of several other glioma cell lines compared to non-targeted EV-AAV2^13^, suggesting the role of VSV-G may vary by recipient cell type. A related line of evidence is that display of rabies virus glycoprotein (RVG) on the surface of EV-AAV9 increased transduction of SH-SY5Y cells and transduction of the brain of mice compared to non-targeted EV-AAV9.^22^ RVG binds to the nicotinic acetylcholine receptor (nAChR), neural cell adhesion molecule (NCAM), p75 neurotrophin receptor, integrin β1, and transferrin receptor 1 (TfR1), promoting clathrin-mediated endocytosis of the particle.^35^ Much like VSV-G, upon encountering the low pH of the endosome, RVG undergoes conformational changes that promote endosomal membrane fusion and escape.^36,37^ Given the similarity in mechanisms used by VSV-G and RVG, the differences in transduction when displayed on EV-AAVs is surprising and may indicate recipient cell type and/or AAV serotype-specific effects. A further consideration is that AAV particles are found on both the surface and interior of EV-AAV particles, adding nuance to the interpretation of these delivery phenomena.^15,17,23^ By some estimates, in EV-AAV preparations, approximately 1-2% of EVs may incorporate AAVs.^38^ Taken together, EV-AAV preparations are a heterogenous mixture of EVs and EV-AAVs. Although EV-AAV vectors are promising, successful bioengineering of such vectors would benefit from mechanistic understanding of delivery mechanisms.

A key technical challenge when evaluating engineered EV-AAV-mediated delivery is distinguishing delivery of protein and/or mRNA originating from transgene expression in producer cells (i.e., “pseudotransduction”), from gene delivery to and resulting transgene expression in recipient cells. Pseudotransduction is also observed with lentiviral and retroviral vectors, which are also enveloped and can incorporate cytosolic proteins from producer cells.^39–42^ In certain applications, pseudotransduction may be useful, as demonstrated in dendritic cell–targeted lentiviral vectors for vaccination, in which incorporated protein antigens are employed to train induced immune reactions.^43^ However, in applications where quantification of gene delivery or transduction is desired, pseudotransduction can inflate the rates of transduction reported. Informed in part by prior observations that protein delivery can complicate evaluation of EV-mediated mRNA delivery^44,45^, we hypothesized that controlling for pseudotransduction may be particularly crucial for measuring EV-AAV transduction because achieving comparable rates of transduction typically requires higher numbers of particles for EV-AAV compared to lentiviral vectors. Thus, distinguishing transduction from pseudotransduction is critical for the engineering and evaluation of EV-AAV vectors.

In this study, we developed and validated a quantitative assay for disambiguating pseudotransduction from transduction by EV-AAVs and applied this method to investigate several biological hypotheses that inform bioengineering. Our assay comprises a modified stoplight system, in which Cre recombinase is expressed in recipient cells where it can modify an incoming engineered AAV vector to shift its transgene expression from one fluorescent protein to another. Using this approach, true transduction (hereafter, simply “transduction”) can be distinguished from pseudotransduction. With this system, we compared various EV surface functionalization strategies that drive uptake and/or membrane fusion with recipient cells and determined how these engineering choices impact pseudotransduction and transduction. Such analysis revealed new insights about EV-AAV transduction that would otherwise be obscured by pseudotransduction. Our assay enables generation and testing of new mechanistic hypotheses for studying EV-AAV biology and bioengineering these vectors.

## METHODS

### General DNA assembly

Plasmids were constructed using standard molecular biology techniques. Plasmid assembly was performed via restriction enzyme cloning. Plasmids were transformed into TOP10 competent Escherichia coli (Thermo Fisher) and grown at 37 °C. AAV vector (transfer) plasmids were transformed into NEB Stable competent Escherichia coli (New England Biolabs C3040) and grown at 37 °C. Plasmid cloning was performed primarily using standard restriction enzyme cloning with restriction enzymes (NEB), T4 DNA Ligase (NEB M0202L), and Antarctic Phosphatase (NEB M0289L). Plasmids with pcDNA-based backbones were transformed into chemically competent TOP10 *Escherichia coli* (Thermo Fisher C404010), and cells were grown at 37°C. Plasmids with AAV transgene vector backbones were transformed into chemically competent NEB Stable *Escherichia coli* (NEB C3040H), and cells were grown at 30°C.

### Source vectors for DNA assembly

The AAV transfer vector (pAAV-GFP SN), and helper plasmid (pHelper) were gifts from David Schaffer (University of California, Berkeley). The LumiScarlet gene was PCR amplified from pcDNA3-LumiScarlet, which was a gift from Huiwang Ai (Addgene plasmid # 126623 ; http://n2t.net/addgene:126623 ; RRID:Addgene_126623)^46^, and cloned into the pAAVGFP SN plasmid using AgeI and NotI. pOTTC1032 - pAAV EF1a Nuc-flox(mCherry)-EGFP (pJB037) was a gift from Brandon Harvey (Addgene plasmid # 112677 ; http://n2t.net/addgene:112677 ; RRID:Addgene_112677).^47^ Gene strings for the other Cre-responsive vectors (pJB038 - No NLS and eGFP, pJB040 - NLS and mNeonGreen, and pJB041 No NLS and mNeonGreen) were codon optimized and synthesized by ThermoFisher (GeneArt), then cloned into the pOCTTC1032 plasmid using KpnI and EcoRI restriction enzymes. RepCap plasmids for AAV2 (pRep2Cap2) and AAV6 (pRep2Cap6) were gifts from the Vector Core at the University of Pennsylvania (Penn Vector Core (RRID: SCR_022432)). pcDNA is plasmid pPD005 (Addgene plasmid # 138749 ; http://n2t.net/addgene:138749 ; RRID:Addgene_138749).^48^ psPAX2 and pMD2.G plasmids were gifted by William Miller from Northwestern University. pHIE822 was created by introducing point mutations (K47Q, R354A) in the VSV-G-encoding gene of pMD2.G using site-directed mutagenesis by PCR. pCMV-VSV-G(P127D)-Myc (pJB042) was a gift from Wesley Sundquist (Addgene plasmid # 80055 ; http://n2t.net/addgene:80055 ; RRID:Addgene_80055).^49^ pHIE963 was created using standard restriction enzyme cloning to insert a Cre recombinase gene into a third-generation lentiviral transfer vector provided by Twist Bioscience (pTwist Lenti SFFV PuroR). The Cre recombinase gene was amplified by PCR from pBS185 CMV-Cre, which was a gift from Brian Sauer (Addgene plasmid # 11916 ; http://n2t.net/addgene:11916 ; RRID:Addgene_11916).^50^ The Ultra-HEK plasmid is a commercial product made by Syenex. The plasmids used to produce Ultra-HEK lentiviral vectors are Syenex plasmids: SNX-mNG-TV3, SNX-Gagpol, SNX-Rev, and SNX-VSV-G. A list of plasmids utilized in this study can be found in **Supplementary Data 1**. Plasmid sequences can be found in **Supplementary Data 2**.

### Plasmid preparation

TOP10 or NEB Stable *E. coli* were grown overnight, shaking at 37°C or 30°C, respectively, in 50-100 mL of LB media with the appropriate selective antibiotic. The next day, DNA was prepped using a ZymoPURE II Plasmid Midiprep Kit (Zymo D4201) following the manufacturer’s instructions. DNA purity and concentrations for relevant experiments were measured with a NanoDrop 2000 (Thermo Fisher Scientific). All plasmid sequences were confirmed by whole plasmid sequencing, including AAV transfer plasmids to ensure ITR integrity.

### Cell culture

FreeStyle 293-F cells (Gibco R79007) cells were cultured in FreeStyle 293 Expression Medium (Gibco 12338018) supplemented with 1% penicillin–streptomycin (Gibco 15140-122). HEK293FT cells (Thermo Fisher R70007) were cultured in Dulbecco’s modified Eagle medium (DMEM, Gibco 31600-091) supplemented with 10% fetal bovine serum (FBS) (Gibco 16140-071), 1% penicillin–streptomycin (Gibco 15140-122) and 4 mM additional L-glutamine (Gibco 25030-081). HEK293T Lenti-X cells (Takara 632180) were cultured in HEK293FT media formulation supplemented with 1 mm sodium pyruvate from Gibco (11360070). Jurkat T cells (ATCC TIB-152) were cultured in Roswell Park Memorial Institute medium (RPMI 1640, Gibco 31800-105) supplemented with 10% FBS, 1% penicillin–streptomycin, and 4 mM L-glutamine.

Sublines generated from these cell lines were cultured in the same way. Cells were subcultured at a 1:5 or 1:10 ratio every 2–3 days, using trypsin–EDTA (Gibco 25300-054) to remove adherent cells from the plate. Freestyle 293-F cells were maintained at 37 °C with 8% CO_2_ rotating at 135 rpm on a shaking platform (Infors HT Multitron – I8000A) with orbital diameter 25 mm. HEK293FT, HEK293T Lenti-X, and Jurkat cells were maintained at 37 °C with 5% CO_2_. Freestyle 293F, HEK293FT, HEK293T Lenti-X and Jurkat cells tested negative for mycoplasma with the MycoAlert Mycoplasma Detection Kit (Lonza LT07-318).

### Cell line generation

To generate lentivirus, HEK293T Lenti-X cells were plated in 10 cm dishes at a density of 5 × 10^6^ cells per dish (6.25 × 10^5^ cells per ml). About 24 h later for Lenti-X, cells were transfected with 10 µg of lentiviral transfer vector plasmid, 8 µg of psPAX2 second generation lentiviral packaging plasmid, and 3 µg pMD2.G VSV-G-encoding plasmid via calcium phosphate transfection. Briefly, DNA was diluted with sterile H_2_O and added to CaCl_2_ (2 M) to achieve a final concentration of 0.3 M CaCl_2_. DNA-containing samples were then added dropwise to an equal-volume of 2X HEPES-buffered saline (280 mM NaCl, 0.5 M HEPES, 1.5 mM Na_2_HPO_4_) and pipetted four times to mix. After 3–4 min, the solution was vigorously pipetted eight times and 2 mL of transfection reagent per 10 cm dish was added dropwise to cells. The plates were gently swirled and incubated overnight at 37°C with 5% CO_2_. The medium was replaced the morning after transfection and cells were incubated for an additional 28–30 h. At 28 h post-medium change, lentivirus was collected from the conditioned medium. The medium was centrifuged at 500 x *g* for 2 min to clear cells, and the supernatant was filtered through a 0.45 µm PES pore filter (VWR). Lentivirus was concentrated from the filtered supernatant by ultracentrifugation in Ultra Clear tubes (Beckman Coulter 344059) at 100,420 x *g* at 4 °C in a Beckman Coulter Optima L-80 XP ultracentrifuge using an SW41Ti rotor. The supernatant was aspirated, leaving virus in ∼100 µl final volume (∼100X concentration), and concentrated lentivirus was left on ice for at least 30 min before resuspension, then used to transduce ∼1 × 10^5^ cells, plated either at the time of transduction or on the day before. When appropriate, drug selection began 2 d post transduction, using antibiotic concentrations of 1 µg ml^−1^ puromycin (Invitrogen ant-pr) on HEK293FT cells or 0.2 µg ml^−1^ puromycin on Jurkat cells. Cells were kept in antibiotics for at least 2 weeks with subculturing every 1–2 d.

### Transfection

For transfection of Freestyle 293-F cells, a transfection protocol previously used to make AAV from suspension HEK cells was utilized.^51^ Briefly, 3 d before transfection cells were plated in fresh FreeStyle 293 Expression Medium at a concentration of 1 × 10^6^ cells per mL. On the day of transfection, Freestyle 293-F cells were spun at 150g x 5 min and resuspended in fresh FreeStyle 293 Expression Medium. The cells were counted and plated at a density of 1 × 10^6^ cells per mL. Cells were transfected with 1.5 μg of DNA per million cells by PEI transfection. Plasmid DNA was added to a volume of room temperature (∼20°C) FreeStyle 293 Expression Medium, corresponding to 5% of the total volume in the flask to be transfected. The tube was mixed to evenly distribute plasmid DNA. 3 μg of PEI (Polysciences, Inc – 24765) per million cells was added to the tube and vortexed. The tube was then incubated for 10 min at room temperature to allow for precomplex formation and then added directly to the flask.

### EV-AAV production, isolation, and concentration

Freestyle 293-F producer cell lines were transfected as described in Transfection. For EV-AAVs, the AAV vector plasmid (transfer), packaging plasmid (rep/cap), and adenoviral helper plasmid (pHelper) were transfected in equal DNA mass ratios up to the maximum DNA amount for the mixture. In transfections used to make EV-AAV vectors including two types of EV surface protein, the mass ratio of transfected plasmids was 1:1:1:1.59:1.41 for the AAV vector plasmid (transfer), packaging plasmid (rep/cap), adenoviral helper plasmid (pHelper), pMD2.G VSV-G plasmid, and Ultra-HEK plasmid respectively. For control vectors in which one such EV surface protein was omitted, pcDNA was included instead (at the equivalent DNA mass) to ensure the same total mass of DNA was transfected into each flask. In experiments comparing various single EV surface functionalizations, an equivalent mass of plasmid encoding each surface protein was used, with the remaining mass made up by the AAV vector plasmid (transfer), packaging plasmid (rep/cap), adenoviral helper plasmid (pHelper) at equivalent mass ratios. For exact DNA doses, see **Supplementary Data 3**. 3 d after transfection, EV-AAVs were collected from the conditioned medium by differential centrifugation. Briefly, conditioned medium was cleared of debris by centrifugation at 300 x g for 10 min to remove cells followed by centrifugation at 2,000 x g for 20 min to remove dead cells and apoptotic bodies in a Sorvall Legend X1R Centrifuge (ThermoFisher Scientific 75004261) with a TX-400 4 x 400 mL Swinging Bucket Rotor. All centrifugation steps were performed at 4 °C. The supernatant underwent tangential flow filtration using a KrossFlo KR2i TFF System with a MICROKROS 20CM 500KD filter (Repligen C02-S500-05-S). Supernatant was flowed through the system at 24.99 mL/min at a constant transmembrane pressure of 6.0 psi. The supernatant was concentrated to approximately 5 mL by TFF. In between samples, the TFF was cleaned with at least 50 mL of 0.2 M NaOH that was sterile filtered by a 0.22 μm bottle top filter (Corning 431118). The system was then washed with at least 100 mL of PBS filtered by a 0.1 μm bottle top filter (Corning 431475). The TFF system was kept in 0.05% sodium azide (EMD SX0299-1) in H_2_O to prevent contamination. After TFF, the EV-AAVs were further concentrated to approximately 0.5 mL (∼800X concentration) in Amicon Ultra-4 100K Centrifugal Filters (Millipore UFC810024) that were pre-washed with 0.1 μm bottle top filtered PBS. 0.5 mL of concentrated EV-AAVs were treated with Benzonase (25 U/mL, EMD Millipore 70664) supplemented with MgCl_2_ at a concentration of 2 mM for 1 h at 37 °C. Samples were then purified to remove free AAV and other non-EV proteins using one of the methods below.

### EV iodixanol purification

Iodixanol gradient purification of EVs was modified from a previous report.^21^ Briefly, 1x PBS-MK buffer was prepared by dissolving 26.3 mg of MgCl2 and 14.91 mg of KCl in 100 mL of 1x PBS. The solution was sterilized by passing it through a 0.22 μm filter (Corning 431118). 1 M NaCl/PBS-MK buffer was prepared by dissolving 5.84 g of NaCl, 26.3 mg of MgCl2 and 14.91 mg of KCl in 1X PBS in a final volume of 100 mL. The solution was sterilized by passing it through a 0.22 μm filter (Corning 431118). Three iodixanol (Sigma Alrich D1556) gradients were prepared from these solutions: 25% iodixanol (5 mL of 60% iodixanol, 7 mL of 1x PBS-MK buffer), 40% iodixanol (6.7 mL of 60% iodixanol and 3.3 mL of 1x PBS-MK buffer) and 60% iodixanol (10 mL of 60% iodixanol). These solutions were layered into UltraClear tubes (Beckman Coulter 344059) in the following order: 1 mL 60% iodixanol, 3 mL 40% iodixanol, and 5 mL 25% iodixanol. EVs were layered on top of the 25% iodixanol layer. The tubes were spun in a Beckman Coulter Optima L-80 XP ultracentrifuge with an SW41 Ti rotor at 190,000 x g for 3 h at 18°C. After the spin, EVs were visible as a white band in the 25% layer and were removed with a syringe with 18G needle. To remove the iodixanol from the EVs, EVs were placed in a new UltraClear tube and diluted in PBS. The tubes were spun at 120,000 x g for 90 min at 4 °C to pellet the EVs. After the spin, the supernatant was aspirated, and EV pellets were left in ∼100–200 µl of PBS. EVs were incubated on ice for at least 30 min after supernatant removal before resuspension in a total of 500 µL of PBS.

### EV size exclusion chromatography purification

Size exclusion chromatography was performed using qEVoriginal (35 nm) Gen2 columns (Izon ICO-35) attached to an Automatic Fraction Collector V1 (Izon). We followed the manufacturer’s instructions by equilibrating the SEC column in PBS sterilized with 0.1 µM bottle top filters (Corning 431475). 0.5 mL of EVs were placed on the column and EV fractions 1-4 were collected based on the manufacturer’s instructions and a previous report of EV-AAV purification with these SEC columns^25^. Fractions 1-4 were combined and concentrated to approximately 250 µL in Amicon Ultra-4 100K Centrifugal Filters (Millipore UFC810024) that were pre-washed with 0.1 µm bottle top filtered PBS.

### EV concentration determination by Nanoparticle Tracking Analysis (NTA)

EV concentration was determined by NanoSight analysis. Samples were diluted in PBS sterilized with 0.1 µM bottle top filters (Corning 431475) to concentrations on the order of 10^8^ particles per ml for analysis. NanoSight analysis was performed on an NS300 (Malvern), software version 3.4. Three 30 s videos were acquired per sample using a 642 nm laser on a camera level of 14, an infusion rate of 30 and a detection threshold of 7. Default settings were used for the blur, minimum track length and minimum expected particle size. EV concentrations were defined as the mean of the concentrations calculated from each video. Size distributions were generated by the software.

### AAV crude cell lysate production

AAV was harvested from the Freestyle 293-F cell lysates of cells transfected to make non-targeted EV-AAVs (no EV surface modifications). Briefly, 72 h post-transfection the EV-AAV containing supernatant was removed and the cells were resuspended and washed with PBS sterilized with 0.1 µM bottle top filters (Corning 431475). The cells were spun at 1000 x g for 7 min in a Sorvall Legend X1R Centrifuge (ThermoFisher Scientific 75004261) with a TX-400 4 x 400 mL Swinging Bucket Rotor. The supernatant was removed, and the cells were mixed with AAV lysis buffer corresponding to 2% of the initial culture volume. AAV lysis buffer was made by combining 2 mL of 5 M NaCl, 0.158 g Tris-HCl and H_2_O up to 100 mL. The solution was pH-adjusted to 8.5 and sterile filtered by a 0.22 µm bottle top filter (Corning 431118). The cell pellet resuspended in AAV lysis buffer was then frozen in a bath of dry ice and ethanol, followed by thawing in a 37°C bead bath. After each thawing the solution was vortexed vigorously for 30 s. The cell pellet underwent three rounds of freeze/thaws. Benzonase (50 U/mL, EMD Millipore 70664) was added to the mixture and incubated for 30 min at 37°C. The solution was transferred to 1.5 mL microcentrifuge tubes and spun at 7000 rpm for 15 min in a microcentrifuge (Eppendorf Centrifuge 5424). The supernatant, clarified AAV crude lysate, was transferred to a clean microcentrifuge tube and stored at 4°C until further processing.

### AAV purification by iodixanol gradient centrifugation

1x PBS-MK buffer was prepared by dissolving 26.3 mg of MgCl2 and14.91 mg of KCl in 100 mL of 1x PBS. The solution was sterilized by passing it through a 0.22 μm filter (Corning 431118). 1 M NaCl/PBS-MK buffer was prepared by dissolving 5.84 g of NaCl, 26.3 mg of MgCl2 and 14.91 mg of KCl in 1X PBS in a final volume of 100 mL. The solution was sterilized by passing it through a 0.22 μm filter (Corning 431118). Four iodixanol (Sigma Alrich D1556) gradients were prepared from these solutions: 15% iodixanol (4.5 mL of 60% iodixanol and 13.5 mL of 1 M NaCl/PBS-MK buffer), 25% iodixanol (5 mL of 60% iodixanol, 7 mL of 1x PBS-MK buffer and 30 μL of phenol red), 40% iodixanol (6.7 mL of 60% iodixanol and 3.3 mL of 1x PBS-MK buffer) and 60% iodixanol (10 mL of 60% iodixanol and 45 μL of phenol red). These solutions were layered into UltraClear ultracentrifuge tubes (Beckman Coulter 344059) in the following order: 1.35 mL 60% iodixanol, 1.35 mL 40% iodixanol, 1.6 mL 25% iodixanol, 2.7 mL 15% iodixanol, and up to 4 mL of AAV Crude Lysate. The tubes were spun in a Beckman Coulter Optima L-80 XP ultracentrifuge with an SW41 Ti rotor at 190,000 x g for 3 h at 18°C. AAV was removed by inserting a syringe with 18G needle at the border between the 60% and the 40% iodixanol fractions, with the needle facing upward to the lower portion of the 40% iodixanol fraction where the viral particles accumulate. The entire 40% iodixanol layer (transparent) was removed without disturbing the layer of protein contaminants that accumulate at the interphase between the 40% and 25% iodixanol layers. To remove the iodixanol this layer was mixed 1:1 by volume with PBS and added to an Amicon Ultra-15 100K spin filter. The spin filter was spun at 3000 x g until the volume remaining in the column was less than 1 mL. PBS was added to the column to the full volume and spun at 3000 x g until 1 mL remained in the column. This was PBS washing was repeated 4 more times to remove iodixanol and concentrate the AAV to its final volume.

### AAV purification by magnetic beads

Free AAV was purified using Dynabeads Capture Select AAVX Magnetic Beads (Thermofisher Scientific 2853522001) by the manufacturer’s protocol. Briefly, 40 μL of bead slurry was added to a low protein binding microcentrifuge tube with 460 μL of PBS. The tubes were placed on a magnetic stand and once the beads were collected on the side of the tube the supernatant was removed. The beads were then mixed with 500 μL of PBS, the tubes were placed on a magnetic stand, and the supernatant was removed. 1 mL of AAV crude lysate was added to the beads and the tubes were put on a rotating platform to continuously mix the beads and lysate for 30 min. After 30 min the beads were collected with the magnetic stand and the supernatant was removed. 500 μL of PBS was mixed with the beads, the beads were collected on the magnetic stand, and the supernatant was removed. This PBS wash was repeated again. 20–50 μL of elution buffer was added to the beads and mixed well. The elution buffer (50 mM citric acid, pH 2.5–3.0) and beads were incubated at room temperature for 10 min with occasional pipetting. The beads were collected on the magnetic stand and the supernatant containing purified AAV was collected. The eluted AAV mixture was neutralized by adding 1 μL of Neutralization Buffer (1 M Tris-HCl, pH 8.7) for each 10 μL of eluate.

### Physical titer determination by qPCR

First, 5 μL of the EV-AAV, purified AAV, or clarified crude lysate samples were mixed with 39 μL nuclease-free H_2_O, 5 μL of 10x DNaseI buffer (ThermoFisher B43) and 1 μL of DNaseI (ThermoFisher EN0521) to eliminate plasmid DNA, incubating at 37°C for 1 h followed by a 15 min inactivation step at 75°C. Samples were placed on ice. Then, viral genomic DNA (vgDNA) was extracted using the High Pure Viral Nucleic Acid Kit (Roche 11858874001), following manufacturer instructions. qPCR analysis was performed by amplifying a sequence within the LumiScarlet reporter using primers CAGGGAGGTTTGACCAGTTT (forward), and GGATATCGATCTTCAGCCCATT (reverse) or the mCherry reporter using the primers AGGACGGCGAGTTCATCTA (forward), and CCCATGGTCTTCTTCTGCATTA (reverse). Primers were designed using IDT’s PrimerQuest Tool. A plasmid containing the LumiScarlet transgene (pJB035) or mCherry transgene (pJB040) were used as a standard; serial dilutions were prepared using EASY dilution (Takara Bio 9160), plating each dilution in duplicate. The vgDNA samples were diluted in nuclease free H_2_O, plating 4 serial dilutions per sample (in duplicate). For each qPCR reaction (each in a 96-well), 5 μL of vgDNA sample were mixed with 10 μL Universal SYBR Master Mix 2x (ThermoFisher K0221), 0.15 μL of 100 μM forward primer, 0.15 μL of 100 μM reverse primer, and 4.7 μL of nuclease free H_2_O. qPCR plates were sealed and spun at 500 x g for 2 min before placing in Bio-Rad C1000 Thermal Cycler with CFX96 Real-Time System. Samples run to detect LumiScarlet were heated to 98°C for 3 min, followed by 40 cycles of: 98°C for 15 s, 62°C for 30 s. Then, a final step of 0.5°C increments every 5 s from 65°C to 95°C was performed. Samples run to detect mCherry were heated to 98°C for 3 min, followed by 40 cycles of: 98°C for 15 s, 55°C for 30 s. Then, a final step of 0.5°C increments every 5 s from 65°C to 95°C was performed.

#### Transduction assays

For most experiments, EV-AAV or free AAV vectors were diluted in 0.1 μm bottle top filtered PBS so that 100 μL of the dilution contained 10^9^ vector genomes. 100 μL of the diluted vector was added to each well of a 24-well plate. Recipient cells were diluted to 10^4^ cells per 900 μL of media and added to the wells with vector, for a final dosage of 10^5^ vector genomes per cell. For the experiments depicted in **Figure 4E-F** and **Figure S17E-F**, 2 x 10^9^ vector genomes were added per well (2 x 10^5^ vector genomes per cell). To provide some rationale, our initial transduction assays (**Figure 4C-D, Figure S17B-C)** yielded fairly low transduction efficiencies, so the dose was increased in subsequent experiments (**Figure 4E-F, Figure S17E-F**). Negative control wells received 100uL of 0.1 μm bottle top filtered PBS. After the addition of all vectors and recipient cells, the plate was vigorously mixed and placed in an incubator. 5 d post transduction, cells were harvested for flow cytometry using 0.05% Trypsin-EDTA with phenol red for HEK293FT recipient cells for 5 min followed by quenching with medium. The resulting cell solution was added to at least 2 volumes of FACS buffer (PBS pH 7.4, 2–5 mM EDTA, 0.1% BSA) supplemented with DAPI (to identify live cells) at a final concentration of 3 μM. Jurkat cells were pipetted from the plate directly into FACS tubes containing FACS buffer supplemented with DAPI. A single FACs tube of PBS treated cells was added to a tube containing FACS buffer without DAPI to identify live cells (DAPI negative). Cells were centrifuged (150 × g for 5 min), supernatant was decanted, fresh FACS buffer, with or without DAPI, was added, and cells were resuspended by briefly vortexing.

For experiments involving lentiviral transduction of HEK293T and HEK293FT by lentiviral vectors including Ultra-HEK, similar methods were used. Cell lines were harvested, counted, and plated in 96-well, TC treated, PBS-moated, flat-bottom plates at a density of 10^5^ cells/mL in a volume of 100 μL per well (10^4^ cells/well). Crude LV supernatant was diluted 5-fold seven times in complete DMEM to generate a seven-point dilution curve starting from the first dilution. Cells were transduced by adding 100 μL/well of LV (each dilution) and were incubated for 3 d at 37°C. Before analysis of transduction by flow cytometry, cells were harvested by aspirating the media, trypsinizing cells for 5 min at room temperature, quenching with complete DMEM, transferring lifted cells to a non-treated, round-bottom 96-well plate, spinning cells at 150 x g for 5 min, and resuspending in 30 μL of FACS buffer (PBS pH 7.4 with 0.05% BSA and 2 mM EDTA) + DRAQ7 viability dye (Thermo Fisher D15106, diluted 1000x). DRAQ7 was used to identify live cells (DRAQ7 negative) for analysis.

#### Microscopy imaging

Microscopy was conducted on a Keyence BZ-X800E fluorescence microscope and a Nikon CFI S Plan Fluor ELWD 20XC objective (MRH08230). LumiScarlet fluorescence was imaged with a Keyence BZ-X Filter TexasRed (Keyence, OP-87765). Images were taken using the Keyence BZ Series Viewer Software (Version 01.03.00.01) in high resolution mode. Scale bars were added to images with the BZ-X800 Analyzer Software (Version 1.1.2.4).

### Transfected producer cell analysis

Freestyle 293-F producer cell lines were transfected as described in Transfection. For each condition, 3 flasks of 20 x 10^6^ cells in 20 mL of Freestyle media were plated. Cells were transfected with 30 μg total of DNA. For each condition and exact DNA doses, see **Supplementary Data 3**. 3 d after transfection, 1 mL of cells and media were removed from each flask, placed in a FACS tube and spun at 150 x g for 5 min. The media was removed, and the cells were washed with PBS. The cells were spun at 150 x g for 5 min and the supernatant was removed. 1 mL of 4% paraformaldehyde in PBS (Thermo Scientific, J61899.AK) was added to each tube, gently vortexed and fixed for 20 min at room temperature. Cells were then washed 3 times with 1 mL of PBS. After the final wash, cells were resuspended in FACS buffer and analyzed by flow cytometry.

### Cycloheximide (CHX) treatment of recipient cells

1.0 x 10^5^ HEK293FT recipient cells per well in 1 mL of DMEM were plated in 24-well plates the night before treatment. Treatments (EV-AAV2, No Rep/Cap EVs, free AAV2) were diluted in 0.1 μm bottle top filtered PBS so that 100 μL of the dilution contained 10^9^ vector genomes. No Rep/Cap EVs were dose-matched to EV-AAV2 by particle count as determined by nanoparticle tracking analysis. 100 μL of the diluted vector was mixed with 900 μL of DMEM. As a negative control, 100 μL of PBS was mixed with 900 μL of DMEM. Three groups of plates were made. Group 1 plates were treated with EV-AAV2, No Rep/Cap EVs, free AAV2 or PBS for 4 h and analyzed by flow cytometry. Group 1 plates were used to measure fluorescence associated with uptake of EVs at the time of cycloheximide or DMSO treatment. Plates in Group 2 and 3 were treated with EV-AAV2, No Rep/Cap EVs, free AAV2 or PBS for 4 h. After 4 h the media was removed and replaced with fresh DMEM containing either 20 μg/mL cycloheximide dissolved in dimethyl sulfoxide (DMSO) for Group 2 or DMSO at an equivalent volume for Group 3. 4 h post CHX or DMSO treatment, Group 2 and Group 3 cells were analyzed by flow cytometry. In each group, the mean PE-Texas Red fluorescence (AU) was determined across 3 treated wells and normalized to the mean fluorescence of PBS treated cells within that group to facilitate comparisons across treatment groups.

### Lentiviral vector production for Ultra-HEK analysis

Lentivirus (LV) samples were produced in adherent Lenti-X cells seeded at a density of 3.2 x 10^5^ cells/mL in a volume of 0.5 mL DMEM in 24-well TC-treated plates 24 h prior to transfection. Cells were transfected with 0.4 μg total DNA comprising 50% transfer vector, 29% gag/pol, 16% rev, and 5% VSV-G for the VSV-G only composition and 46% transfer vector, 29% gag/pol, 16% rev, 5% VSV-G, and 4% Ultra-HEK for the VSV-G and Ultra-HEK composition. The transfer vector used encodes constitutive expression of mNeonGreen. Plasmids used for lentivirus production are Syenex’s commercial plasmids: SNX-mNG-TV3, SNX-Gagpol, SNX-Rev, SNX-VSV-G, and Ultra-HEK. Ultra-HEK is a plasmid designed to increase lentiviral transduction of HEK293-derived cells. Plasmids and PEI MAX transfection reagent (Polysciences 24765) were diluted in Opti-MEM and mixed in equal volumes at a 4:1 mass ratio of PEI to DNA and complexed for 24 min prior to transfection. Medium was changed 12–16 h later. At least 28 h post media change, LV was harvested from the conditioned medium and centrifuged at 1000 x g for 10 min at 4°C, then aliquoted and stored at -80°C until use.

### Analytical flow cytometry and analysis

Most flow cytometry was performed on a BD LSR Fortessa Special Order Research Product using the 552 nm laser for mCherry or LumiScarlet (610/20 filter) and the 488 nm laser for eGFP or mNeonGreen (530/30 filter). At least 10,000 live cells were collected per sample for analysis. Data were analyzed using FlowJo v10 (FlowJo, LLC). Briefly, cells were identified using a forward scatter area (FSC-A) versus side scatter area (SSC-A) plot and gated for single cells using an FSC-A versus forward scatter height (FSC-H) plot (**Supplementary Information (Figures S1,S2,S3)**). Freestyle 293 cells were analyzed in the single cells gate (**Figure S3**). HEK293FT (**Figure S1**) and Jurkat (**Figure S2**) live single cells were identified using a DAPI-A versus SSC-A plot. Transduction was determined by the percentage of live single cells that were brighter than approximately the 0.1% of the brightest cells in the untransduced (PBS treated) cell samples (**Figure S1**). Mixed delivery was quantified by the mean of PE-Texas Red fluorescence of live single cells. This mean fluorescence intensity was converted to MEPTRs by using UltraRainbow Calibration Particles (Spherotech). Briefly, to determine the conversion from the arbitrary unit (au) of fluorescence recorded by the flow cytometer to a standardized fluorescence unit (Molecules of Equivalent PE-TexasRed (MEPTR)), UltraRainbow Calibration Particles were run as part of each flow cytometry experiment, except the AAV-LumiScarlet experiment (**Figure 1**). This reagent contains nine subpopulations of beads, each of a specific size and with a known intensity of various fluorophores, which are provided with calibration data for each lot. The total bead population was identified by SSC-A vs. FSC-A gating, and subpopulations were identified through two fluorescent channels (**Figure S4**). A calibration curve was generated for the experimentally determined au vs. manufacturer supplied MEPTR, and a linear regression was performed with the constraint that 0 au equals 0 MEPTR (**Figure S4B,** left). The slope from the regression was used as the conversion factor. For experiments involving the quantification of mNeonGreen fluorescence intensity, the mean FIT-C fluorescence of live single cells was converted to MEFLs by using UltraRainbow Calibration Particles (Spherotech). Briefly, to determine the conversion from the arbitrary unit (au) of fluorescence recorded by the flow cytometer to a standardized fluorescence unit (Molecules of Equivalent Fluorescein (MEFLs)), UltraRainbow Calibration Particles were run as part of each flow cytometry experiment. A calibration curve was generated for the experimentally determined au vs. manufacturer supplied MEFL, and a linear regression was performed with the constraint that 0 au equals 0 MEFL (**Figure S4B,** right). The slope from the regression was used as the conversion factor.

**Figure 1:**
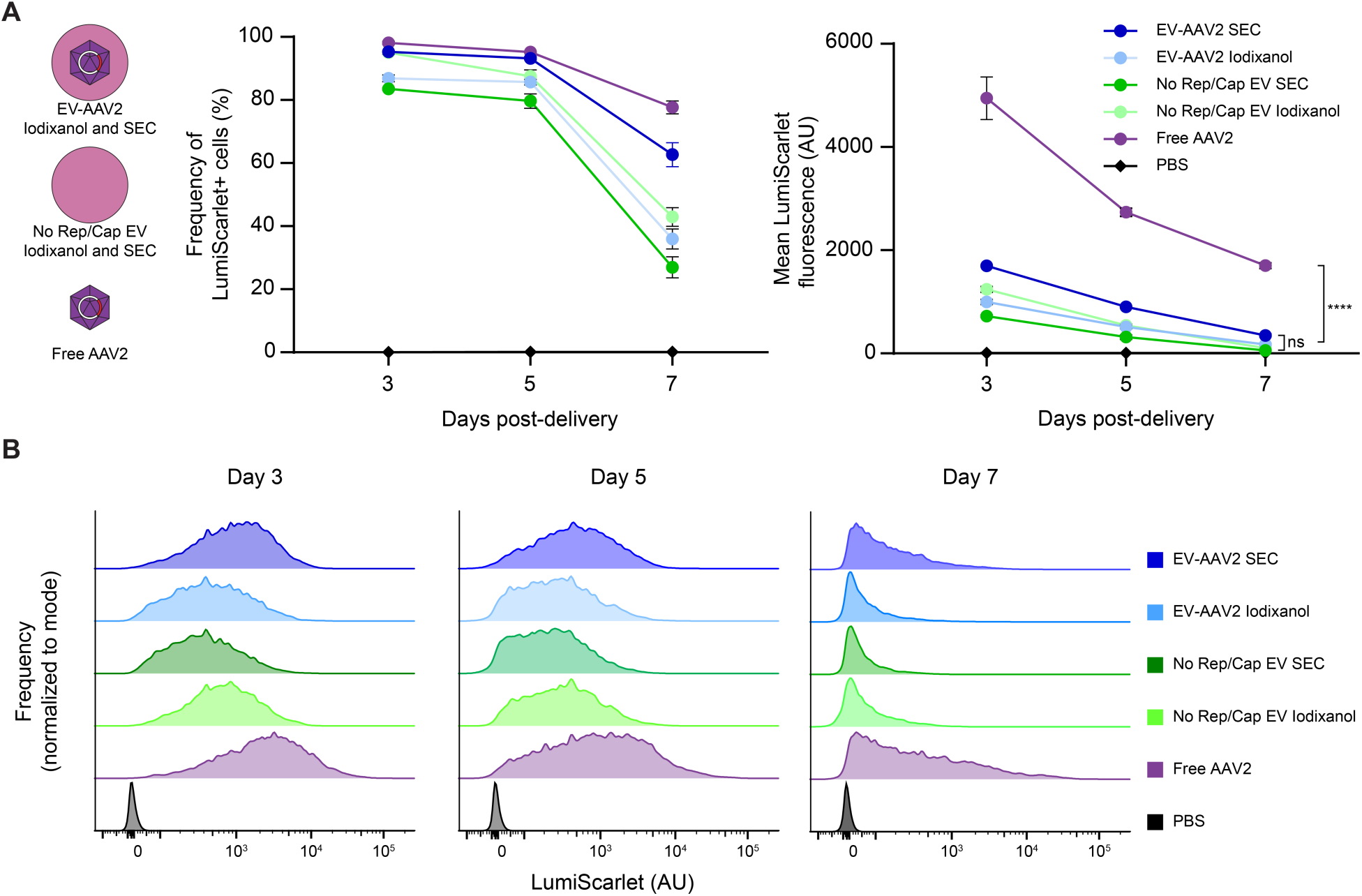
EV-AAV pseudotransduction complicates distinguishing true transduction. **(A)** EV-AAV-mediated protein delivery, mRNA and/or expression in HEK293FT recipient cells. Vector compositions and EV-AAV purification methods employed to separate EVs and free AAV are summarized in cartoons at left. Free AAV (which should confer minimal pseudotransduction) conferred significantly greater fluorescence than did EV-AAV controls. EV-AAV vectors were generally similar to the No Rep/Cap (protein delivery only) control at all time points post-delivery. **(B)** Representative histograms of LumiScarlet fluorescence from samples reported in (A). No subpopulations are apparent that can rigorously distinguish protein delivery from transduction. Experiments were performed in biological triplicate. Data shown are from one of two independent experiments (second experiment: **Figures S8**,**9**). Error bars indicate standard error of the mean. Multicomparison statistical analysis was performed using a two-way ANOVA test, followed by Tukey’s multiple comparison test to evaluate specific comparisons (\*\*\*\**p* < 0.0001). ns; not significant; PBS: phosphate-buffered saline; SEC: size exclusion chromatography.

For experiments involving lentiviral transduction of HEK293T and HEK293FT by Ultra-HEK, similar methods were used. Flow cytometry was performed on a Novocyte Penteon using the 488 nm laser for mNeonGreen (525/45 filter) and the 637 nm laser for DRAQ7 (725/40 filter). Approximately 3,000 live cells were collected per sample for analysis. Data were analyzed using FlowJo v10 (FlowJo, LLC). Briefly, cells were identified using an FSC-A vs SSC-A plot and gated for singlets using an FSC-A vs FSC-H plot. Live cells were identified using a DRAQ7 negative gate drawn on a non-DRAQ7-stained sample, and transduced cells were identified by drawing an mNeonGreen positive gate on a non-transduced sample.

### Transmission Electron Microscopy (TEM)

For transmission electron microscopy (TEM), samples were diluted in PBS sterilized with 0.1 μm bottle top filters (Corning 431475) to concentrations on the order of 10^8^ particles per μL as determined by Nanoparticle Tracking Analysis. Grids were plasma treated, and 3 µL of diluted sample were placed on the carbon side of the grid for 30 s. The grids were then briefly washed on 3 droplets of distilled water and placed on a droplet of stain solution (2% ammonium molybdate) for a few seconds. The staining solution was blotted off and the grid was placed on a second droplet of staining solution for 30 s. Finally, the excess staining solution was removed, and the grid was air-dried. Samples were observed with a JEOL 3200FS TEM at an acceleration voltage of 300 kV. A K2 camera was used for recording images.

### Western blots

For western blots comparing protein in cell lysates and protein in vesicles, a fixed number of vesicles and a fixed mass of cell lysate were loaded into each well: 4.5 × 10^8^ EVs/lane and 2 µg protein/lane, respectively. An established western blot protocol^48^ was followed with the following modifications. In most cases, the following reducing Laemmli composition was used to boil samples (60 mM Tris-HCl pH 6.8, 10% glycerol, 2% sodium dodecylsulphate, 100 mM dithiothreitol (DTT) and 0.01% bromophenol blue); in some cases, a nonreducing Laemmli composition (without DTT) was used, as previously reported^52^. After transfer, membranes were blocked whilst rocking for 1 h at room temperature in 5% milk in Tris-buffered saline with Tween (TBST) (pH: 7.6, 50 mm Tris, 150 mm NaCl, HCl to pH 7.6, 0.1% Tween 20). Primary antibody was added in 5% milk in TBST, rocking, for 1 h at room temperature and then washed three times with TBST for 5 min each. Secondary antibody in 5% milk in TBST was added at room temperature for 1 h or overnight at 4°C. Membranes were then washed three times with TBST for 5 min each. The membrane was incubated with Clarity Western ECL substrate (Bio-Rad, #1705061) and imaged on an Azure c280 running Azure cSeries Acquisition software v1.9.5.0606. Specific antibodies, antibody dilution, heating temperature, heating time and Laemmli composition for each antibody was used as previously reported^52^.

### Statistical analysis

Statistical tests employed are described in relevant figure legends. Unless otherwise stated, three independent technical replicates were analyzed per condition, and the mean fluorescence intensity of at least 10,000 live single cells were analyzed per sample. Each figure represents a single experiment that was repeated (biological replicate) at least twice. Unless otherwise indicated, error bars represent the standard error of the mean. GraphPad Prism 10 was used to analyze the data. Multicomparison statistical analysis was performed using a one-way ANOVA test, followed by Tukey’s multiple comparison (i.e., honestly significant difference, HSD) test to evaluate specific comparisons. Significance threshold: \**p* < 0.05, \*\**p* < 0.01, \*\*\**p* < 0.001, \*\*\*\**p* < 0.0001.

## RESULTS

### Distinguishing pseudotransduction from transduction by EV-AAV is challenging

We first sought to quantify the magnitude and dynamics of pseudotransduction for EV-AAV delivery to HEK293FTs. It is commonly expected that the fluorescence conferred to recipient cells by EV-mediated delivery of producer cell-derived protein and/or mRNA (i.e., pseudotransduction) will decrease over time due to protein degradation and/or cell division, while fluorescence originating from transduction (gene delivery and resulting transgene expression in recipient cells) will persist and increase to steady state. However, the dynamics of such contributions are poorly explored and are likely to be system specific. To characterize these phenomena in our model system, we utilized an AAV vector that constitutively expresses a bright fluorescent protein, LumiScarlet, and we generated both EV-AAV (i.e., heterogeneous preps expected to contain a mixture of EVs lacking any AAV particle and EVs associated with AAV particles) and free AAV preparations (**Figure S5**). As a control for pseudotransduction, we included conditions lacking Rep and Cap genes (“No Rep/Cap”). We sought to investigate both large and small EVs within our EV and EV-AAV preparations, as both subsets have been shown to incorporate and deliver producer cell-derived protein to recipient cells^28^. Therefore, a given EV-AAV or No Rep/Cap EV preparation contains both large and small EVs. We also investigated several EV purification methods, including iodixanol gradient purification^21,23,24^ and size exclusion chromatography^25^, reasoning that some fluorescent protein made by producer cells may be excreted into the media along with free AAVs and EVs, and different EV purification methods may remove these free proteins to different extents. Both purification methods yielded EVs with expected protein markers (**Figures S6A,B**), appropriate size distributions (**Figure S6C**), and morphology by TEM (**Figure S6D**). Purified vectors were dose-normalized (by genome count for EV-AAV and AAV with No Rep/Cap EVs matched to EV-AAV by particle count) and delivered to HEK293FT cells (which actively endocytose EVs). Fluorescence was monitored over 7 days by flow cytometry (**Figure 1, Figure S8**) and microscopy (**Figure S7, Figure S9**). In general, fluorescence attributable to EV-AAV transduction was not readily distinguished from fluorescence attributable to pseudotransduction (e.g., No Rep/Cap control). Free AAV2 conferred distinguishable transgene expression in one experiment, which was distinguished by quantifying the magnitude of fluorescence per cell (**Figure 1A, right**). Representative histograms of LumiScarlet fluorescence from each sample also demonstrate that fluorescence attributable to EV-AAV transduction was not readily distinguished from fluorescence attributable to protein delivery (**Figure 1B, Figure S8B**). We noted that LumiScarlet fluorescence shifts leftward in these histograms over the time course. Since this trend is observed across conditions, we hypothesize that cell division by these proliferative HEK293FT cells dilutes both AAV genomes and LumiScarlet protein. Our analyses were visually confirmed by microscopy (**Figure S7A-C, Figure S9A-C**), in which free AAV2 conferred the expected whole cell fluorescence associated with de novo transgene expression, while EV conditions exhibited puncta (which likely indicate the accumulation of multiple EVs in compartments such as endosomes) that are readily apparent at 3 days post-treatment and remain detectable at 7 days post-treatment. Transduction is apparent in images of EV-AAV and free AAV treated cells (but not No Rep/Cap EV treated cells) by day 7, which is identifiable by diffuse and bright fluorescence throughout the cells. Our analyses indicate that delivered protein is detectable (i.e., as puncta) in recipient cells for up to a week, such that the time scale for delivered protein depletion and degradation is not readably separated from the timescale of AAV episomal vector dilution (which may be particularly pronounced in these rapidly dividing recipient cells). Overall, protein delivery by EV-AAVs may lead to significant pseudotransduction and is challenging to distinguish from transduction regardless of EV purification method.

### A Cre recombinase-based dual reporter system enables discrimination between pseudotransduction and transduction

To address the challenge of pseudotransduction, we developed a novel Cre recombinase (Cre)-based system to enable quantitative and unambiguous evaluation of gene and protein delivery by EV-AAVs. We utilized a Cre-responsive AAV genome encoding a constitutive mCherry fluorescent protein that, upon recombination, is excised to instead drive expression of a green fluorescent protein.^47^ This dual-fluorescence system enables visualization of producer cell-derived protein and/or mRNA delivery (mCherry) and transduction (green) in Cre-expressing recipient cells. We initially investigated vector design in this system using AAV and No Rep/Cap crude lysates; only the former should include transduction-competent AAV particles (**Figure 2A**). Given a preceding Cre-responsive construct developed for a distinct purpose^47^, we generated several vectors to identify design choices that are useful for this application (**Figure 2B**). We explored changing the transduction reporter from eGFP to mNeonGreen, hypothesizing that a brighter fluorophore would help identify transduction. We also hypothesized that removing the nuclear localization sequences (NLS) from the transduction reporter protein may increase accumulation in the recipient cell, again facilitating detection. To evaluate these designs, we made Cre-expressing recipient HEK293FT cells by lentiviral transduction.

**Figure 2:**
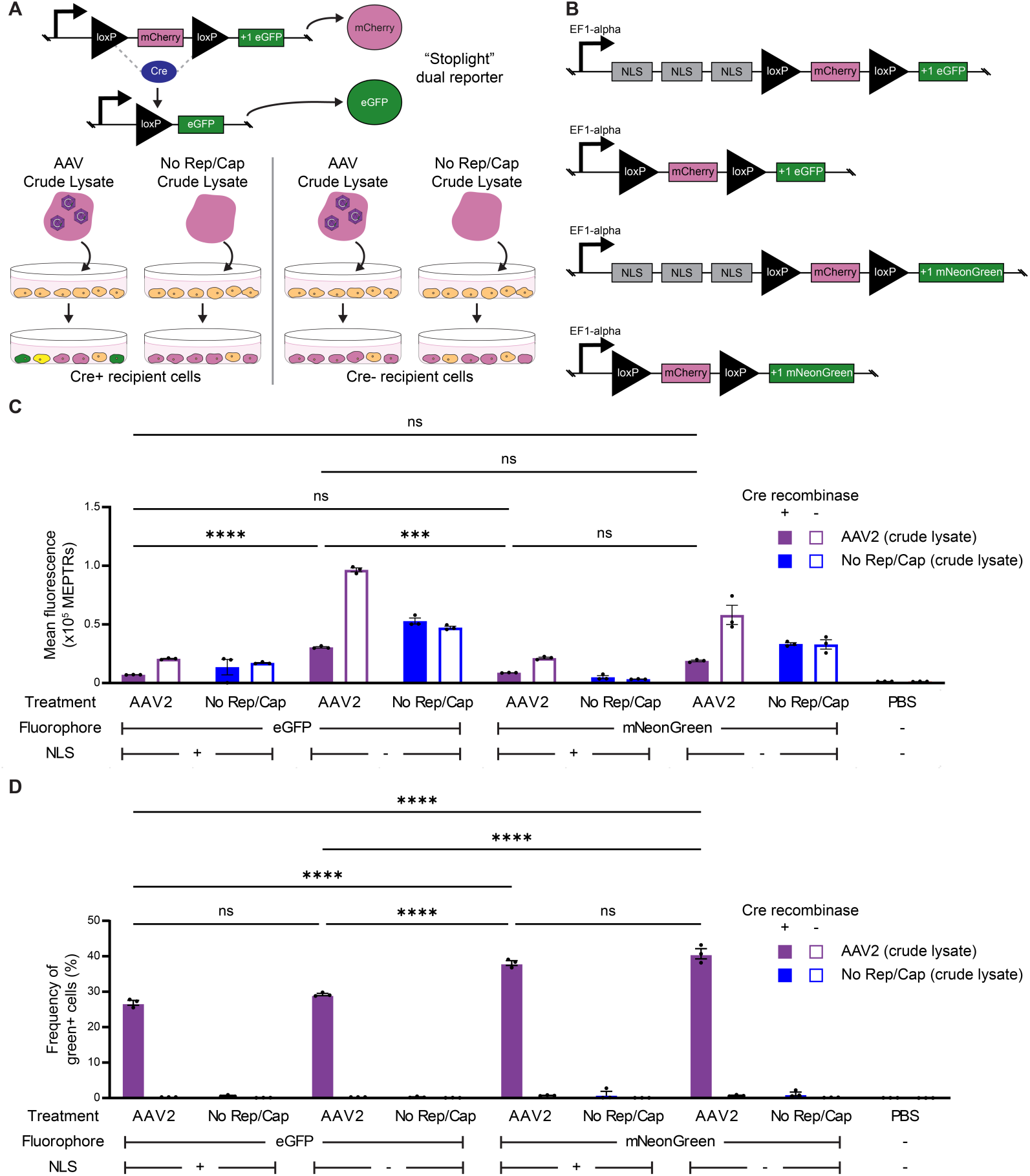
A Cre recombinase-based dual reporter system distinguishes mixed delivery and transduction. **(A)** Schematic depicting the expected function of the stoplight dual-reporter system in which Cre expressed in recipient cells acts upon incoming AAV genomes to switch them from expression of a red reporter to a green reporter. Free AAV mediates transduction only (green); EV-AAV mediates both mixed delivery (red) and transduction (green) and sometimes both (yellow); No Rep/Cap EVs can only mediate mixed delivery (red). **(B)** Cre-responsive plasmids evaluated in this experiment. **(C,D)** Mixed delivery (C) and transduction (D) conferred by various vector compositions. Note: some mCherry gene expression could occur in recipient cells prior to Cre-mediated recombination (or in the possible absence of recombination), such that (C) includes both mixed delivery and this ambiguous de novo gene expression. Samples were normalized to include 1e9 vector genomes per well (for AAV crude lysate conditions) or a volume-equivalent of the AAV2 crude lysate condition for No Rep/Cap crude lysate conditions (10^4^ recipient cells). Experiments were performed in biological triplicate. One of two independent experiments is shown (second experiment: **Figure S10**). Error bars indicate standard error of the mean. Multicomparison statistical analysis was performed using a two-way ANOVA test, followed by Tukey’s multiple comparison test to evaluate specific comparisons (\**p* < 0.05, \*\**p* < 0.01, \*\*\**p* < 0.001, \*\*\*\**p* < 0.0001). NLS: nuclear localization sequence; ns: not significant; PBS: phosphate-buffered saline.

We next evaluated the potential of each reporter system to ultimately detect pseudotransduction and transduction. Each variant was packaged as either AAV2 crude lysate—which should include a mix of free AAV2 and cell-derived fluorescent protein/mRNA—or a No Rep/Cap crude lysate control which should include only cell-derived fluorescent protein/mRNA (and no transduction-competent AAV particles). Each was delivered to Cre-expressing recipient cells or parental cell controls. Removal of the 3 x NLS generally increased mCherry protein delivery, as expected, since the NLS sequesters some protein in the nucleus of the producer cells (**Figure 2C, Figure S10A**). Greater mCherry fluorescence was observed in Cre-negative than Cre-positive recipient cells, in samples treated with AAV2 crude lysate (i.e., which are expected to include transduction competent AAV2 particles). A plausible explanation is that in recipient cells lacking Cre, more mCherry gene expression occurs than when Cre is present, because in the former case a larger fraction of the transgenes delivered by AAV transduction remain unrecombined. This suggests that the decrease in mCherry expression in Cre+ cells (vs. Cre- cells) is consistent with recombination of the transgene, switching from mCherry to green fluorescent protein production. The absence of this Cre-expression-dependence in samples lacking AAV2 particles (i.e., No Rep/Cap conditions) supports this interpretation. Most importantly, green fluorescent protein was detected only with AAV2 preps (i.e., crude lysates containing transduction-competent vectors) delivered to Cre-expressing recipient cells (**Figure 2D, Figure S10B**), validating this reporter system for unambiguously indicating transduction and de novo transgene expression in recipient cells. Overall, the mNeonGreen vectors conferred higher percentages of green positive live cells than did eGFP vectors, with the mNeonGreen vector without an NLS performing the best. On balance, going forward we selected the vector with mNeonGreen and the 3 x NLS, since it conferred more green-positivity in Cre-expressing recipient cells compared to eGFP vectors (i.e., greater signal), and it conferred less mCherry fluorescence compared to vectors without an NLS (i.e., lower background).

We next interrogated several aspects of our reporter system. First, we probed whether any leaky mNeonGreen expression occurred in producer cells, as spontaneous recombination has been reported in other Cre responsive plasmids^53,54^. If spontaneous recombination occurs, mNeonGreen protein may be expressed and loaded into EVs, as could mNeonGreen mRNA, resulting in mNeonGreen detection in recipient cells. To test this possibility we examined EV-AAV producer cells 3 d after transfection (**Figure S11**). We observed very low (but detectable) mNeonGreen expression in ∼11% of cells transfected with our AAV plasmids, and a similar frequency was observed in the Stoplight only (No Rep/Cap) condition, such that this leakiness was not impacted by AAV genome replication and packaging. In contrast, controls including Cre recombinase in EV-AAV producer cells yielded approximately 60% mNeonGreen positive cells and much higher magnitudes of mNeonGreen expression. Thus, despite some evidence of spontaneous recombination of our stoplight plasmid in EV producer cells, not much mNeonGreen protein is produced during the timescale of our transfections, which helps to explain why no mNeonGreen was detected in recipient cells except when both AAV and Cre were present (**Figure 2D, Figure S10B**). To further test this interpretation, we repeated our initial delivery experiment based upon crude lysate (**Figure 2D, Figure S10B**) using SEC purified EVs and purified free AAV (**Figure S12**) and again found that mNeonGreen transfer (protein and/or mRNA) by EV-AAV preps (and by No Rep/Cap control EVs) is minimal.

Finally, we explored the source of the mCherry signal observed in recipient cells. mCherry expression in recipient cells could result from delivery of producer cell derived protein by EVs, producer cell derived mRNA and subsequent translation in recipient cells, or de novo mCherry produced by the unrecombined transgene delivered by EV-AAVs. To investigate the possibility that producer cell derived mRNA contributes to mCherry signal, we utilized a protocol previously used to investigate mRNA delivery by engineered EVs^44^ (**Figure S13A**). Cells were treated with EV-AAVs, No Rep/Cap EVs, or Free AAV and fluorescence was assessed after 4 hours. As second group of cells were similarly treated with vectors for 4 h, but then 20 μg/mL of the translation inhibitor cycloheximide (CHX) was added and fluorescence was assessed after another 4 h had elapsed. Overall, there was minimal decrease in fluorescence in cells treated with cycloheximide and either EV-AAVs or No Rep/Cap EVs compared to DMSO-treated controls (**Figure S13B,C**). These results suggest that at least at early timepoints, most mCherry fluorescence observed in recipient cells treated with EVs or EV-AAVs originates from producer cell-derived mCherry protein. Nonetheless, to encompass the possibility that mCherry signal in recipient cells could derive from protein delivery, mRNA delivery, and/or AAV delivery (for conditions contain transduction-competent AAV particles), we henceforth conservatively describe these phenomena collectively as “mixed delivery”. We carried this validated reporter system forward to probe EV-AAV bioengineering questions.

### EV-AAV2 surface functionalization substantially increases mixed delivery but not transduction of HEK293FTs

We next investigated how EV functionalization strategies that have been reported to increase delivery of lentiviruses or EVs impact mixed delivery and transduction by EV-AAV. VSV-G is often used in pseudotyping lentiviruses^55,56^ and EVs^57,58^, and it increases the protein delivered by EVs.^58^ VSV-G binds to the LDL receptor (LDL-R) on the surface of recipient cells^30^, increasing particle uptake as a result. Within the low pH of the endosome, VSV-G undergoes conformational changes that enable it to promote fusion between the viral (or EV) membrane and the endosomal membrane to release the vector’s luminal contents into the cytosol of the recipient cell.^59^ To separate the impacts of enhancing uptake and mediating fusion, we also utilized Ultra-HEK, a commercial product (Syenex) that increases lentiviral transduction of HEK293-based cells when combined with a fusogen such as VSV-G. Ultra-HEK is described as interacting with proteins on the surface of recipient HEK293 cells to cause increased uptake of the particle (but not fusion). In our hands, packing lentiviral vectors with Ultra-HEK led to increased transduction (i.e., functional titer) for delivery to both HEK293T cells and HEK293FT cells (**Figure 3A–C**).

**Figure 3:**
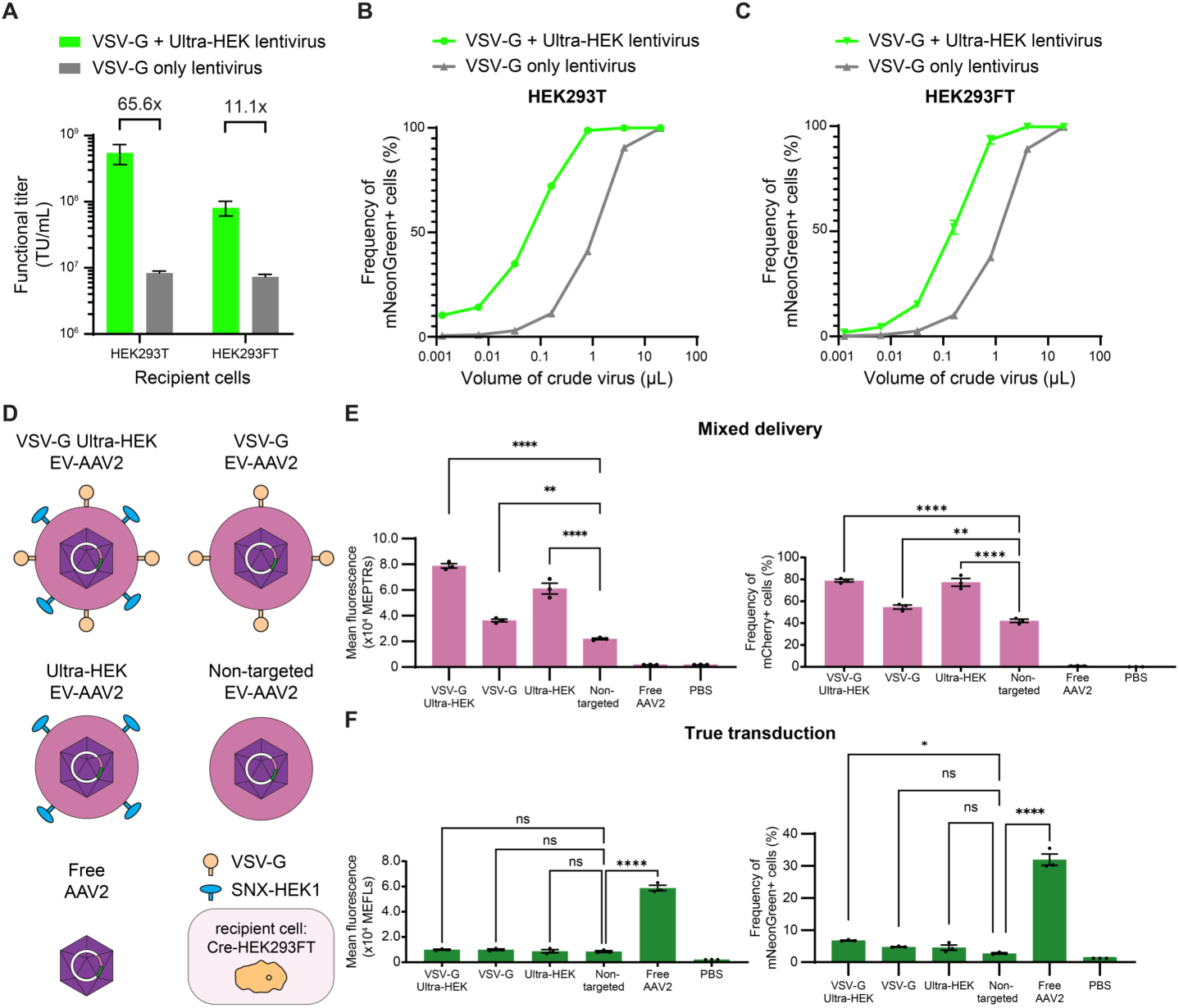
EV-AAV2 functionalization increases mixed delivery but not transduction of HEK293FTs. (A-C) Validation of Ultra-HEK (Syenex) for enhancing lentiviral delivery to HEK293T and HEK293FT cells. Relative fold-improvements in functional titer compared to a standard VSV-G-pseudotyped lentiviral vector are indicated. (**D**) Schematic depicting vectors used in (E,F) (Cre-HEK293FT recipient cells). (**E,F**) Mixed delivery quantified by mean fluorescence in MEPTRs (left) and frequency of mCherry positive cells (right) (E) and transduction quantified by mean fluorescence in MEFLs (left) and frequency of mNeonGreen positive cells (right) (F) to HEK293FT cells (5 days post-delivery). Samples were normalized to include 1e9 vector genomes per well for each condition (10^4^ recipient cells). VSV-G and Ultra-HEK-functionalized EV-AAV increase mixed delivery but only modestly increase transduction and only when combined. Experiments were performed in biological triplicate. One of two independent experiments is shown (second experiment: **Figure S15**). Error bars indicate standard error of the mean. Multicomparison statistical analysis was performed using a one-way ANOVA test, followed by Tukey’s multiple comparison test to evaluate specific comparisons (\**p* < 0.05, \*\**p* < 0.01, \*\*\**p* < 0.001, \*\*\*\**p* < 0.0001). PBS: phosphate-buffered saline.

**Figure 4:**
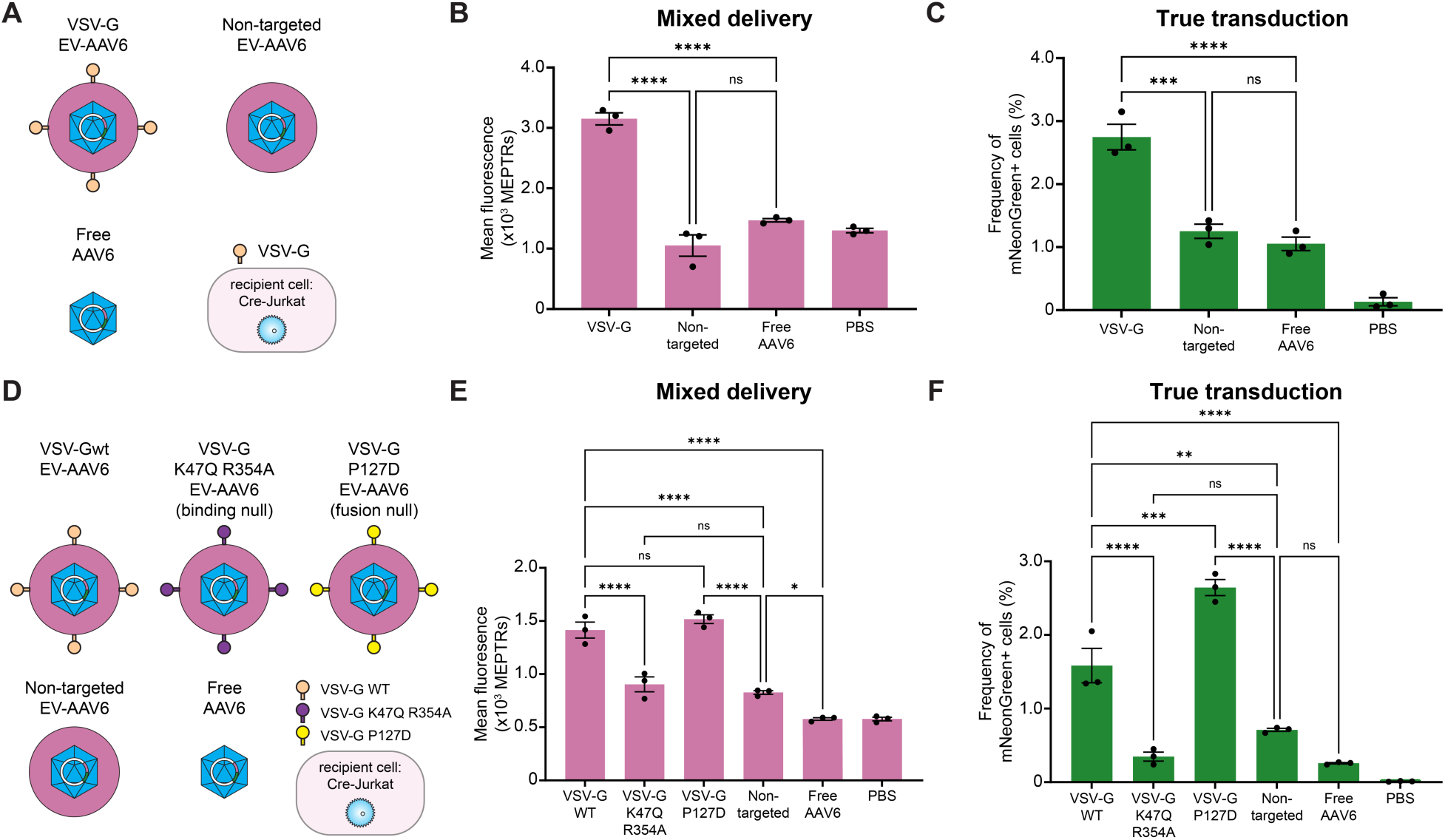
VSV-G functionalization of EV-AAV6 increases transduction and mixed delivery to Jurkat T cells by increasing uptake. **(A)** Schematic depicting vectors used in (B,C) (Cre-Jurkat T cell recipient cells). (**B,C**) Mixed delivery (B) and transduction (C) in Jurkat T cells (5 days post-delivery). Samples were normalized to include 1e9 vector genomes per well for each condition (10^4^ recipient cells). VSV-G functionalization significantly increases mixed delivery to and transduction of Jurkats. **(D)** Schematic depicting vectors used in (E,F) (Cre-Jurkat T cell recipient cells). (**E,F**) Mixed delivery (E) and transduction (F) in Jurkat T cells (5 days post-delivery). Samples were normalized to include 2e9 vector genomes per condition (10^4^ recipient cells). VSV-G functionalization significantly increases mixed delivery and transduction of Jurkats. Mixed delivery and transduction are increased only by enhanced binding of recipient cells (VSV-G WT and VSV-G P127D vectors). Experiments were performed in biological triplicate. Data shown are from one of two independent experiments (second experiments in **Figure S18**). Error bars indicate standard error of the mean. Multicomparison statistical analysis was performed using a one-way ANOVA test, followed by Tukey’s multiple comparison test to evaluate specific comparisons (\**p* < 0.05, \*\**p* < 0.01, \*\*\**p* < 0.001, \*\*\*\**p* < 0.0001). PBS: phosphate-buffered saline.

Using our reporter system and this menu of surface functionalization strategies (**Figure 3D, Figure S14A**), we revisited the question of how driving EV-AAV uptake (and/or fusion) impacts transduction efficiency. All functionalization strategies increased mixed delivery, as measured by either the magnitude mCherry fluorescence (**Figure 3E**, left and **Figure S14B,** left) or frequency of mCherry positive cells (**Figure 3E**, right and **Figure S14B,** right), likely due to increased particle uptake by recipient cells. Ultra-HEK combined with VSV-G functionalization increased mixed delivery to the greatest extent compared to non-targeted EV-AAV2. Since doses of EV-AAVs were normalized by genome copy, it is possible that equivalent genome doses comprise distinct numbers of EVs. To exclude the possibility that this could explain the mixed delivery trend, we plotted mixed delivery by EV dose (in particle count), and we found that this does not well explain our observations (**Figure S15**). Thus, differences in mixed delivery are best explained by the biological effects of our functionalization strategies. Interestingly, surface functionalization did not increase transduction to the same degree as it did protein delivery, with transduction measured by either magnitude of mNeonGreen fluorescence (**Figure 3F**, left and **Figure S14C,** left) or frequency of mNeonGreen positive cells (**Figure 3F**, right and **Figure S14C,** right). Only the dual functionalization strategy modestly increased the amount of transduction of HEKs over non-targeted vectors, and there was no evidence that fusion mediated by VSV-G enhanced or blocked EV-AAV transduction. Given this comparison, going forward, we chose metrics that are most aligned with the phenomenon being evaluated. We reasoned that degree of fluorescence conferred by mixed delivery is a continuous variable, and therefore mean fluorescence captures variation in that signal. Conversely, transduction is digital (transduced vs. not transduced), even if transduced cells vary in their degree of mNeonGreen expression; therefore, frequency of mNeonGreen+ cells is the metric most aligned with that measurement. Finally, we noted that the vast majority of cells that were truly transduced (mNeonGreen+) also were mCherry+ (i.e., mNeonGreen+/mCherry-cells were vanishingly rare), and this pattern was true across functionalization strategies (**Figure S16**). These findings led us to investigate whether these trends extend to other serotypes and recipient cells.

### EV-AAV6 surface functionalization increases mixed delivery and transduction of Jurkat T cells

We next explored EV-AAV delivery to a cell type with low spontaneous EV uptake. We chose to use human Jurkat T cells, which are somewhat recalcitrant to EV-based delivery due (at least in part) to low endocytosis.^28^ We transduced Jurkats with the Cre recombinase gene to make a recipient line compatible with our dual reporter system. We used AAV6 as it infects both primary T cells^60^ and Jurkats^61^. We focused on VSV-G functionalization which mediates lentiviral transduction of T cells via LDL-R binding and uptake (**Figure 4A, Figure S17A**).^62^ Similar to HEK293FT delivery (**Figure 3, Figure S14**), VSV-G functionalization increased mixed delivery to Jurkats compared to non-targeted EV-AAV6 (**Figure 4B, Figure S17B**). Surprisingly, and unlike in HEK293FT recipient cells, VSV-G functionalization also increased transduction of Jurkats (**Figure 4C, Figure S17C**).

This led us to investigate what mechanism conferred by VSV-G (uptake vs. fusion) contributed to increased transduction. We employed two VSV-G mutants to dissect these mechanisms (**Figure 4D, Figure S17D**). P127D retains the ability to bind to LDL-R but is deficient in membrane fusion.^49,63,64^ Conversely, mutations K47Q-R354A block LDL-R binding but still enable VSV-G to confer membrane fusion.^65–67^ Wild-type (WT) and P127D VSV-G functionalization resulted in more mixed delivery than did K47Q-R354A or non-targeted vectors (**Figure 4E, Figure S17E**). Interestingly, the K47Q-R354A VSV-G conferred less transduction than did the WT or P127D VSV-G vectors, indicating that binding to LDL-R on recipient cells is the dominant mechanism by which VSV-G increases transduction of Jurkat T cells (**Figure 4F, Figure S17F**). Notably, P127D VSV-G, which is fusion incompetent, conferred more transduction than did VSV-G WT vectors. Altogether, these results may indicate that for EV-AAV6 delivery to Jurkats, increased LDL-R binding increases particle uptake and transduction, while membrane fusion limits transduction.

We next investigated whether the trend of double positive cells we observed in our HEK delivery experiments (**Figure S16)** held for Jurkat delivery experiments (**Figure S18**). We utilized the mutant VSV-G data sets to explore this question. Surprisingly, and contrasting with our HEK delivery experiments, most truly transduced Jurkat cells (mNeonGreen+) were mCherry-(**Figure S18B,C).** This difference may be explained by the relatively low uptake (endocytosis) rate of EVs by Jurkats compared to HEKs, as evidenced by the low frequency of mixed delivery signal overall compared to HEKs. Additionally, the vectors with functionalization that improves uptake (VSV-Gwt and P127D) demonstrated more cells that were double positive than the uptake incompetent vectors (VSV-G K47Q 354A and non-targeted). Overall, this analysis illustrates how the stoplight vector enables one to probe multiple aspects of delivery and compare across cell types.

**Recipient cell type but not capsid serotype drives the impact of functionalization on EV-AAV delivery** To reconcile the distinct patterns observed in our HEK293FT and Jurkat delivery experiments, we investigated whether differences are driven primarily by recipient cell type or AAV serotype. We chose to investigate transducing HEK293FT cells with AAV6 since AAV6 is much less efficient at infecting HEK293FTs compared to AAV2.^61^ This difference may be due to differences in receptor density on the surface of HEKs. AAV2 primarily uses heparan sulfate proteoglycan to mediate binding and uptake by recipient cells.^68^ In contrast, AAV6 primarily uses sialic acid as its receptor^68^, which may be less abundant or accessible on HEK293 cells compared to heparan sulfate proteoglycan. Therefore, we hypothesized that increasing uptake of EV-AAV6 (e.g., via VSV-G functionalization) would increase transduction compared to both non-targeted EV-AAV6 or free AAV6 (**Figure 5A, Figure S19A**). VSV-G functionalization increased mixed delivery compared to non-targeted EV-AAV6, as expected (**Figure 5B, Figure S19B**). However, VSV-G mediated uptake (and fusion) did not increase transduction by EV-AAV6 (**Figure 5C, Figure S19C**), as was seen for EV-AAV2 (**Figure 3F, Figure S14D**). Thus, increasing uptake and membrane fusion in a cell type that is permissible to EV delivery, such as HEK293FTs, did not increase transduction, regardless of AAV capsid choice. With the set of conditions evaluated, EV surface functionalization strategies conferred cell type-dependent but not capsid-dependent effects on transduction by EV-AAVs.

**Figure 5:**
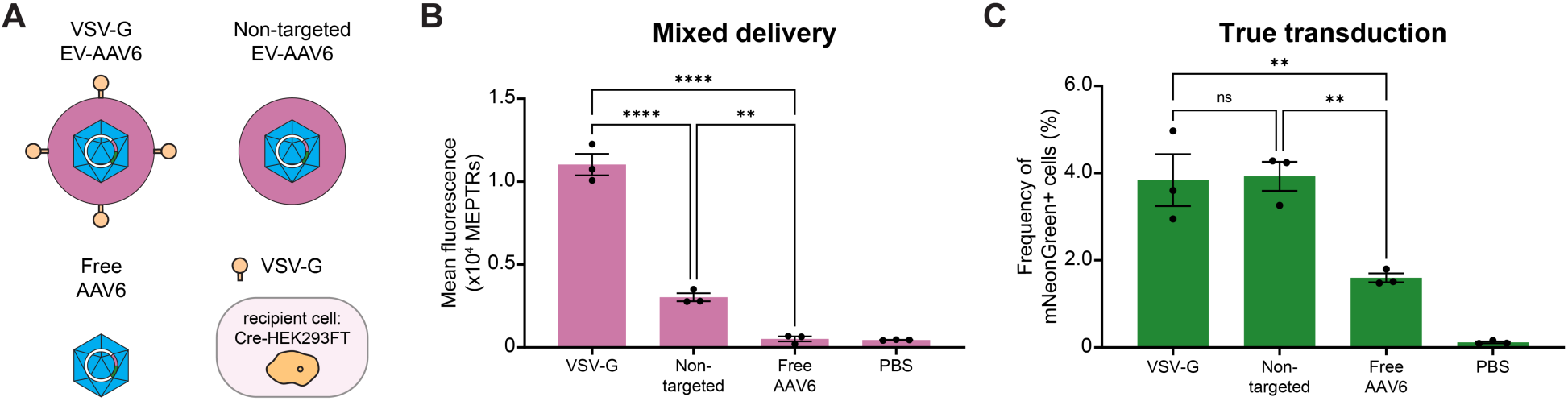
Determinants of EV-AAV delivery to HEK293FT cells apply across two AAV serotypes. **(A)** Schematic depicting vectors used in (B,C) (Cre-HEK293FT recipient cells). (**B,C**) Mixed delivery (B) and transduction (C) to HEK29FT cells (5 days post-delivery). Samples were normalized to include 1e9 vector genomes per well for each condition (10^4^ recipient cells). VSV-G functionalization enhances mixed delivery but not transduction by EV-AAV6, concordant with observations made using EV-AAV2. Experiments were performed in biological triplicate. One of two independent experiments is shown (second experiment: **Figure S20**). Error bars indicate standard error of the mean. Multicomparison statistical analysis was performed using a one-way ANOVA test, followed by Tukey’s multiple comparison test to evaluate specific comparisons (\**p* < 0.05, \*\**p* < 0.01, \*\*\**p* < 0.001, \*\*\*\**p* < 0.0001). PBS: phosphate-buffered saline.

## DISCUSSION

Engineered EVs are promising delivery vehicles for AAV vectors, enabling tissue-specific targeting and neutralizing antibody evasion. However, analyzing and bioengineering such vectors is made challenging by the difficulty associated with distinguishing pseudotransduction from transduction. To address this need, in this study we developed and validated a new assay that unambiguously distinguishes transduction from mixed delivery (including pseudotransduction), while enabling quantification of both phenomena. This assay helped to generate new insights that inform both interpretation of past studies and identify opportunities for future investigation.

Our observations provide new insights that help to interpret apparent contradictions in the EV-AAV literature. A prior report indicated that pseudotyping EV-AAV2 with VSV-G decreased the transduction of HEK293 cells compared to non-targeted EV-AAV2.^69^ Notably, this analysis employed a luminescence-based plate reader assay, and it may have been confounded by pseudotransduction artifacts. Our results suggest that VSV-G neither improves, nor hinders EV-AAV2 transduction of HEKs (**Figure 3**, **Figure S14**). Interestingly, our results suggest that VSV-G does improve transduction of EV-AAV6 in Jurkats by increasing particle uptake (**Figure 4, Figure S17**). Overall, our work suggests that the effects of surface functionalization on EV-AAV transduction is cell type dependent.

Some previous studies report that EV-AAVs are more potent than AAV vectors for *in vivo* delivery (on a per genome basis)^14–16,22^, but other work focused on delivery to neuronal and primary astrocyte cell cultures seemingly contradicts this finding^70^. It is possible that improved *in vivo* efficacy of EV-AAV particles is primarily attributable not to the enhanced interaction between the AAV and the target cells, but rather to shielding of AAVs from neutralizing antibodies and/or biodistribution effects such as increased circulation time or improved delivery through tissue barriers. Our observations add another layer of possible explanation, given our finding that target cell type and capsid choice may impact the efficacy of EV-AAV compared to free AAV. Free AAV2 was significantly more potent for transduction in HEK293FTs compared to non-targeted EV-AAV2 (**Figure 3, Figure S14**). However, non-targeted EV-AAV6 outperformed (**Figure 5)** or was equivalent to (**Figure S19**) free AAV6 in transduction in HEK293FTs. This suggests that capsid choice may impact the relative transduction efficacy of EV-AAV and free AAV. EV-AAV6 was not statistically significantly different from free AAV6 in transduction of Jurkat T cells (**Figure 4, Figure S17**), further suggesting that the capsid choice may impact efficacy comparisons between EV-AAV and free AAV. Overall, our findings indicate that the impact of EV-encapsulation of AAV confers case-specific effects that should be evaluated for each application, and which may drive application-specific bioengineering.

This study focused on identifying high level determinants of functional delivery, yet it also identified opportunities for further subsequent mechanistic investigations. Pseudotransduction was more evident in HEK293FT recipient cells, which actively endocytose EVs, compared to Jurkat recipient cells, which exhibit less endocytosis. Increasing EV uptake via surface functionalization also increased pseudotransduction in both cell types. Increasing the uptake of EV-AAV to HEK293FT recipient cells, both with or without promoting membrane fusion, increased mixed delivery, but this did not substantially increase transduction for either EV-AAV2 or EV-AAV6. However, VSV-G functionalization of EV-AAV6 increased both mixed delivery and transduction of Jurkat T cells, and this effect was driven primarily by increased uptake rather than fusion; indeed, blocking VSV-G-mediated fusion via the P127D mutant enhanced transduction. The basis for these differences is not known, but a possible explanation is that AAV requires processing in the endosome—including capsid cleavage by endosomal cathepsins^31^ and acidification-mediated autoproteolytic processing^32^—to expose a phospholipase domain promoting virion escape and subsequent nuclear trafficking^33,34^. In prior work, microinjection of AAV2 particles directly into the cytosol resulted in no nuclear trafficking and subsequent transduction^71^, suggesting that endosomal processing of the AAV capsid is required for successful transduction. This requirement is well-characterized for free AAV2 and may extend to other serotypes.

Combining these observations enables one to propose potential explanatory mechanisms as to how different EV surface functionalizations impact transduction. First, it is possible that VSV-G confers improved EV-AAV6 uptake by Jurkats, overcoming one barrier to delivery, but VSVG-mediated membrane fusion circumvents the endosomal processing of the AAV6 capsid that is required to complete transduction. Thus, the VSV-G P127D mutant balances these effects to benefit delivery to T cells. It would be informative to probe how each step in capsid processing is impacted by encapsulation in EVs, and how such processing steps may differentially impact AAV particles on the EV surface versus the EV interior. In addition, directly tracking internalization of both receptors and particles could help to test the mechanisms offered as speculative explanations here. A second, non-exclusive hypothesis for explaining improved transduction of Jurkats by EVs functionalized with the VSV-G P127D mutant is that this vector confers enhanced EV uptake compared to WT VSV-G, as evidenced by increased mixed delivery signal (**Fig 4E** and **Fig S17E**). No such mechanism is yet known, but it is possible that the P127D mutant is expressed at higher levels on EVs, is internalized more rapidly, or drives different trafficking after uptake. A final non-exclusive hypothesis involves innate immune activation. It is possible that WT VSV-G induces membrane fusion to prompt or enhance innate immune responses such as interferon production, which dampen transgene expression in cells transduced by AAV^72^. Elucidating such contributors to successful delivery (e.g., transgene expression magnitude as a function of vgDNA dose) could build upon these observations to improve EV-AAV engineering.

It will be interesting to extend these studies to the many AAV capsid variants that exist, including natural serotypes as well as novel forms developed through directed evolution and protein engineering^73^, and to a variety of recipient cell types including those which endocytose EVs more readily than do HEK293FTs^74^. This work could also enable new AAV bioengineering campaigns specific to enhancing EV-AAV delivery, such as exploring mutations that destabilize the capsid structure to enhance transduction after EV fusion and delivery of the AAV core to the cytosol, bypassing endosomal processing.

There exist a number of opportunities for extending the approaches and methods reported here to study aspects of EV biology beyond those probed in this investigation. First and foremost, there exist many different methods for producing and purifying EV-AAV particles, and choice of method is likely to affect the biology, composition, and function of the resulting particles. Our EV harvesting method utilized differential centrifugation to remove cells (300 g x 10 min) and cell debris/apoptotic bodies (2000 g x 20 min), followed by tangential flow filtration to concentrate the supernatant. This choice intentionally includes a mixture of EVs in our samples (our EV characterizations by nanoparticle tracking analysis, western blots, and TEM (**Figure S6**) demonstrate the purity of our preparations). Although the MISEV2023 guidelines for EV research indicate that EV preparations pelleted at different centrifugation speeds (i.e., “small” and “large” EVs) contain EVs with overlapping size profiles^75^, and EVs are heterogenous in size and associated contents regardless of purification method^76,77^, it would nonetheless be interesting to employ our stoplight assay to evaluate whether EV-AAV fractions that pellet at different speeds, or which are isolated with different purification methods, confer different levels of pseudotransduction and transduction. Our EV-AAV preparations certainly contain both EVs that lack AAV and EVs that are associated with AAV.^17^ If methods to isolate EVs containing AAVs were available, and to segregate by EV-surface versus EV-interior loading of AAV, it would be particularly interesting to analyze how these subsets contribute to transduction and pseudotransduction. It would also be interesting to study how EV-AAV delivery phenomena change over a broader dose range. In our studies, the MOI as defined by transducing units per cell is rather low, such that we were far from the saturating regime of EV-AAV transduction. Accessing that higher MOI regime could be made feasible through improvements such as recently reported methods for enhancing AAV loading into EVs.^38^ Each of these represents an opportunity for leveraging our stoplight method to improve understanding of EV-AAV mediated delivery.

Our analysis provides an opportunity to speculate as to how pseudotransduction may, or may not, influence other investigations of EV-AAV. Many *in vitro* and *in vivo* studies of EV-AAVs have been performed, delivering transgenes including luciferases, fluorescent proteins, and therapeutic genes. A potential factor in detecting pseudotransduction by EV-AAVs is the half-life of the transgenic protein in recipient cells. EV-AAV preparations encoding firefly luciferase but lacking the capsid protein (equivalent to our No Rep/Cap EV samples) have previously been compared to EV-AAV-FLuc (with the capsid protein) in transduction of HELA cells^23^. Capsid-less EV-AAV-FLuc conferred low FLuc activity (2-fold above background levels) whereas EV-AAV conferred 9-fold higher FLuc activity compared with the capsid-less EV-AAV genomes. Since this suggests a lower degree of pseudotransduction, it is possible that compared to FLuc, fluorescent proteins may resist degradation in the endosomal lysosomal compartment (which is distinct from half-life measured in the cytosol). Thus, choice of reporter transgene may mitigate, but not necessarily avoid, pseudotransduction artifacts, at least on the relatively short time scales of *in vitro* delivery experiments using rapidly dividing cells (where dilution of AAV genomes constrains the time one can wait post-delivery to analyze transduction). Our stoplight system may be particularly enabling for such experiments.

An exciting prospect is extending this assay for *in vivo* EV-AAV delivery analyses. The stoplight vector inspiring our approach was originally employed to detect Cre expression *in vivo*^47^. Given that transgenic animals can be engineered to express Cre in a tissue-restricted or general fashion^78^, our assay could be employed to unambiguously identify EV-AAV transduction *in vivo*. Such a capability could be particularly useful for directly comparing AAV and EV-AAV-mediated delivery, and for enabling comparative evaluation of bioengineering approaches, *in vivo* multiplexed screening, and other efforts for which pseudotransduction is misleading or reduces signal-to-noise to preclude feasibility. Because most *in vivo* studies of EV-AAV wait days to weeks after EV-AAV treatment to analyze transduction, pseudotransduction artifacts may have diminished by the time of analysis, but that is unproven and merits examination, especially for slow-dividing recipient cells. This stoplight tool could probe whether pseudotransduction is indeed negligible, and even if it is, this tool could enable faster, unambiguous evaluation of *in vivo* transduction. We expect that the methods and findings presented here can help advance the understanding and use of EV-AAV vectors for diverse gene delivery applications.

## AUTHOR CONTRIBUTIONS

J.D.B., D.M.S., N.P.K., D.T.E., and J.N.L. conceived the initial project. J.D.B. performed the experiments. D.M.S. and H.I.E. performed the Ultra-HEK lentiviral vector experiment. J.D.B. and J.N.L. planned and analyzed experiments. J.D.B and J.N.L. wrote the manuscript. J.N.L. supervised the work.

## ACKNOWLEDGEMENTS

J.D.B. was supported by the National Institutes of Health Training Grant (T32GM008449) through Northwestern University’s Biotechnology Training Program. J.D.B was supported by the National Science Foundation Graduate Research Fellowship under grant number (DGE-1842165). This project was partially supported by an award from the McCormick Catalyst Fund (J.N.L.) and by a Cornew Innovation Award from the Chemistry of Life Processes Institute at Northwestern University (J.N.L.). Flow Cytometry was performed in the Northwestern University Flow Cytometry Core Facility supported by Cancer Center Support Grant (NCI CA060553) and in the Single Cell Genomics Facility at Northwestern University (RRID:SCR_026652), graciously supported by the Department of Neurobiology, the Department of Molecular Biosciences, and the NU Office for Research. This work made use of the BioCryo Core (RRID:SCR_021288) of Northwestern University’s NUANCE Center, which has received support from the SHyNE Resource (NSF ECCS-2025633), the IIN, and Northwestern’s MRSEC program (NSF DMR-2308691). Biological analysis] was performed in the Analytical bioNanoTechnology Equipment Core Facility of the Center for Regenerative Nanomedicine at Northwestern University. ANTEC receives partial support from the Soft and Hybrid Nanotechnology Experimental (SHyNE) Resource (NSF ECCS-2025633) and Feinberg School of Medicine, Northwestern University. This work was supported by the Northwestern University Sanger Sequencing Facility. Any opinion, findings and conclusions or recommendations expressed in this material are those of the authors and do not necessarily reflect the views of the National Science Foundation. The authors kindly thank the Leonard lab members for useful discussions throughout the planning, experimental, analysis and writing phases of this project. The authors would like to thank Dmitriy Bobrovnikov and Roxi Mitrut for discussions of Cre-responsive AAV vectors.

