## Supplementary Information for "Distinguishing Pseudotransduction and True Transduction Enables Characterization and Bioengineering of Extracellular Vesicle-Adeno-Associated Virus Vectors"

**Supplementary Information**
**for**

### Distinguishing Pseudotransduction and True Transduction Enables Characterization and Bioengineering of Extracellular Vesicle-Adeno-Associated Virus Vectors

Jonathan D. Boucher[1,2,3], Devin M. Stranford[1,3,4], Hailey I. Edelstein[1,3,4], Danielle Tullman-Ercek[1-3,5], Neha P. Kamat[3,5,6], Joshua N. Leonard[1-5,7*]

[1] Department of Chemical and Biological Engineering, Northwestern University, Evanston, Illinois 60208, United States
[2] Interdisciplinary Biological Sciences Program, Northwestern University, Evanston, Illinois 60208, United States
[3] Center for Synthetic Biology, Northwestern University, Evanston, Illinois 60208, United States
[4] Syenex Inc, Evanston, Illinois 60201, United States
[5] Chemistry of Life Processes Institute, Northwestern University, Evanston, Illinois 60208, United States
[6] Department of Biomedical Engineering, Northwestern University, Evanston, IL, 60208 USA
[7] Member, Robert H. Lurie Comprehensive Cancer Center, Northwestern University, Evanston, Illinois 60208, United States

**<u>Contents</u>**

**Supplementary Figures 1-19**

#### SUPPLEMENTARY FIGURES

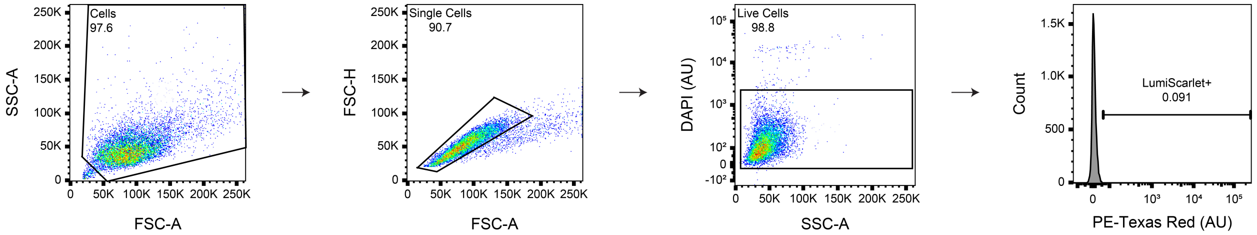


**Supplementary Figure 1: Conventional flow cytometry gating scheme for HEK293FTs.** The plots show a sample of cells incubated with PBS (negative control for vector transduction). In the gating procedure, cells were identified based on the FSC-A vs SSC-A profile. From this population, single cells were identified based on the FSC-A vs FSC-H profile. Cells were stained with 3 μM DAPI, and live cells were identified on the SSC-A vs DAPI-A profile. The LumiScarlet+ cell population was defined as all single live cells with a greater PE-Texas Red signal than the top 0.1% of the sample of PBS incubated cells. For experiments with Cre-responsive AAV vectors, gene delivery was defined as all single live cells with a greater FITC signal than the top 0.1% of the sample of PBS incubated cells. These % positive gates were drawn such that they did not encompass more than 0.1% of this non-fluorescent population of cells averaged across three replicates.

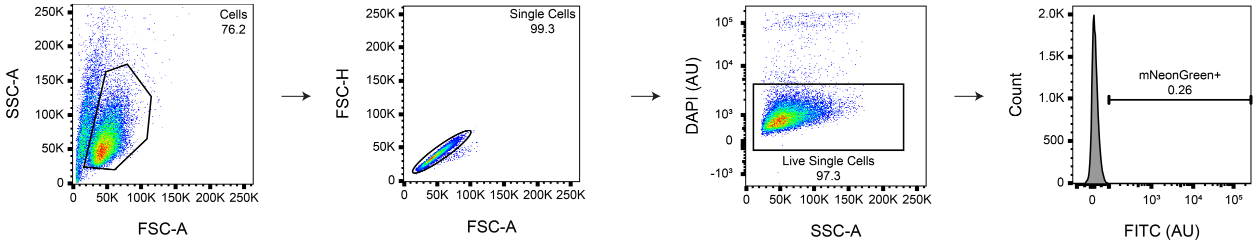


**Supplementary Figure 2: Conventional flow cytometry gating scheme for Jurkat T Cells.** The plots show a sample of cells incubated with PBS (negative control for vector transduction). In the gating procedure, cells were identified based on the FSC-A vs SSC-A profile. From this population, single cells were identified based on the FSC-A vs FSC-H profile. Cells were stained with 3 μM DAPI, and live cells were identified on the SSC-A vs DAPI-A profile. Gene delivery was defined as all single live cells with a greater FITC signal than the top 0.1% of the sample of PBS incubated cells. These % positive gates were drawn such that they did not encompass more than 0.1% of this non-fluorescent population of cells averaged across three replicates.

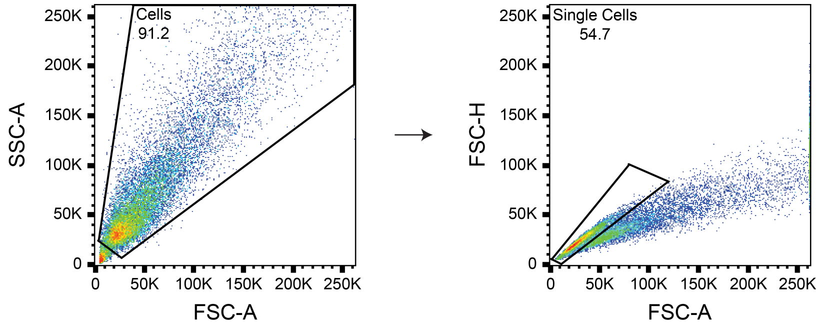


**Supplementary Figure 3: Conventional flow cytometry gating scheme for Freestyle 293-F cells.** The plots show a sample of cells transfected with pcDNA (negative control for producer cell analysis). Cells were fixed with 4% paraformaldehyde before flow cytometry. In the gating procedure, cells were identified based on the FSC-A vs SSC-A profile. From this population, single cells were identified based on the FSC-A vs FSC-H profile.

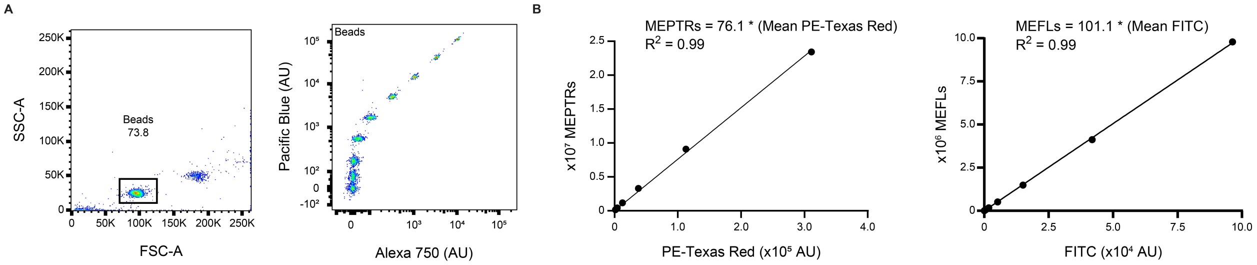


**Supplementary Figure 4**: **Calibration procedure used to convert fluorescence to absolute units. (A)** UltraRainbow Calibration Particles have nine fluorescent bead populations. Beads were identified based on the FSC-A vs. SSC-A profile. The beads are fluorescent in the majority of fluorescent channels on the flow cytometer. For each experiment, two channels were used to identify the bead populations. **(B)** The mean intensity of each population in the PE Texas Red (mCherry, left) channel in arbitrary units was recorded and plotted against the manufacturer-supplied number of fluorophores on the beads for each population (MEPTRs). The mean intensity of each population in the FITC (mNeonGreen, right) channel in arbitrary units was recorded and plotted against the manufacturer-supplied number of fluorophores on the beads for each population (MEFLs). To generate the calibration curve, linear regressions were performed with the constraint that the y-intercept equals zero. In each experiment, except those in **Figure 1** (LumiScarlet experiments were retained in arbitrary units), MFI were converted to MEPTRs or MEFLs by using the multiplier on MFI obtained from the regression.

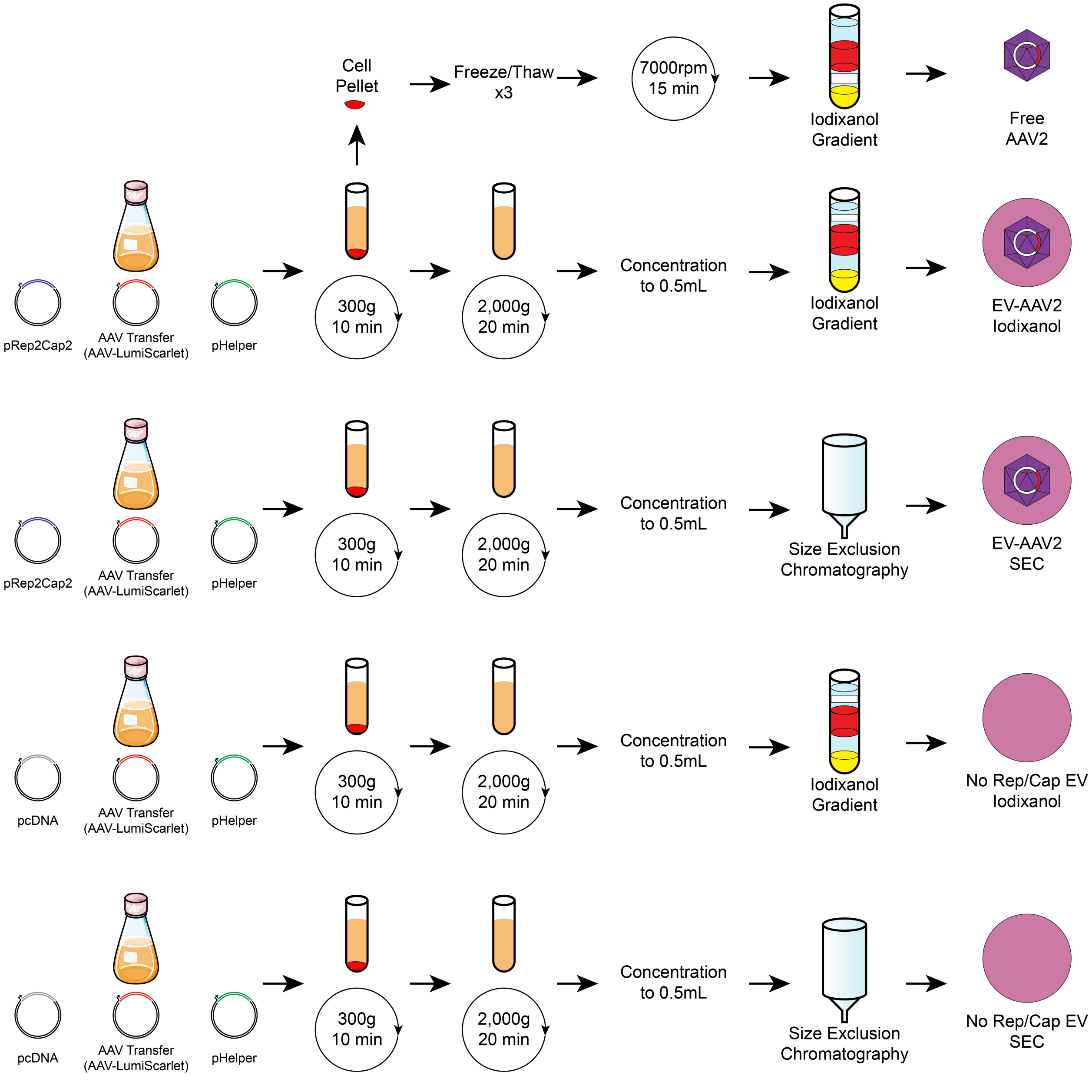


**Supplemental Figure 5: Production methods for samples used in LumiScarlet time course.** EV-AAVs were made by transfection of the standard AAV plasmids. No Rep/Cap EVs were made by transfection of the pHelper, pAAV-LumiScarlet, and pcDNA replacing the pRep2Cap2 plasmid. Two centrifugations were performed to pellet cells and cellular debris. The supernatant was concentrated to 0.5 mL final volume by tangential flow filtration followed by spin filtration. Samples were treated with benzonase and purified by iodixanol gradient centrifugation or size exclusion chromatography, as indicated. Free AAV was produced from cells transfected with standard AAV plasmids. The cell pellet was resuspended in AAV lysis buffer and underwent 3 freeze/thaws in an ethanol and dry ice bath. The mixture was spun to remove cellular debris, and the supernatant contained AAV crude lysate. AAV crude lysate was purified by iodixanol gradient centrifugation to remove cellular contaminants. bath_flask icon by Servier https://smart.servier.com/ is licensed under CC-BY 3.0 Unported https://creativecommons.org/licenses/by/3.0/

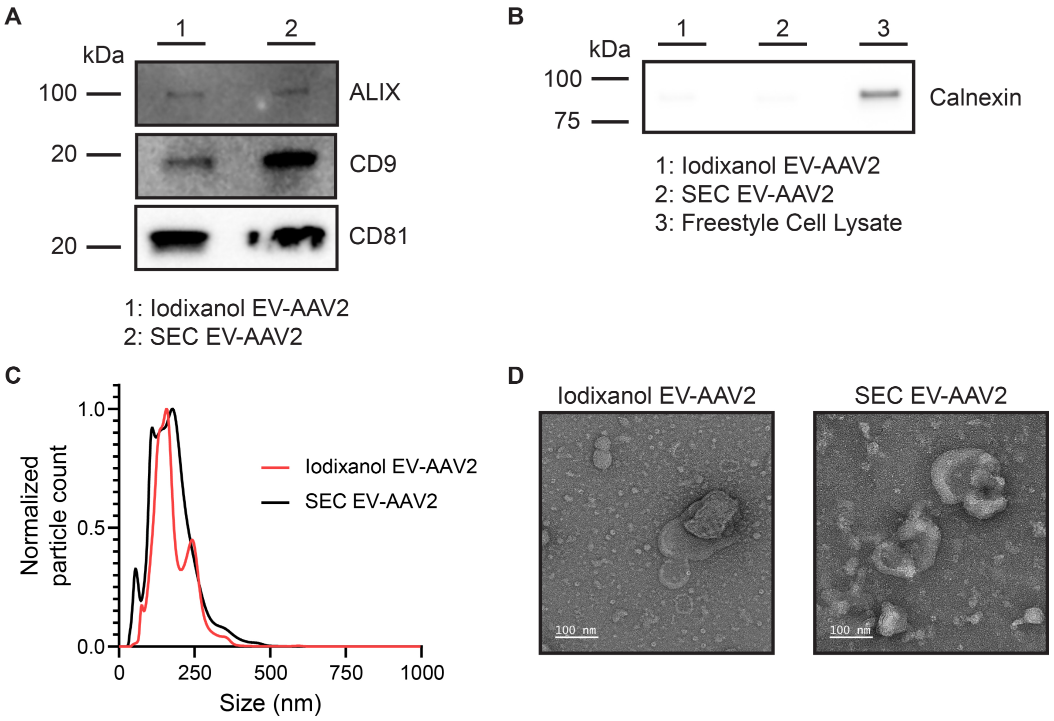


**Supplementary Figure 6: Standard extracellular vesicle quality control analyses. (A,B)** Western blots yielding expected patterns of common EV markers in purified vesicles versus producer cells. EVs contain expected markers, and calnexin is only present in cell lysate. **(C)** Representative histogram of nanoparticle tracking analysis of EV-AAV2s derived from Freestyle 293F cells and purified by iodixanol gradient centrifugation (red) and size exclusion chromatography (black), normalized to the mode in each population. **(D)** EV-AAVs morphology analysis by transmission electron microscopy. EV-AAV2 show standard cup-shaped morphology expected for this method. Left: Iodixanol gradient centrifugation purified EV-AAV2. Right: Size exclusion chromatography purified EV-AAV2. Scale bar: 100 nm.

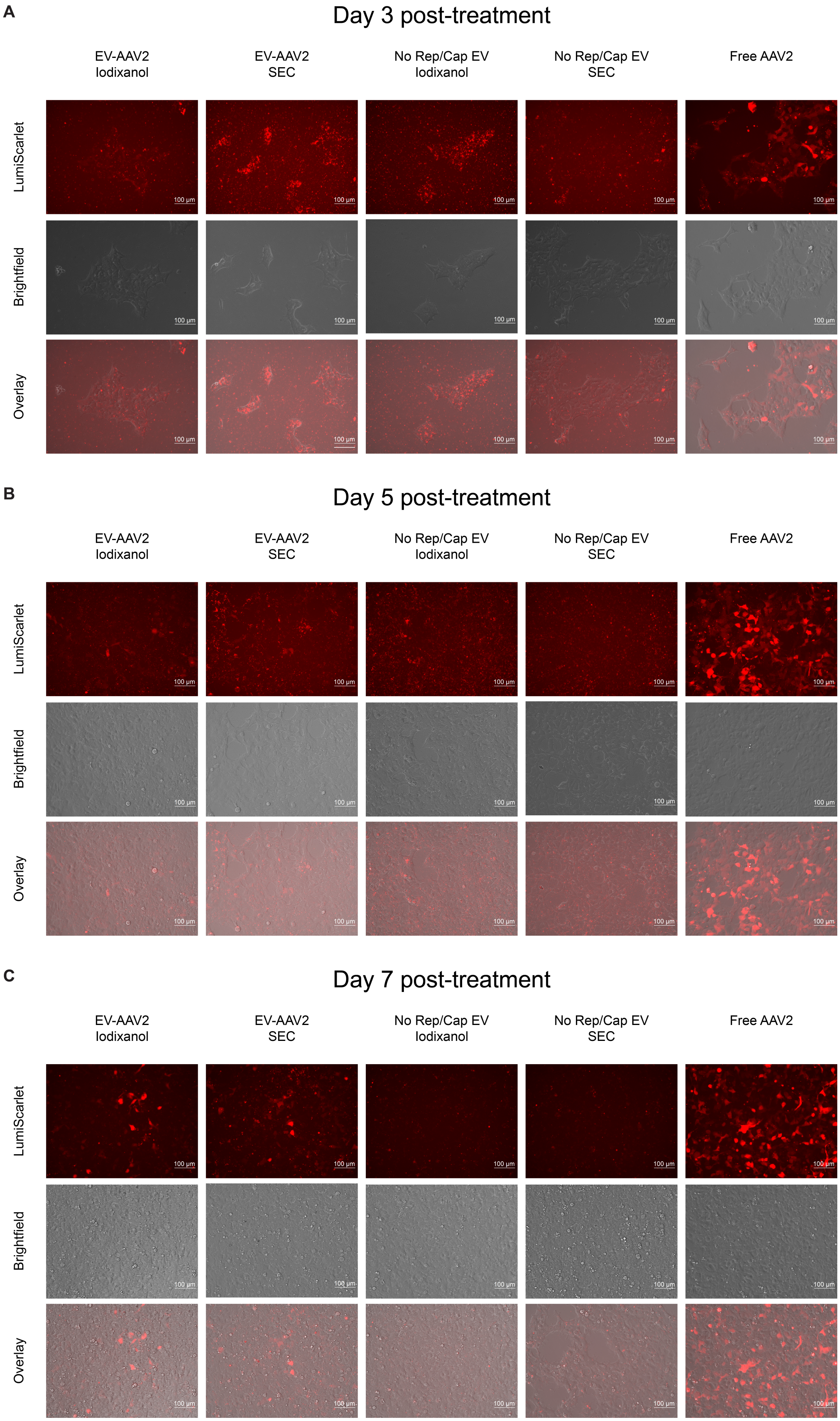

A
Day 3 post-treatment
B
Day 5 post-treatment
C
Day 7 post-treatment
EV-AAV2 Iodixanol
EV-AAV2 SEC
No Rep/Cap EV Iodixanol
No Rep/Cap EV SEC
Free AAV2
LumiScarlet
Brightfield
Overlay
100 µm

**Supplementary Figure 7: Illustrative micrographs of EV-AAV and AAV pseudotransduction and transduction**. **(A-C)** Microscopy images of HEK293FT cells treated with various vectors, Scale bars: 100 µm. Whole cell fluorescence is apparent for free AAV2 conditions, and puncta are evident in EV-treated conditions. Images were taken of cells analyzed in **Figure 1.**

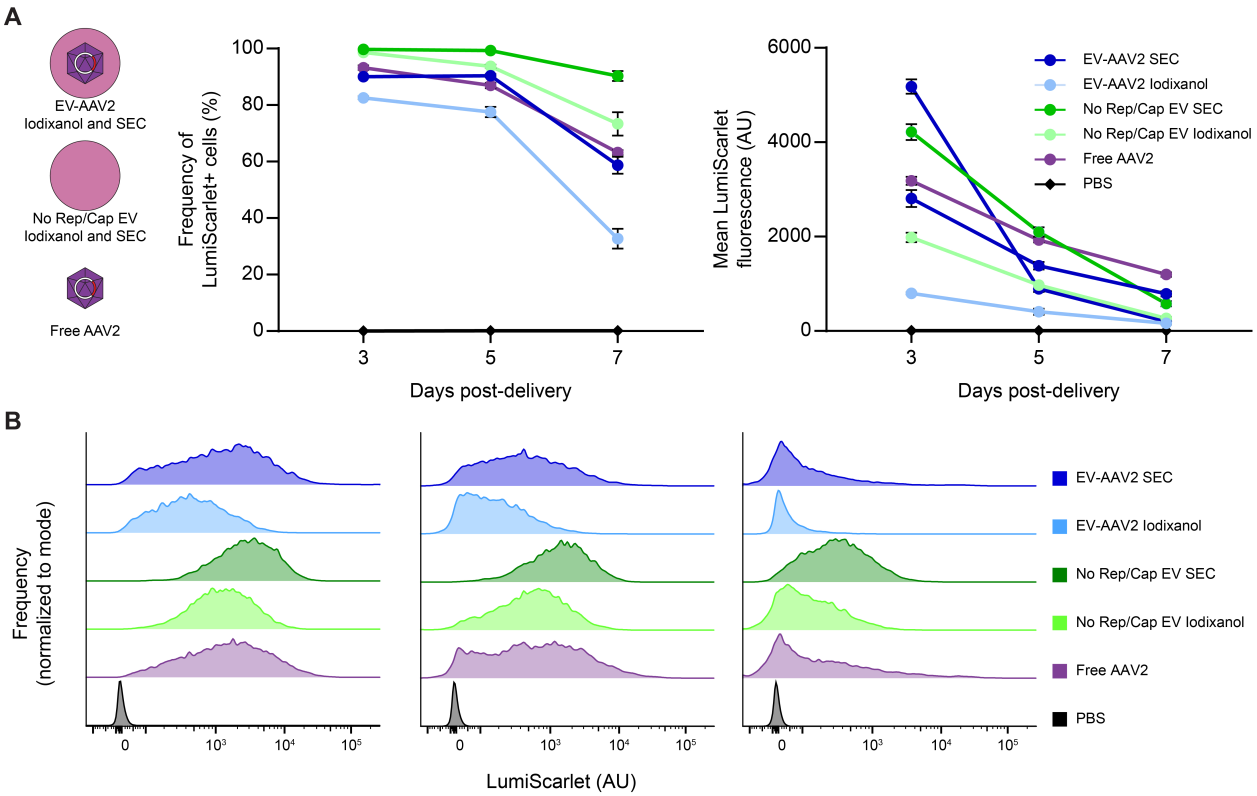


**Supplementary Figure 8: EV-AAV pseudotransduction complicates distinguishing true transduction. (A)** EV-AAV-mediated protein delivery, mRNA and/or expression in HEK293FT recipient cells. Vector compositions and EV-AAV purification methods employed to separate EVs and free AAV are summarized in cartoons at left. EV-AAV vectors were generally similar to the No Rep/Cap (protein delivery only) control at all time points post-delivery. **(B)** Representative histograms of LumiScarlet fluorescence from samples reported in (A). No subpopulations are apparent that can rigorously distinguish protein delivery from gene delivery. Experiments were performed in biological triplicate. Data shown are from one of two independent experiments (first experiment data are in **Figure 1**). PBS: phosphate-buffered saline; SEC: size exclusion chromatography.

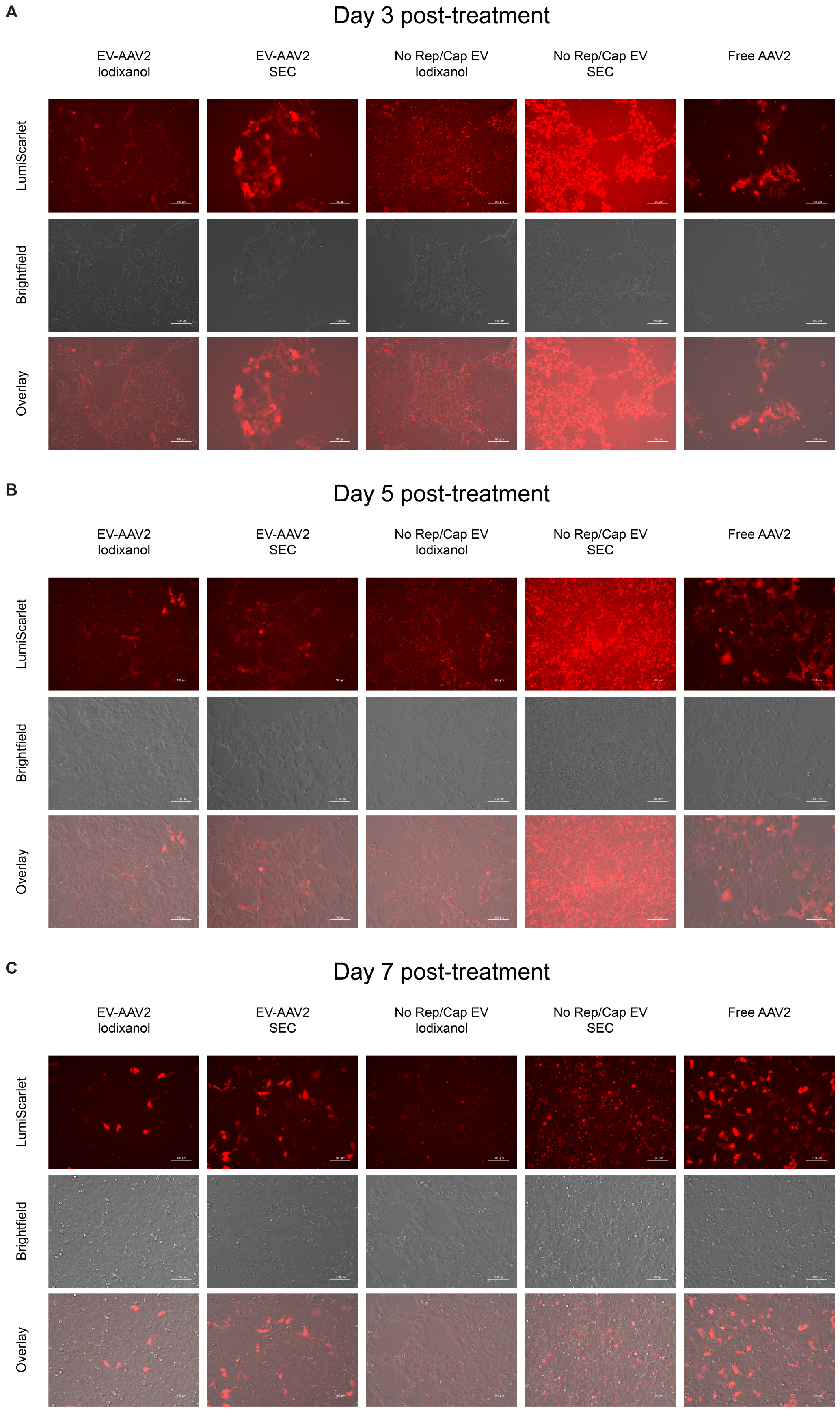
A
Day 3 post-treatment
EV-AAV2
Iodixanol
EV-AAV2
SEC
No Rep/Cap EV
Iodixanol
No Rep/Cap EV
SEC
Free AAV2
LumiScarlet
Brightfield
Overlay
B
Day 5 post-treatment
EV-AAV2
Iodixanol
EV-AAV2
SEC
No Rep/Cap EV
Iodixanol
No Rep/Cap EV
SEC
Free AAV2
LumiScarlet
Brightfield
Overlay
C
Day 7 post-treatment
EV-AAV2
Iodixanol
EV-AAV2
SEC
No Rep/Cap EV
Iodixanol
No Rep/Cap EV
SEC
Free AAV2
LumiScarlet
Brightfield
Overlay

**Supplementary Figure 9: Illustrative micrographs of EV-AAV and AAV pseudotransduction and transduction**. **(A-C)** Microscopy images of HEK293FT cells treated with various vectors, Scale bars: 100 µm. Whole cell fluorescence is apparent for free AAV2 conditions, and puncta are evident in EV-treated conditions. Images were taken of cells analyzed in **Figure S8.**

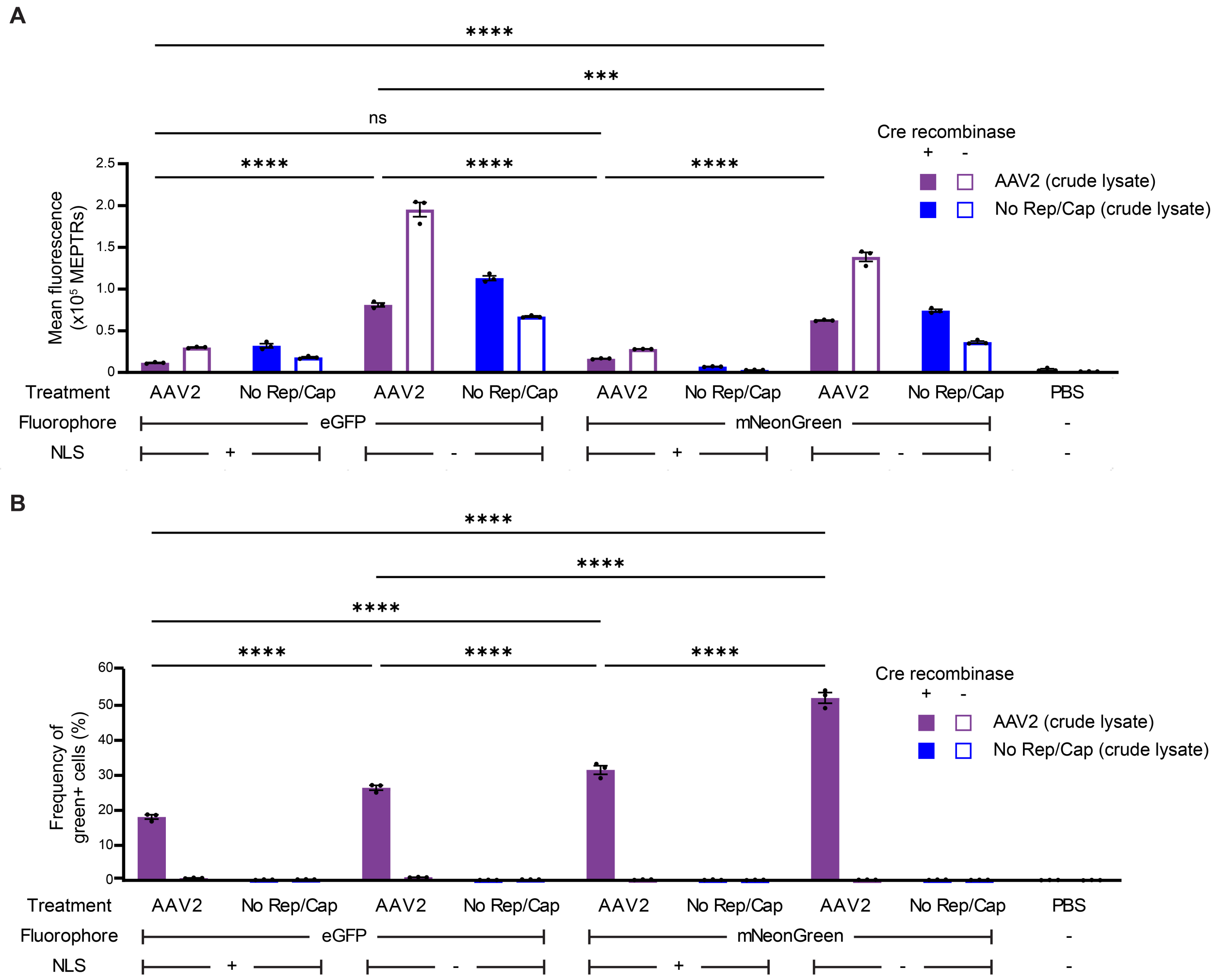


**Supplementary Figure 10: A Cre recombinase-based dual reporter system distinguishes mixed delivery and true transduction. (A,B)** Mixed delivery (A) and true transduction (B) conferred by various vector compositions. Note: some mCherry gene expression could occur in recipient cells prior to Cre-mediated recombination (or in the possible absence of recombination in some cells), such that (C) includes both mixed delivery and this ambiguous de novo gene expression. Samples were normalized to include 1e9 vector genomes per well (for AAV crude lysate conditions) or a volume-equivalent of the AAV2 crude lysate condition for No Rep/Cap crude lysate conditions ($10^4$ recipient cells). Experiments were performed in biological triplicate. Data shown are from one of two independent experiments (first experiment data are in **Figure 2**). Error bars indicate standard error of the mean. Multicomparison statistical analysis was performed using a two-way ANOVA test, followed by Tukey's multiple comparison test to evaluate specific comparisons (**p* < 0.05, ***p* < 0.01, ****p* < 0.001, *****p* < 0.0001). NLS: nuclear localization sequence; ns: not significant; PBS: phosphate-buffered saline.

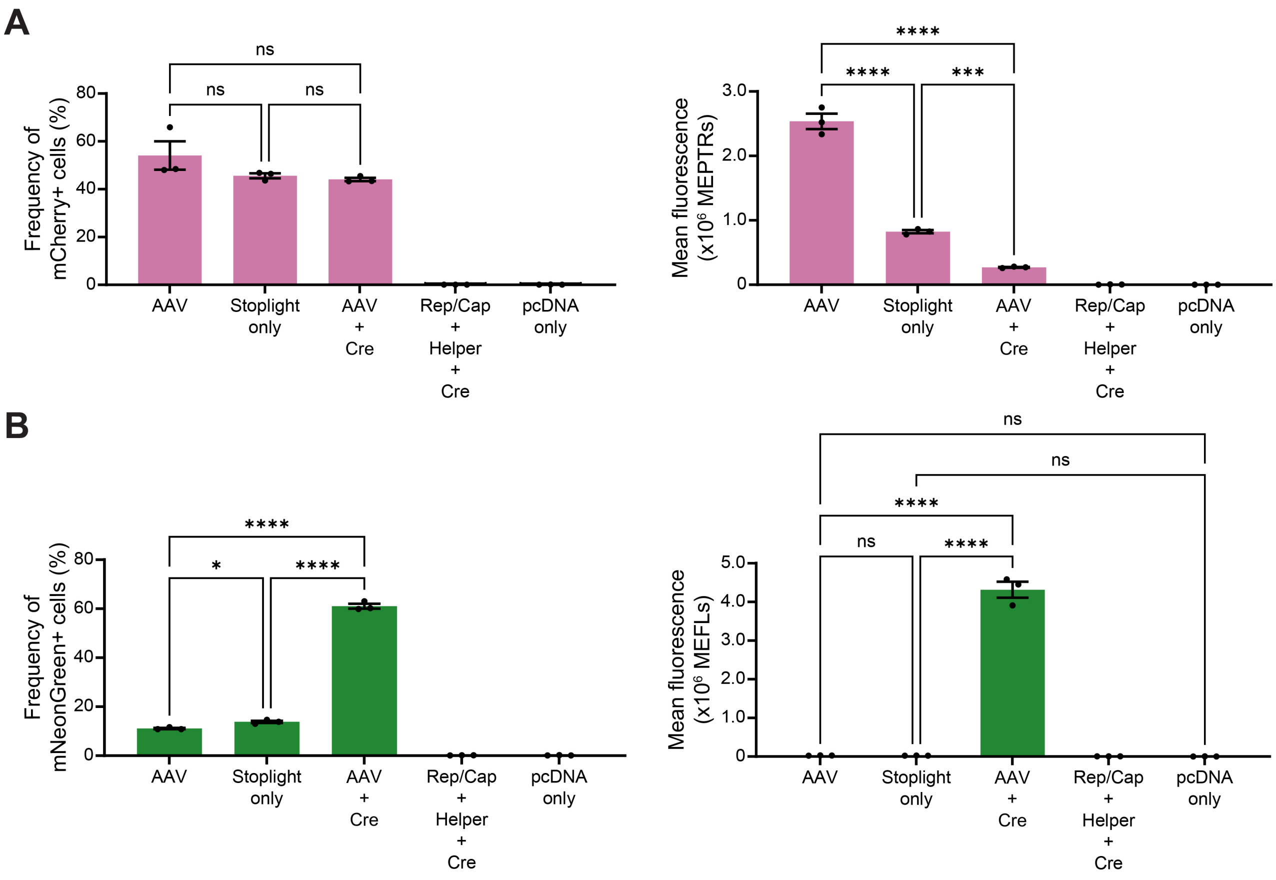


**Supplementary Figure 11: The stoplight plasmid undergoes low levels of spontaneous recombination in producer cells. (A)** mCherry expression in producer cells quantified by the frequency of mCherry positive cells (left) and mean fluorescence (right). **(B)** mNeonGreen expression in producer cells quantified by the frequency of mNeonGreen positive cells (left) and mean fluorescence (right). Freestyle 293-F producer cells were transfected with an equivalent mass of DNA, treated with 4% PFA and analyzed by flow cytometry 3 d post transfection. Data shown are from one independent experiment. Bars represent the mean of three individual transfections. Error bars indicate standard error of the mean. Multicomparison statistical analysis was performed using a one-way ANOVA test, followed by Tukey's multiple comparison test to evaluate specific comparisons (*$p$ < 0.05, **$p$ < 0.01, ***$p$ < 0.001, ****$p$ < 0.0001).

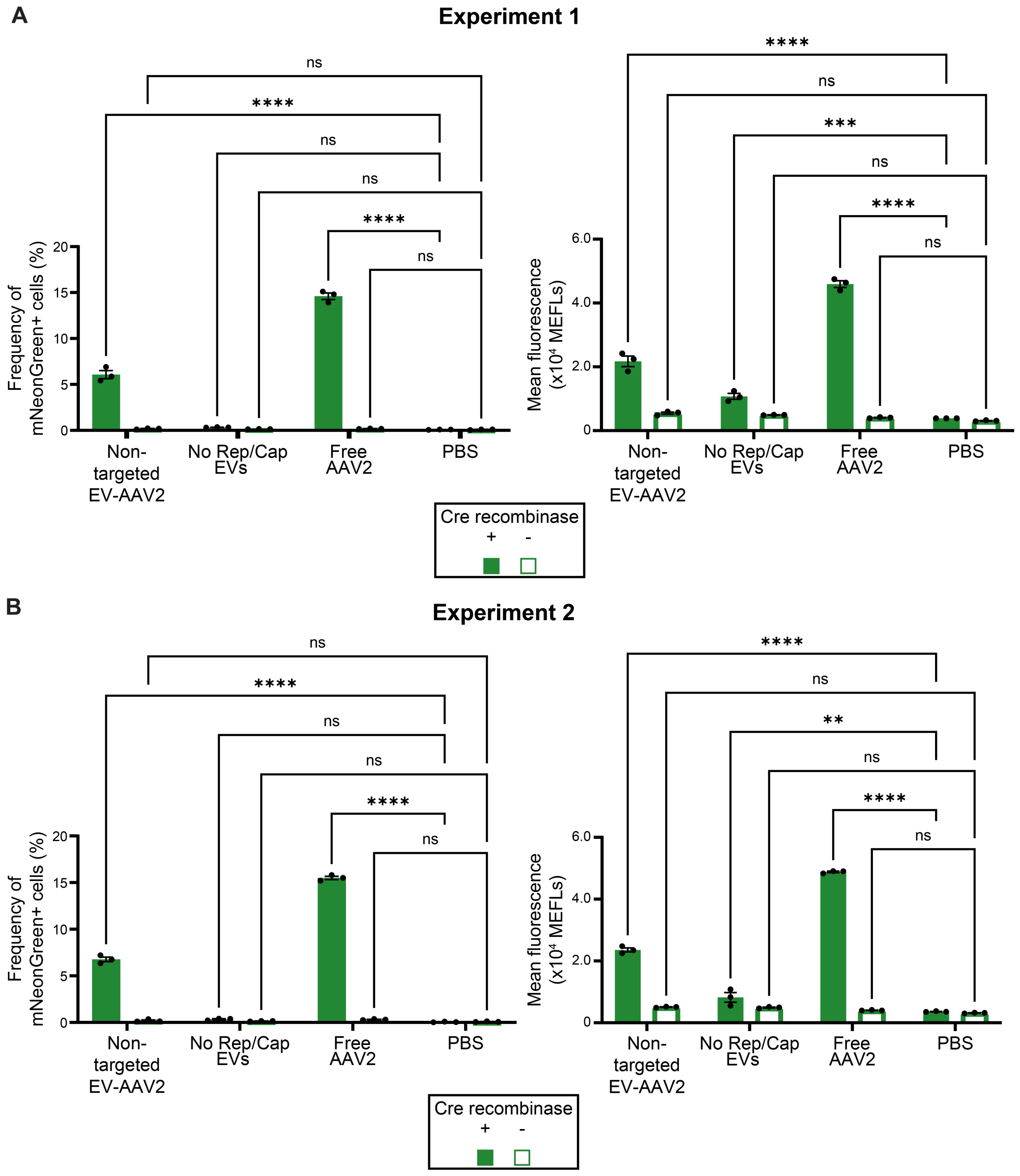


**Supplementary Figure 12: mNeonGreen protein transfer by EVs is minimal. (A,B)** mNeonGreen expression quantified by frequency of mNeonGreen positive cells (left) and mean fluorescence in MEFLs (right) in recipient cells (+/- Cre). Samples were normalized to include 1e9 vector genomes (for EV-AAV2 and Free AAV2 samples) per well or an EV count equivalent of the EV-AAV2 condition for the No Rep/Cap EVs ($10^4$ recipient cells). Experiments were performed in biological triplicate. Data shown are from two independent experiments **(A,B)**. Error bars indicate standard error of the mean. Multicomparison statistical analysis was performed using a one-way ANOVA test, followed by Tukey's multiple comparison test to evaluate specific comparisons (*$p < 0.05$, **$p < 0.01$, ***$p < 0.001$, ****$p < 0.0001$).

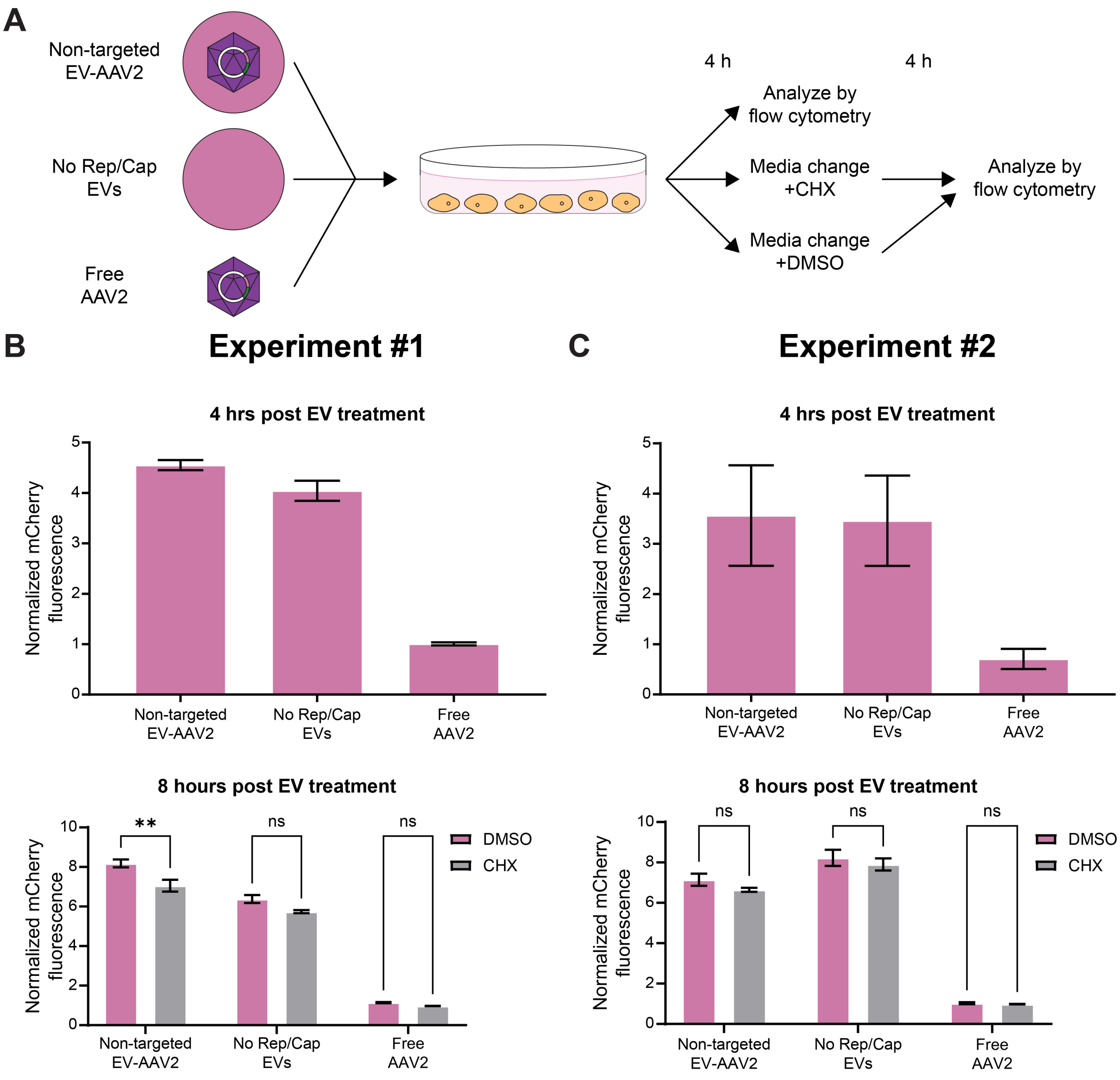


**Supplementary Figure 13: mCherry detected in recipient cells is primarily attributable to protein transfer by EVs. (A)** HEK293FT recipient cells were treated with vector preps or controls as indicated for 4 h, and then fresh medium containing cycloheximide (CHX) or DMSO (solvent control) was added. Samples were normalized to include 1e9 vector genomes (for EV-AAV2 and Free AAV2 samples) per well or an EV count equivalent of the EV-AAV2 condition for the No Rep/Cap EVs ($10^5$ recipient cells plated ~15 h before). **(B,C)** Fluorescence of recipient cells 4 h post EV treatment normalized to PBS treated cells (top). Fluorescence of recipient cells 4 h post media change (8 h post EV treatment) and normalized to fluorescence from PBS treated cells also treated with CHX or DMSO containing medium. Experiments were performed in biological triplicate. Data shown are from two independent experiments (**B** and **C**). Error bars indicate standard error of the mean. Multicomparison statistical analysis was performed using a two-way ANOVA test, followed by Šídák's multiple comparisons test to evaluate specific comparisons (*$p < 0.05$, **$p < 0.01$, ***$p < 0.001$, ****$p < 0.0001$).

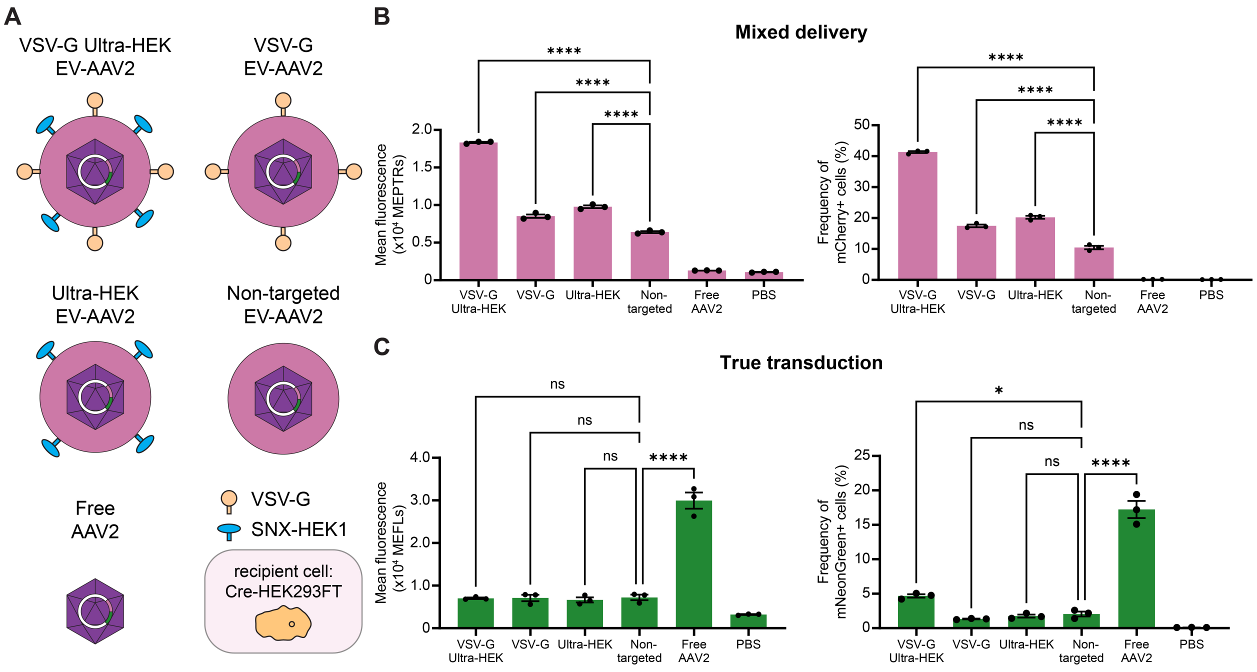


**Supplementary Figure S14: EV-AAV2 functionalization increases mixed delivery but not true transduction to HEK293FTs.** (**A**) Schematic depicting vectors used in (B,C) (Cre-HEK293FT recipient cells). (**B,C**) Mixed delivery quantified by mean fluorescence in MEPTRs (left) and frequency of mCherry positive cells (right) (E) and true transduction quantified by mean fluorescence in MEFLs (left) and frequency of mNeonGreen positive cells (right) (F) to HEK293FT cells (5 d post-delivery). Samples were normalized to include 1e9 vector genomes per well for each condition ($10^4$ recipient cells). VSV-G and Ultra-HEK-functionalized EV-AAV increase protein delivery but only modestly increase gene delivery and only when combined. Experiments were performed in biological triplicate. Data shown are from one of two independent experiments (first experiment data are in **Figure 3**). Error bars indicate standard error of the mean. Multicomparison statistical analysis was performed using a one-way ANOVA test, followed by Tukey's multiple comparison test to evaluate specific comparisons (*$p < 0.05$, **$p < 0.01$, ***$p < 0.001$, ****$p < 0.0001$). PBS: phosphate-buffered saline.

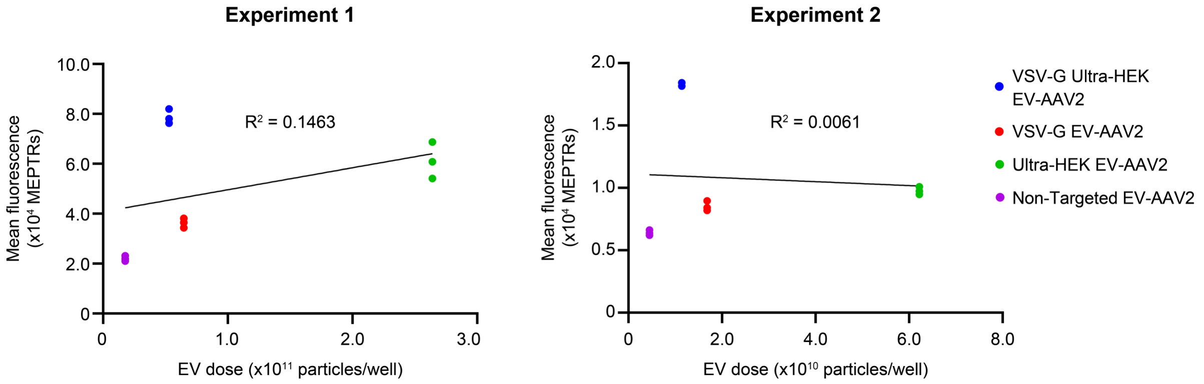


**Supplementary Figure 15: EV dose alone is not an explanatory variable for mixed delivery signal**. Data reported in **Figure 3** (left) or **Figure S15** (right) were replotted to investigate whether EV dose alone can explain protein delivery trends. Each dot represents a flow cytometry data of a single biological replicate (one well of cells) five days post transduction. Linear regression was performed on this dataset to determine if the mixed delivery signal (MEPTRs) is explained mostly by the dose of vesicles delivered to each well of cells. Since EV dose explains only about 15% (left) and 0.6% (right) of the variation in mixed delivery, a significant amount of the mixed delivery signal derives from biological differences attributable to EV functionalization strategies rather than EV dose. Horizontal error bars (occluded by data points) indicate standard error of the mean in EV dose.

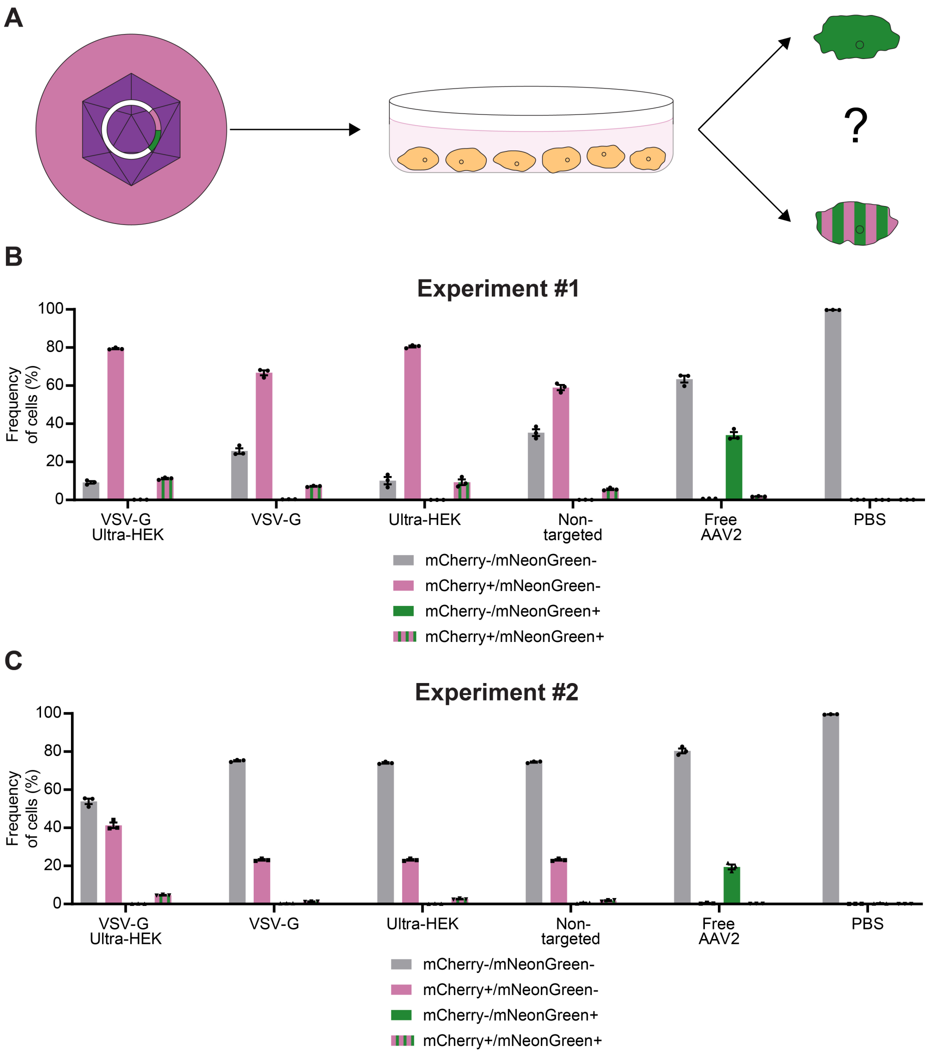


**Supplementary Figure 16: Truly transduced HEK293FT recipient cells treated by EV-AAVs also are positive for mixed delivery mCherry expression. (A)** Schematic representing the question as to whether HEK293FT recipient cells that were transduced are a separate population or a subset of those that experience mixed delivery. **(B,C)** Frequency of recipient cells analyzed for no fluorescence expression (mCherry-/mNeonGreen-), mCherry expression only (mCherry+/mNeonGreen-), mNeonGreen expression only (mCherry-/mNeonGreen+), or both mCherry and mNeonGreen expression (mCherry+/mNeonGreen+). Data are re-plotted from **Figure 3** (B) and **Figure S15** (C). Almost all HEK293FT cells that are truly transduced (mNeonGreen+) by EV-AAVs also experience mixed delivery (mCherry+), regardless of functionalization strategy. Experiments were performed in biological triplicate. Data shown are from two independent experiments **(B**,**C)**. Error bars indicate standard error of the mean.

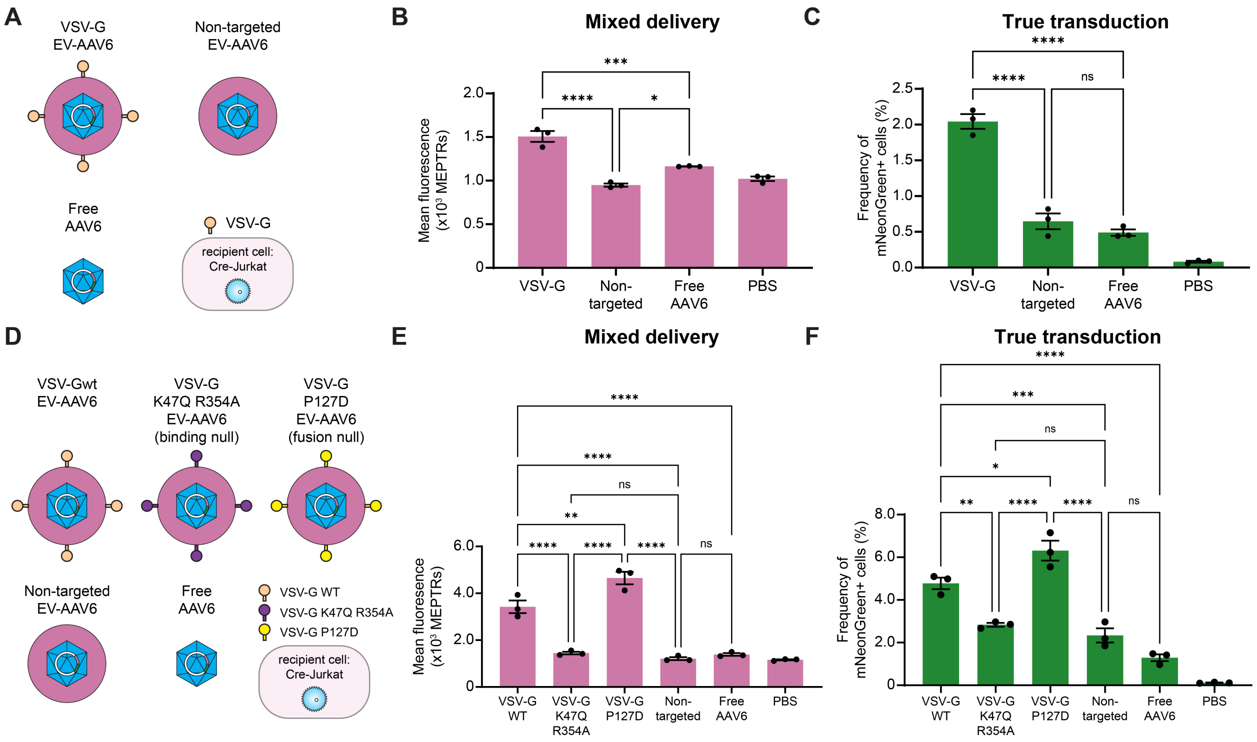


**Supplementary Figure 17: VSV-G functionalization of EV-AAV6 increases true transduction and mixed delivery to Jurkat T cells by increasing uptake. (A)** Schematic depicting vectors used in (B,C) (Cre-expressing Jurkat T cell recipient cells). (**B,C**) Mixed delivery (B) and true transduction (C) in Jurkat T cells (5 d post-delivery). Samples were normalized to include 1e9 vector genomes per well for each condition ($10^4$ recipient cells). VSV-G functionalization significantly increases mixed delivery to and transduction of Jurkats. **(D)** Schematic depicting vectors used in (E,F) (Cre-Jurkat T cell recipient cells). (**E,F**) Mixed delivery (E) and true transduction (F) in Jurkat T cells (5 d post-delivery). Samples were normalized to include 2e9 vector genomes per well for each condition ($10^4$ recipient cells). VSV-G functionalization significantly increases mixed delivery to and true transduction of Jurkats. Mixed delivery and true transduction are increased only by enhanced binding of recipient cells (VSV-G WT and VSV-G P127D vectors). Experiments were performed in biological triplicate. Data shown are from one of two independent experiments (first experiments in **Figure 4**). Error bars indicate standard error of the mean. Multicomparison statistical analysis was performed using a one-way ANOVA test, followed by Tukey's multiple comparison test to evaluate specific comparisons (*$p < 0.05$, **$p < 0.01$, ***$p < 0.001$, ****$p < 0.0001$). PBS: phosphate-buffered saline.

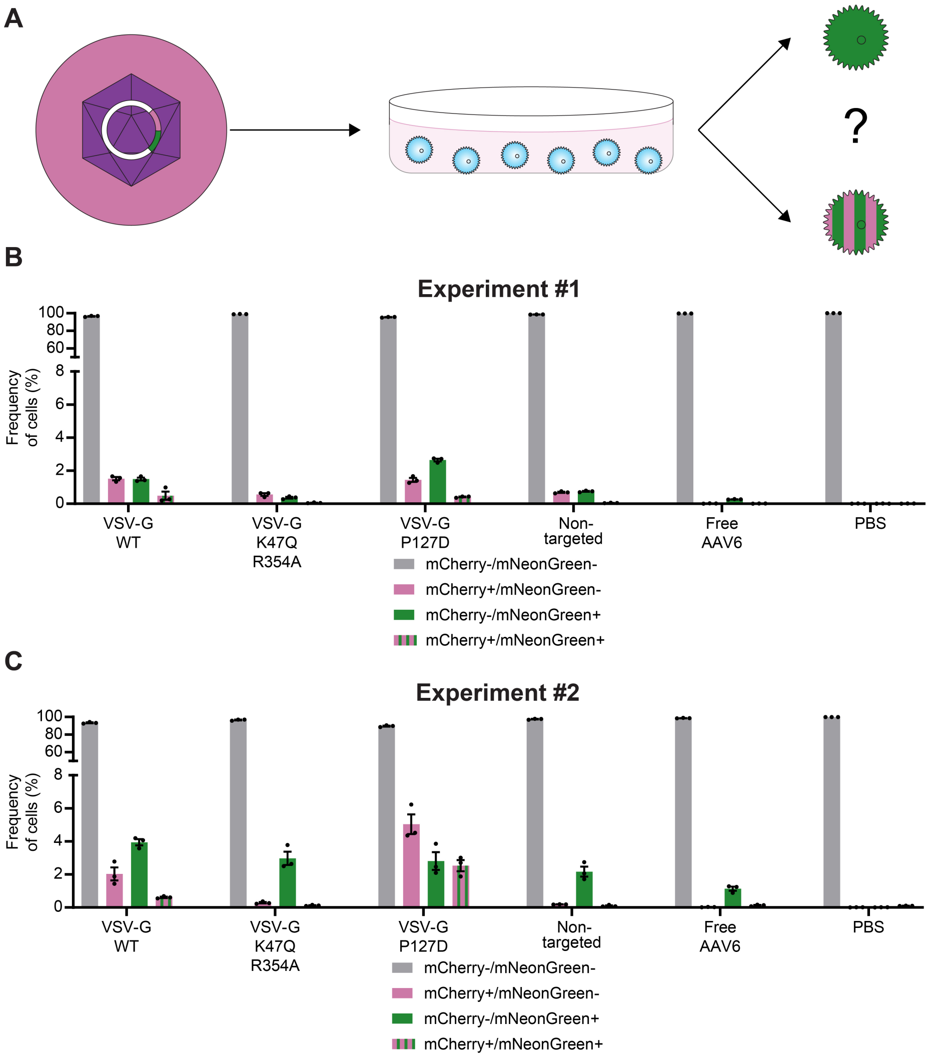


**Supplementary Figure 18: Some truly transduced Jurkat recipient cells treated by EV-AAVs are a distinct population from Jurkats positive for mixed delivery mCherry expression. (A)** Schematic representing the question as to whether Jurkat recipient cells that were transduced are a separate population or a subset of those that experience mixed delivery. **(B,C)** Frequency of recipient cells analyzed for no fluorescence expression (mCherry-/mNeonGreen-), mCherry expression only (mCherry+/mNeonGreen-), mNeonGreen expression only (mCherry-/mNeonGreen+), or both mCherry and mNeonGreen expression (mCherry+/mNeonGreen+). Data re-plotted from **Figure 4** (B) and **Figure S18** (C). Some truly transduced (mNeonGreen+) Jurkats treated by EV-AAVs do not express mCherry. The frequency of mCherry and mNeonGreen expression (mCherry+/mNeonGreen+) increases with enhanced binding of recipient cells (VSV-G WT and VSV-G P127D vectors). Experiments were performed in biological triplicate. Data shown are from two independent experiments **(B**,**C)**. Error bars indicate standard error of the mean.

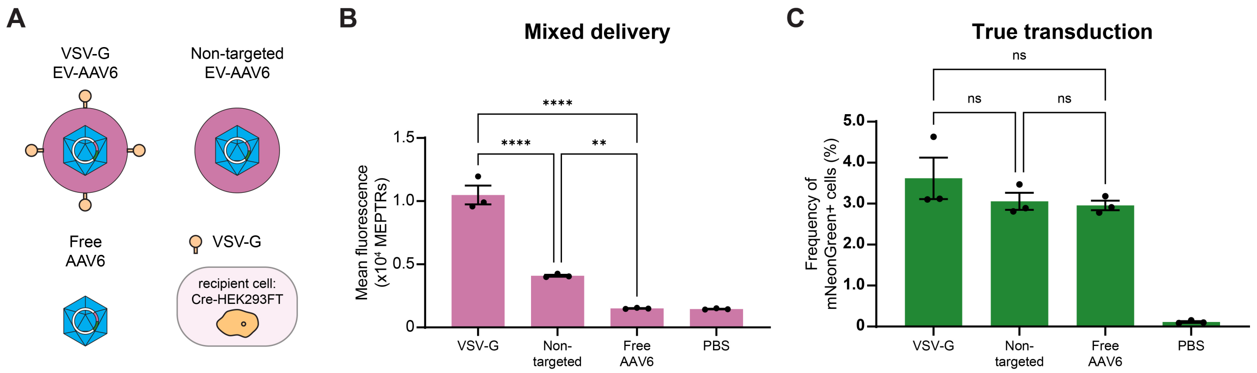


**Supplementary Figure 19: Determinants of EV-AAV delivery to HEK293FT cells apply across two AAV serotypes. (A)** Schematic depicting vectors used in (B,C) (Cre-HEK293FT recipient cells). (**B,C**) Mixed delivery (B) and true transduction (C) to HEK29FT cells (5 d post-delivery). Samples were normalized to include 1e9 vector genomes per well for each condition ($10^4$ recipient cells). VSV-G functionalization enhances mixed delivery but not true transduction by EV-AAV6, concordant with observations made using EV-AAV2. Experiments were performed in biological triplicate. Data shown are from one of two independent experiments (first experiment data are in **Figure 5**). Error bars indicate standard error of the mean. Multicomparison statistical analysis was performed using a one-way ANOVA test, followed by Tukey's multiple comparison test to evaluate specific comparisons (*$p < 0.05$, **$p < 0.01$, ***$p < 0.001$, ****$p < 0.0001$). PBS: phosphate-buffered saline.
