## Supplementary figures and images for "Distinguishing Pseudotransduction and True Transduction Enables Characterization and Bioengineering of Extracellular Vesicle-Adeno-Associated Virus Vectors"

### EV-AAV2 Iodixanol_Brightfield_Day3.tif

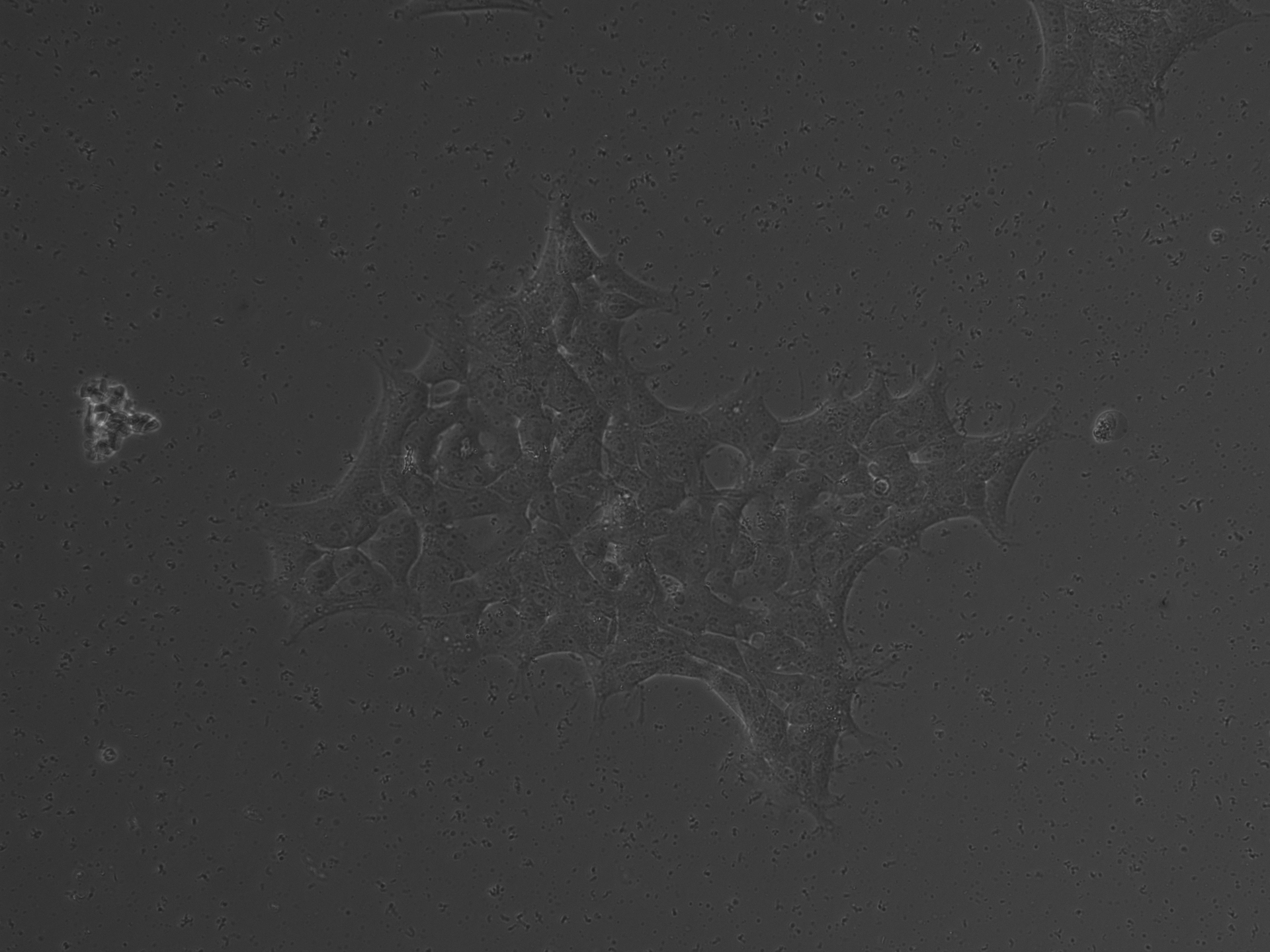

### EV-AAV2 Iodixanol_Brightfield_Day3.tif

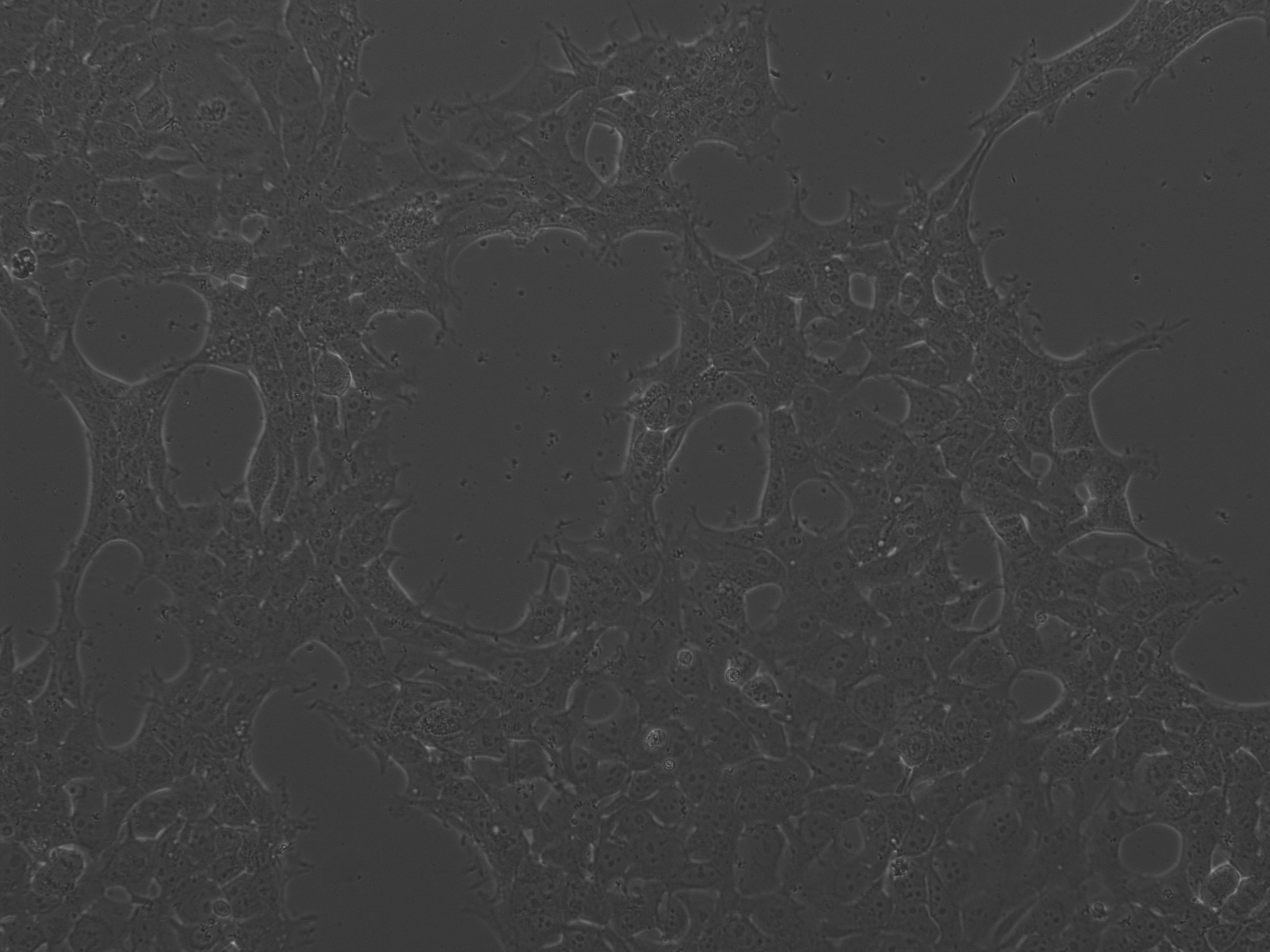

### EV-AAV2 Iodixanol_Brightfield_Day5.tif

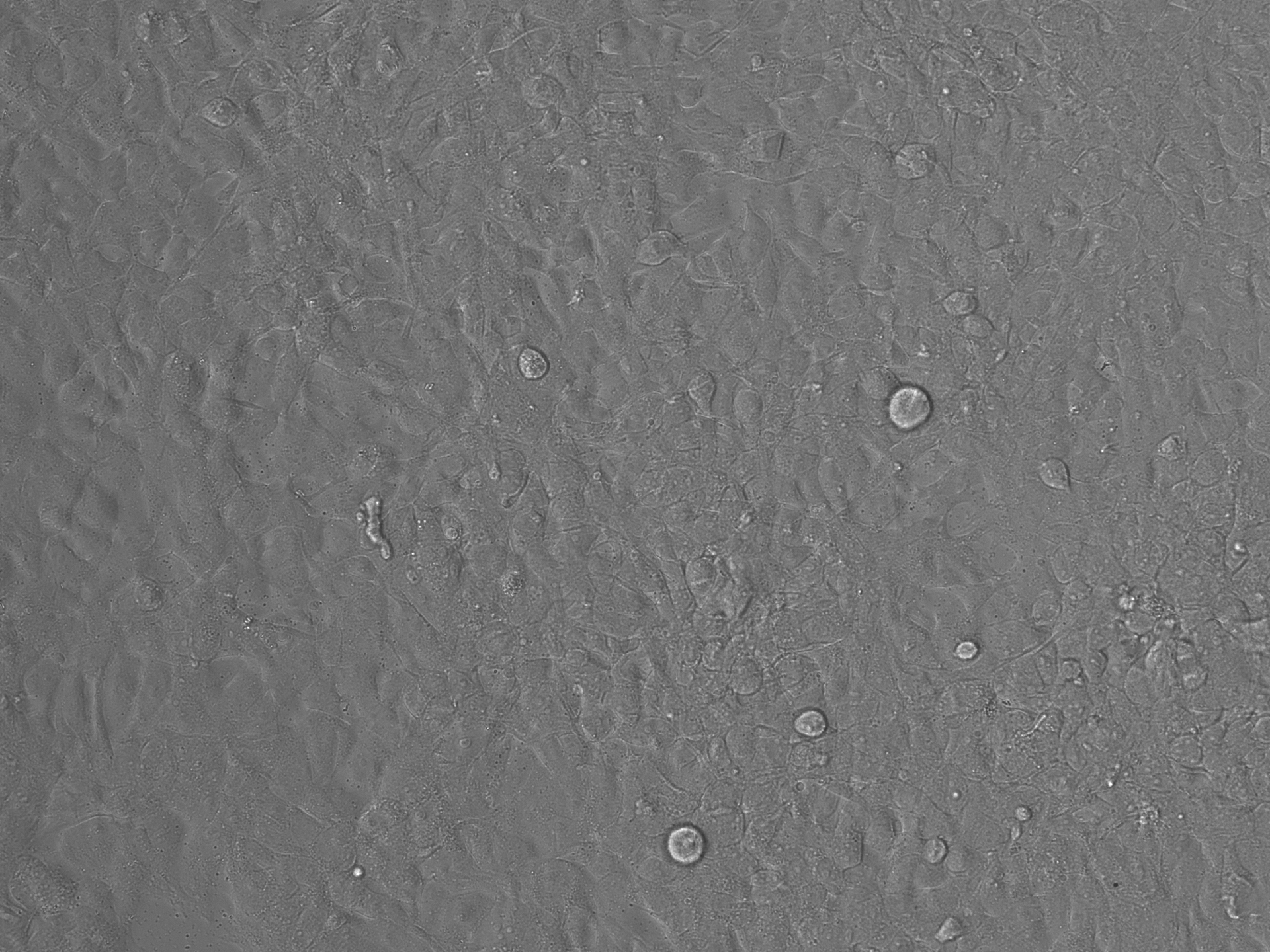

### EV-AAV2 Iodixanol_Brightfield_Day5.tif

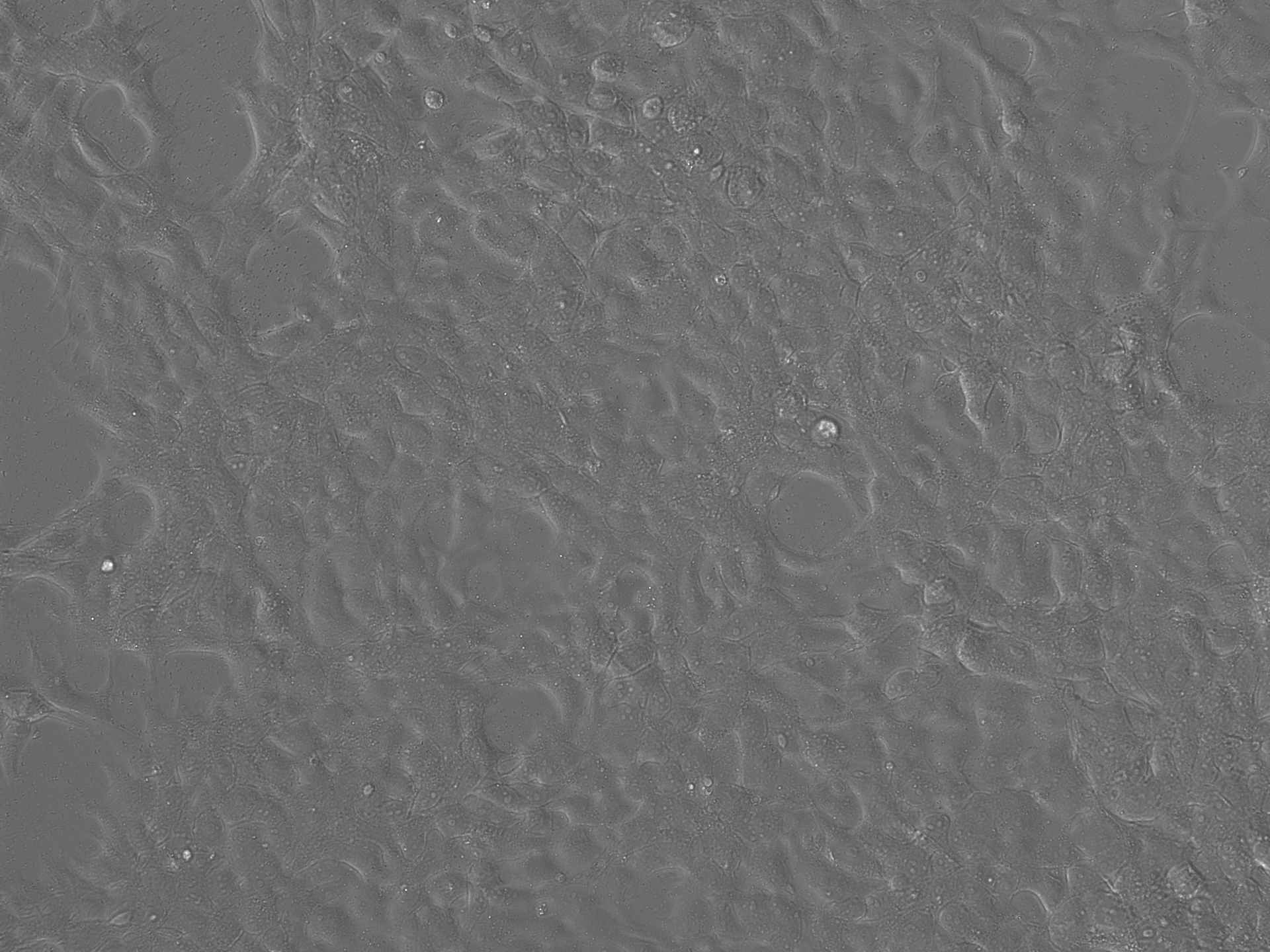

### EV-AAV2 Iodixanol_Brightfield_Day7.tif

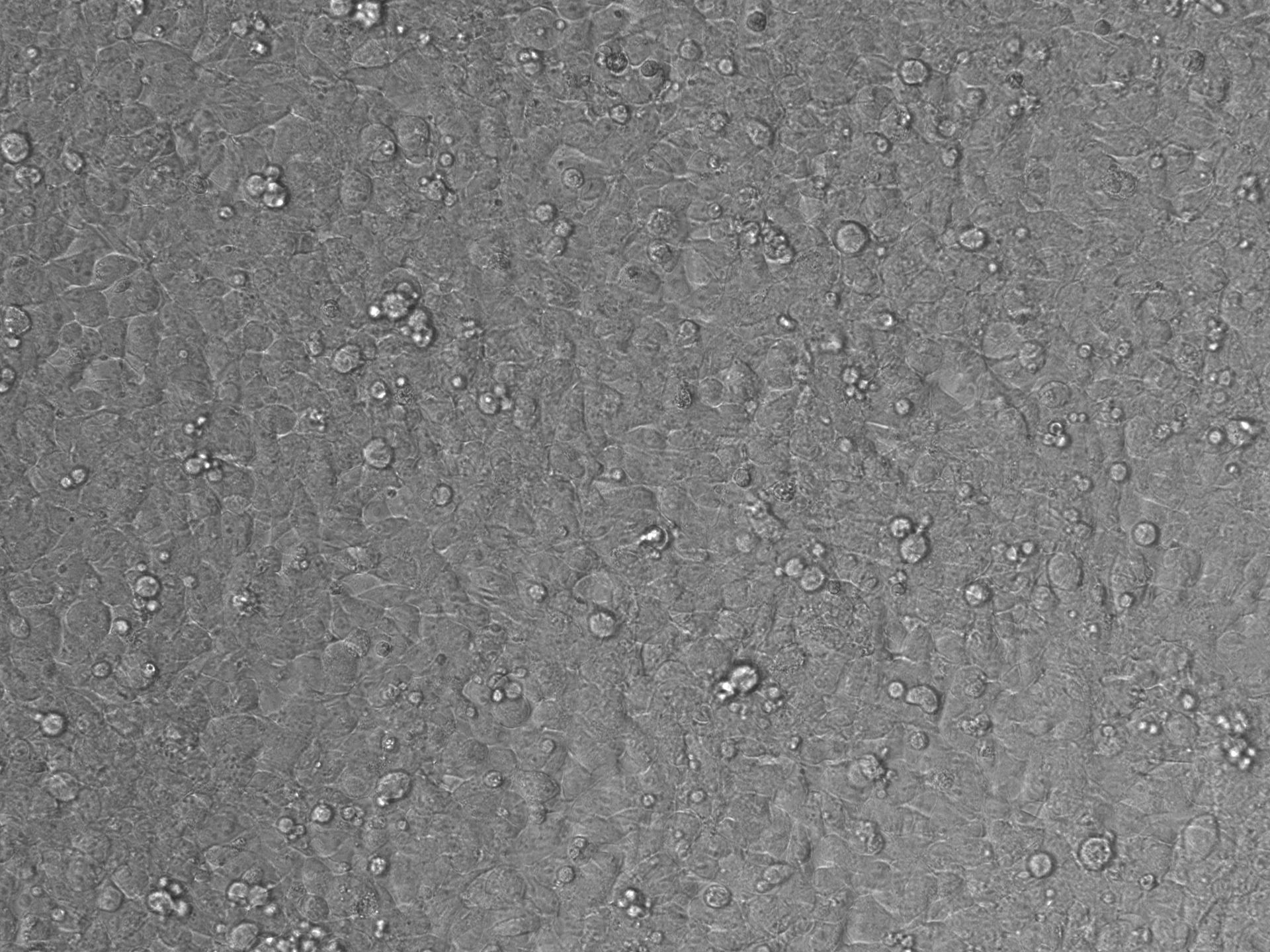

### EV-AAV2 Iodixanol_Brightfield_Day7.tif

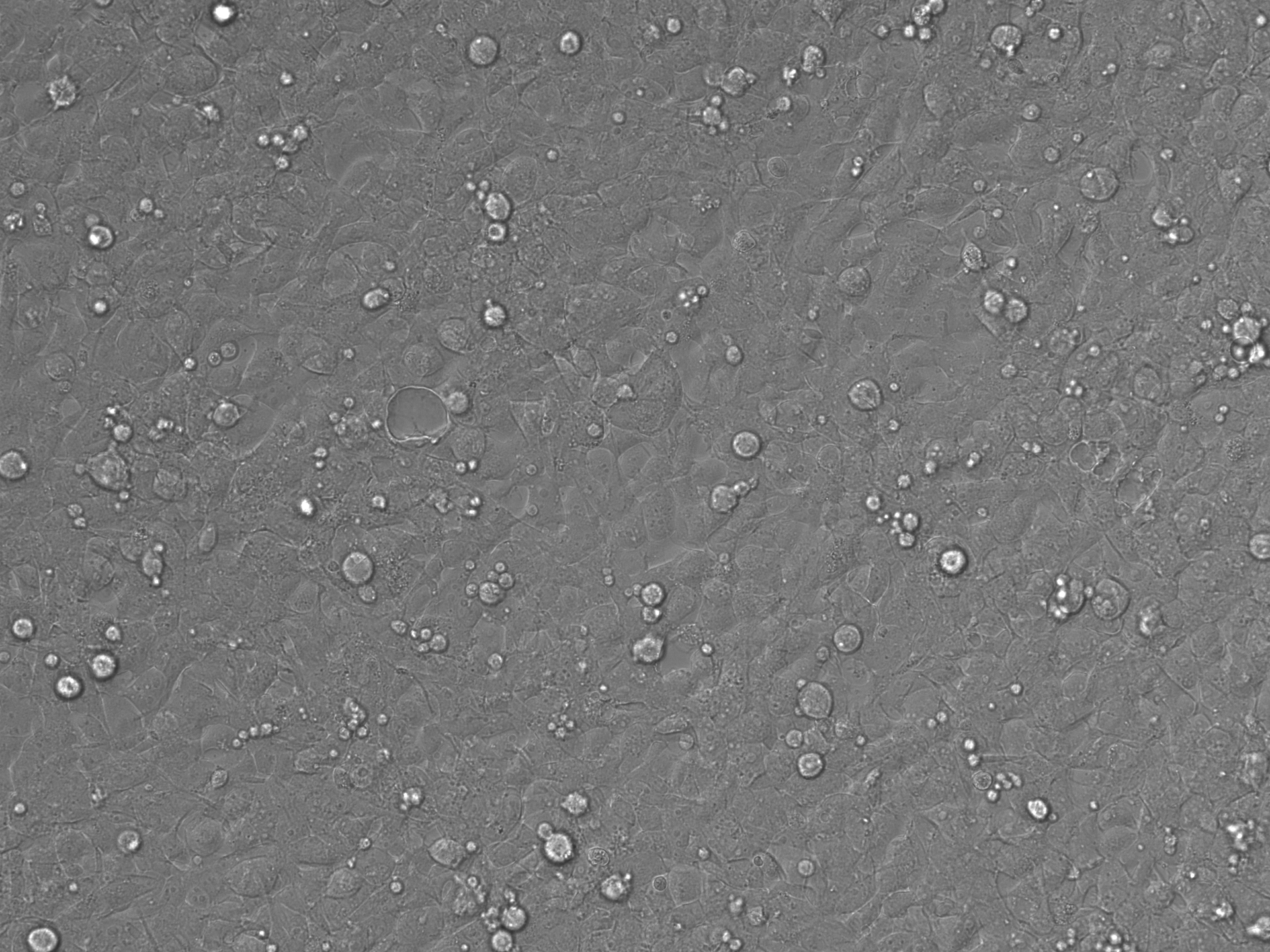

### EV-AAV2 Iodixanol_LumiScarlet_Day3.tif

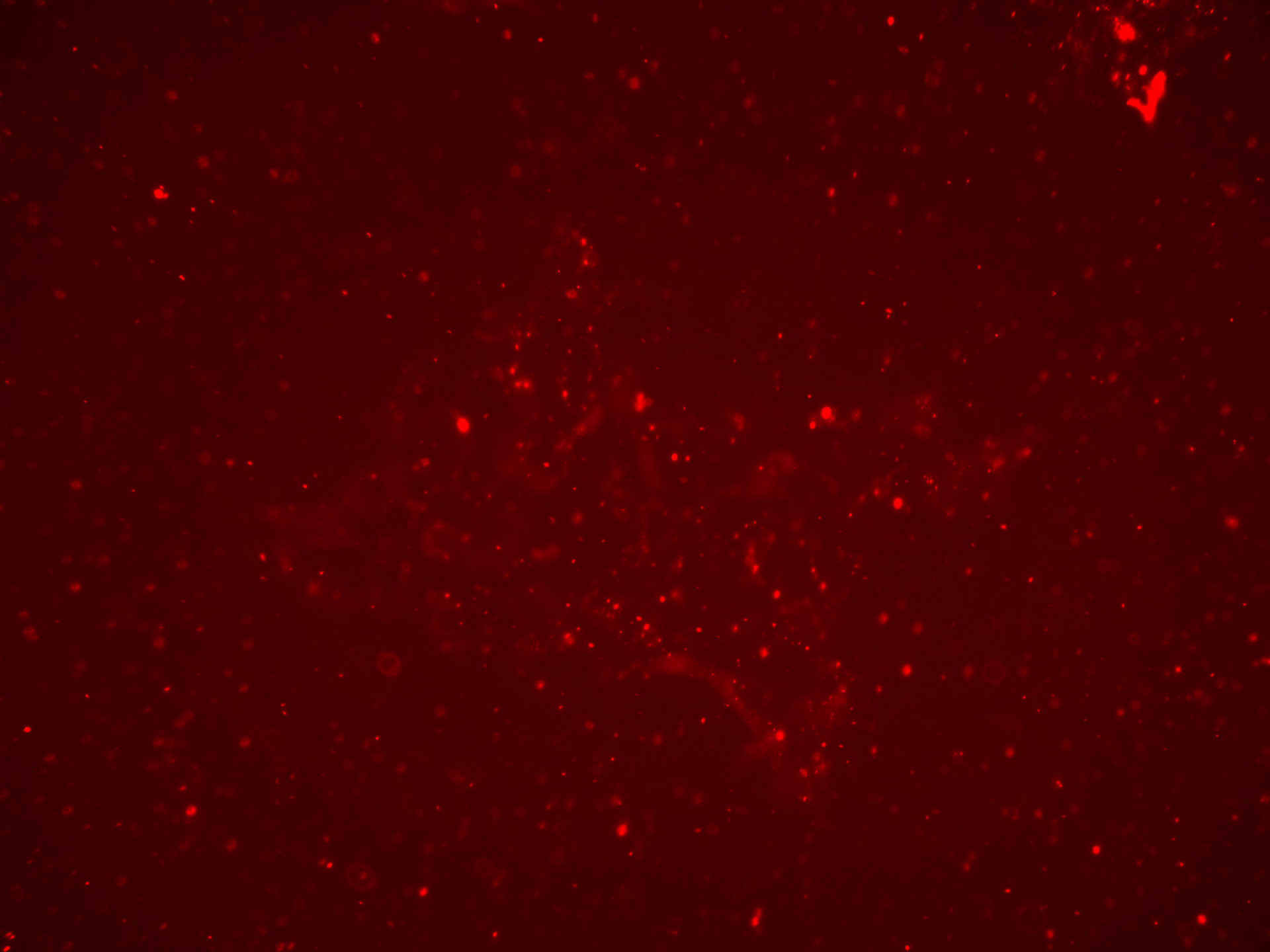

### EV-AAV2 Iodixanol_LumiScarlet_Day3.tif

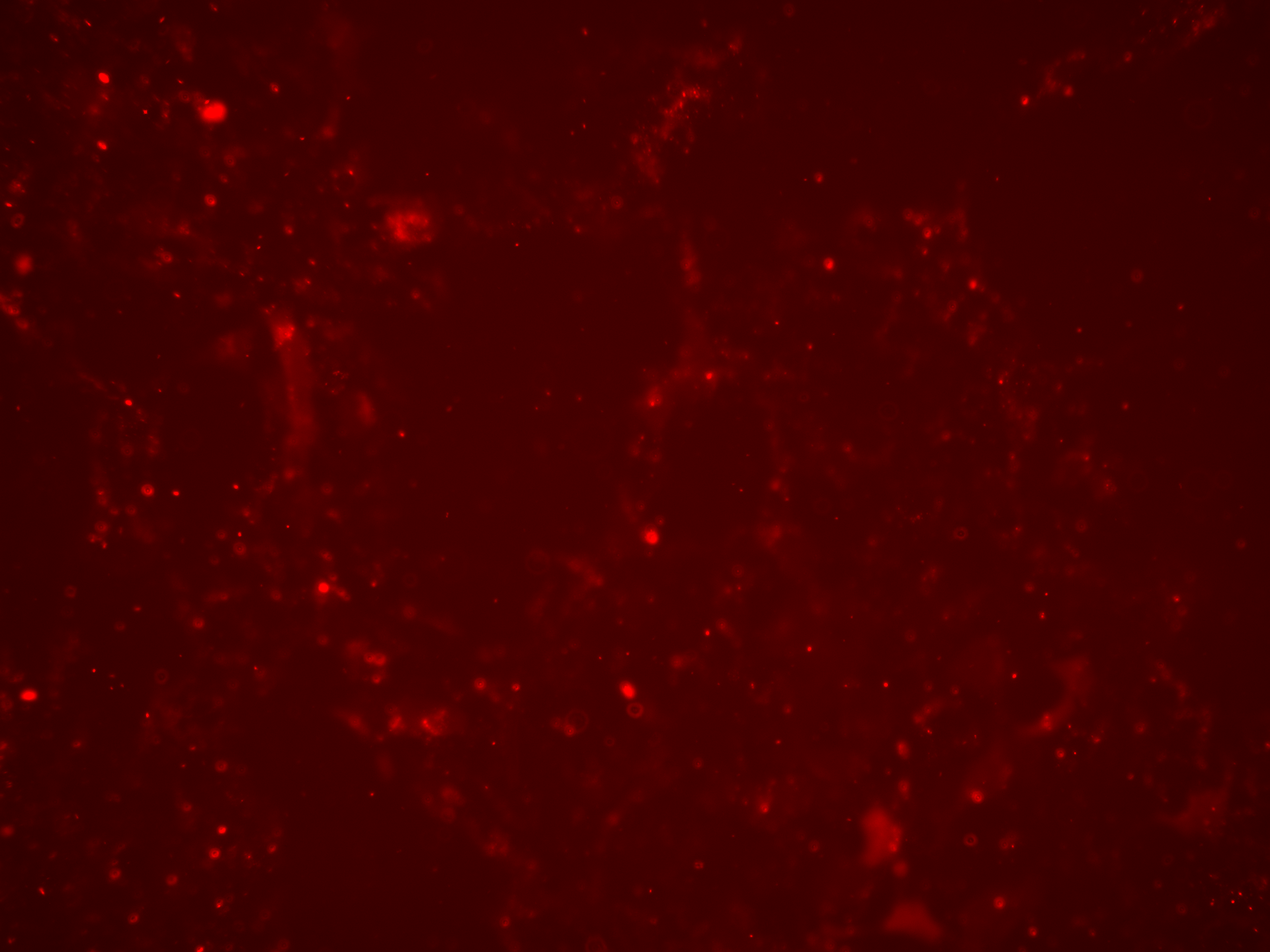

### EV-AAV2 Iodixanol_LumiScarlet_Day5.tif

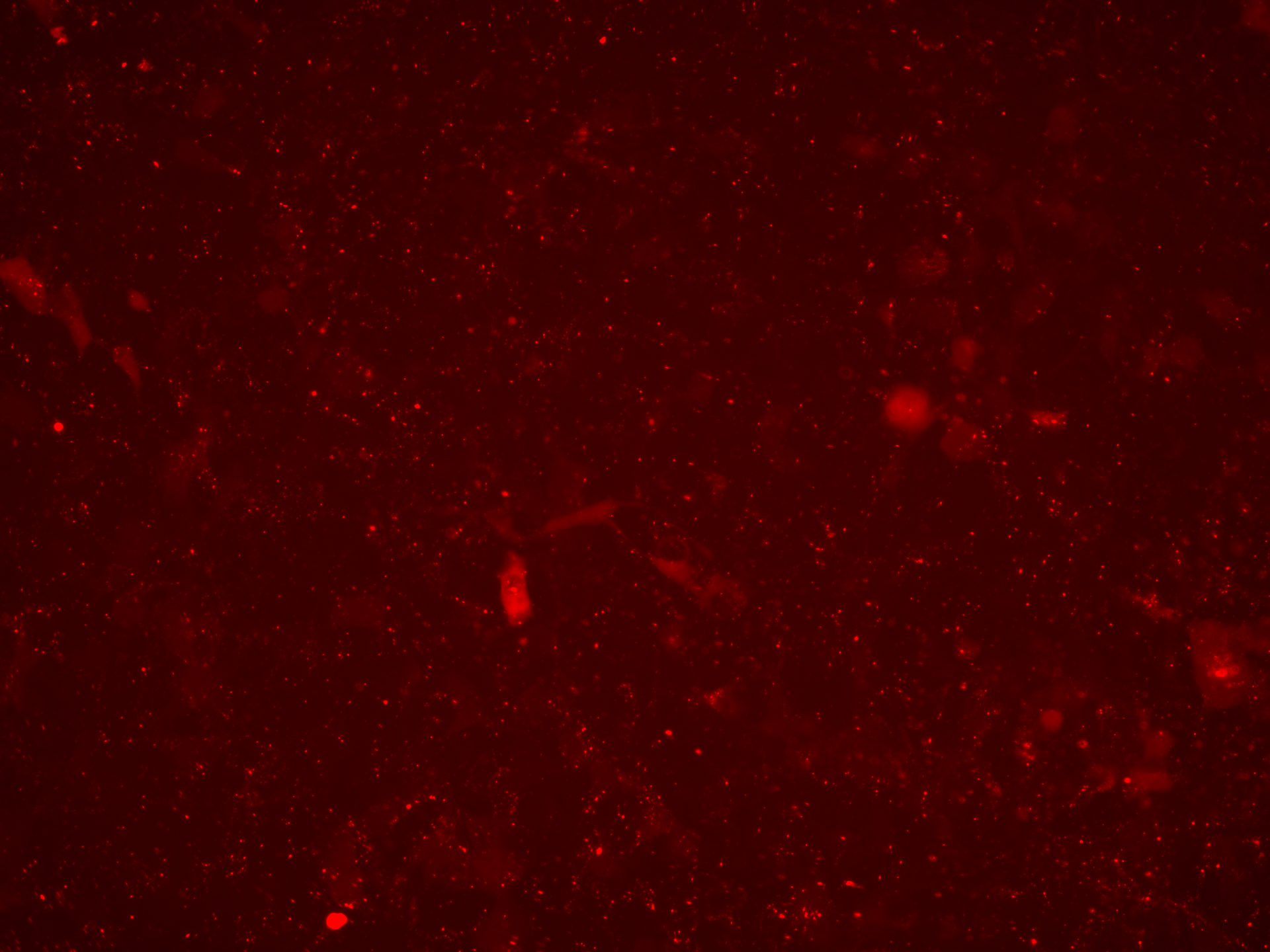

### EV-AAV2 Iodixanol_LumiScarlet_Day5.tif

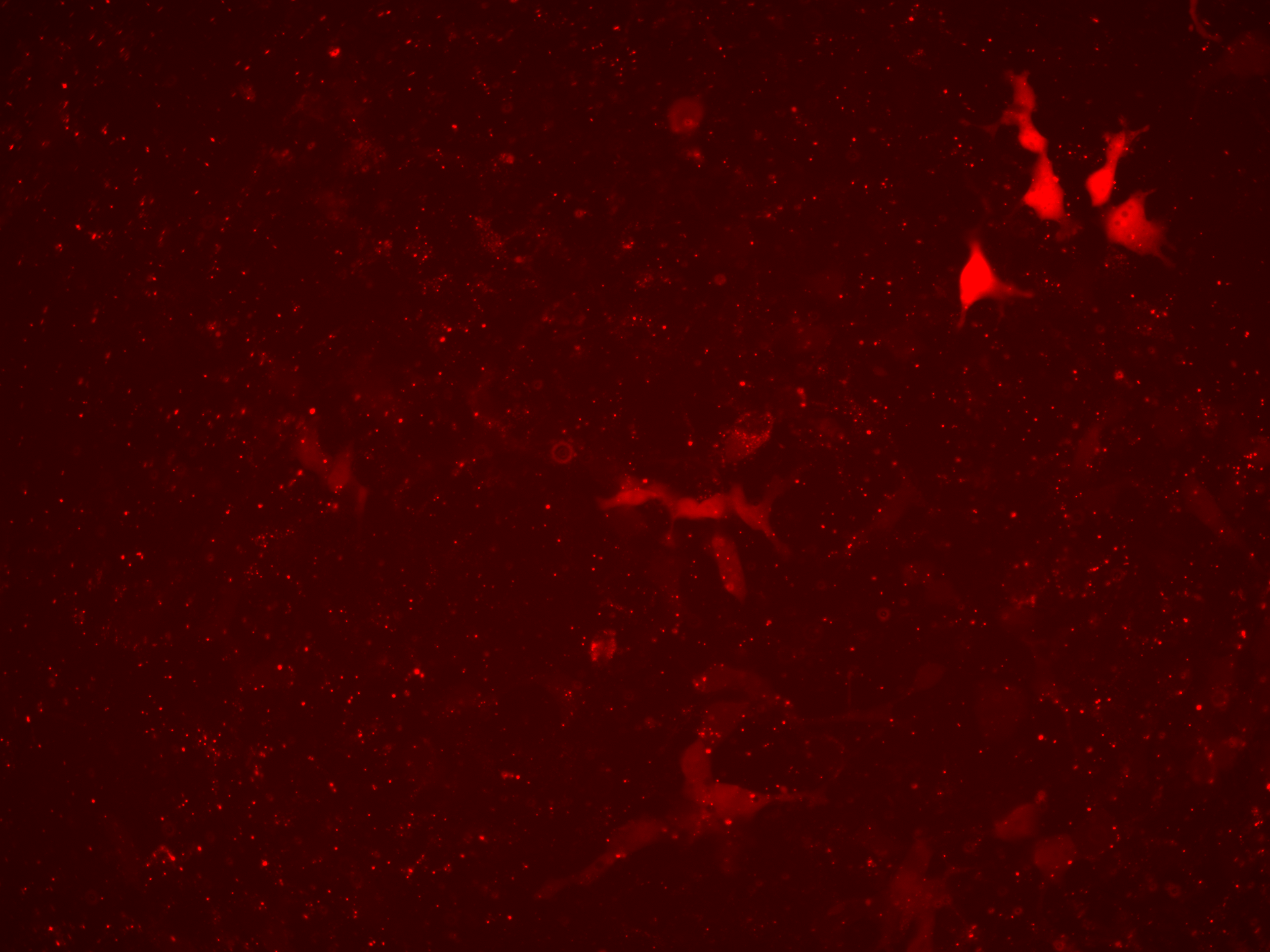

### EV-AAV2 Iodixanol_LumiScarlet_Day7.tif

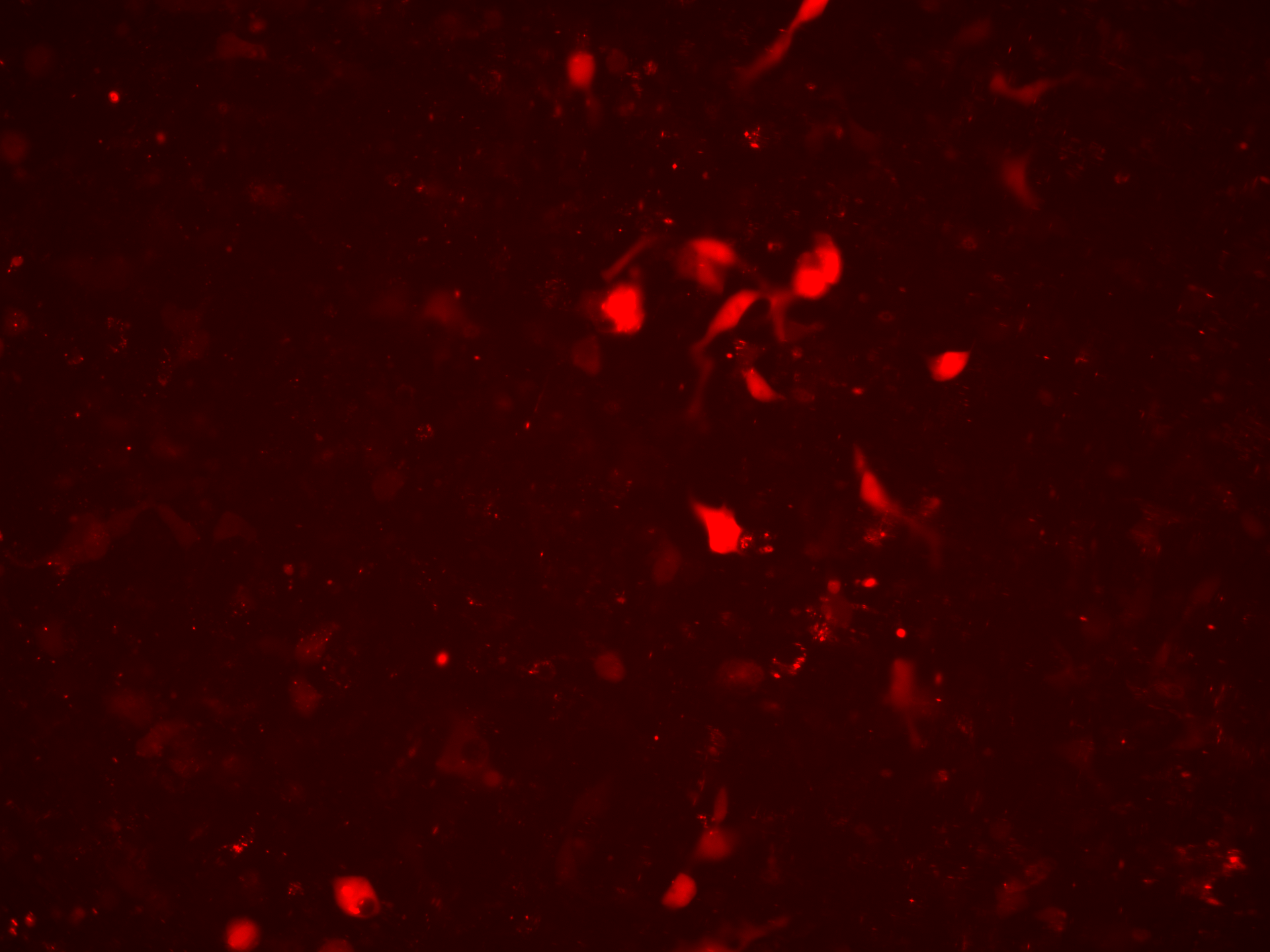

### EV-AAV2 Iodixanol_LumiScarlet_Day7.tif

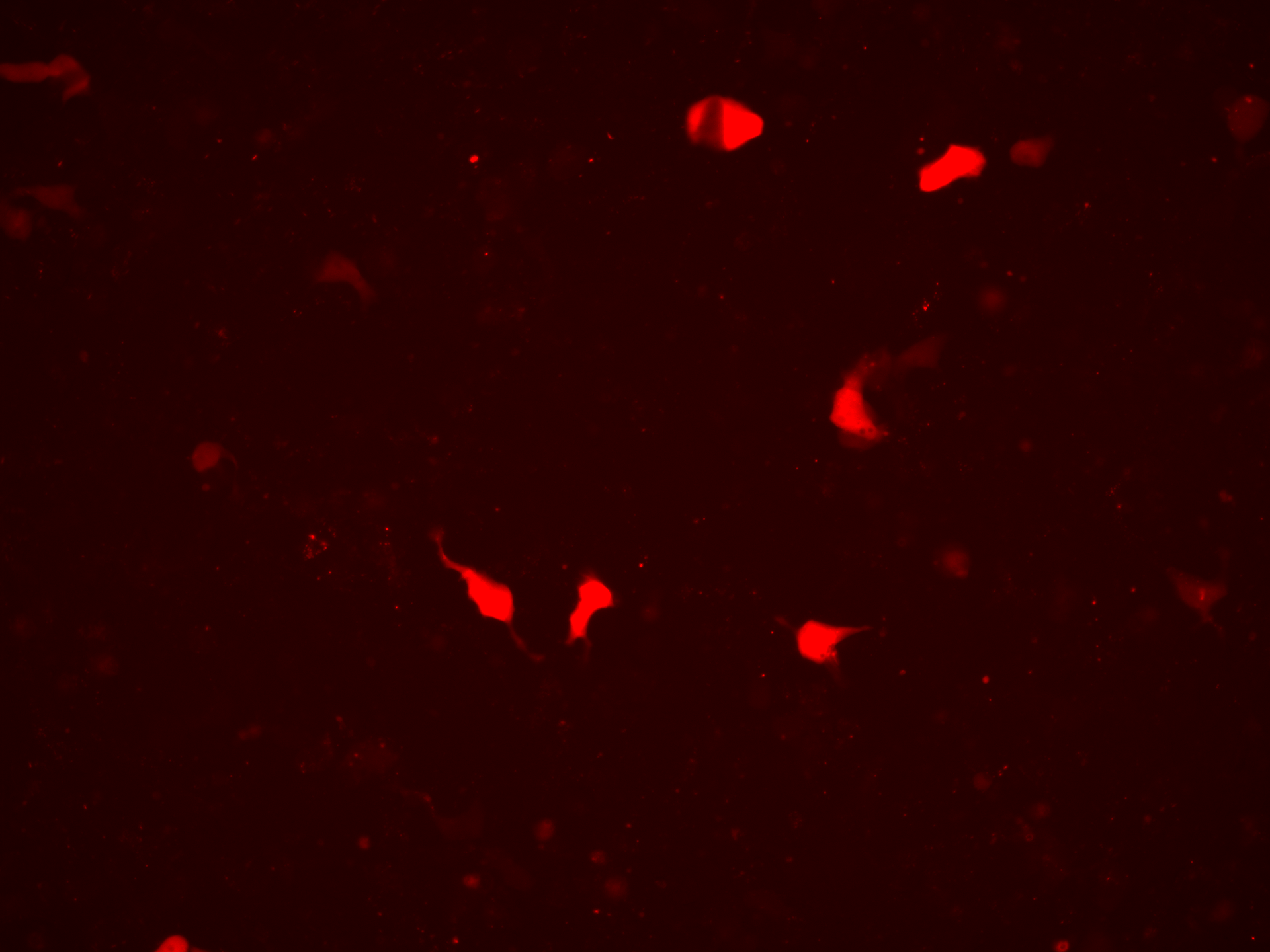

### EV-AAV2 Iodixanol_Overlay_Day3.tif

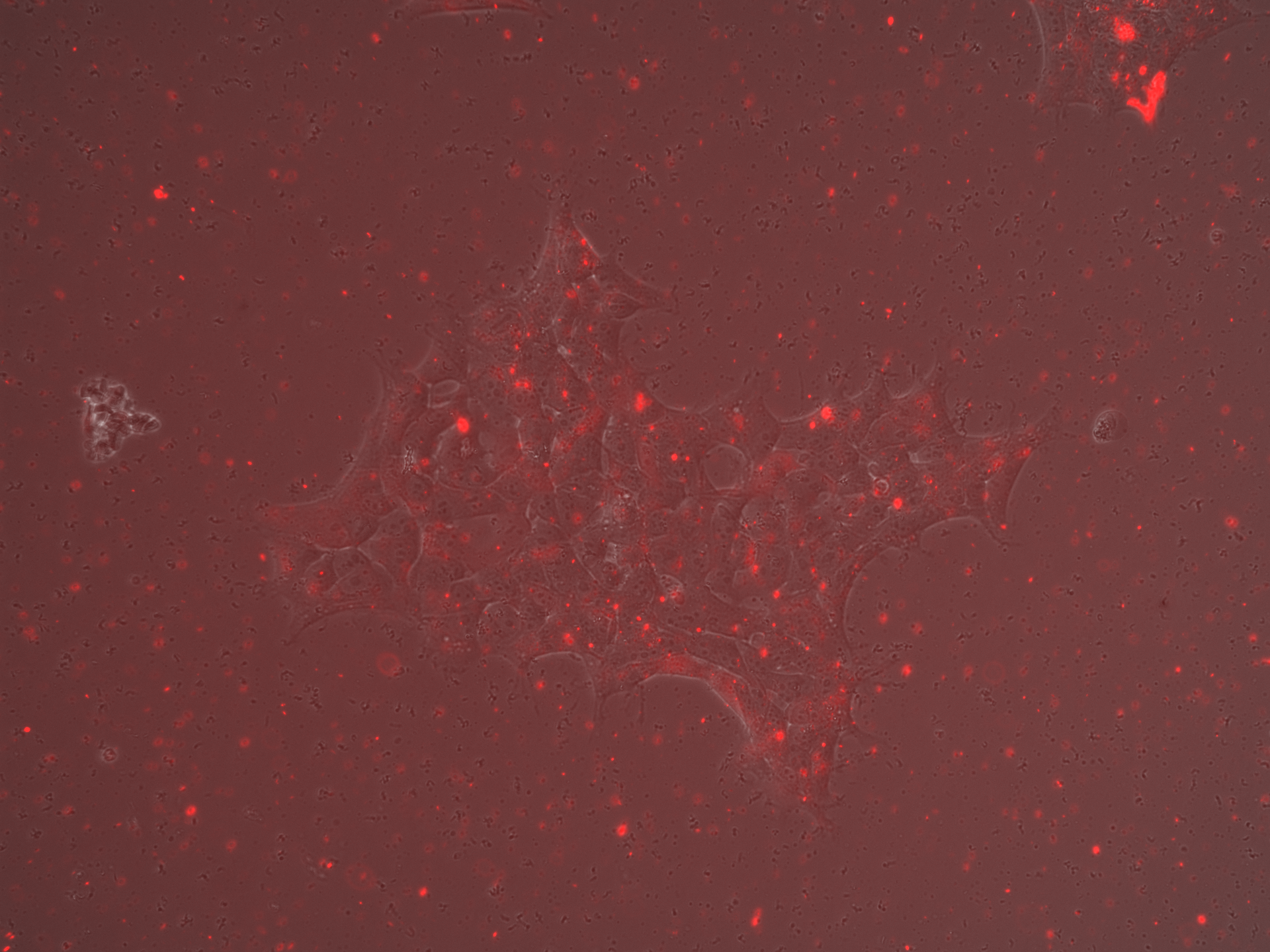

### EV-AAV2 Iodixanol_Overlay_Day3.tif

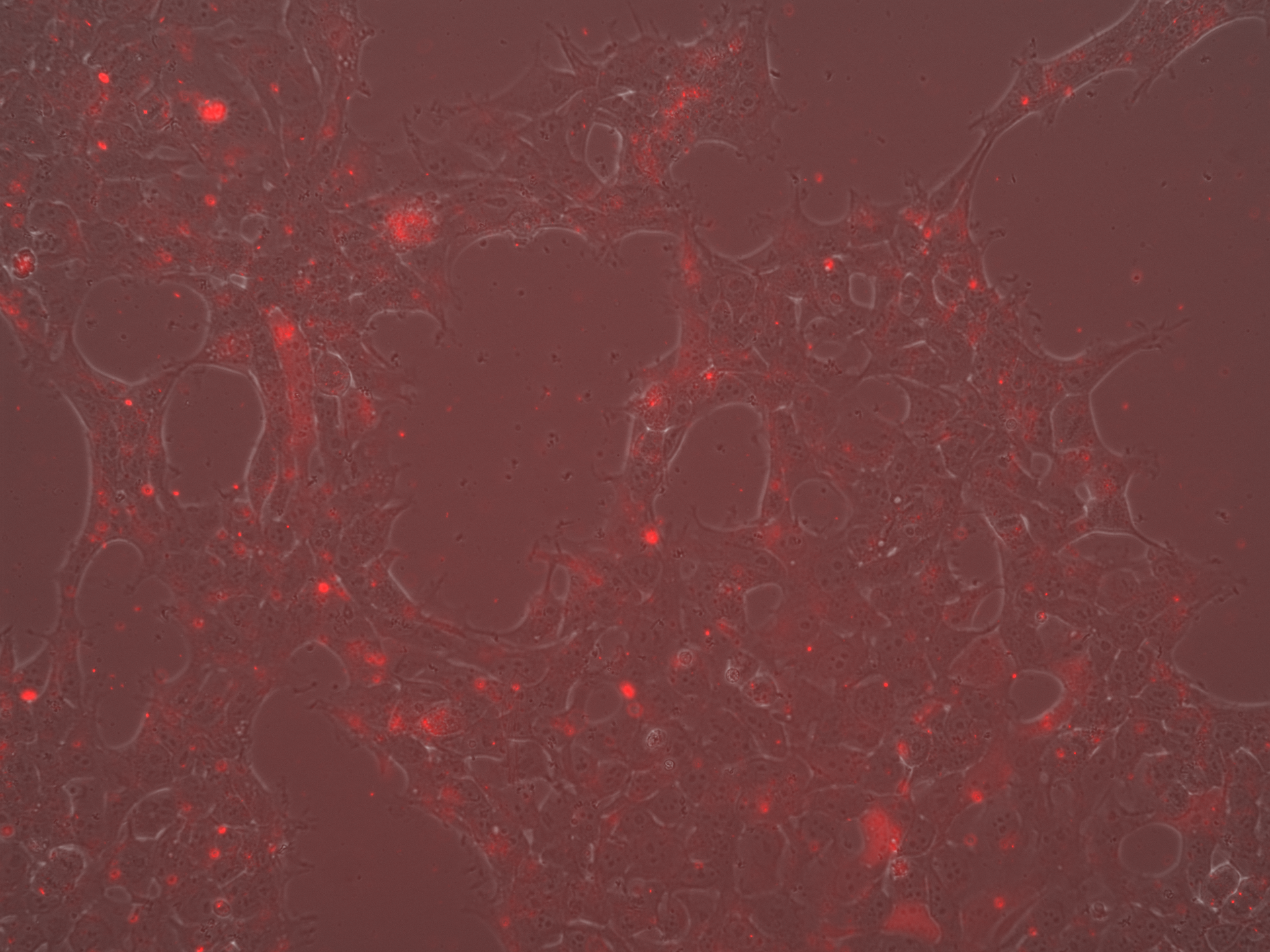

### EV-AAV2 Iodixanol_Overlay_Day5.tif

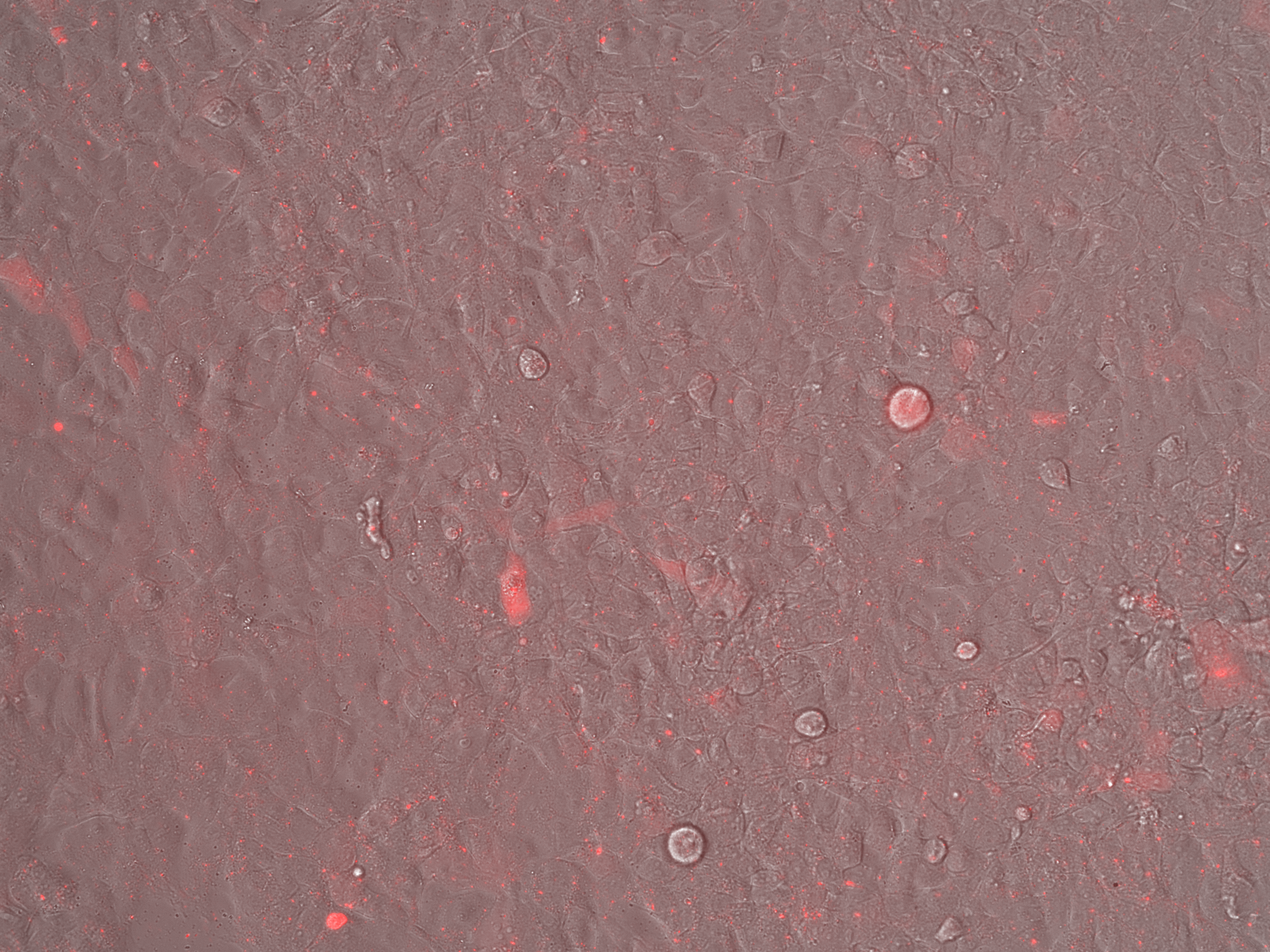

### EV-AAV2 Iodixanol_Overlay_Day5.tif

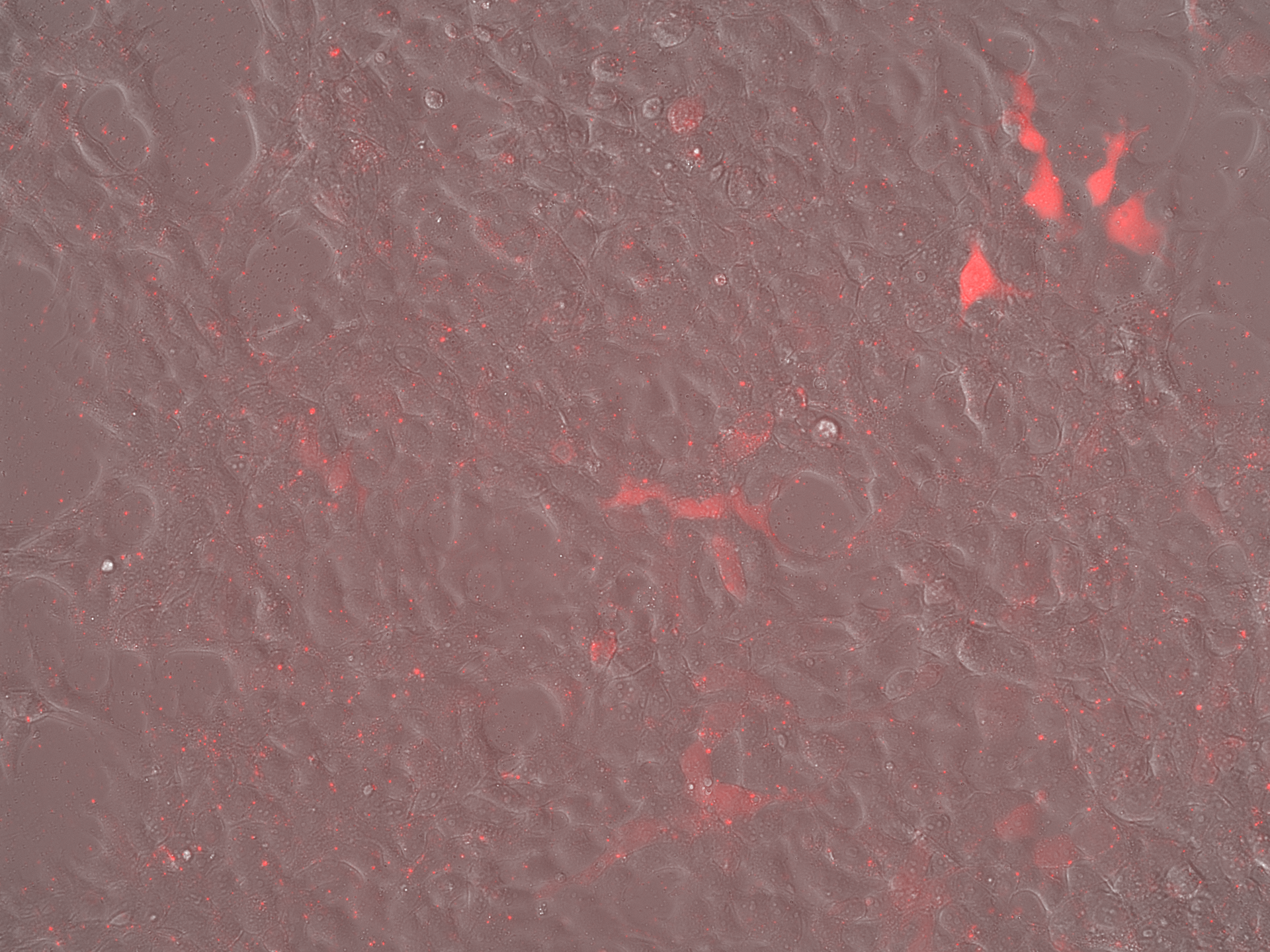

### EV-AAV2 Iodixanol_Overlay_Day7.tif

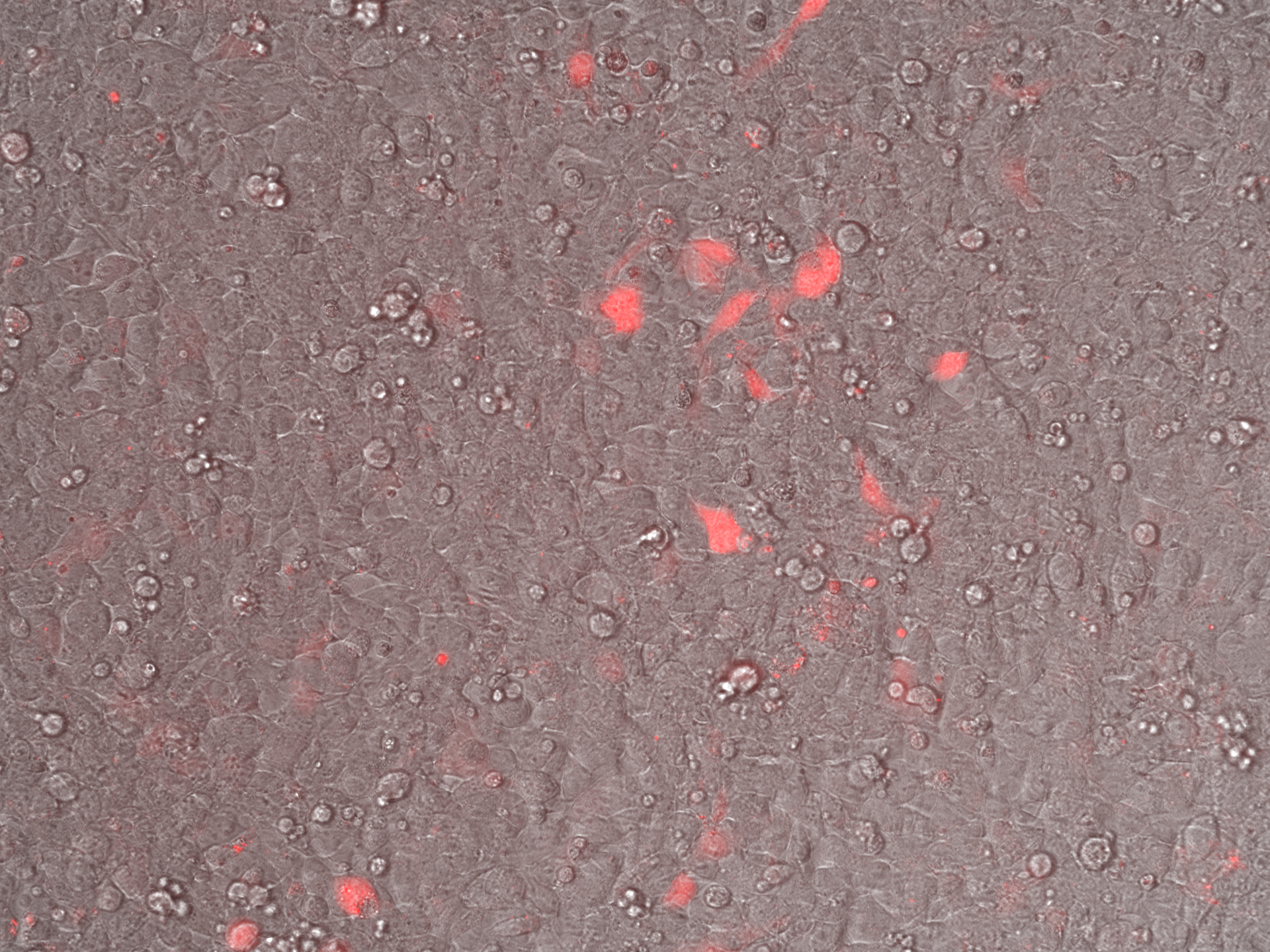

### EV-AAV2 Iodixanol_Overlay_Day7.tif

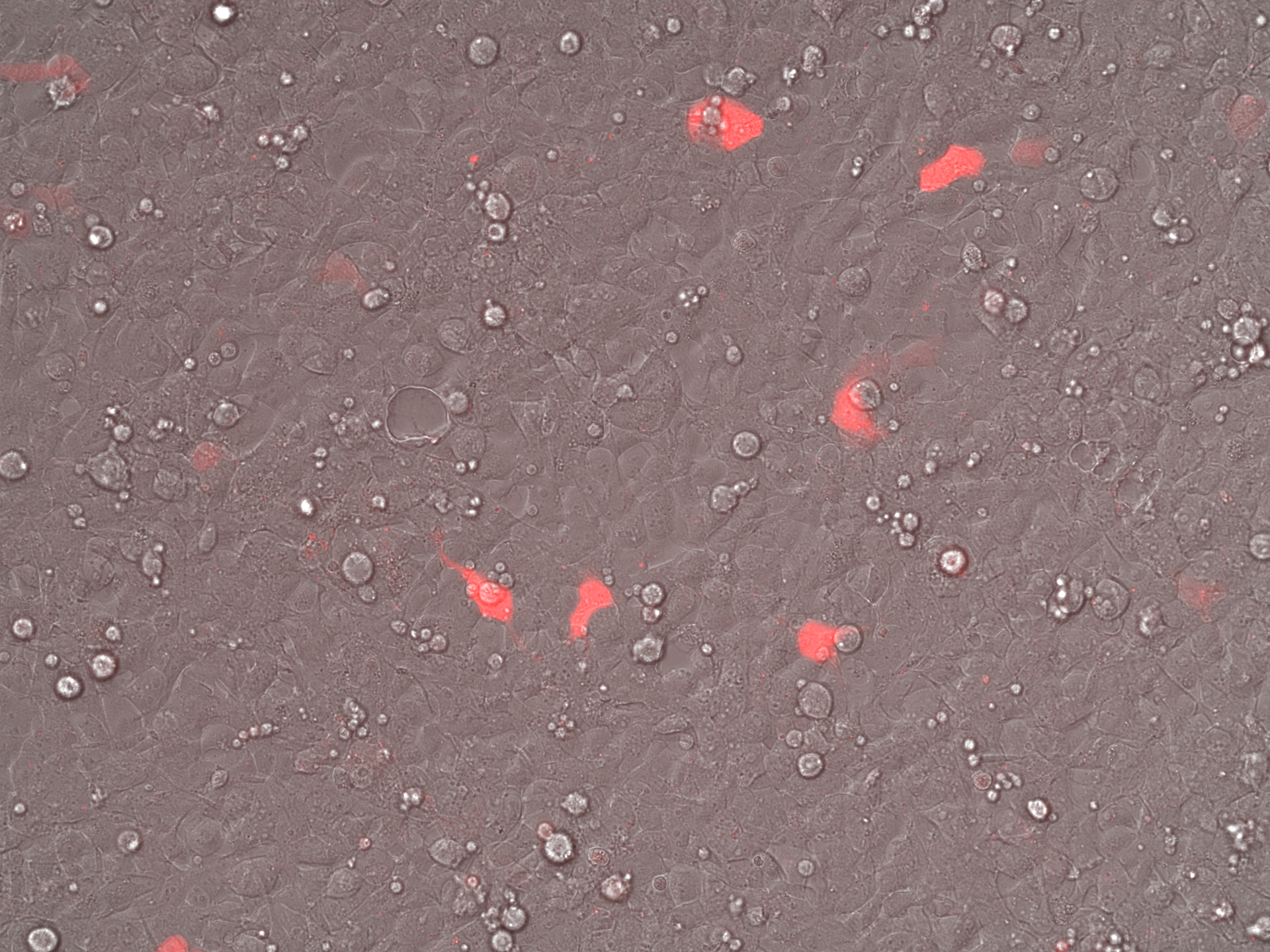

### EV-AAV2 SEC_Brightfield_Day3.tif

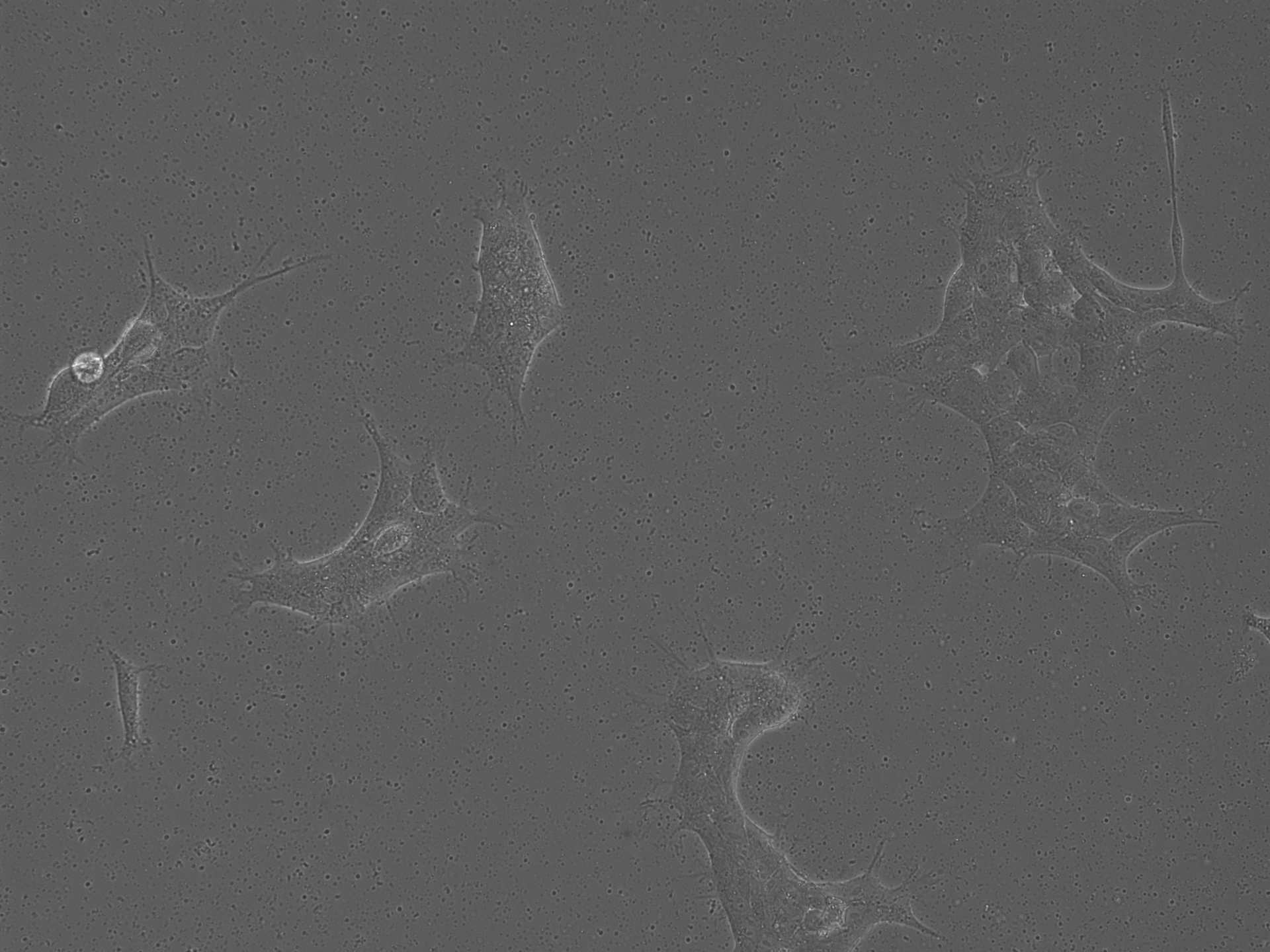

### EV-AAV2 SEC_Brightfield_Day3.tif

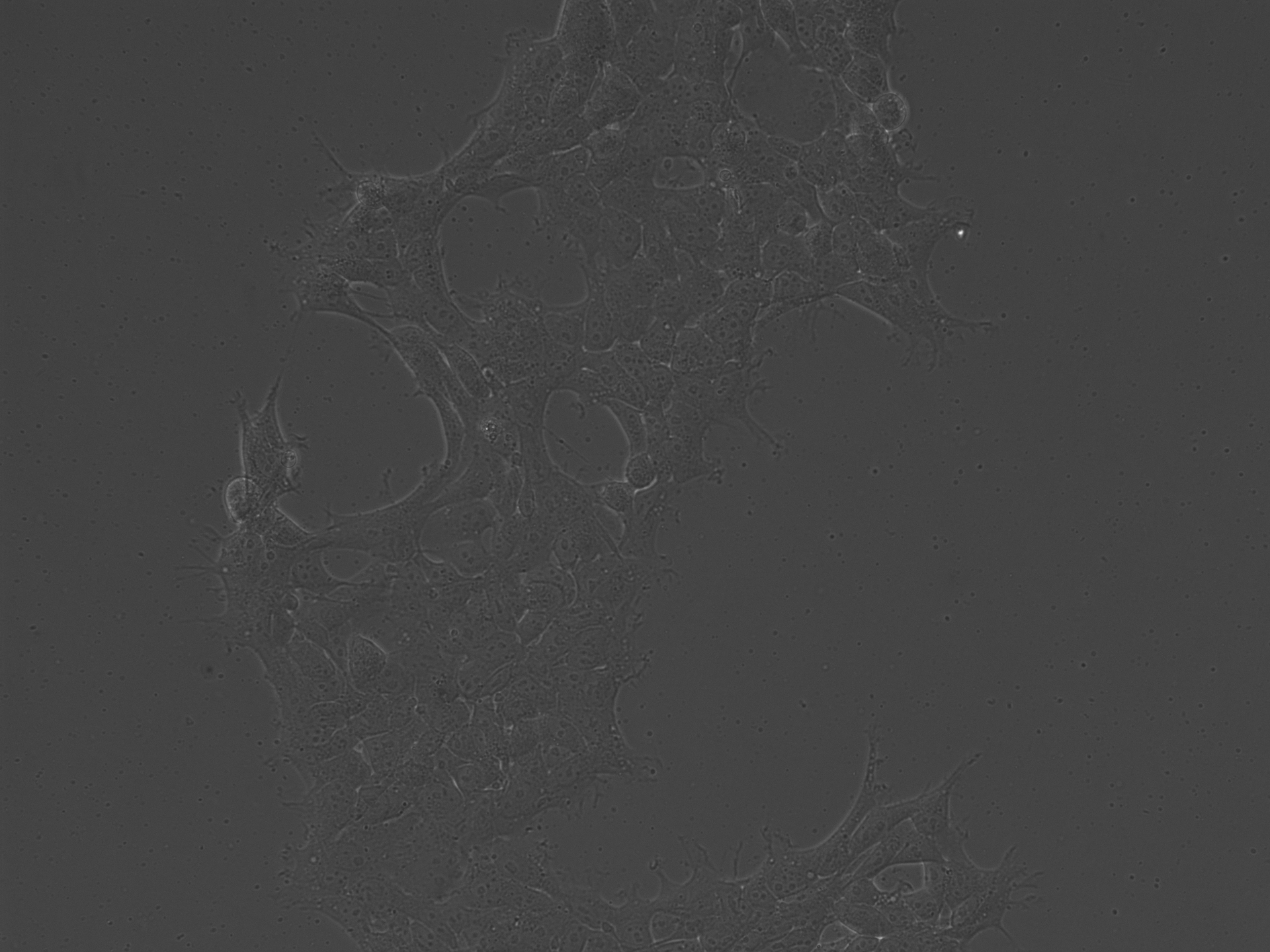

### EV-AAV2 SEC_Brightfield_Day5.tif

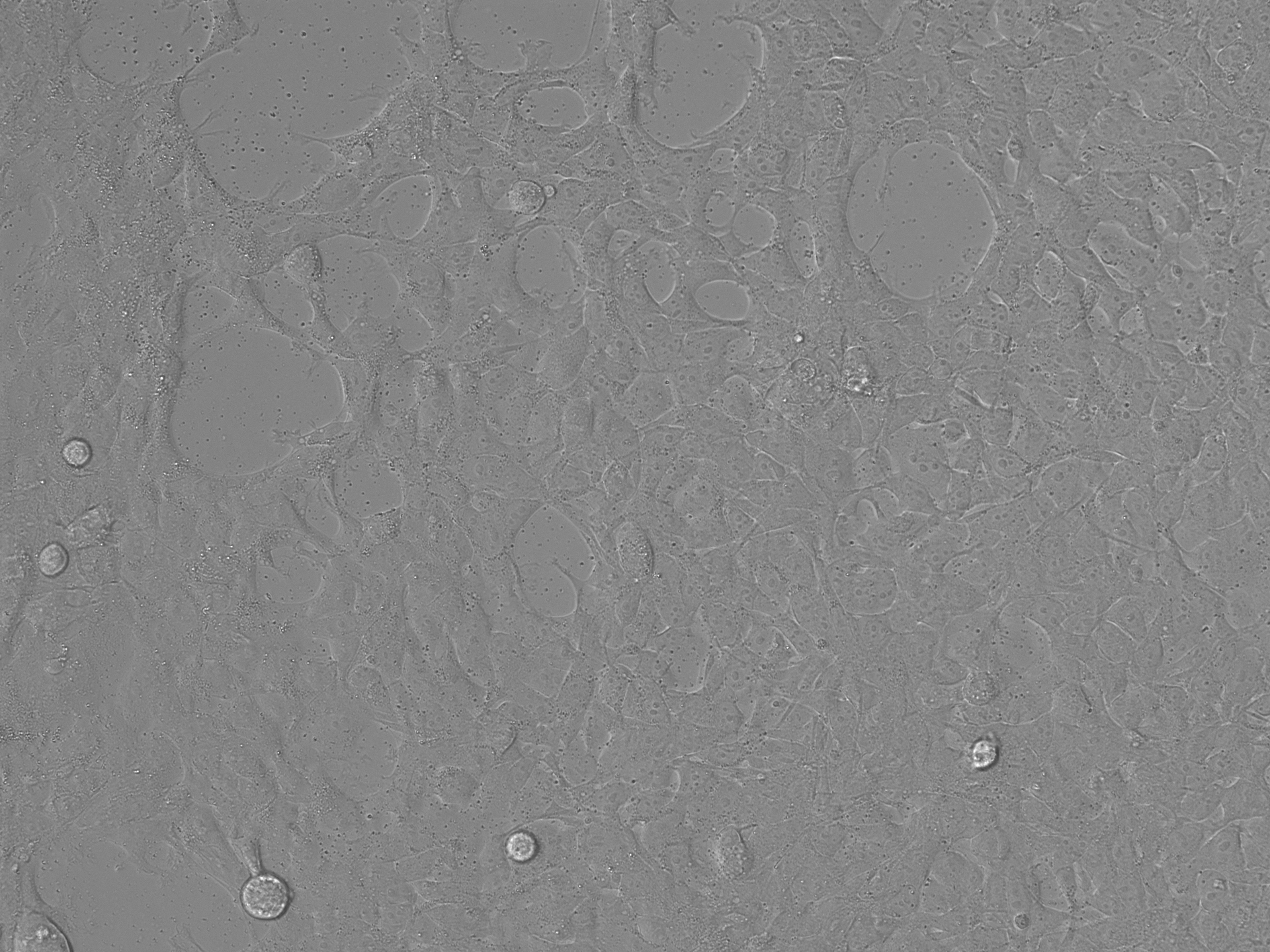

### EV-AAV2 SEC_Brightfield_Day5.tif

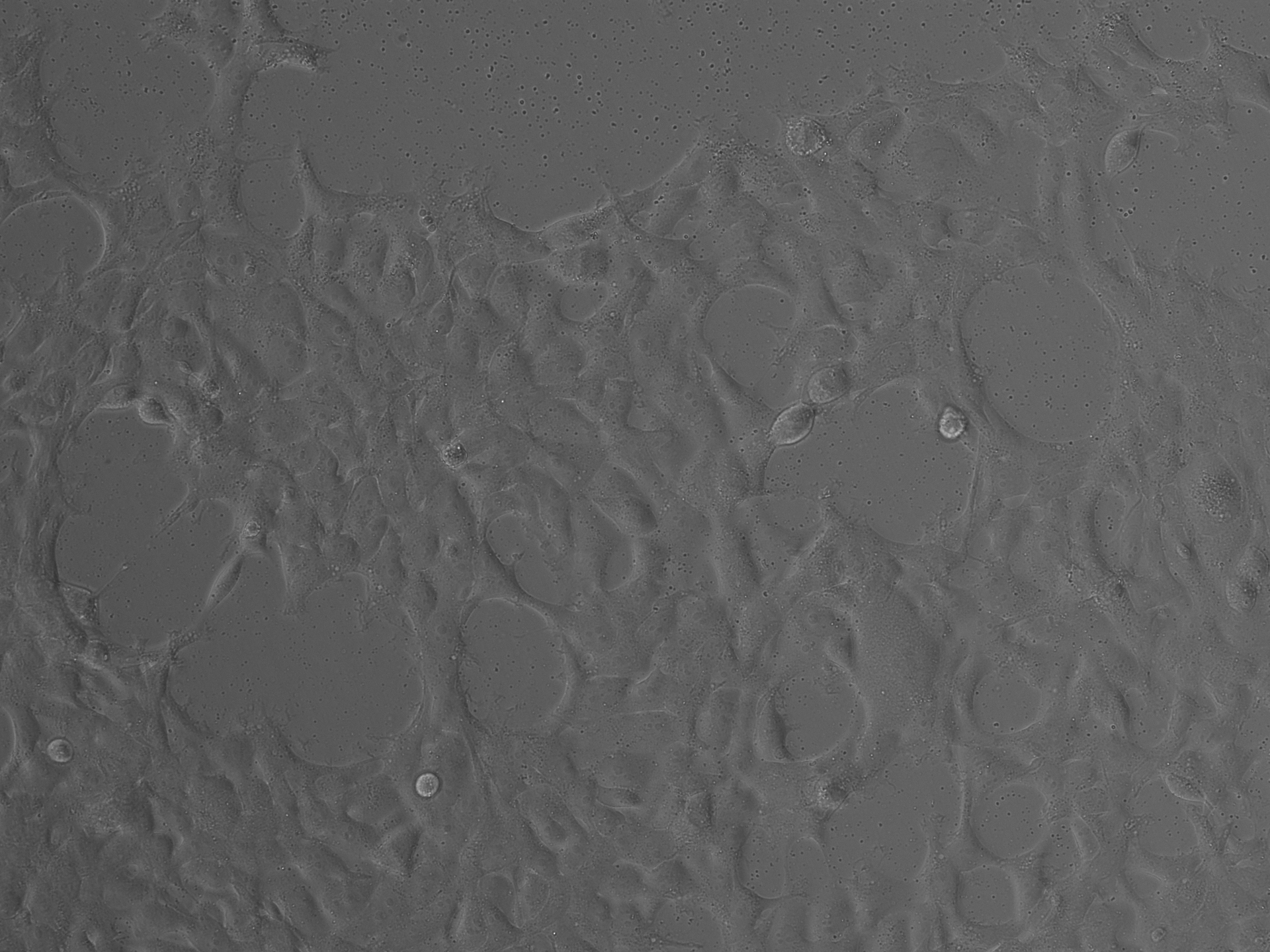

### EV-AAV2 SEC_Brightfield_Day7.tif

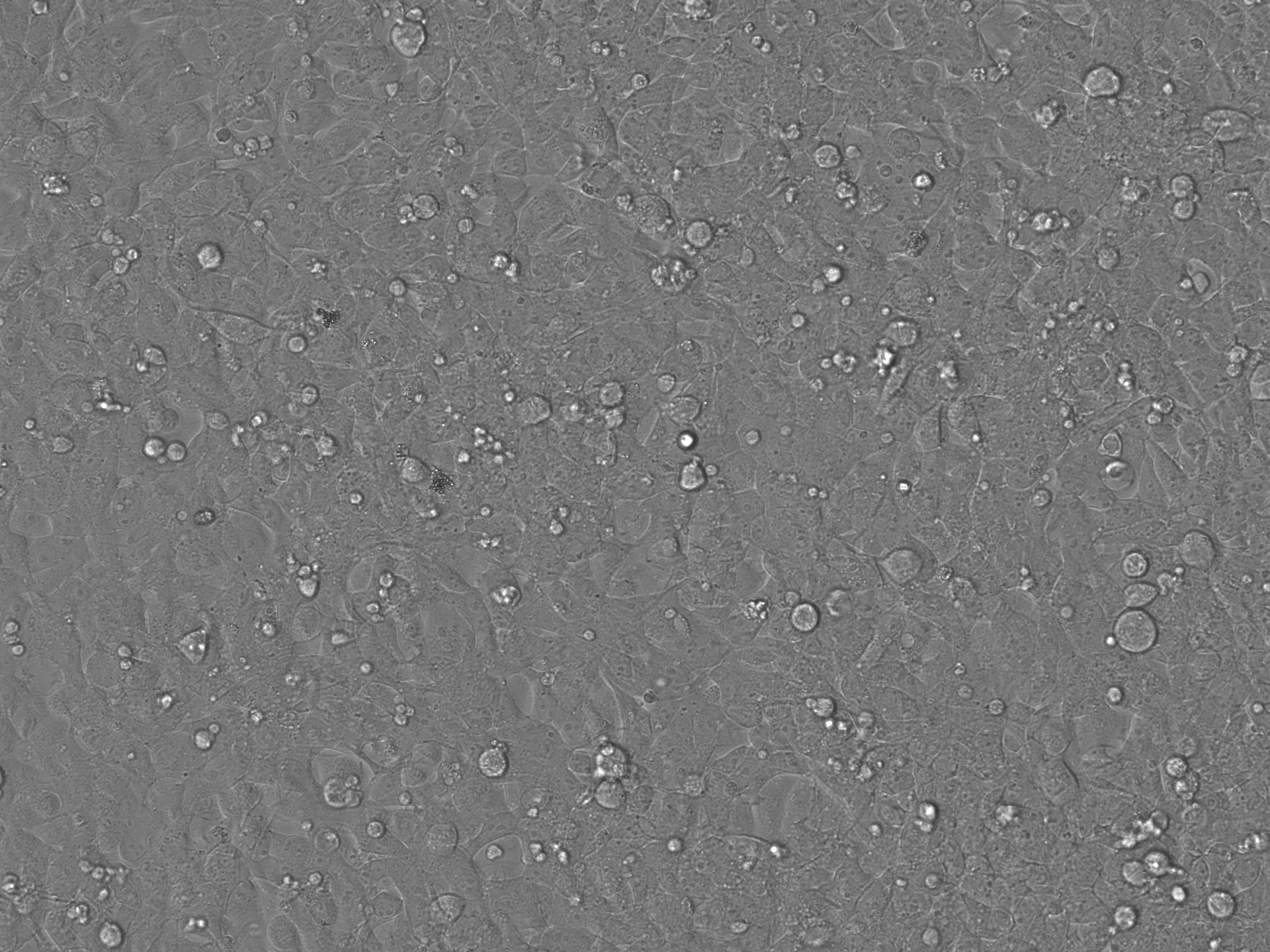

### EV-AAV2 SEC_Brightfield_Day7.tif

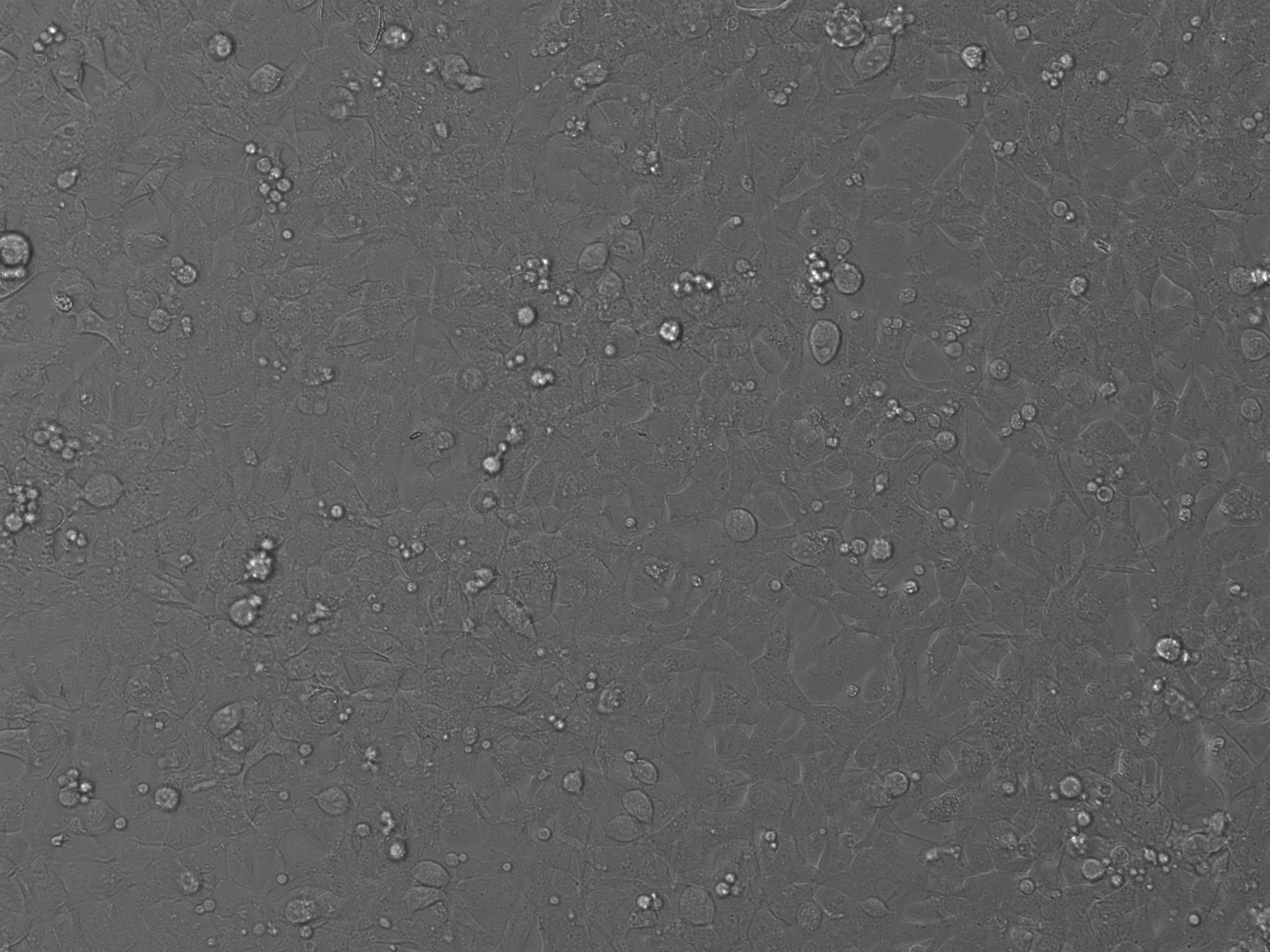

### EV-AAV2 SEC_LumiScarlet_Day3.tif

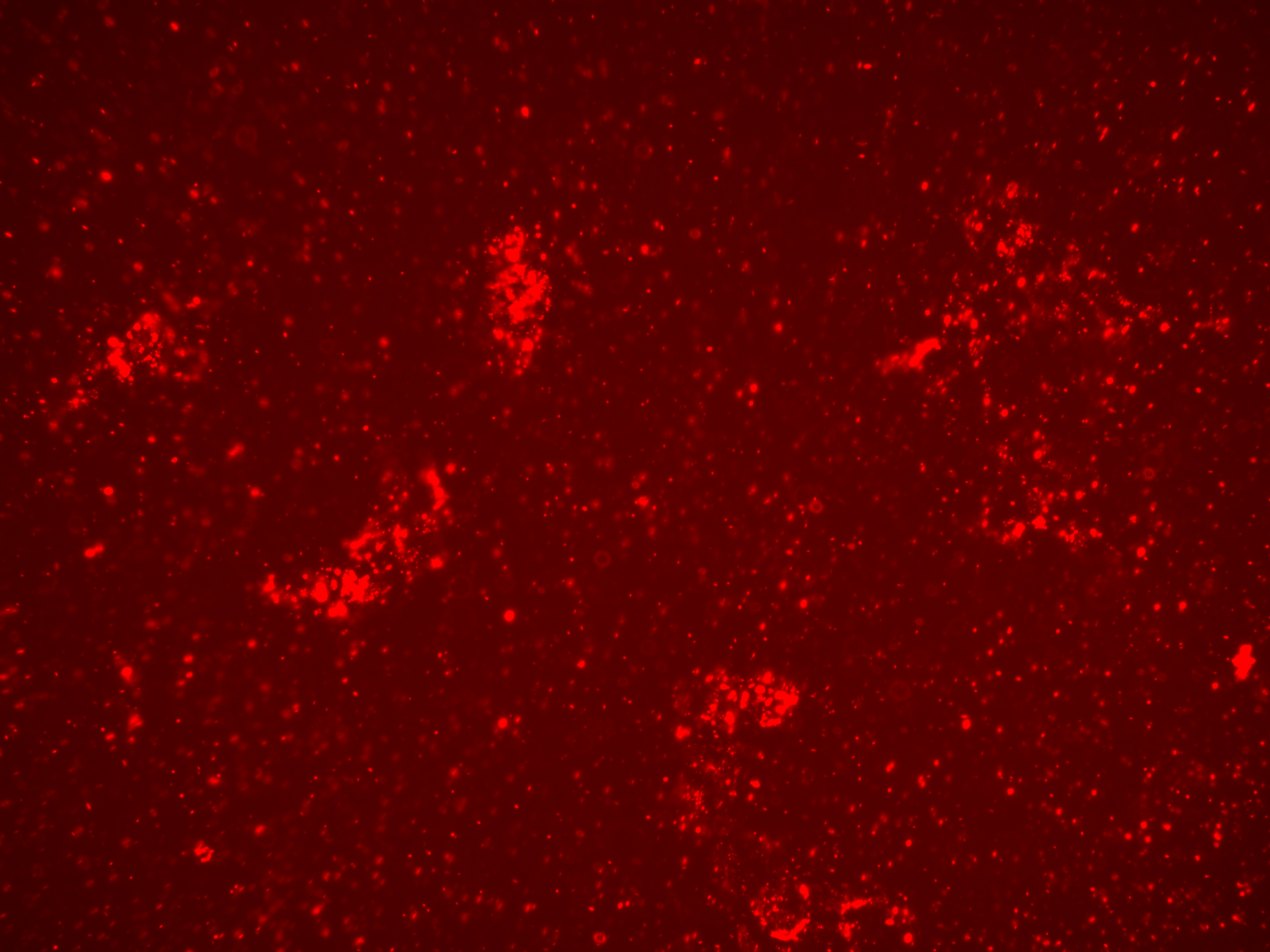

### EV-AAV2 SEC_LumiScarlet_Day3.tif

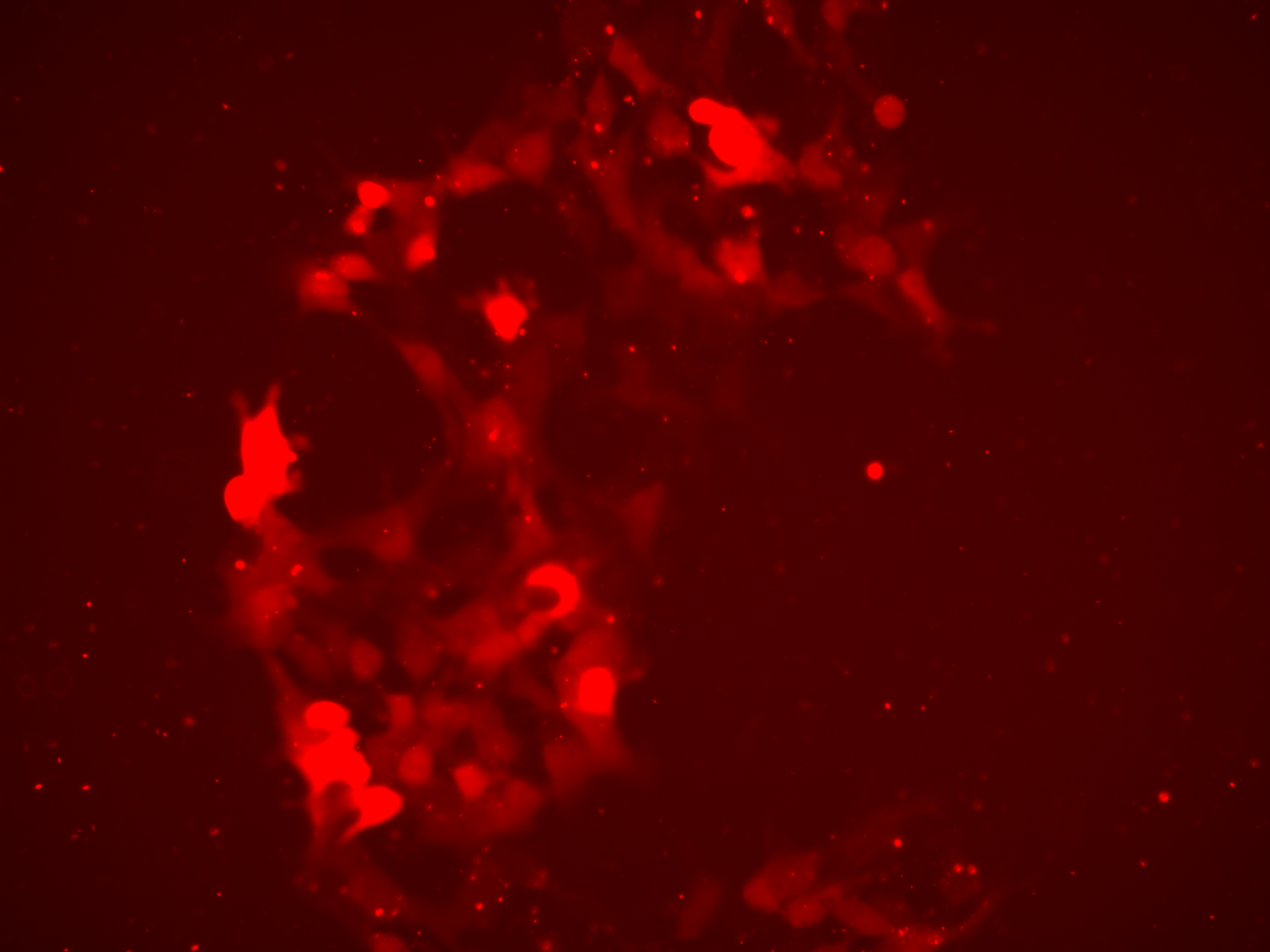

### EV-AAV2 SEC_LumiScarlet_Day5.tif

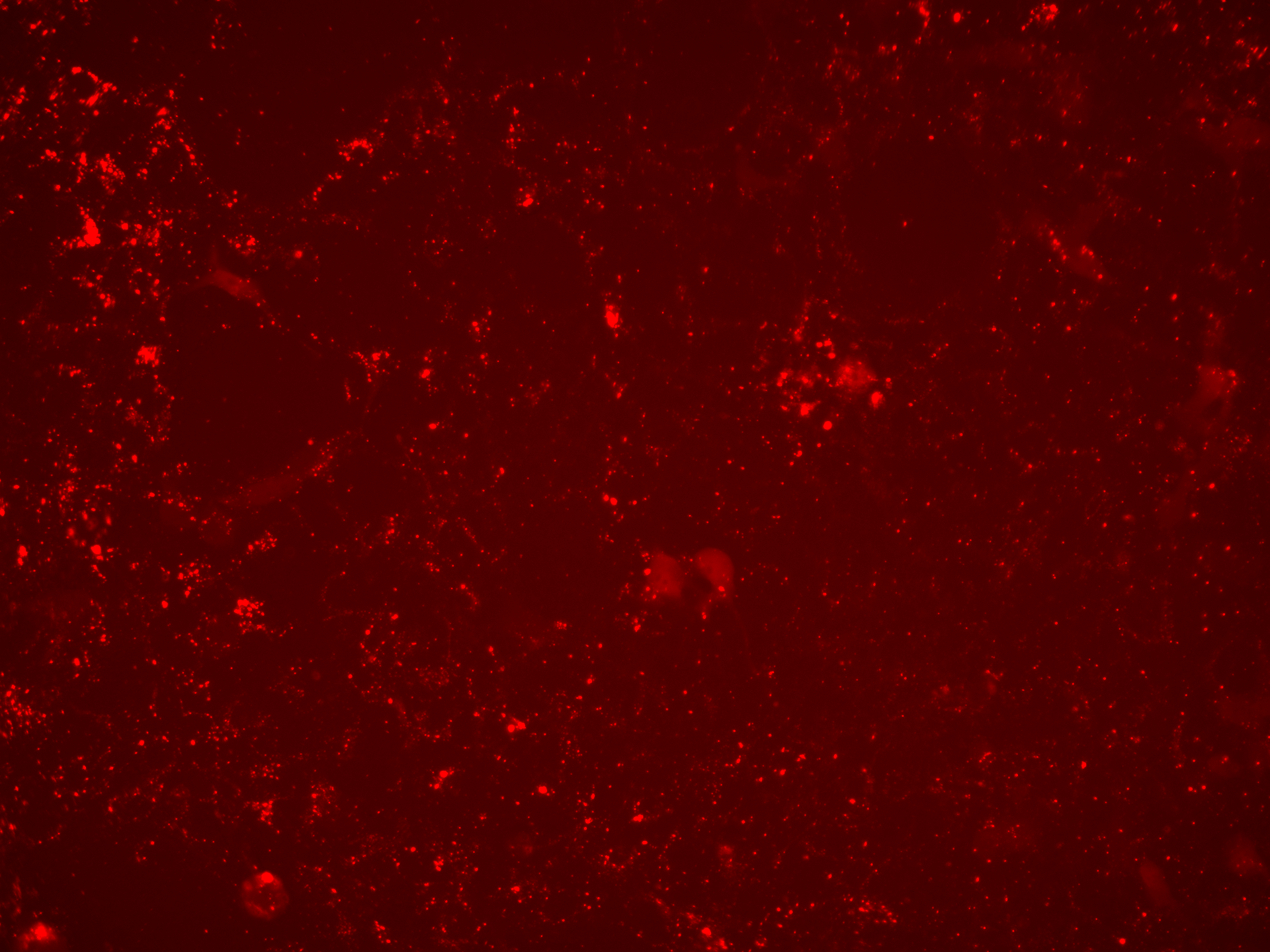

### EV-AAV2 SEC_LumiScarlet_Day5.tif

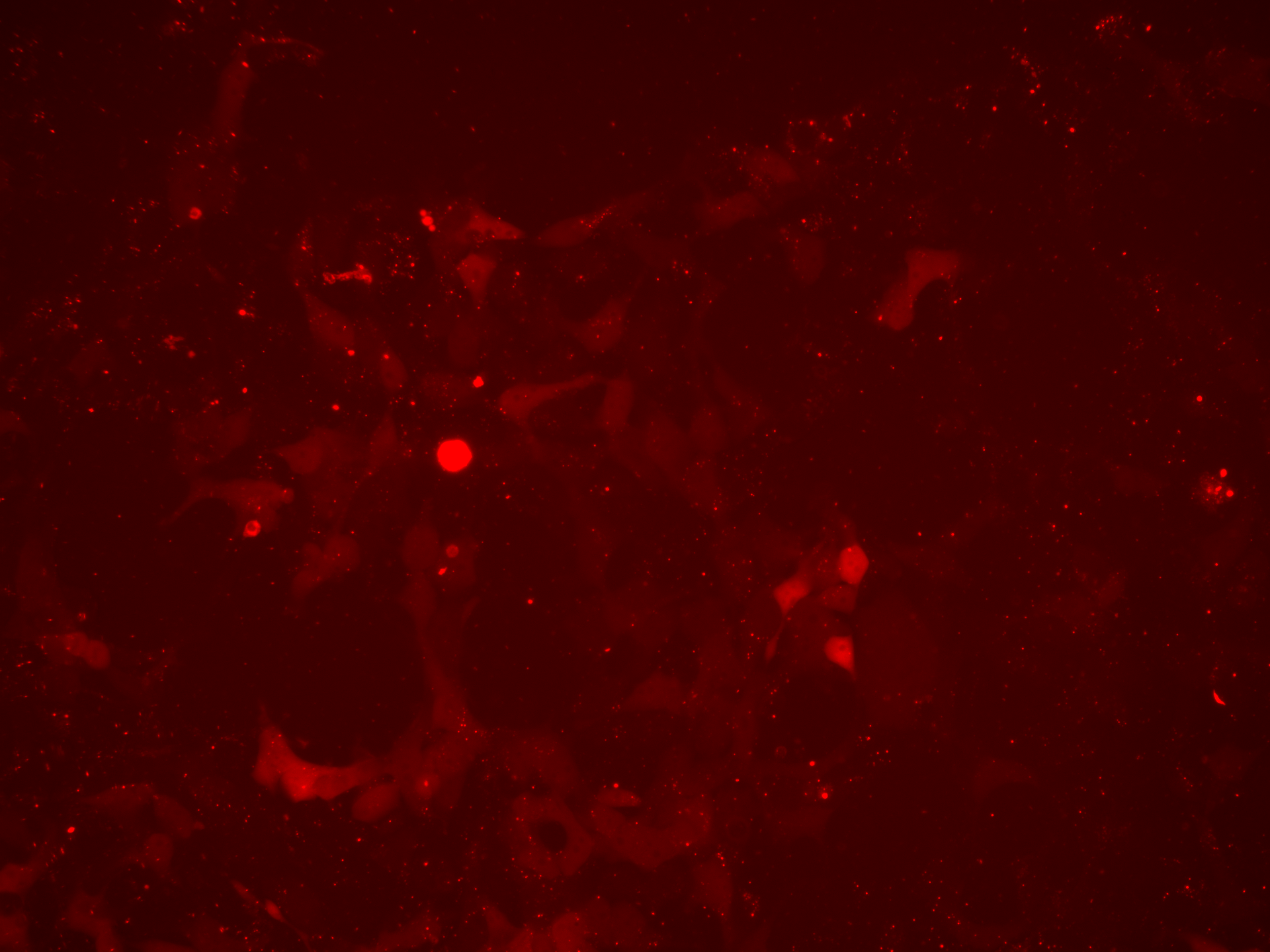

### EV-AAV2 SEC_LumiScarlet_Day7.tif

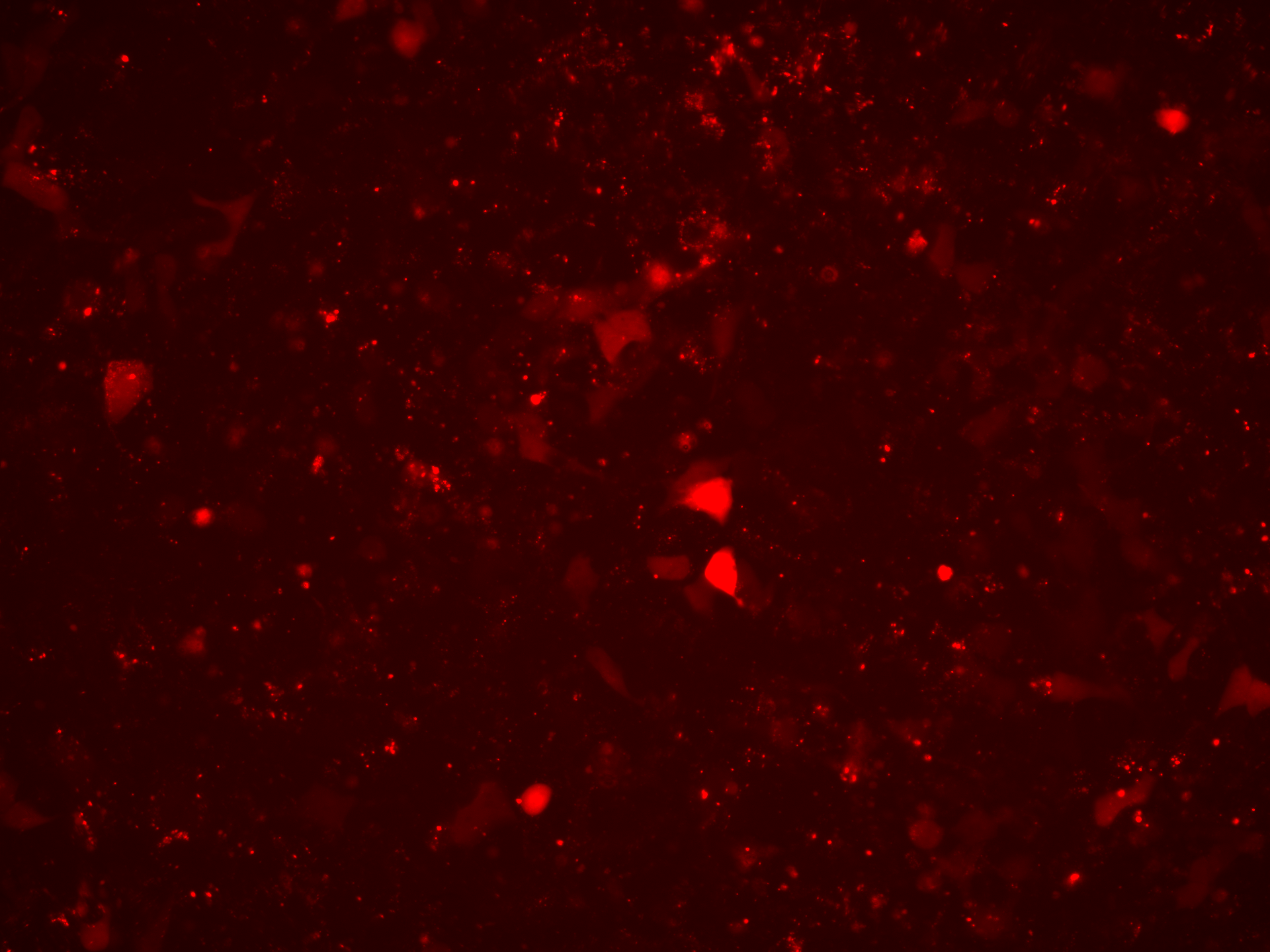

### EV-AAV2 SEC_LumiScarlet_Day7.tif

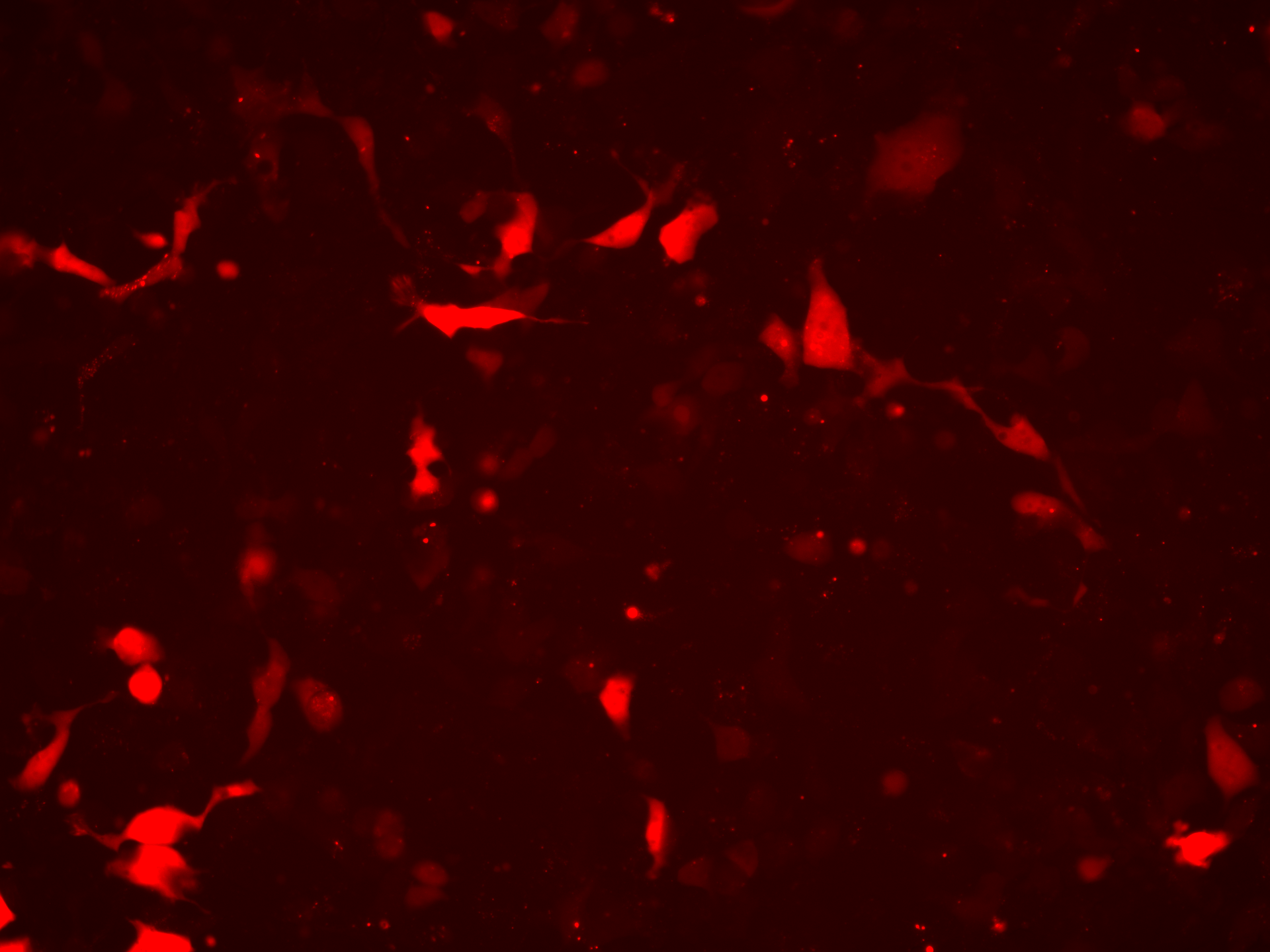

### EV-AAV2 SEC_Overlay_Day3.tif

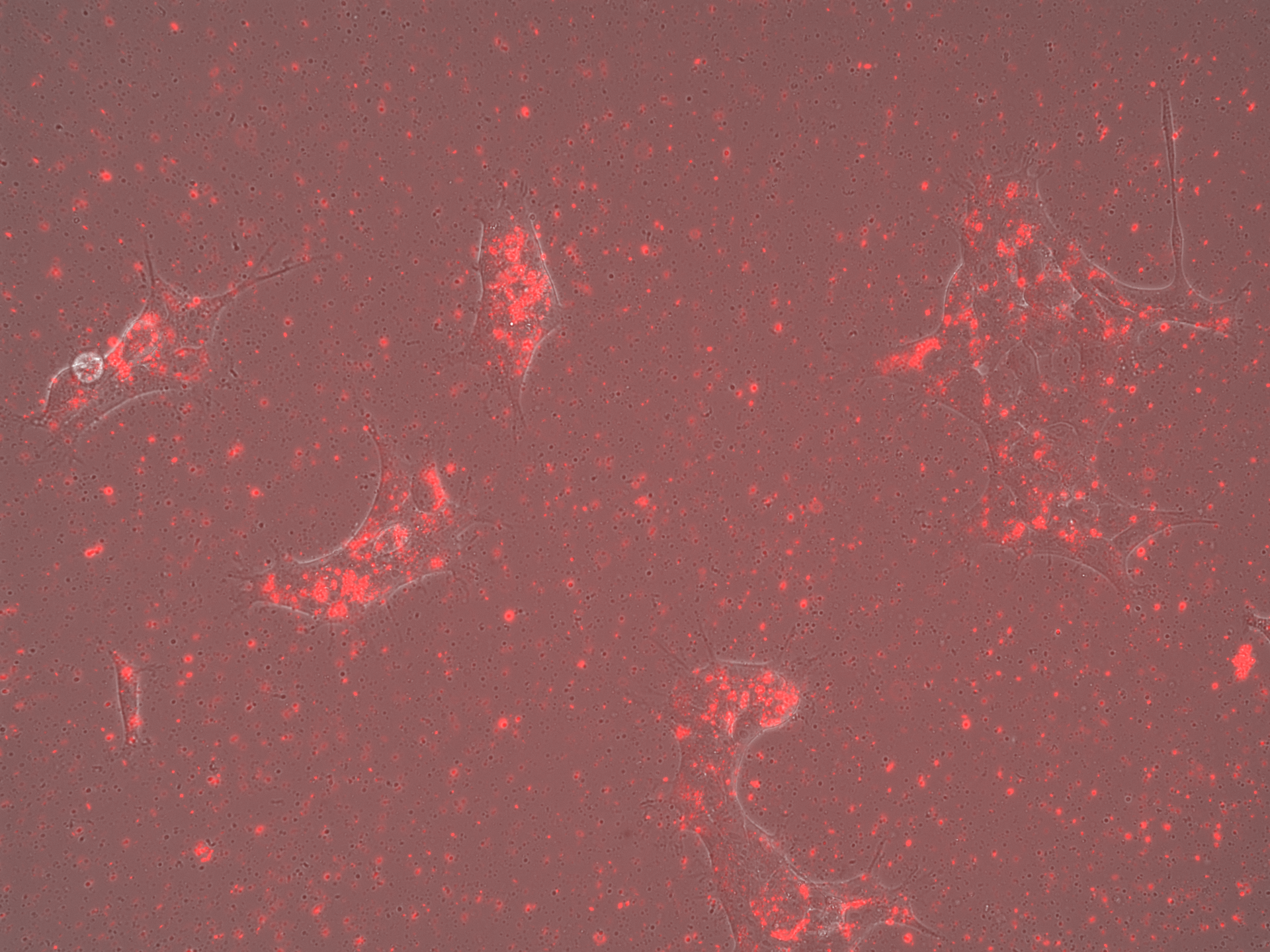

### EV-AAV2 SEC_Overlay_Day3.tif

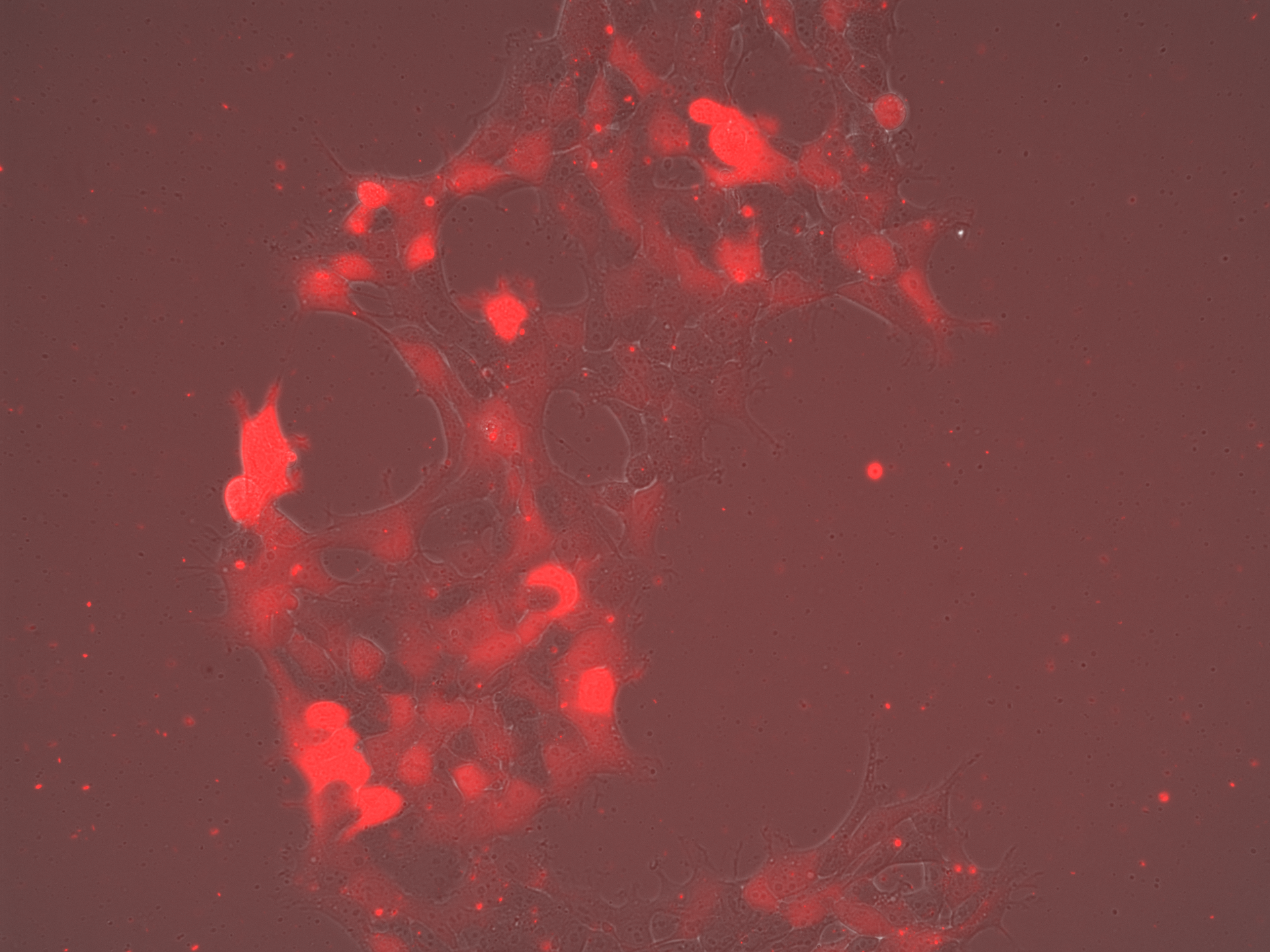

### EV-AAV2 SEC_Overlay_Day5.tif

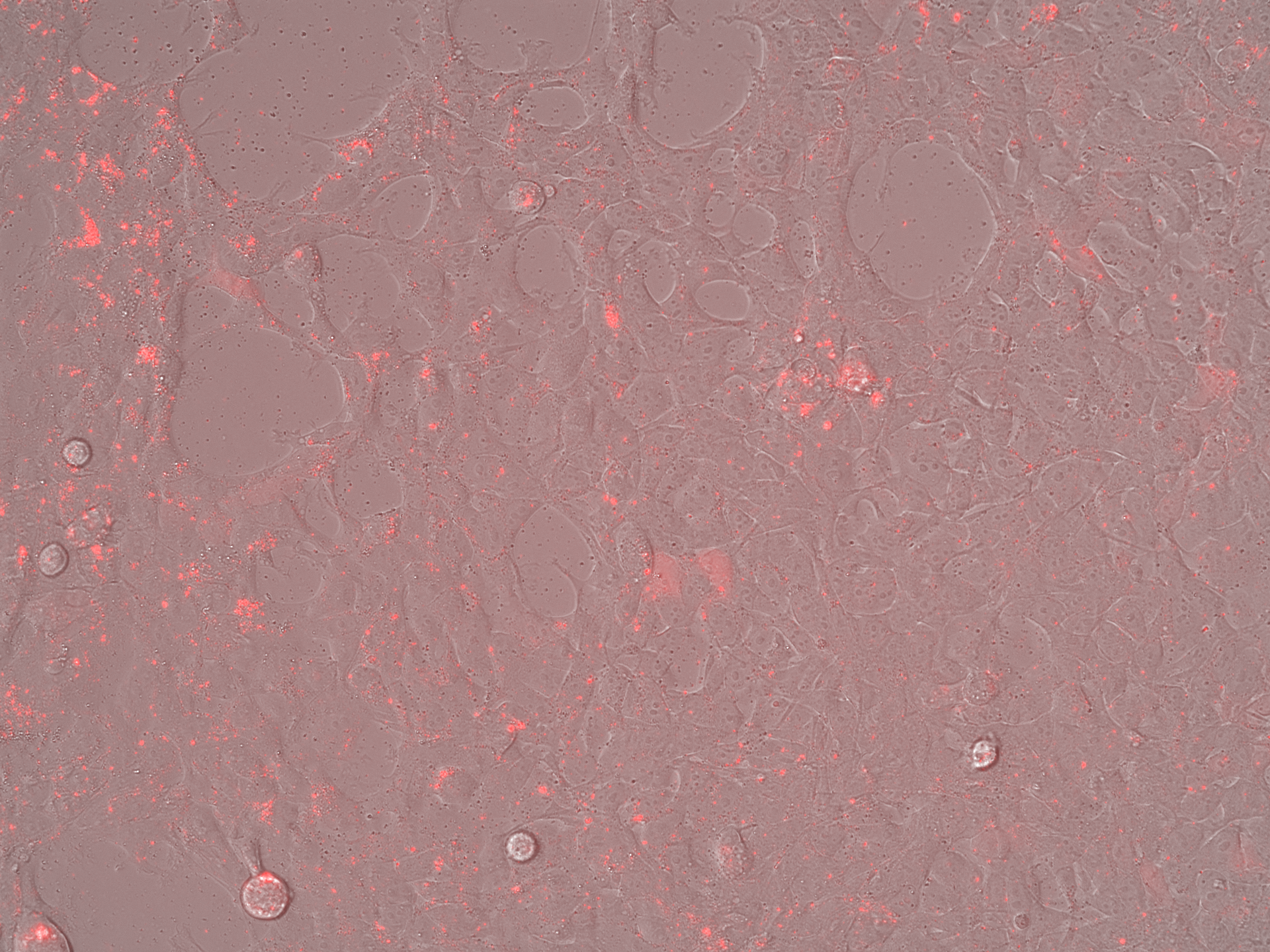

### EV-AAV2 SEC_Overlay_Day5.tif

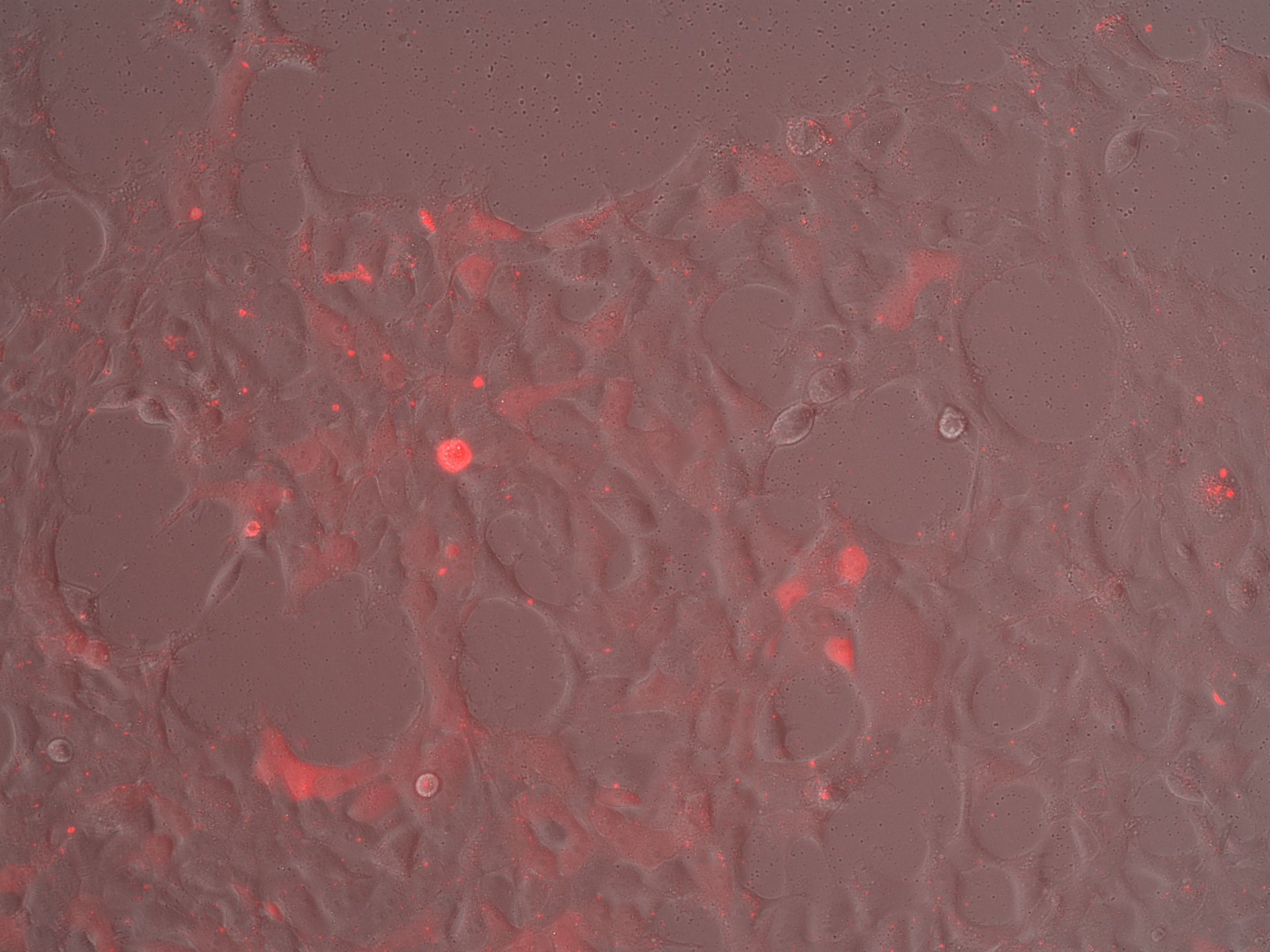

### EV-AAV2 SEC_Overlay_Day7.tif

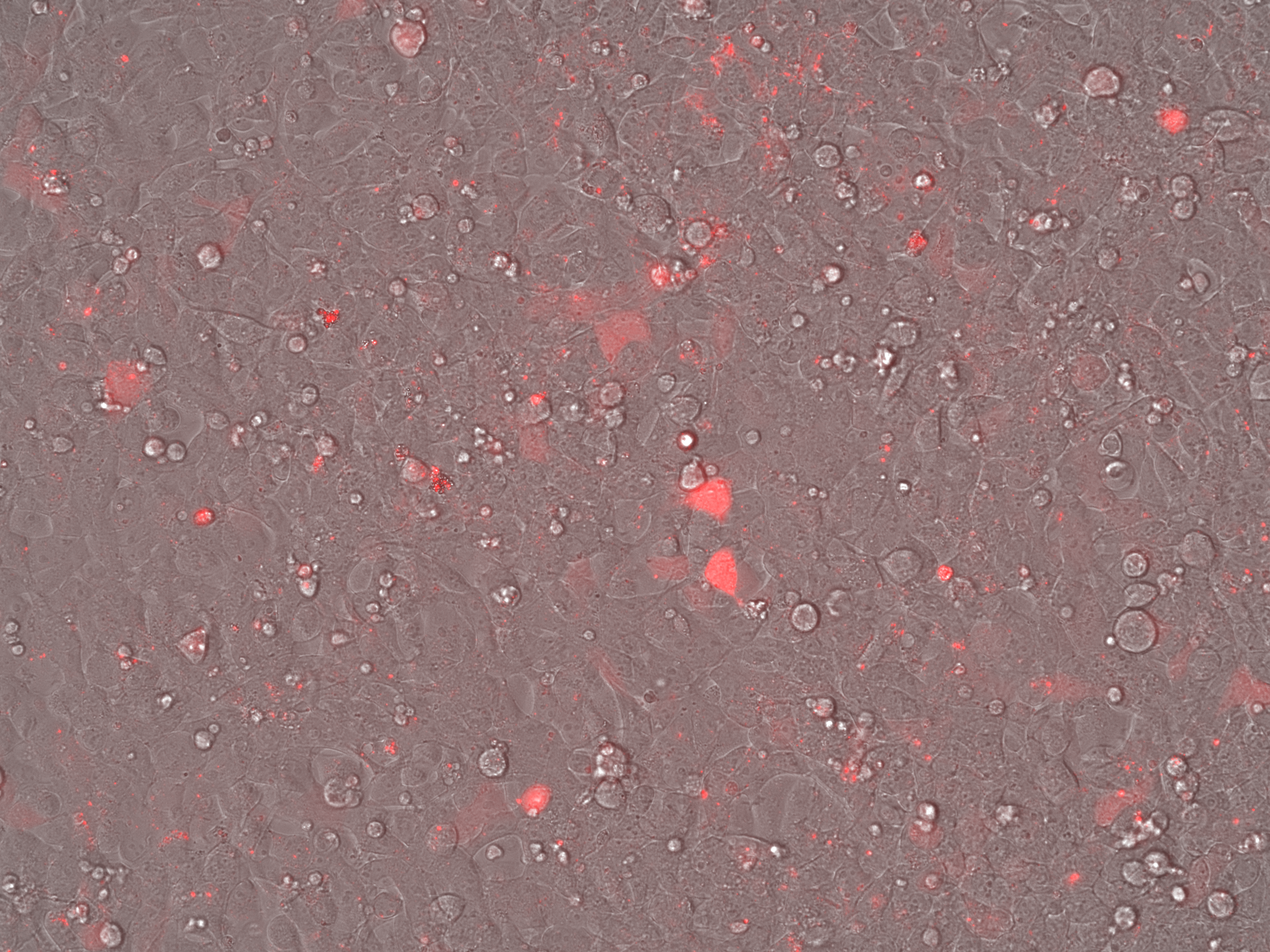

### EV-AAV2 SEC_Overlay_Day7.tif

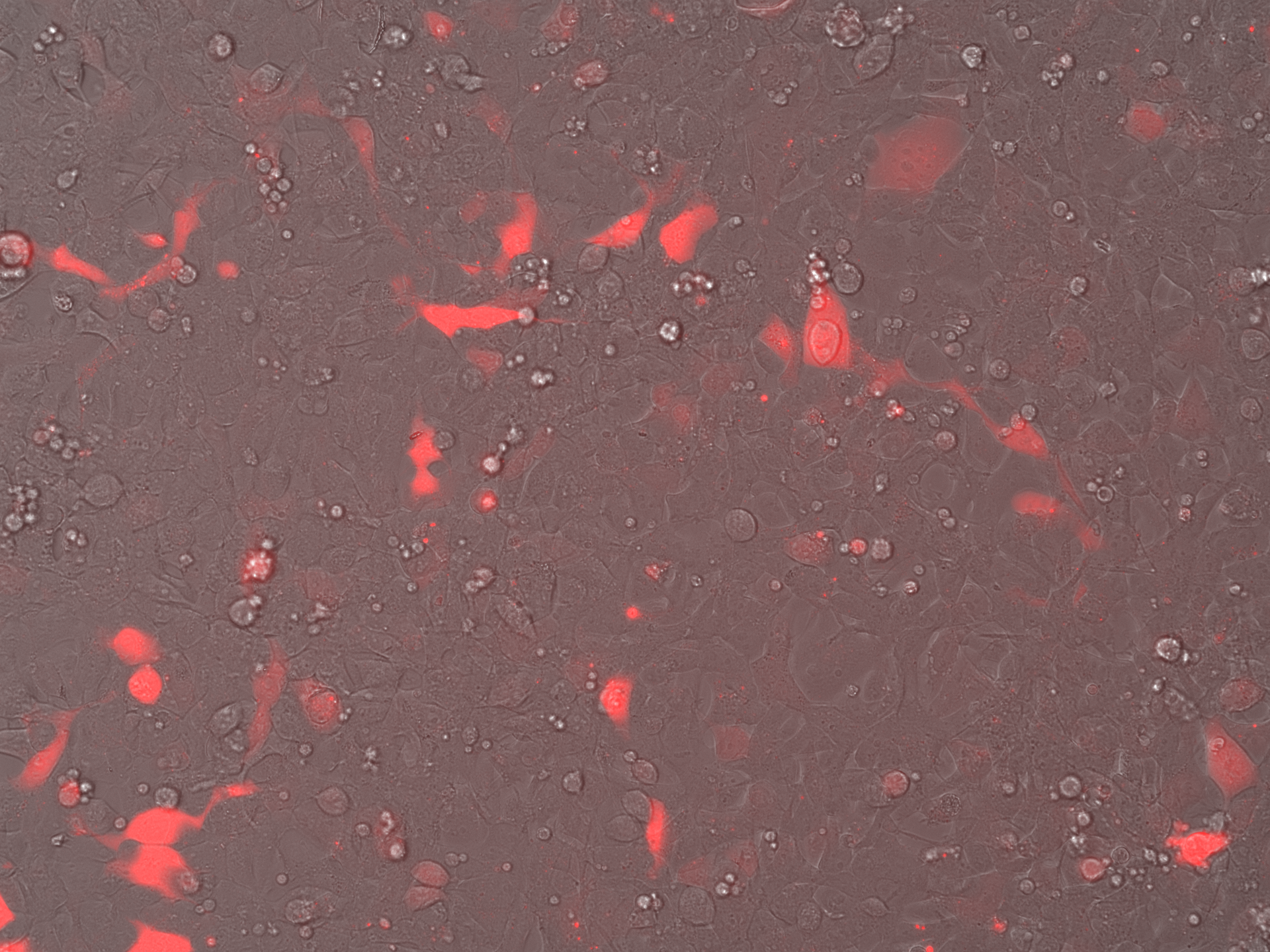

### Free AAV2_Brightfield_Day3.tif

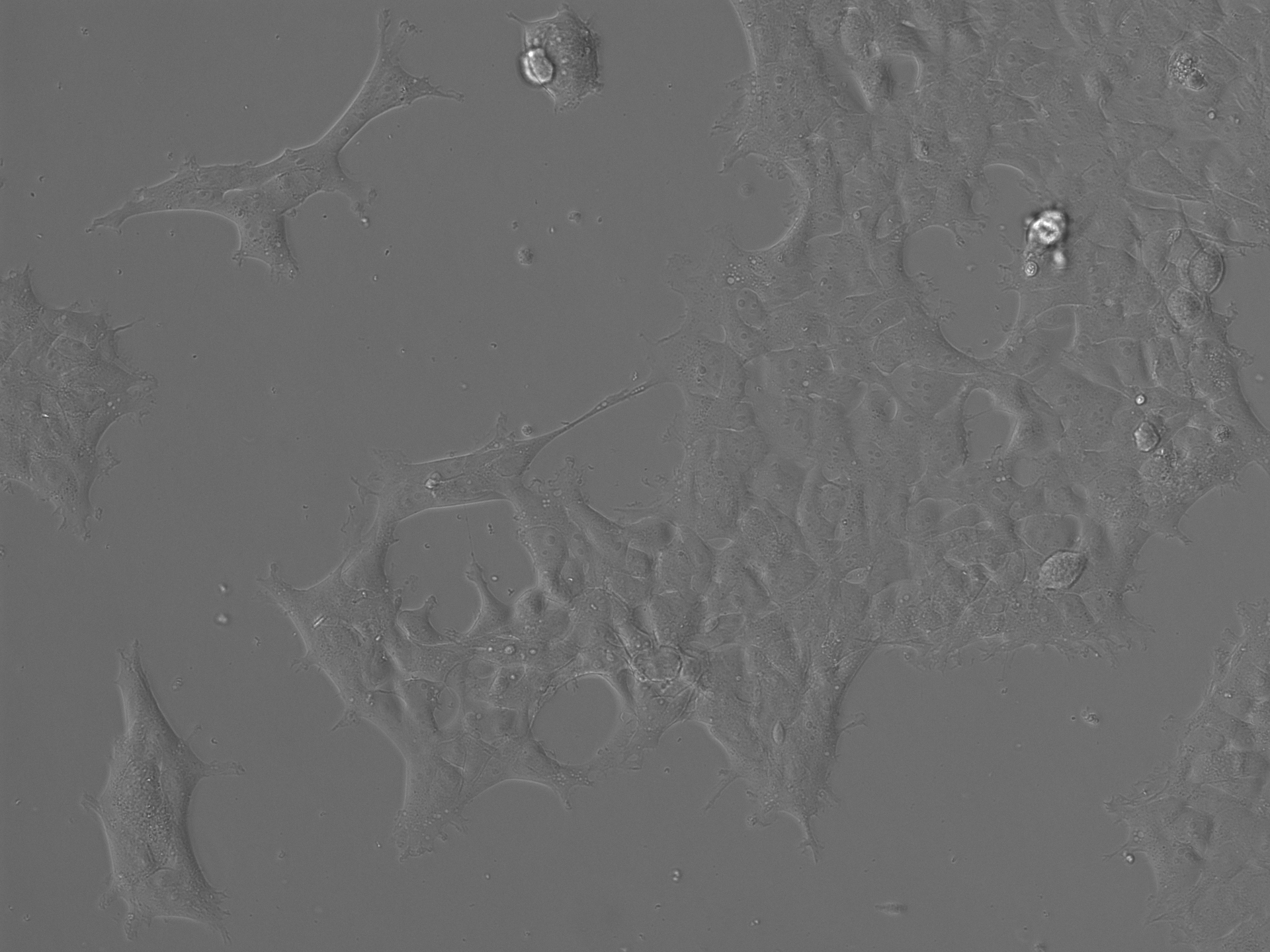

### Free AAV2_Brightfield_Day3.tif

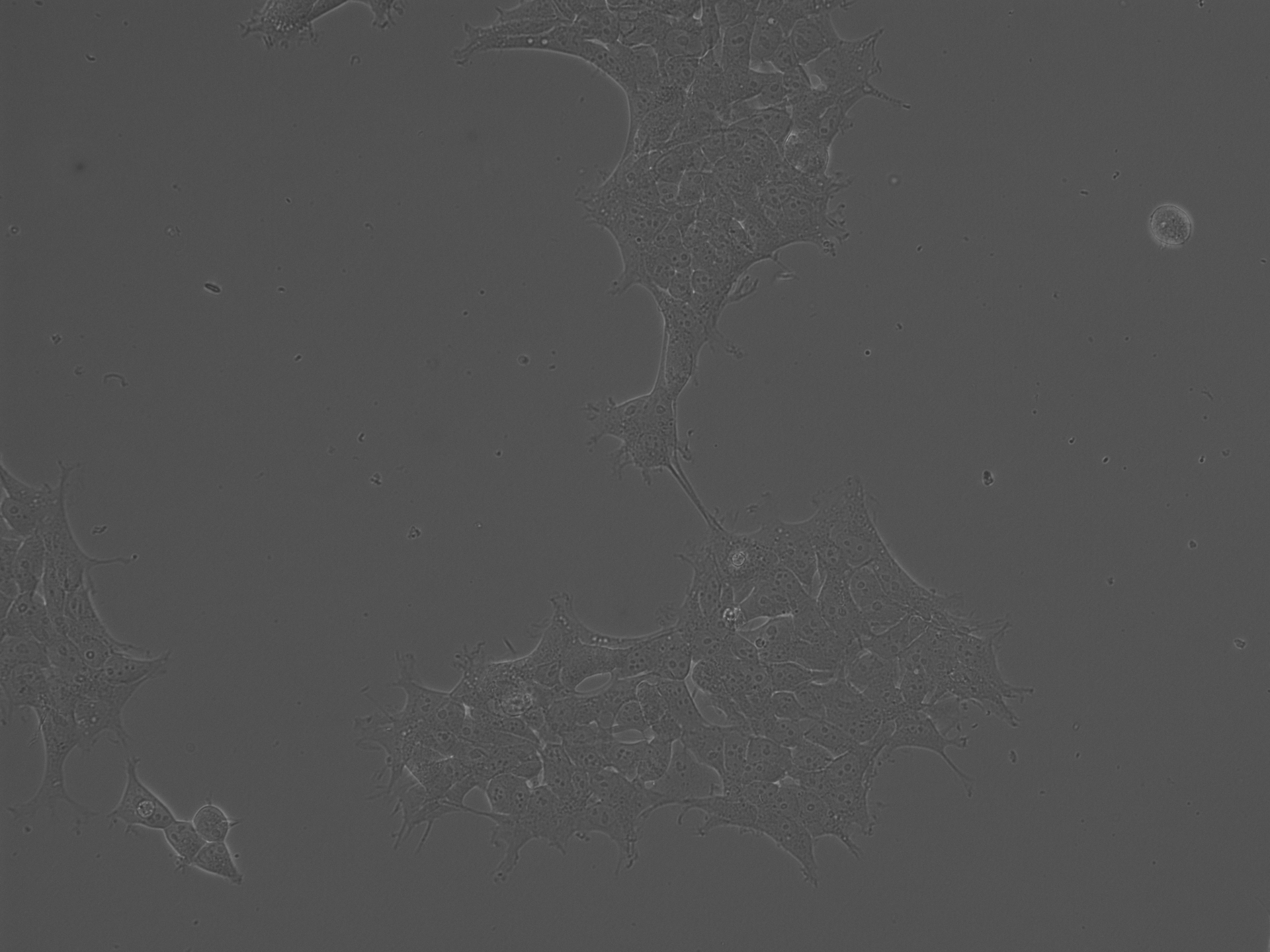

### Free AAV2_Brightfield_Day5.tif

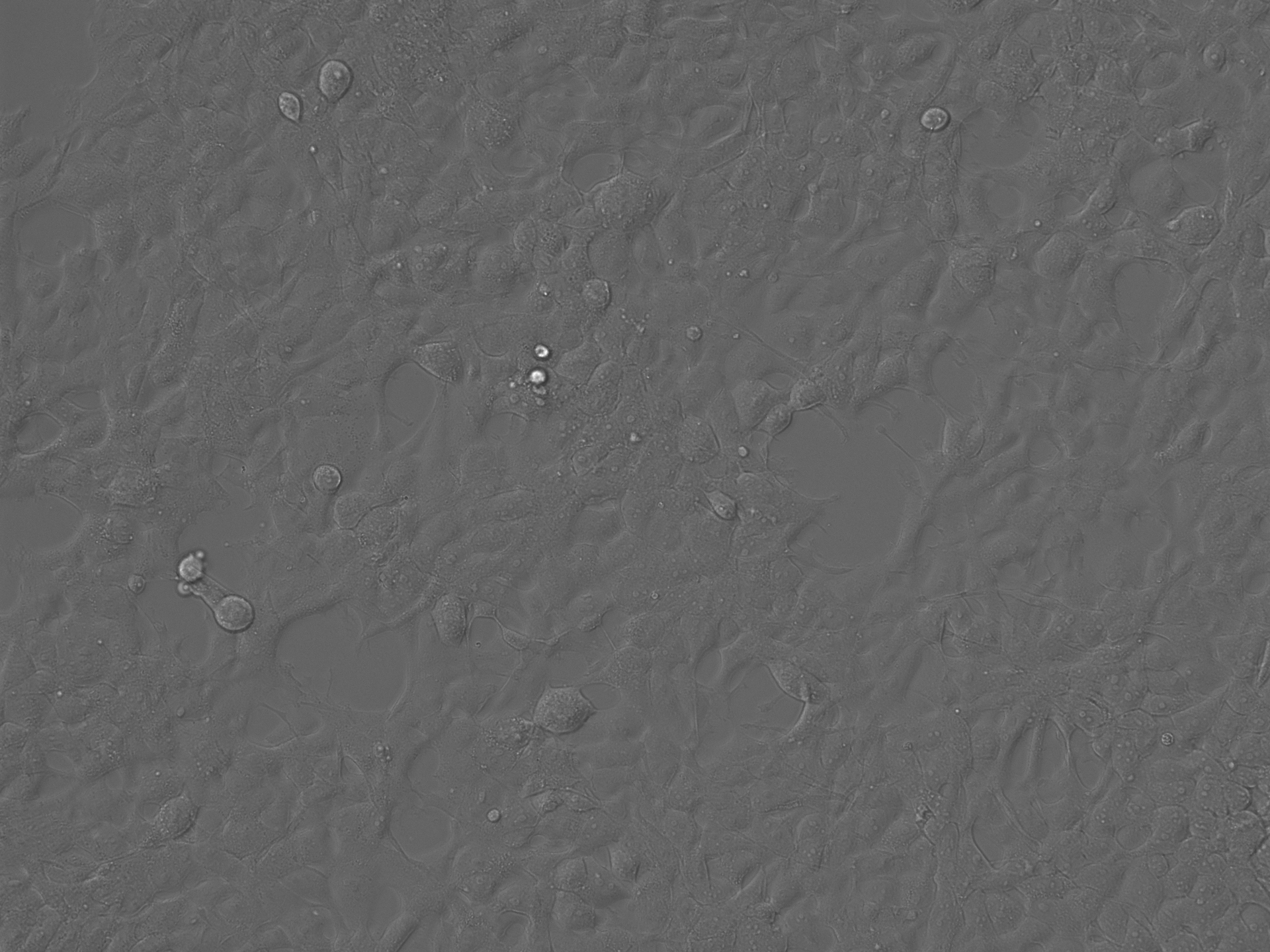

### Free AAV2_Brightfield_Day5.tif

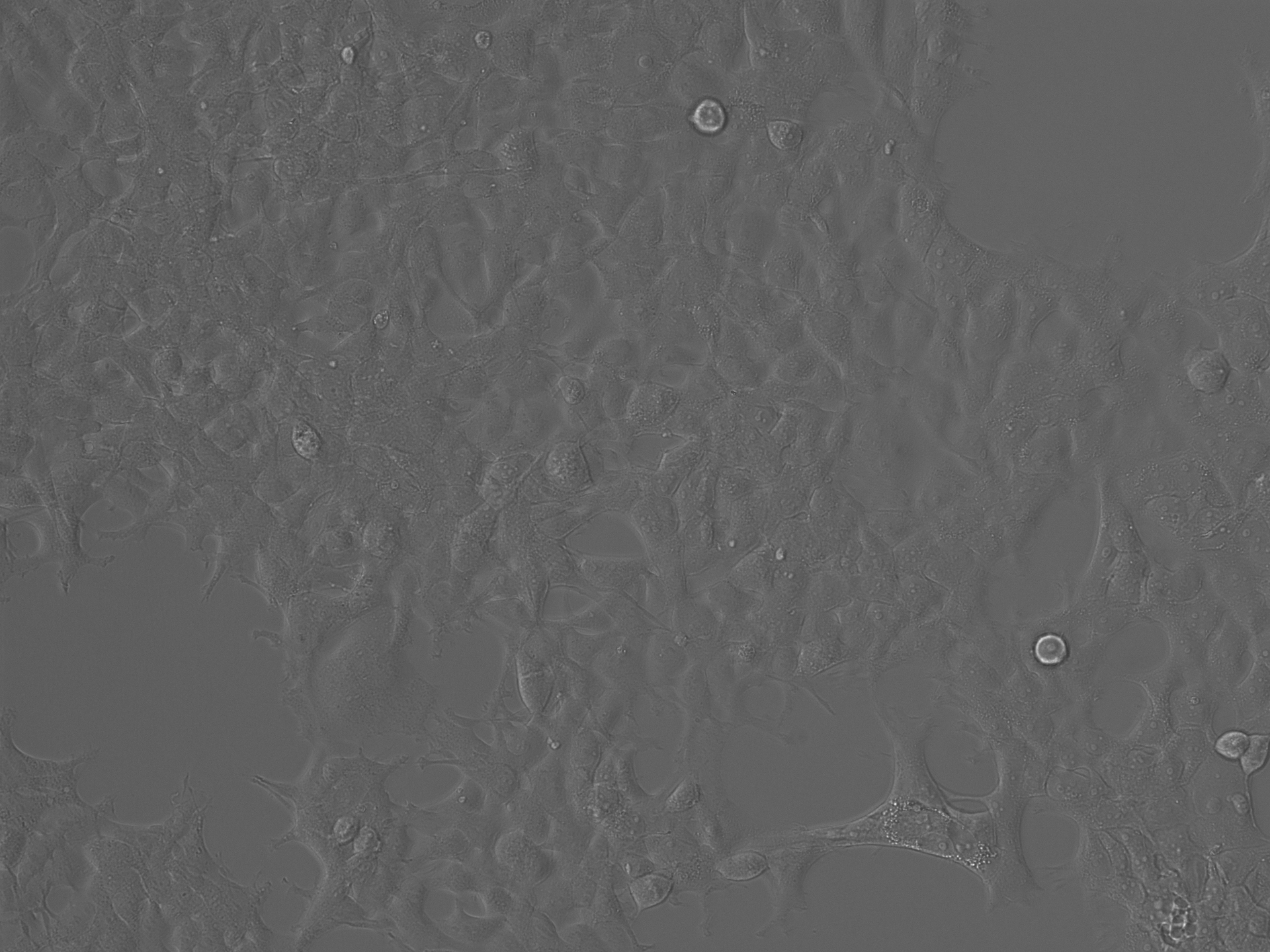

### Free AAV2_Brightfield_Day7.tif

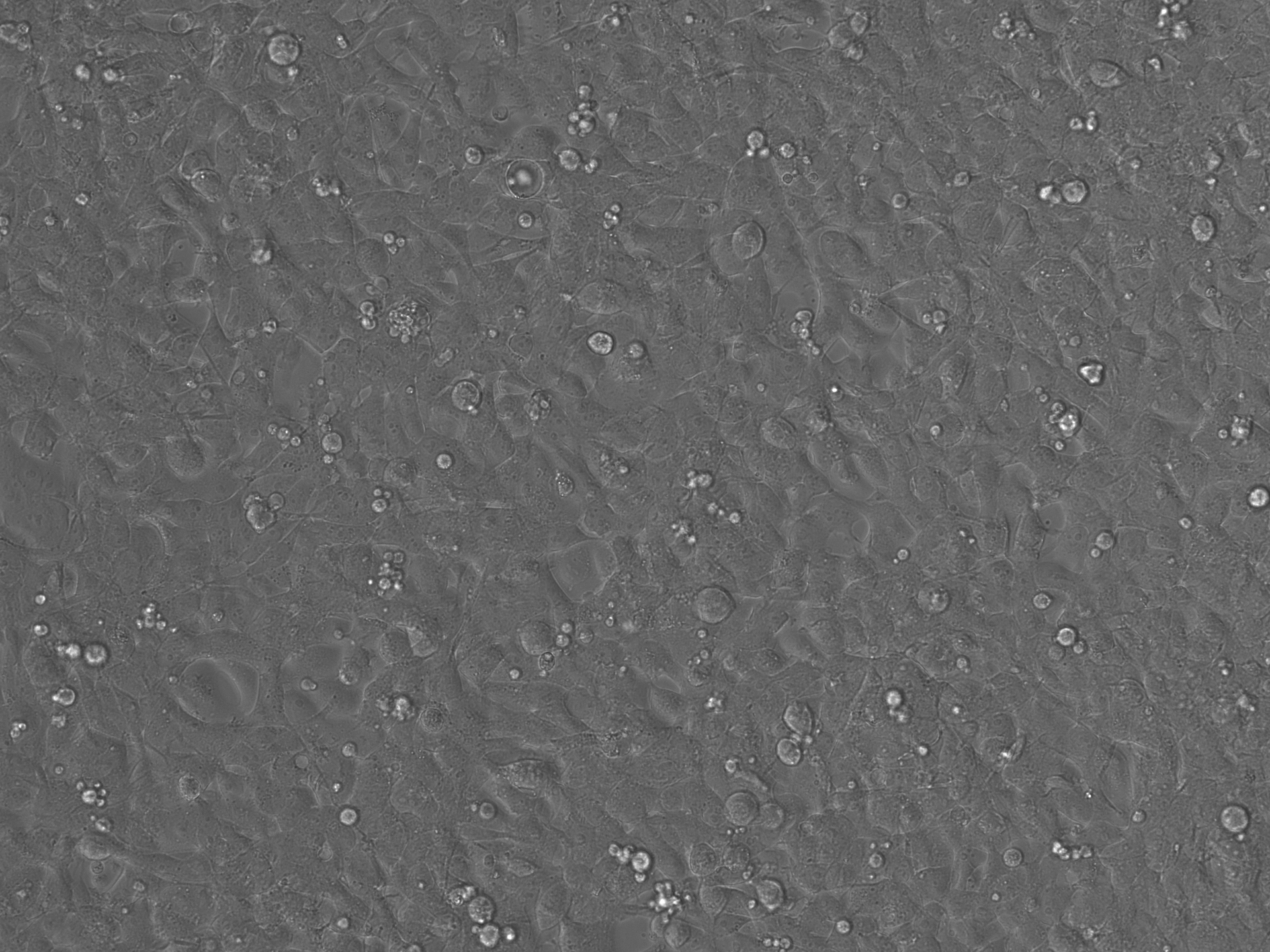

### Free AAV2_Brightfield_Day7.tif

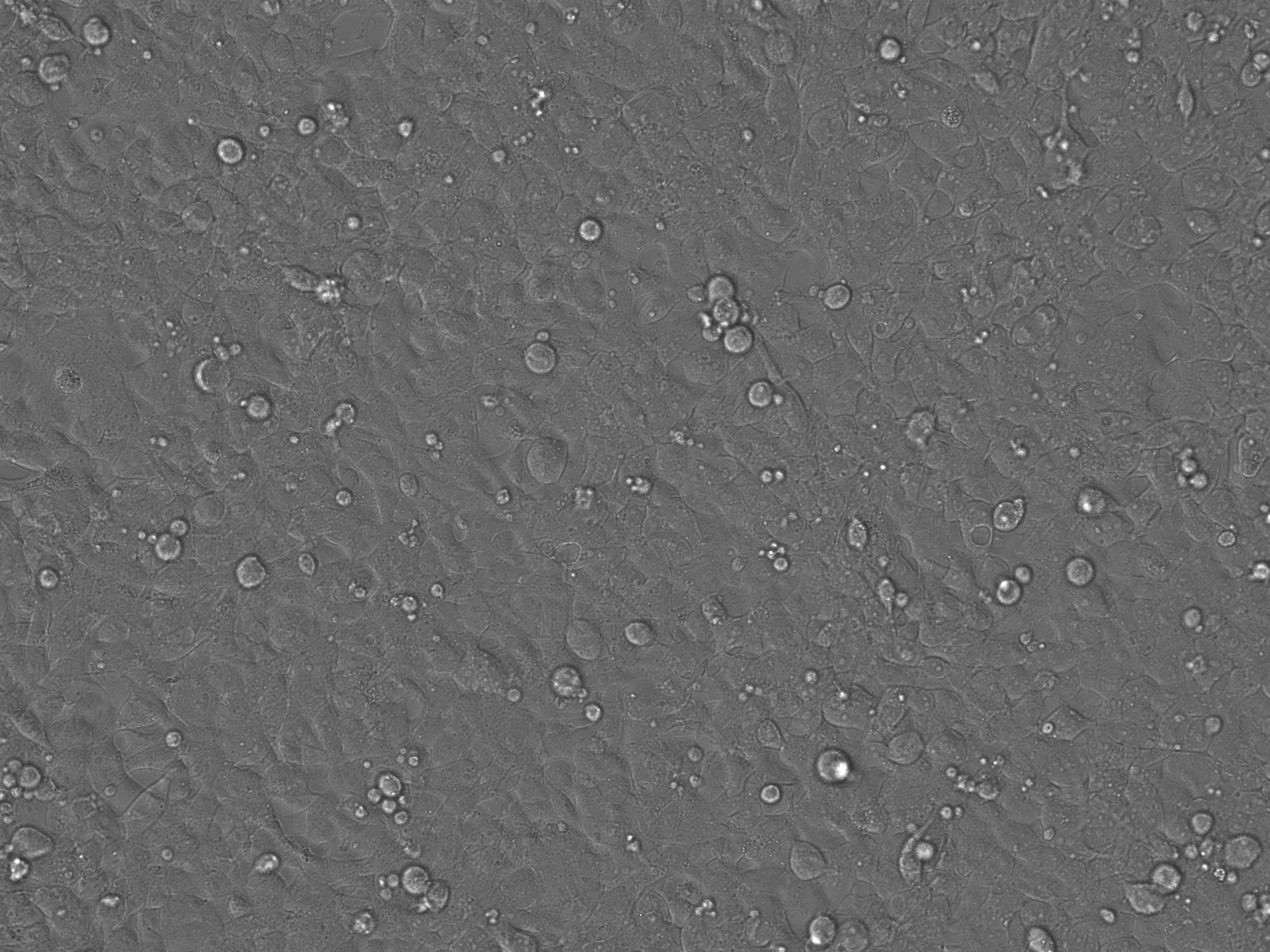

### Free AAV2_LumiScarlet_Day3.tif

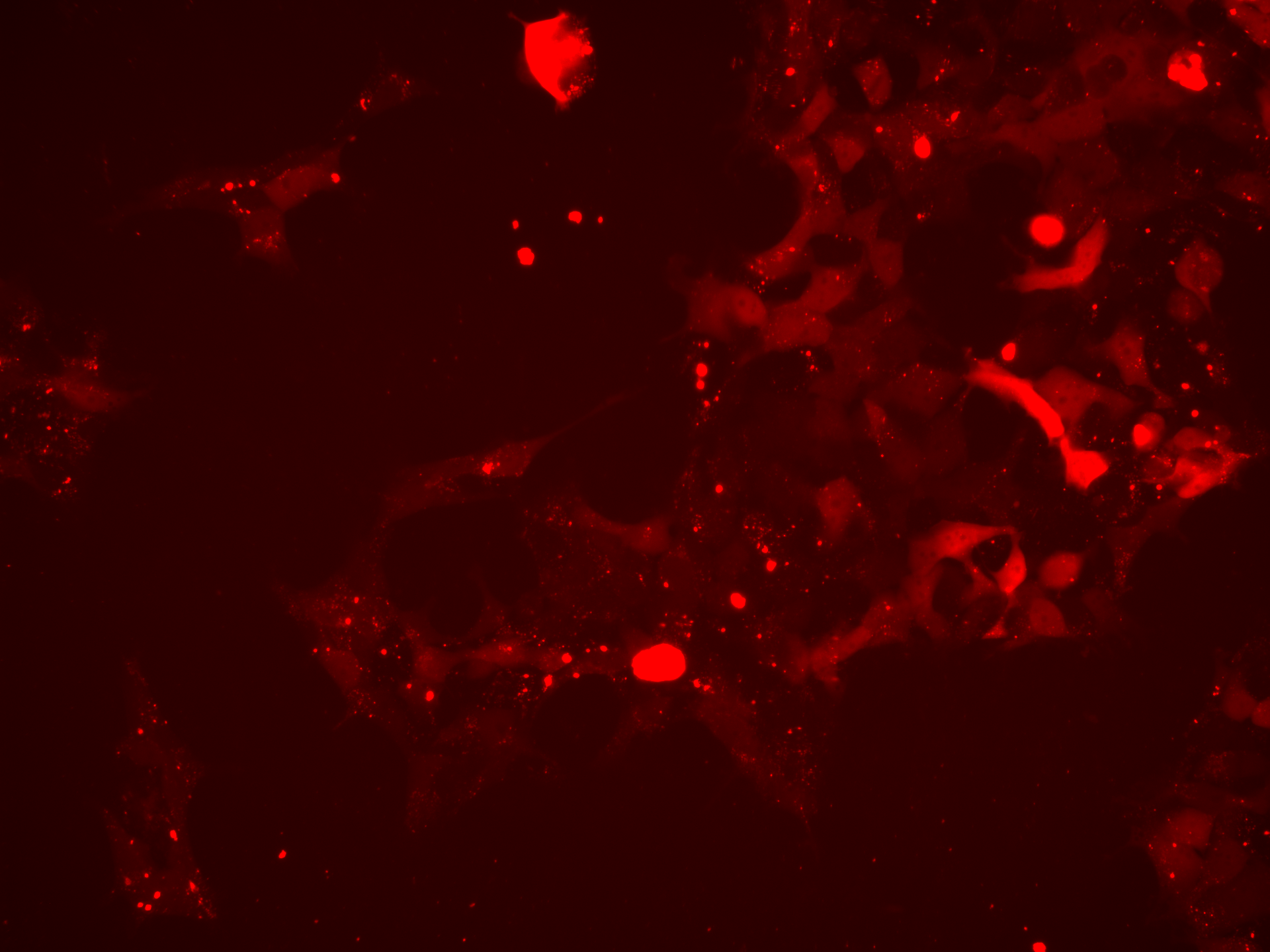

### Free AAV2_LumiScarlet_Day3.tif

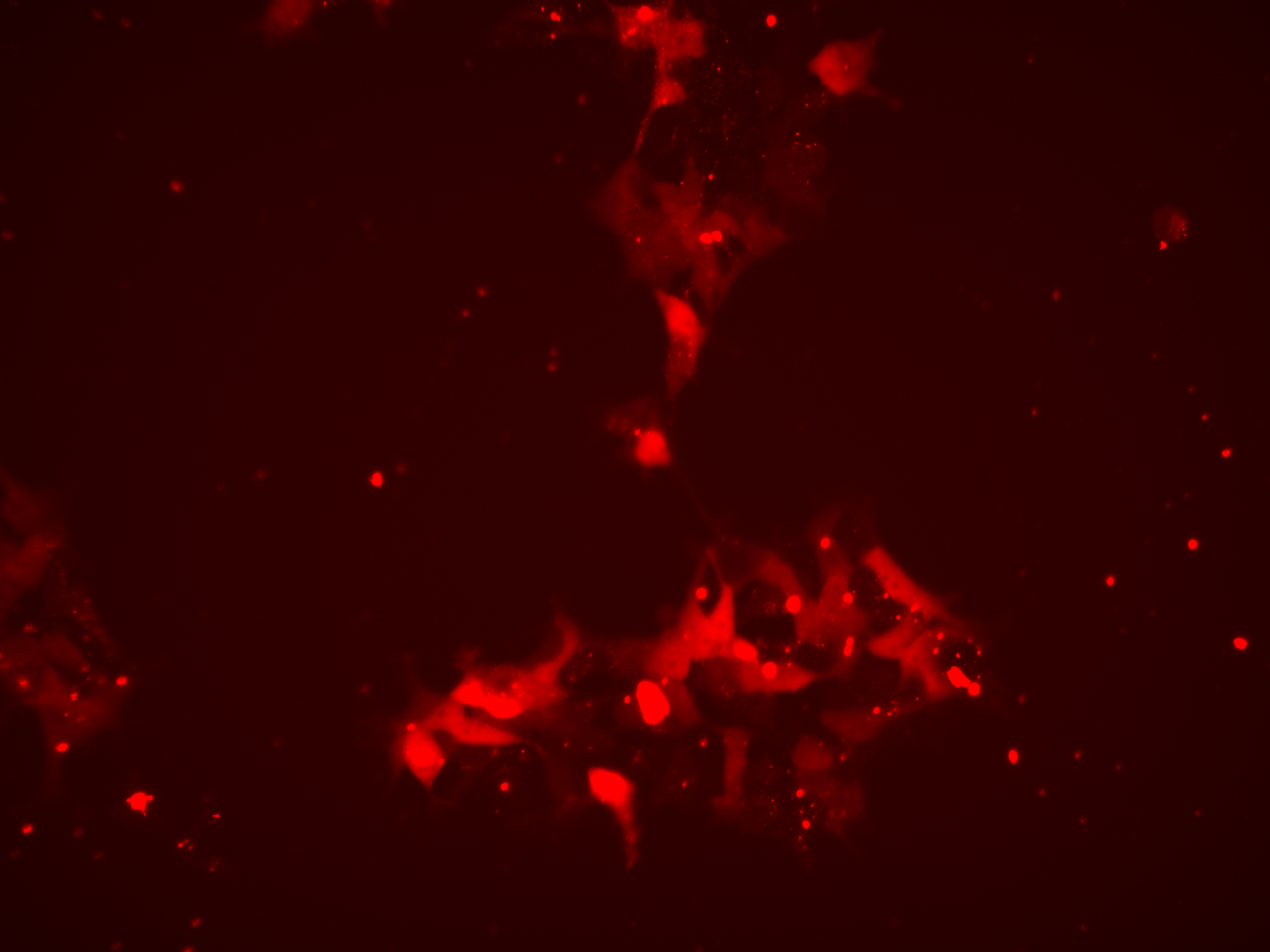

### Free AAV2_LumiScarlet_Day5.tif

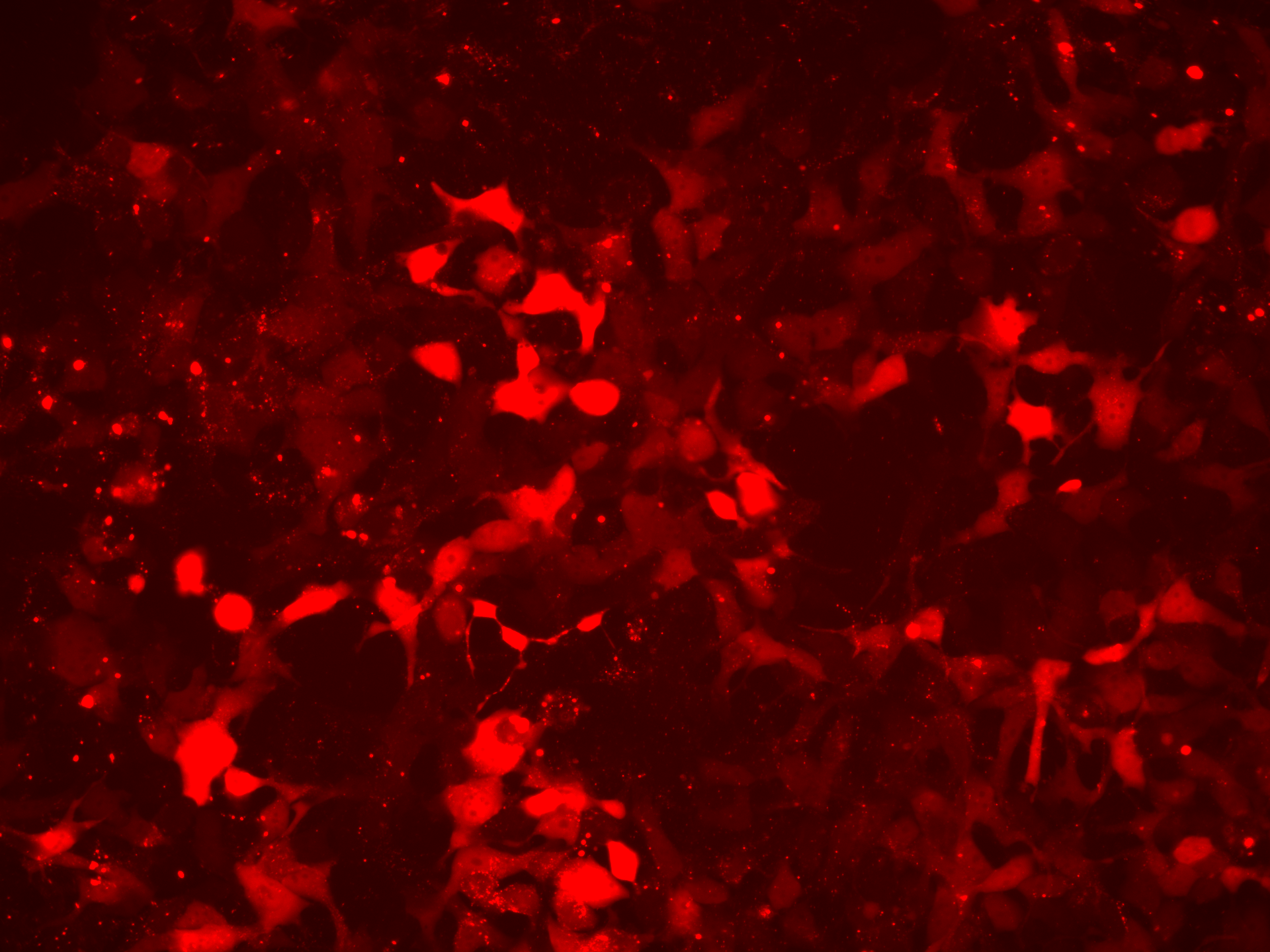

### Free AAV2_LumiScarlet_Day5.tif

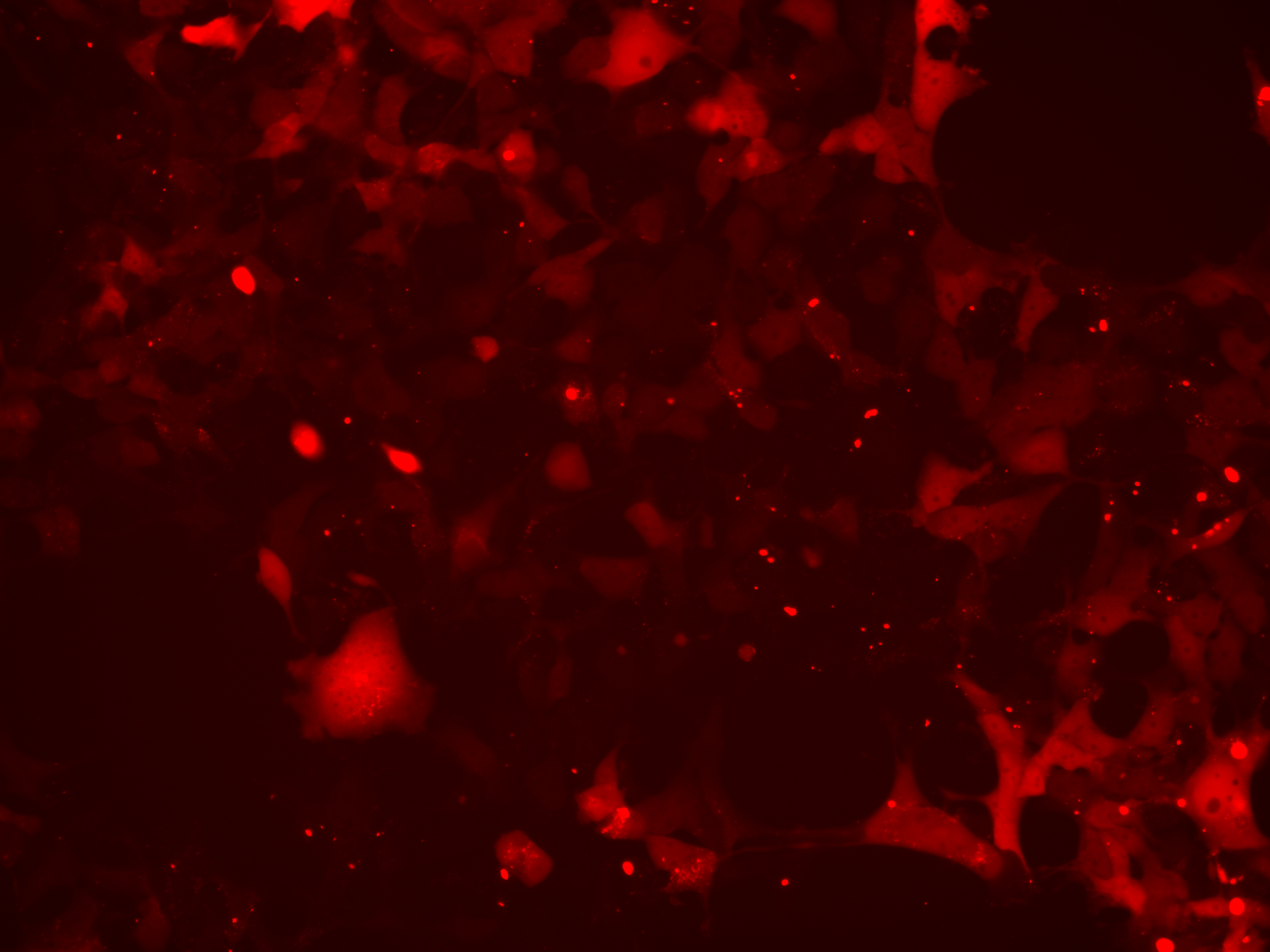

### Free AAV2_LumiScarlet_Day7.tif

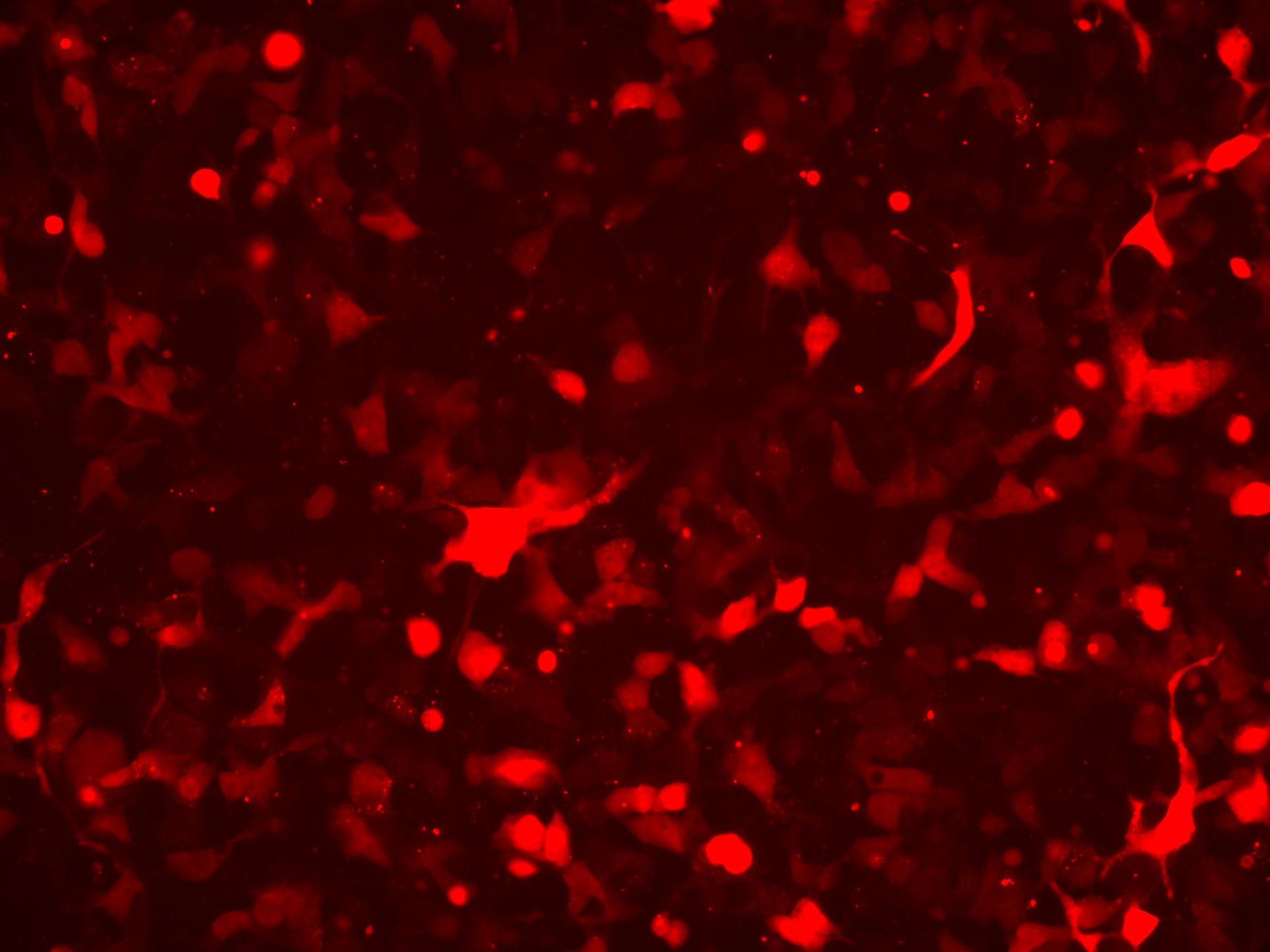

### Free AAV2_LumiScarlet_Day7.tif

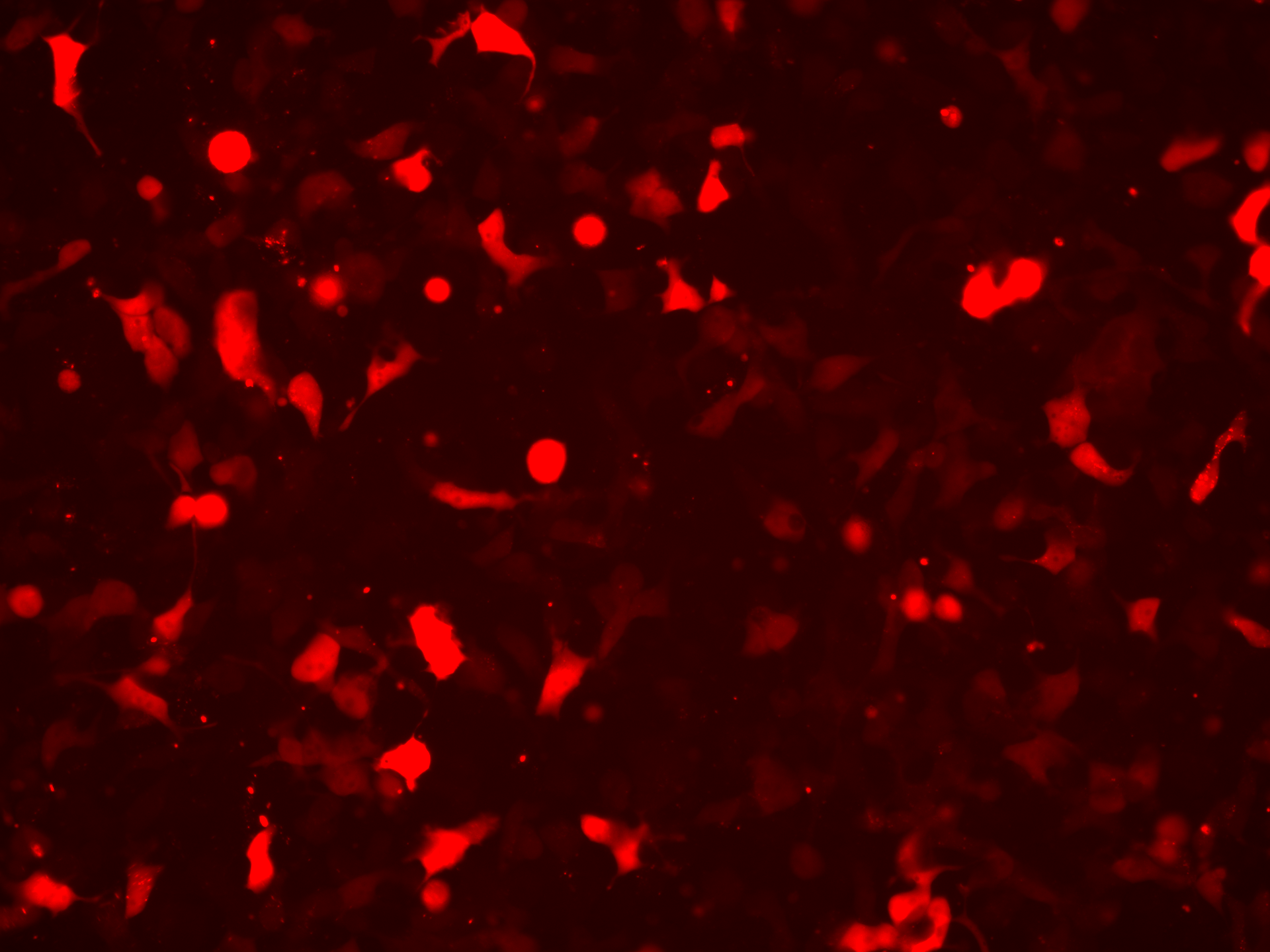

### Free AAV2_Overlay_Day3.tif

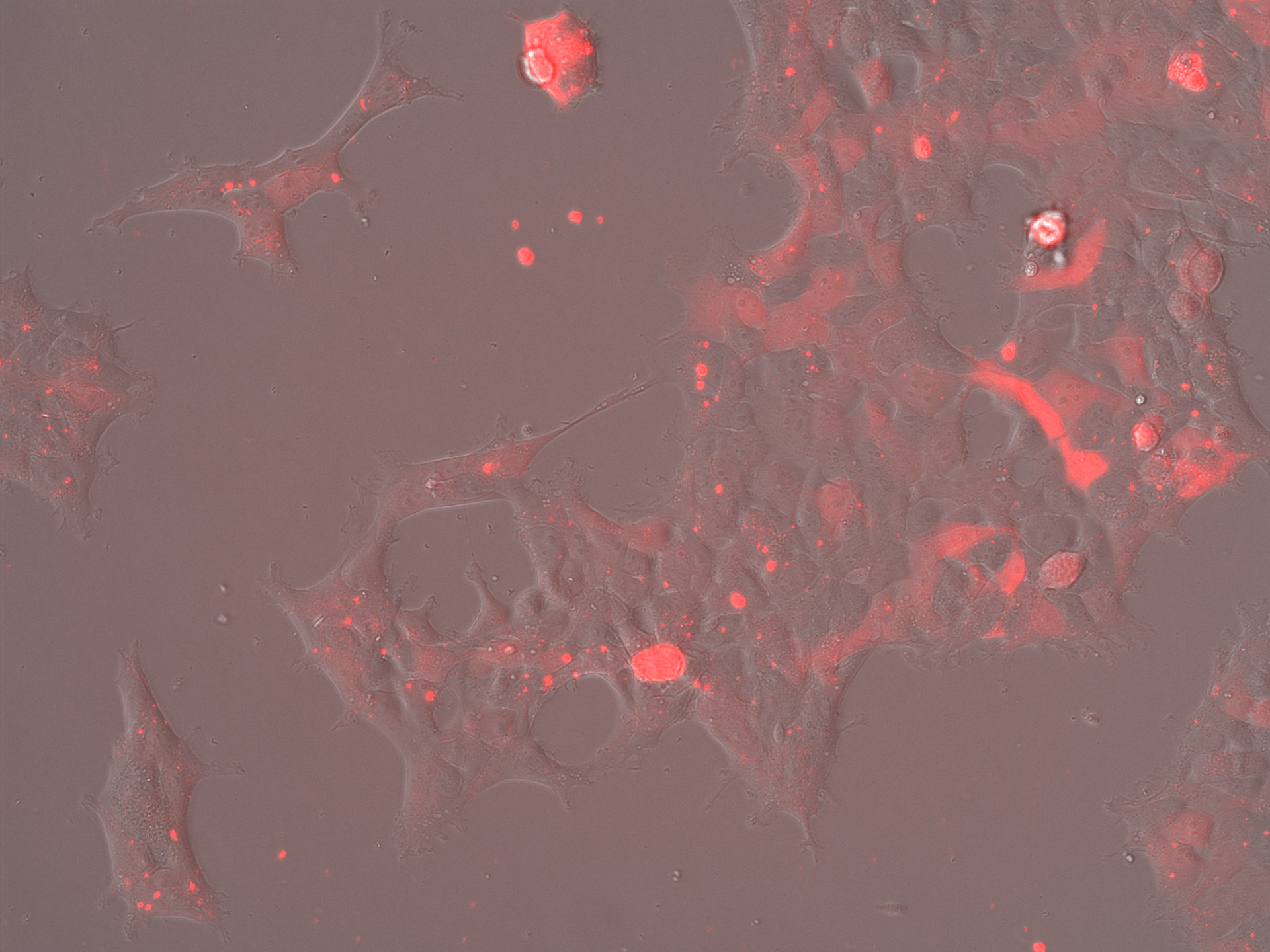

### Free AAV2_Overlay_Day3.tif

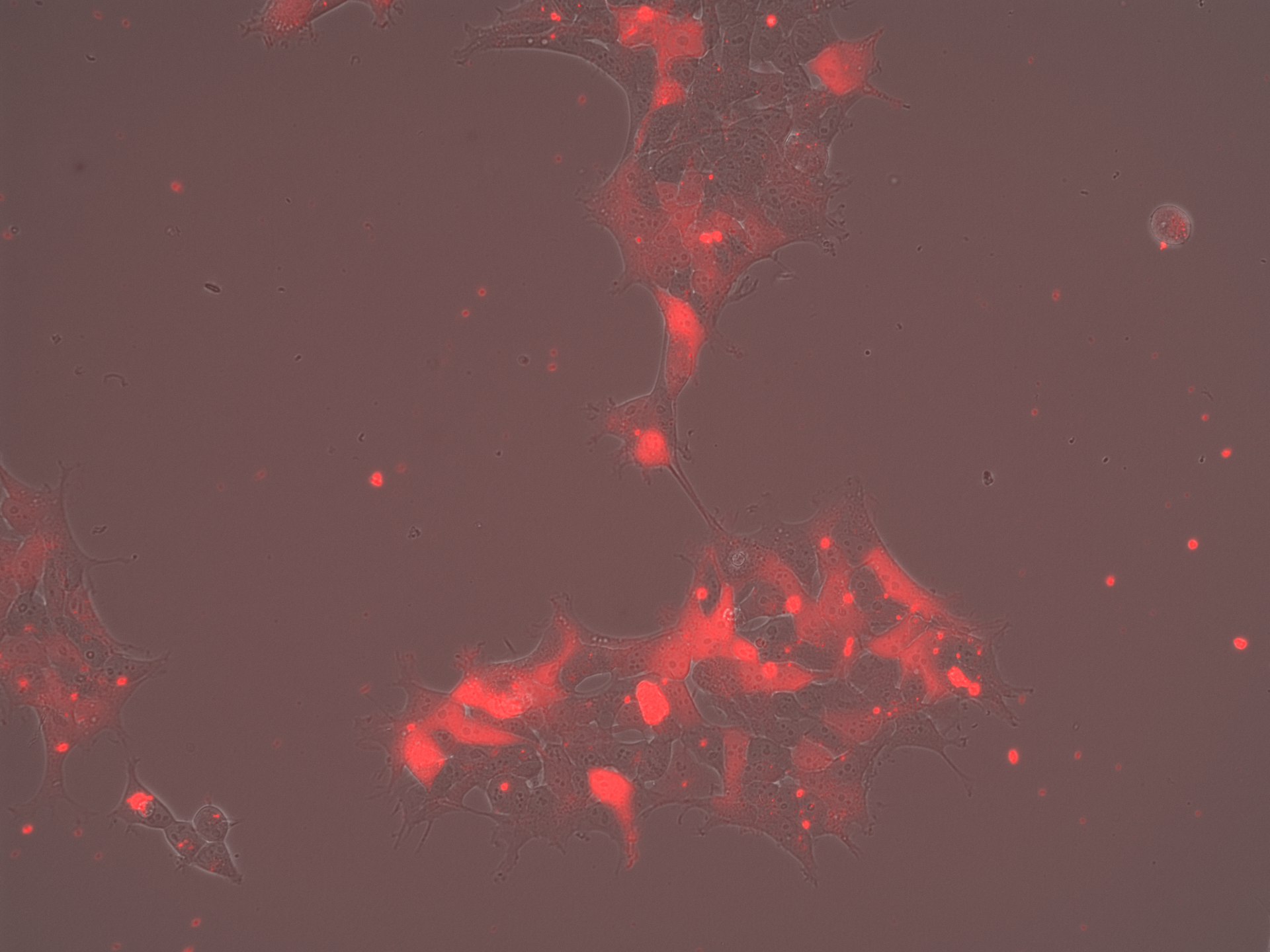

### Free AAV2_Overlay_Day5.tif

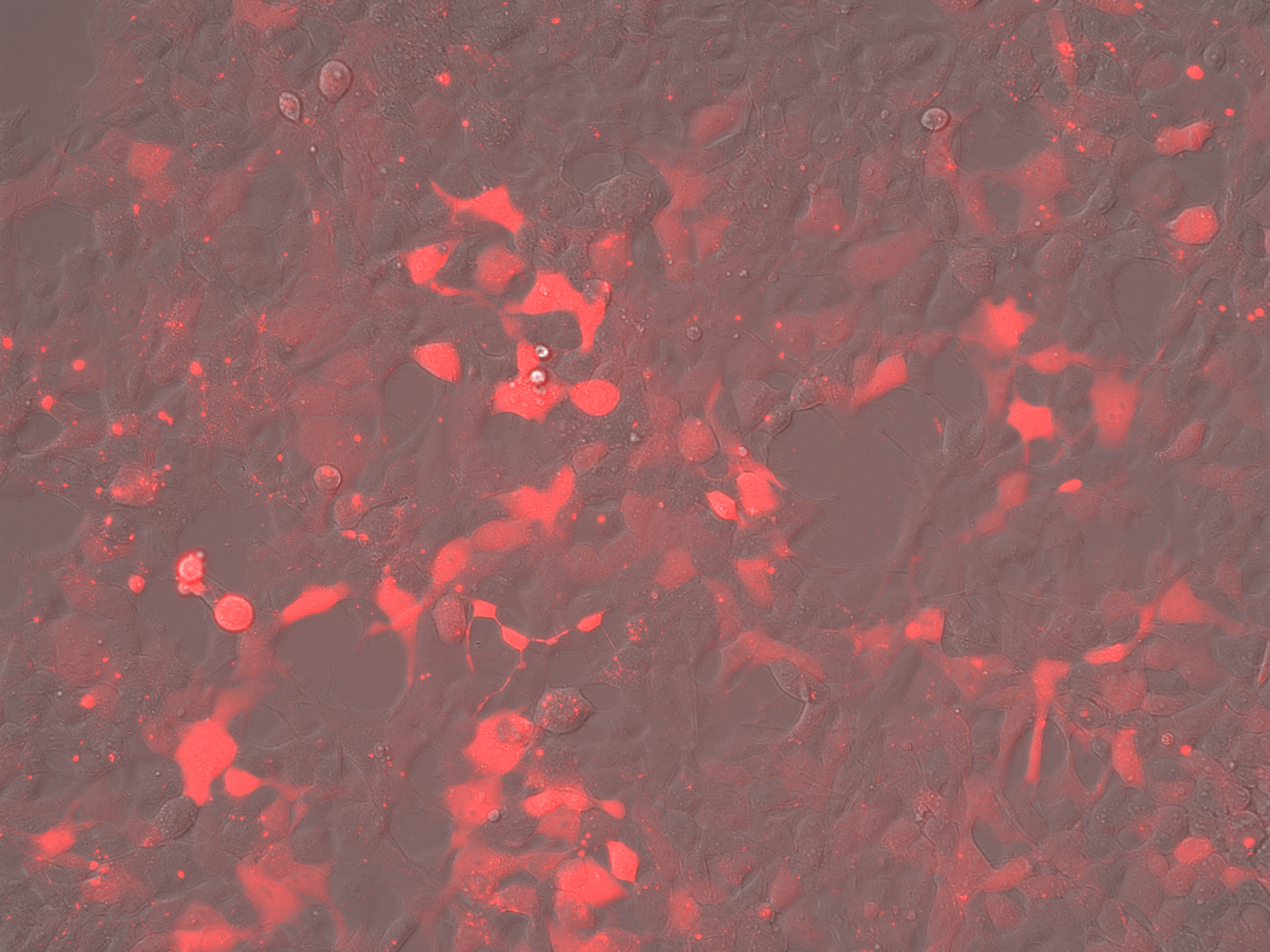

### Free AAV2_Overlay_Day5.tif

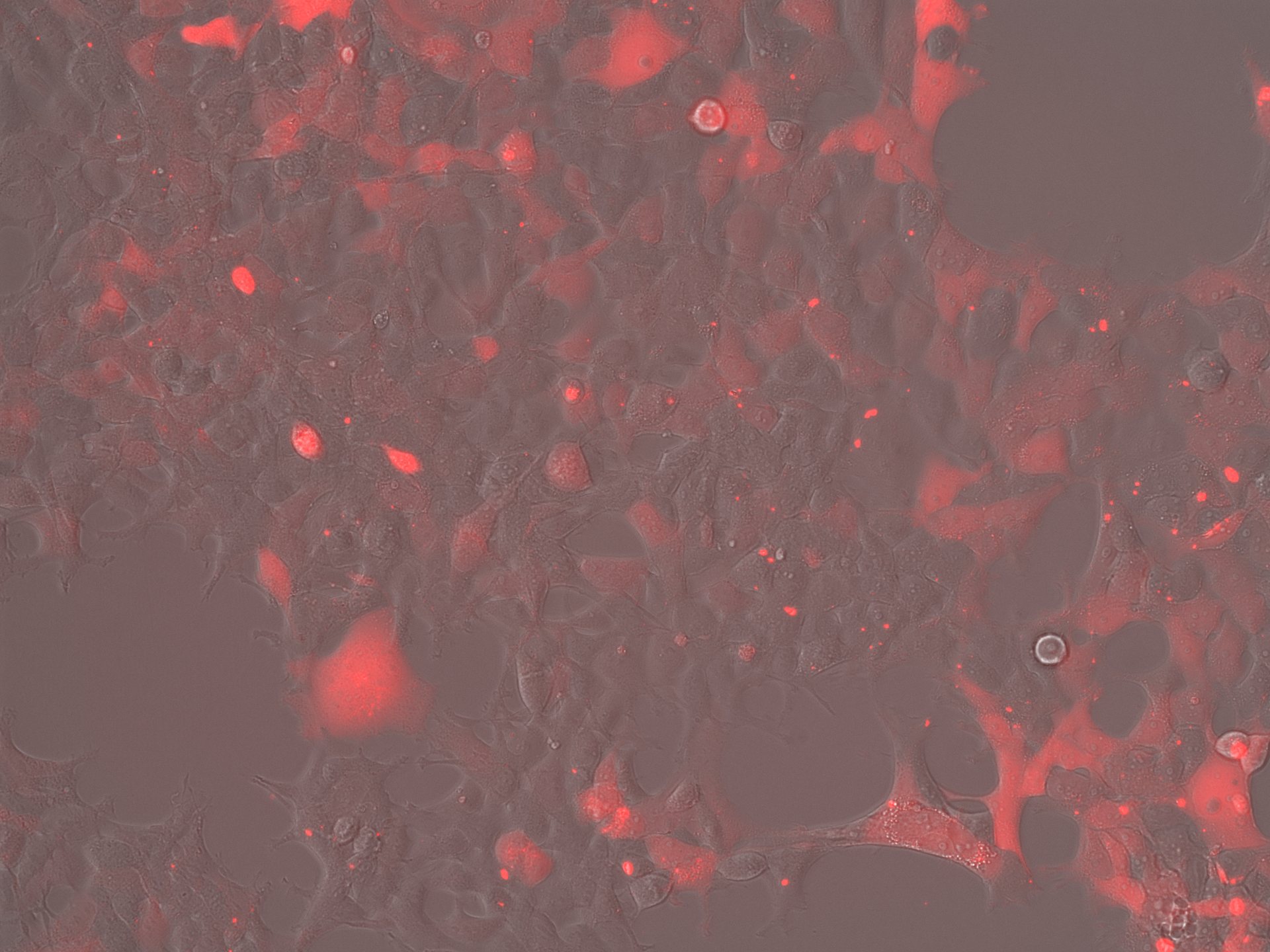

### Free AAV2_Overlay_Day7.tif

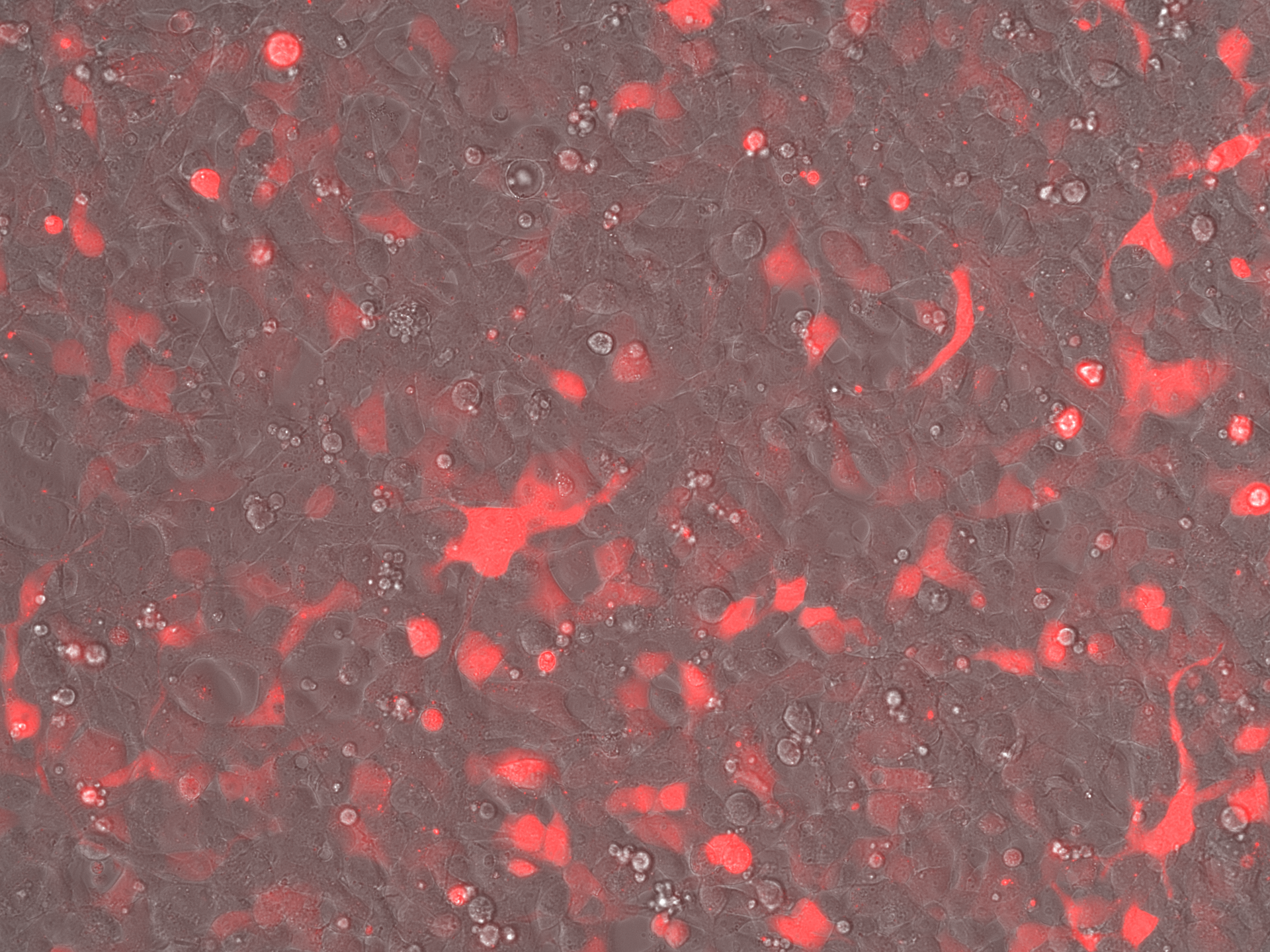

### Free AAV2_Overlay_Day7.tif

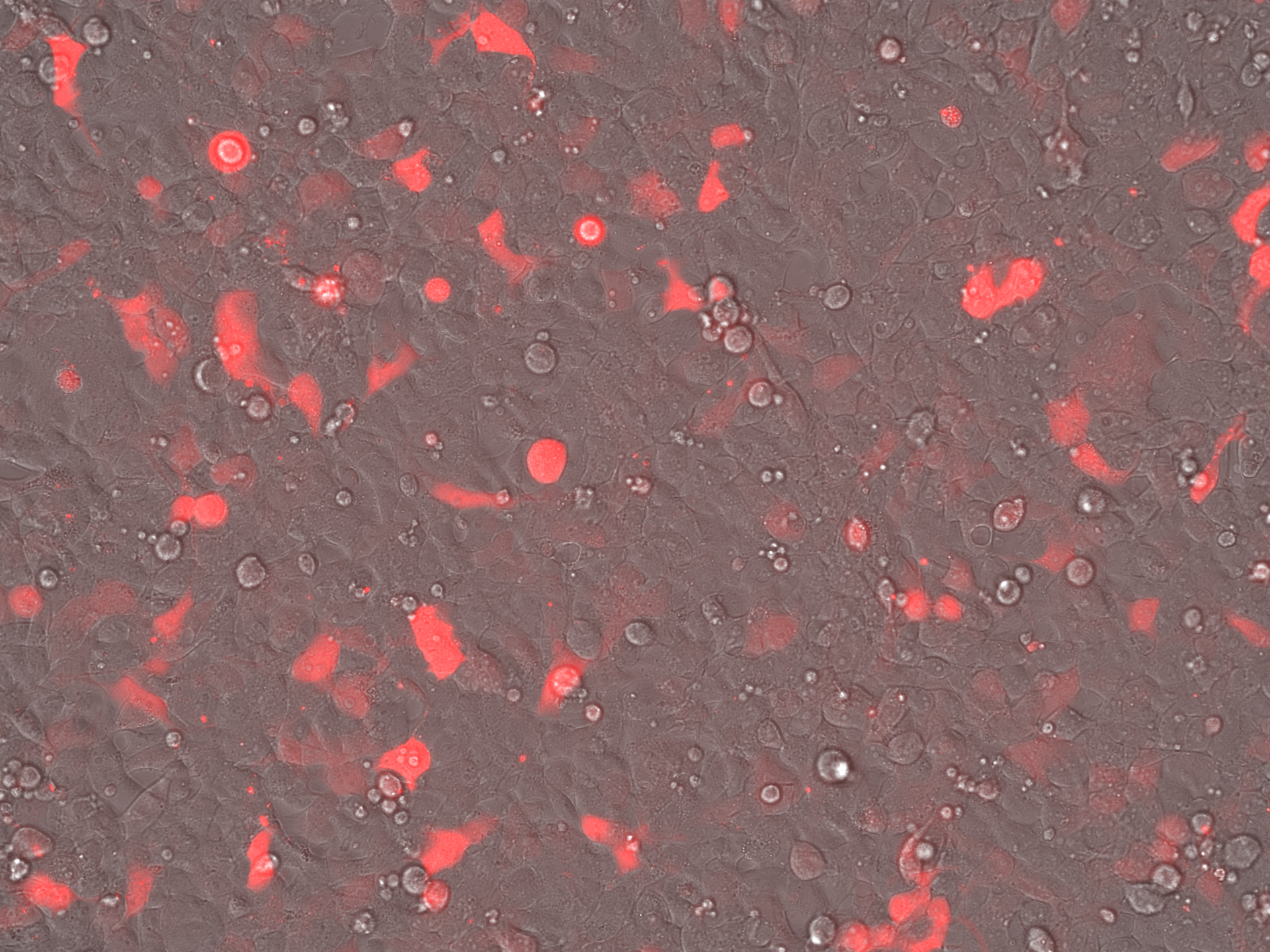

### No Rep-Cap Iodixanol_Brightfield_Day3.tif

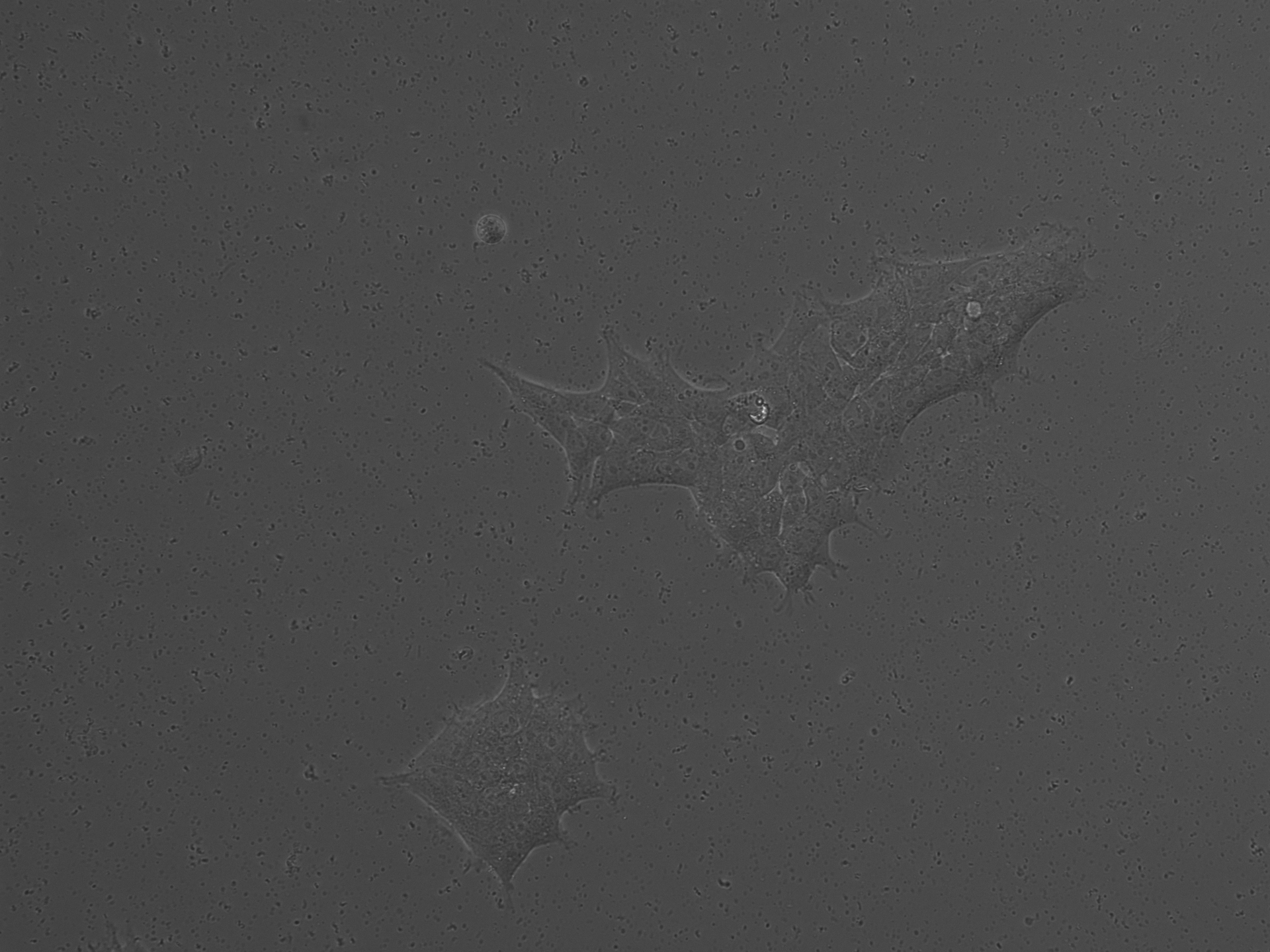

### No Rep-Cap Iodixanol_Brightfield_Day3.tif

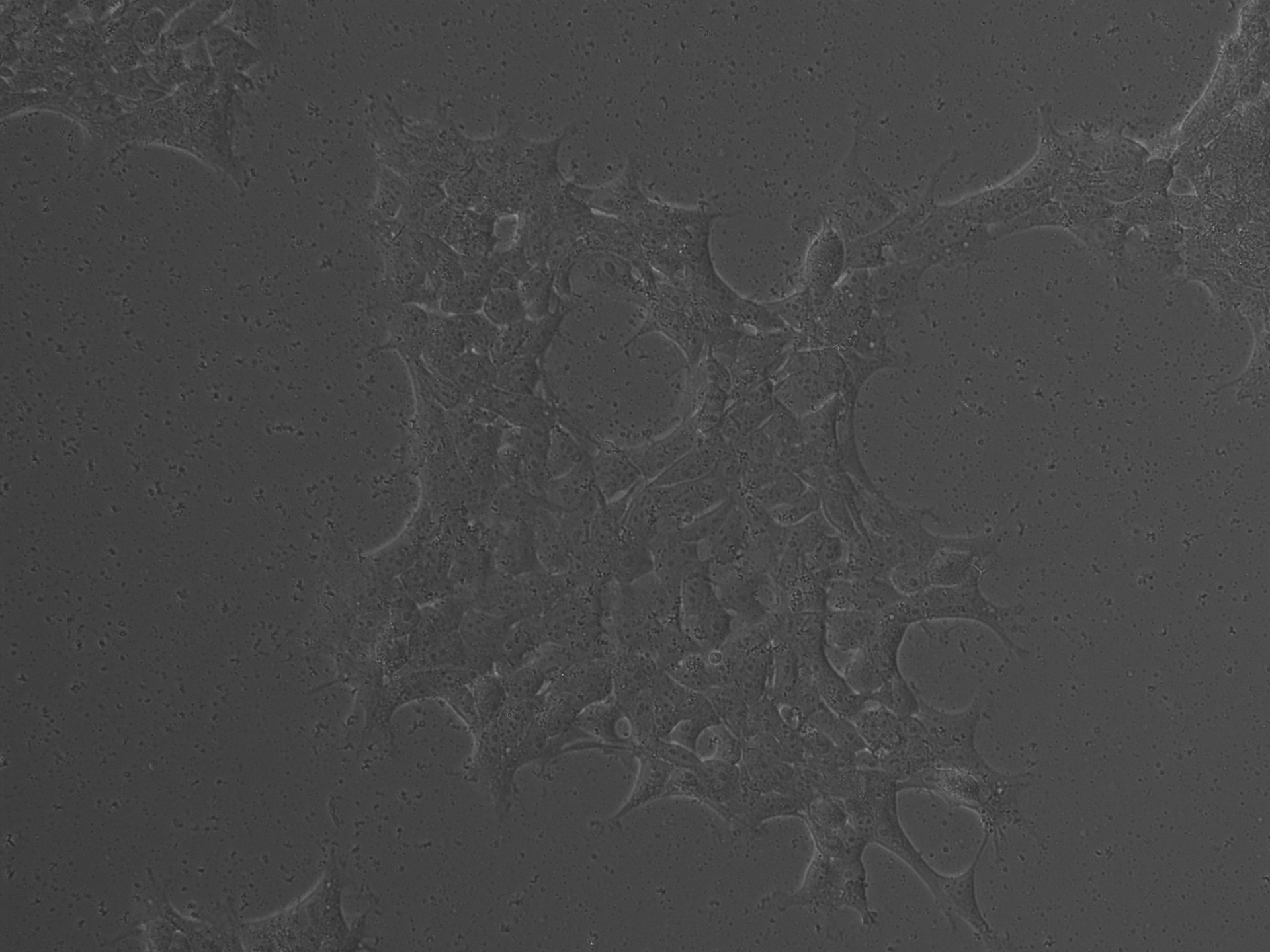

### No Rep-Cap Iodixanol_Brightfield_Day5.tif

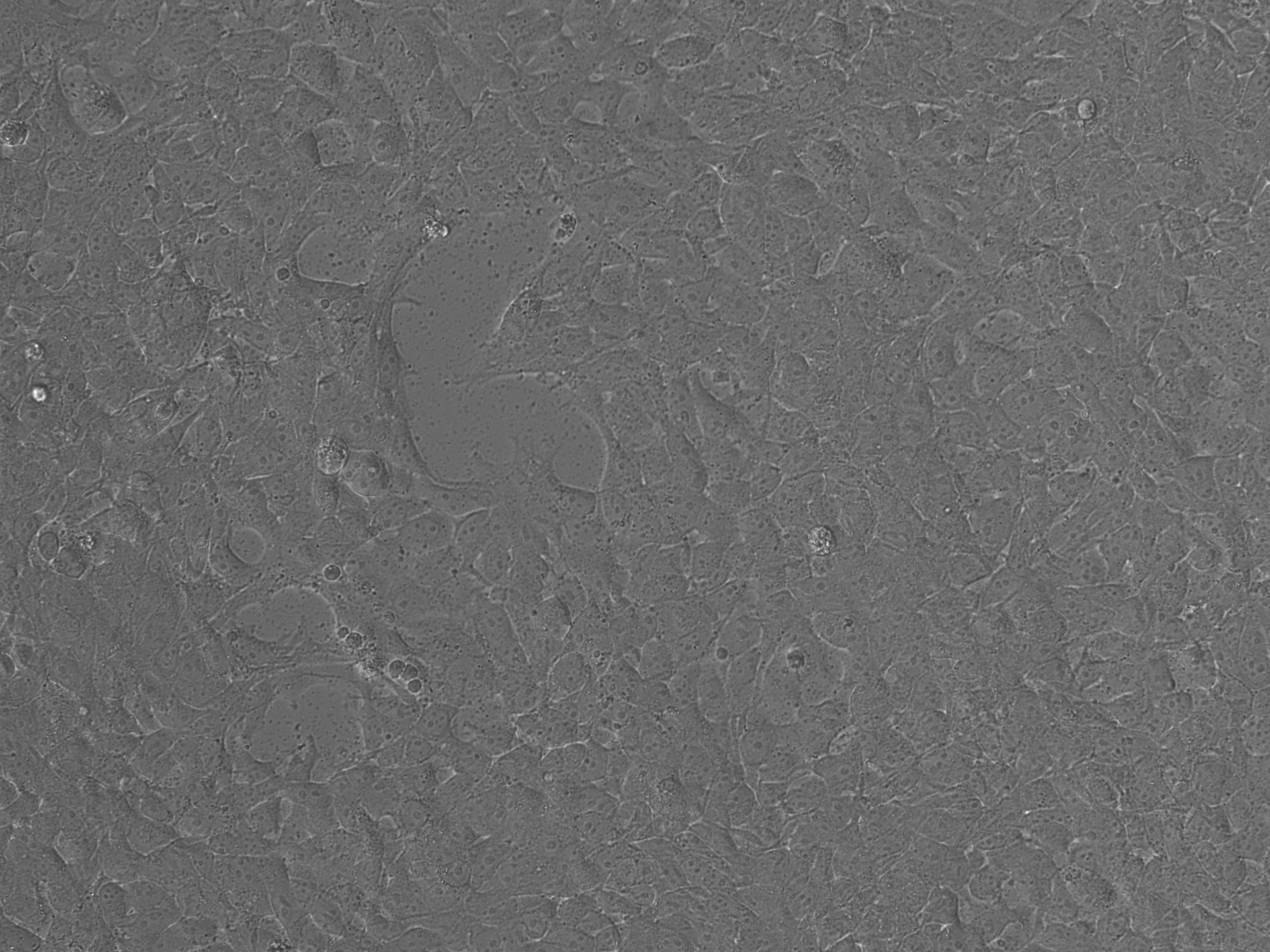

### No Rep-Cap Iodixanol_Brightfield_Day7.tif

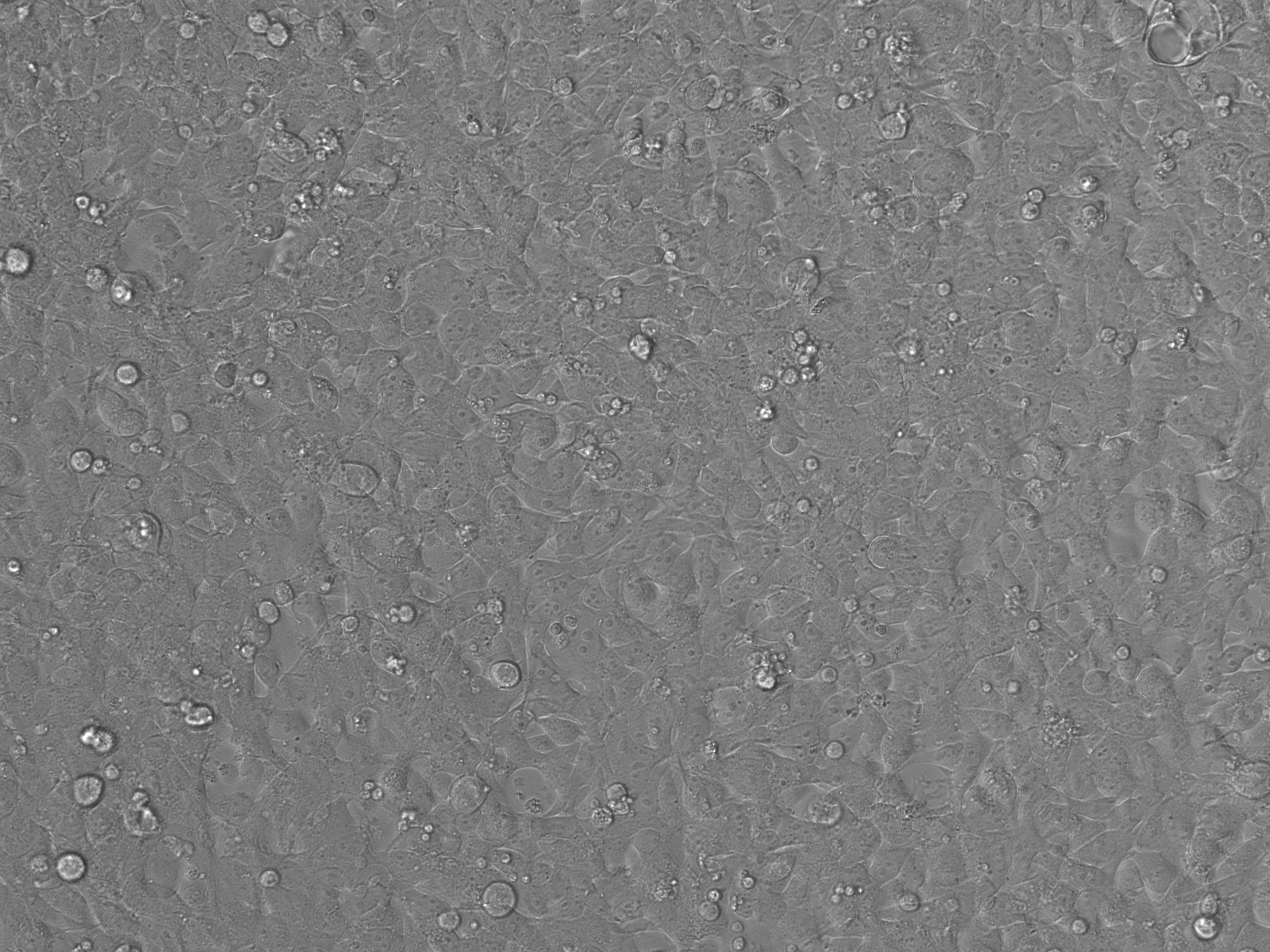

### No Rep-Cap Iodixanol_Brightfield_Day7.tif

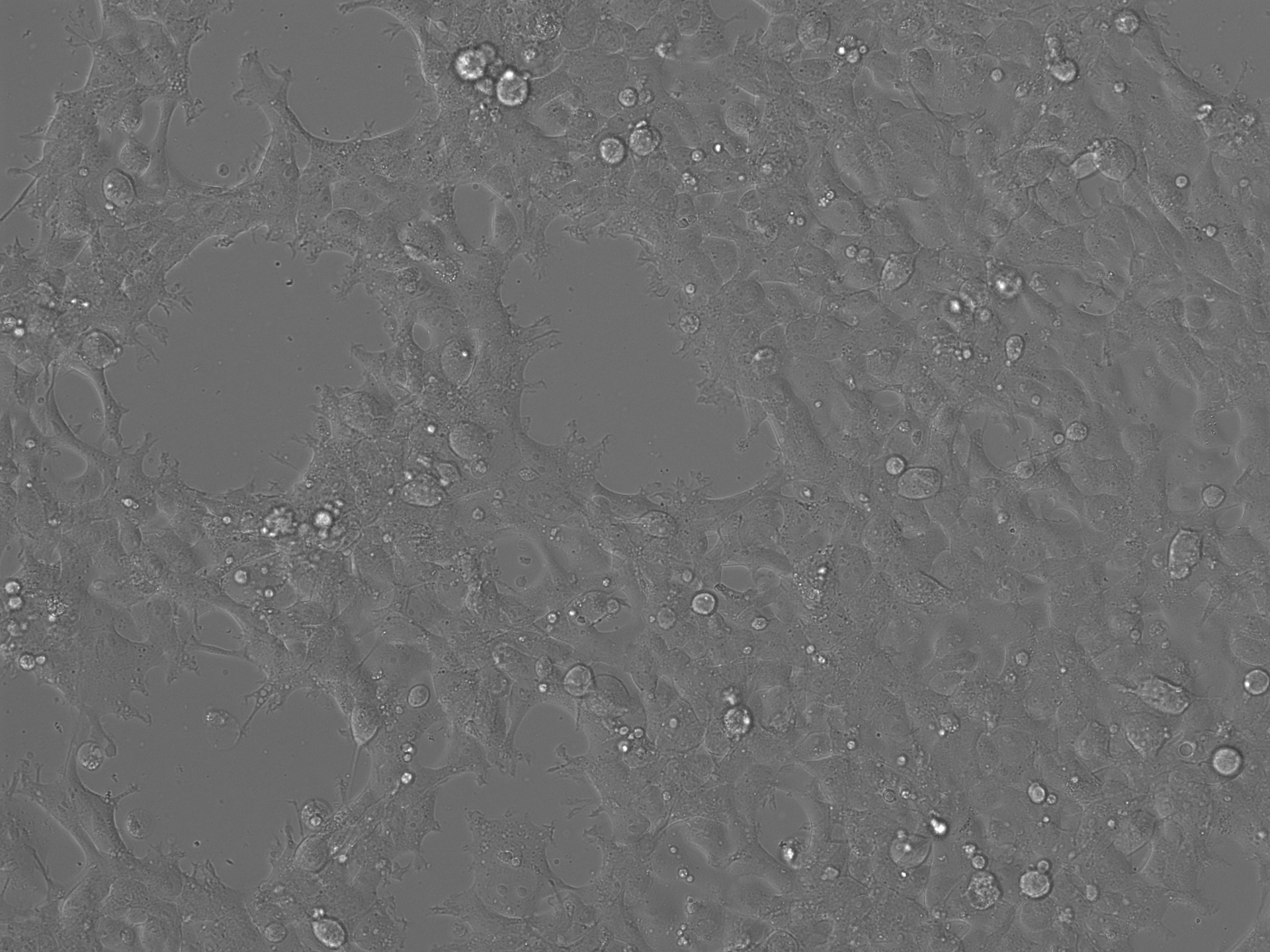

### No Rep-Cap Iodixanol_LumiScarlet_Day3.tif

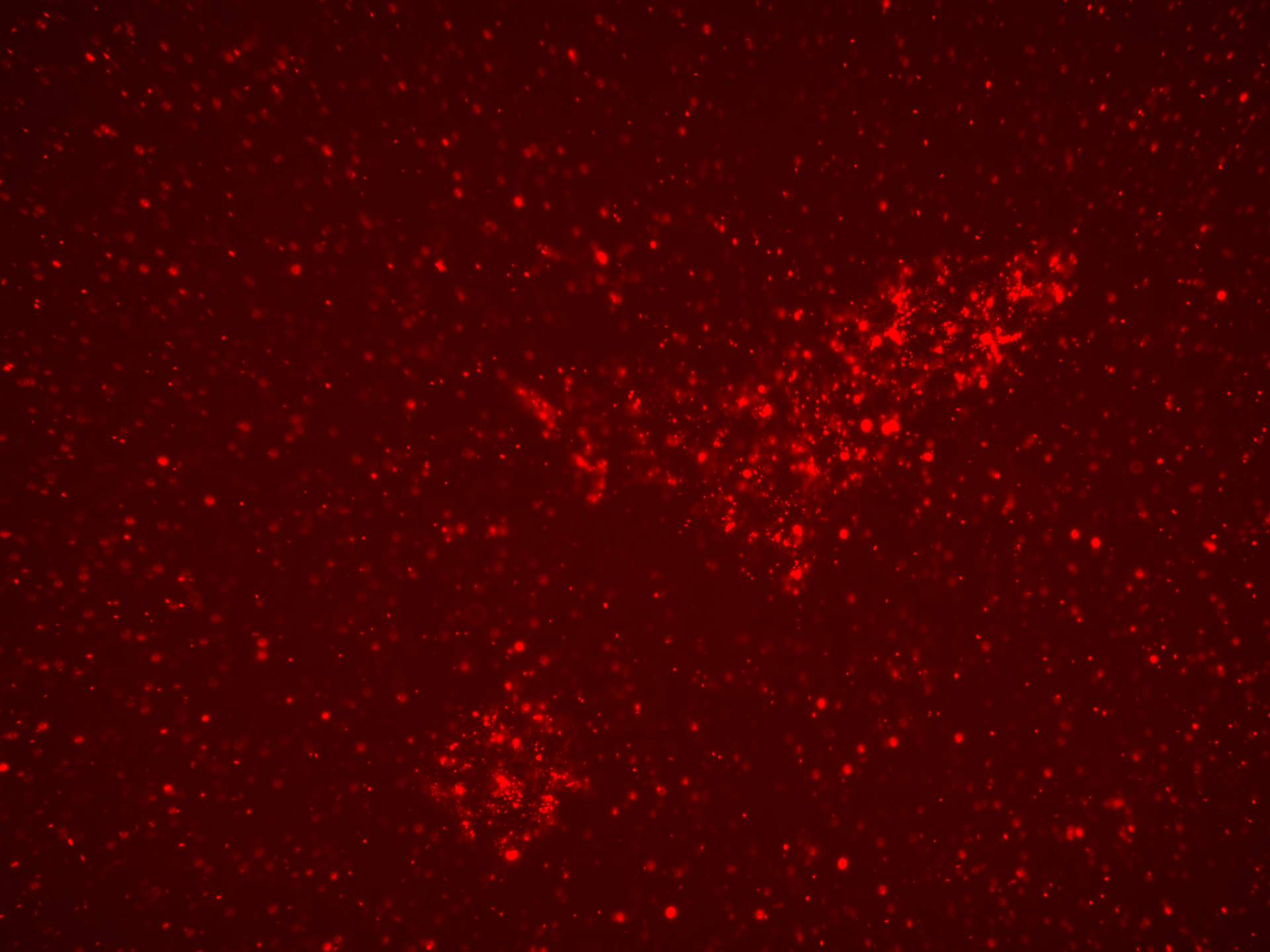

### No Rep-Cap Iodixanol_LumiScarlet_Day3.tif

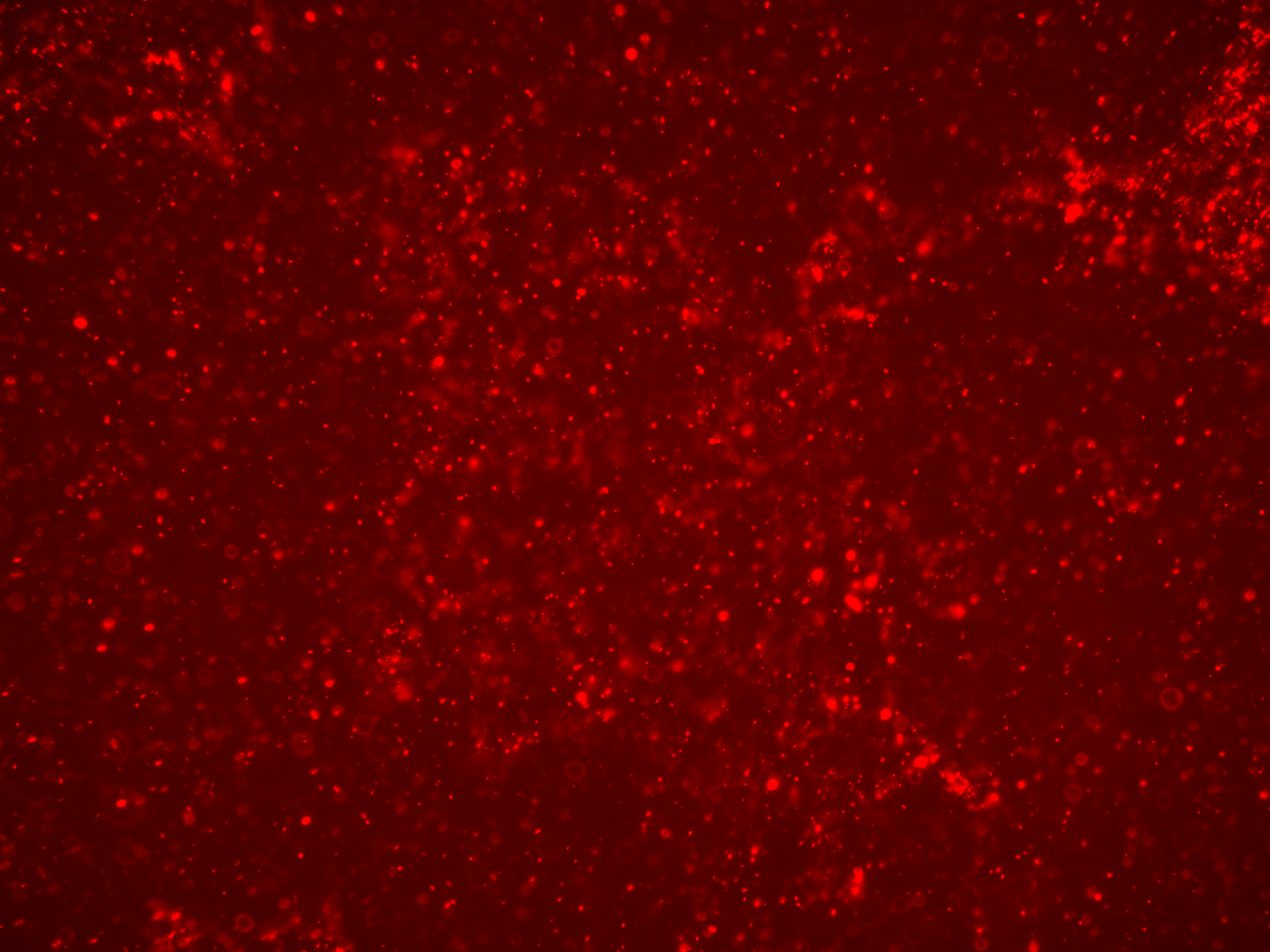

### No Rep-Cap Iodixanol_LumiScarlet_Day5.tif

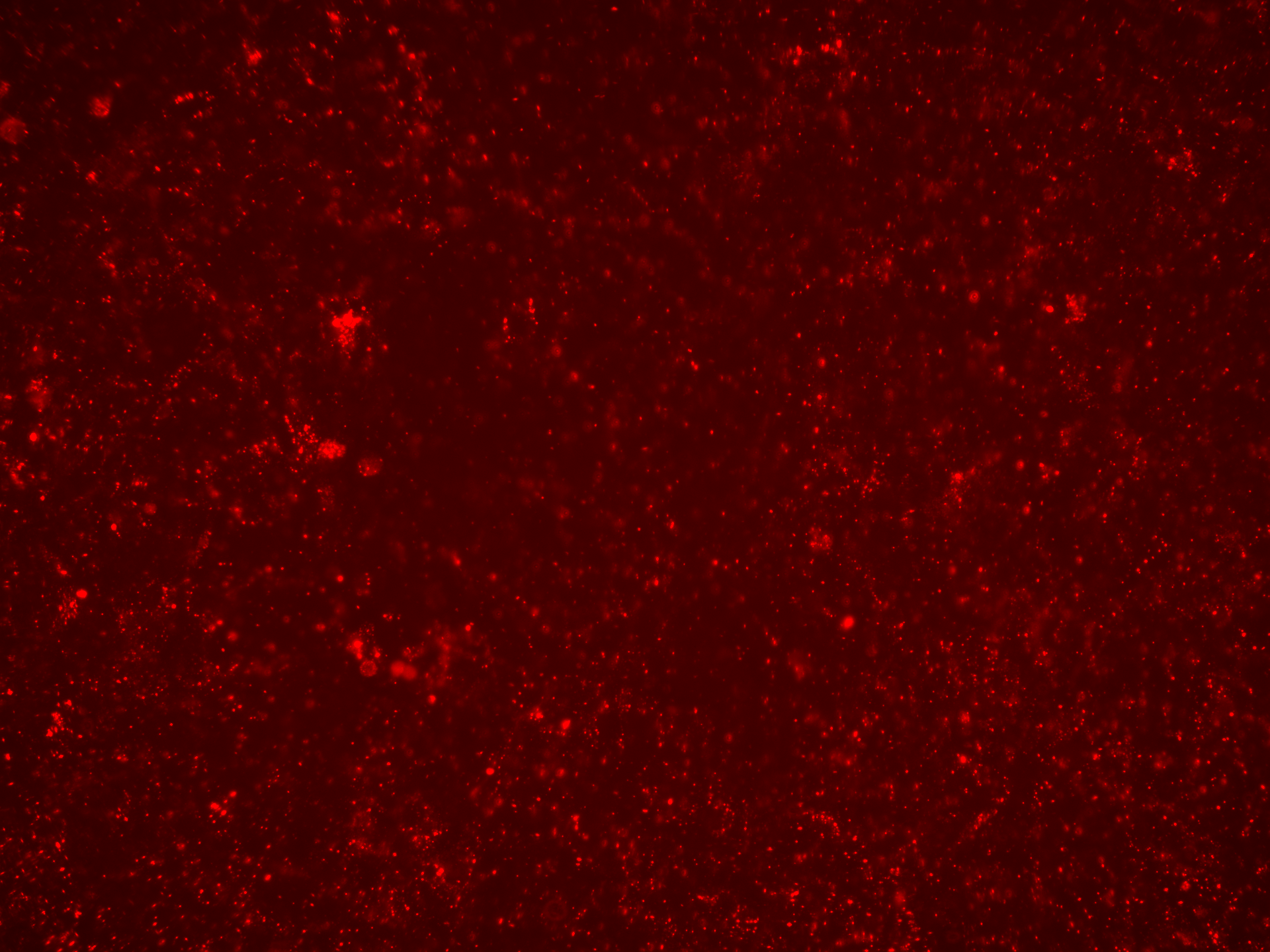

### No Rep-Cap Iodixanol_LumiScarlet_Day7.tif

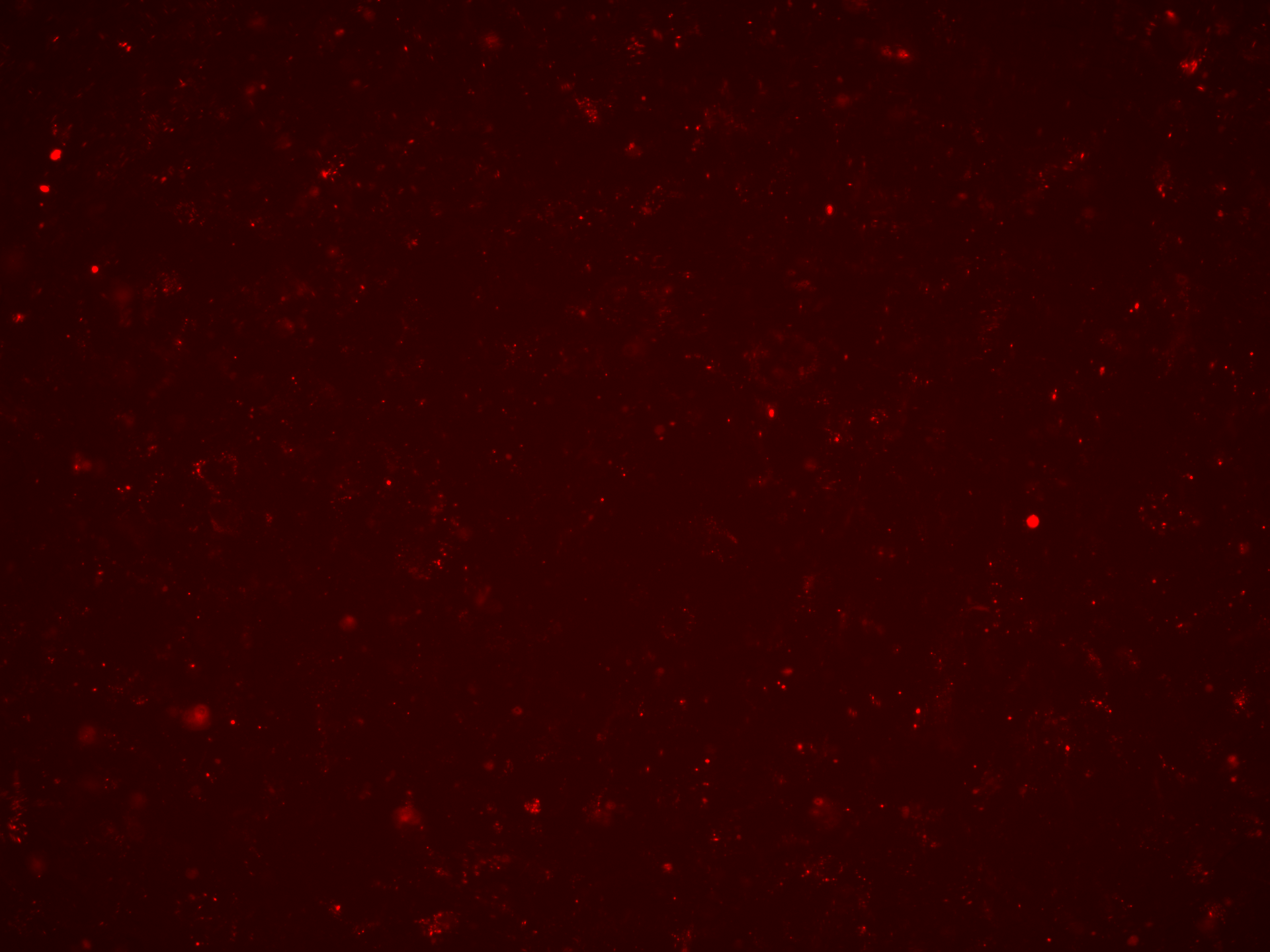

### No Rep-Cap Iodixanol_LumiScarlet_Day7.tif

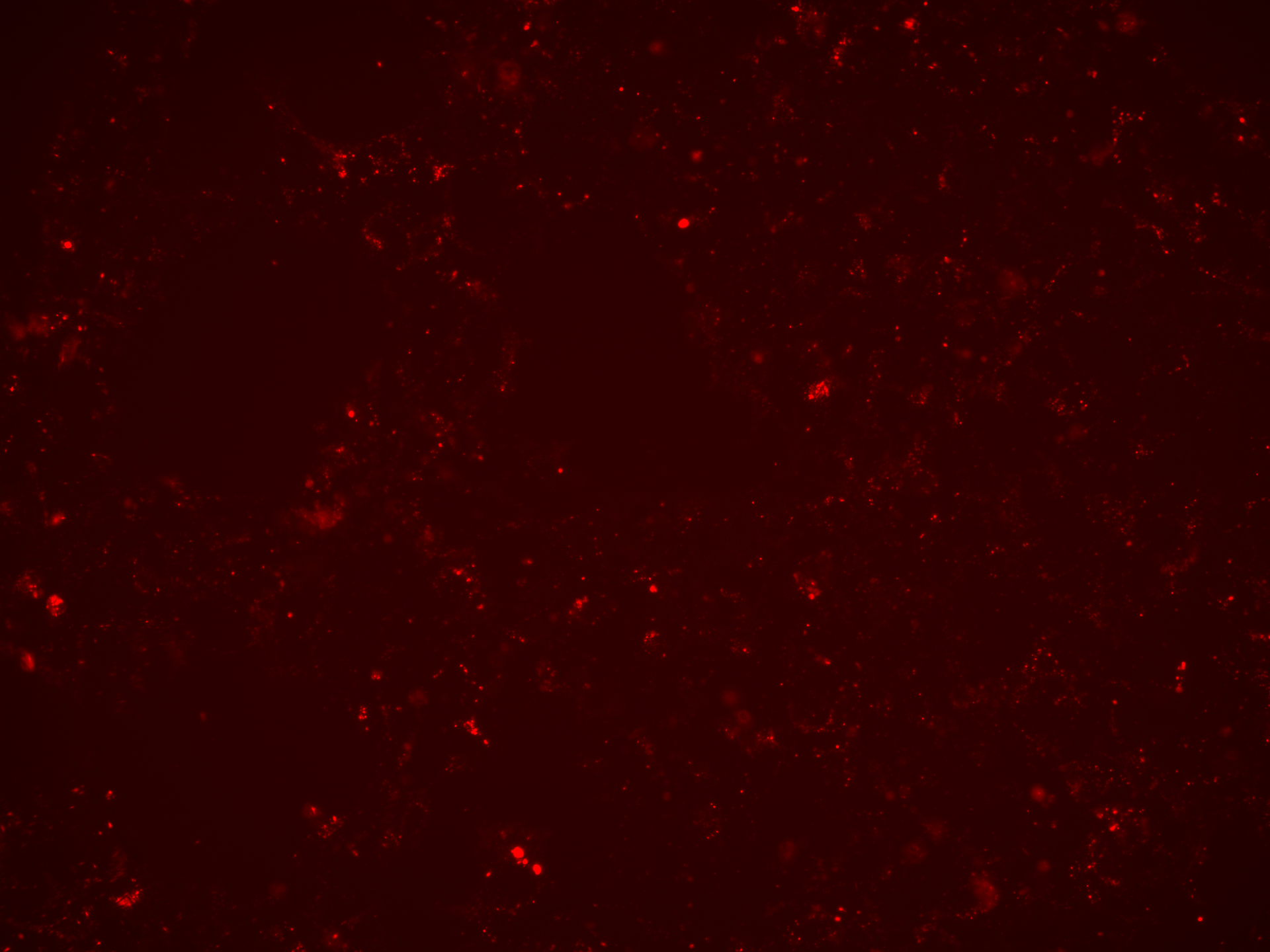

### No Rep-Cap Iodixanol_Overlay_Day3.tif

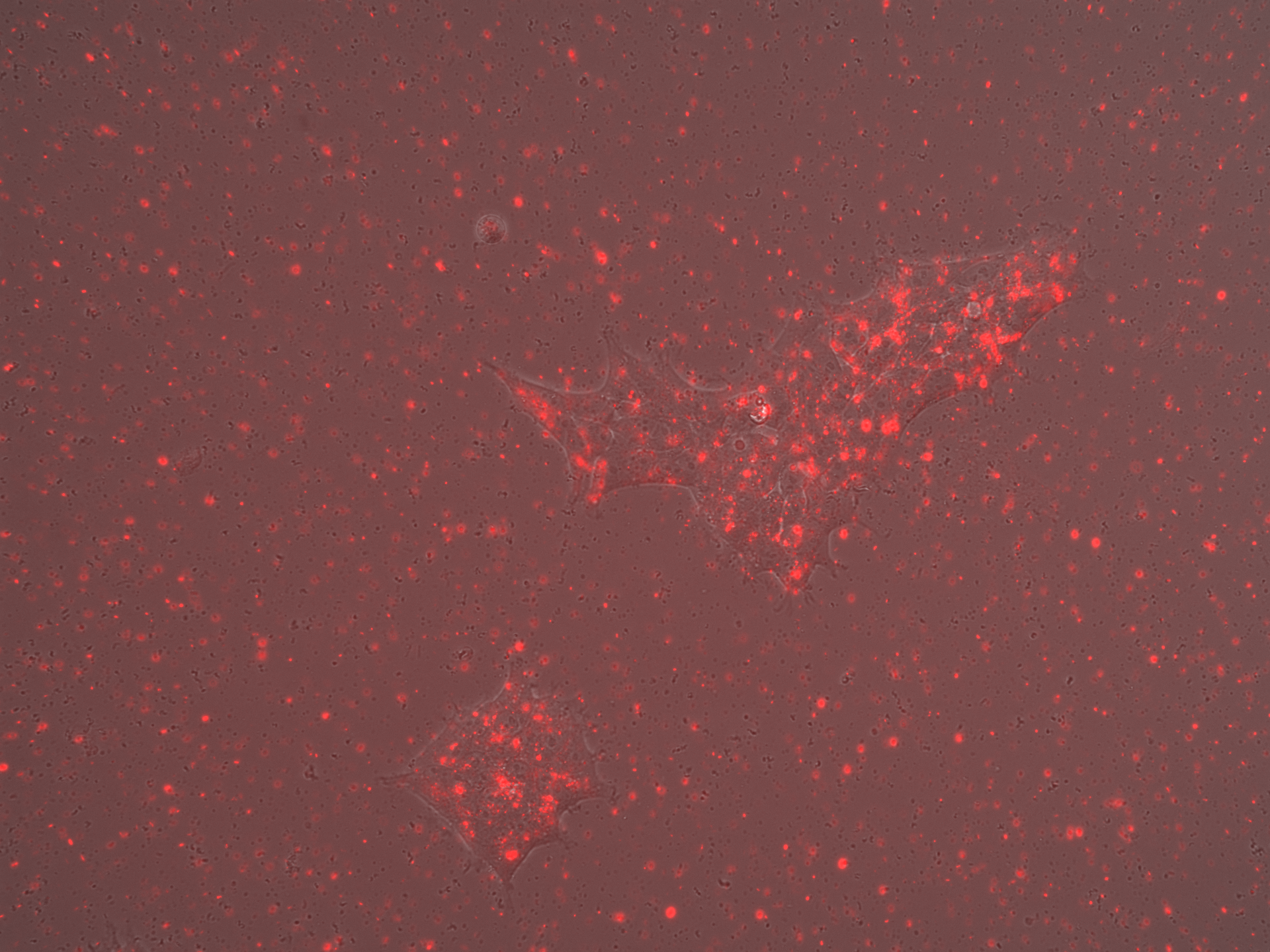

### No Rep-Cap Iodixanol_Overlay_Day3.tif

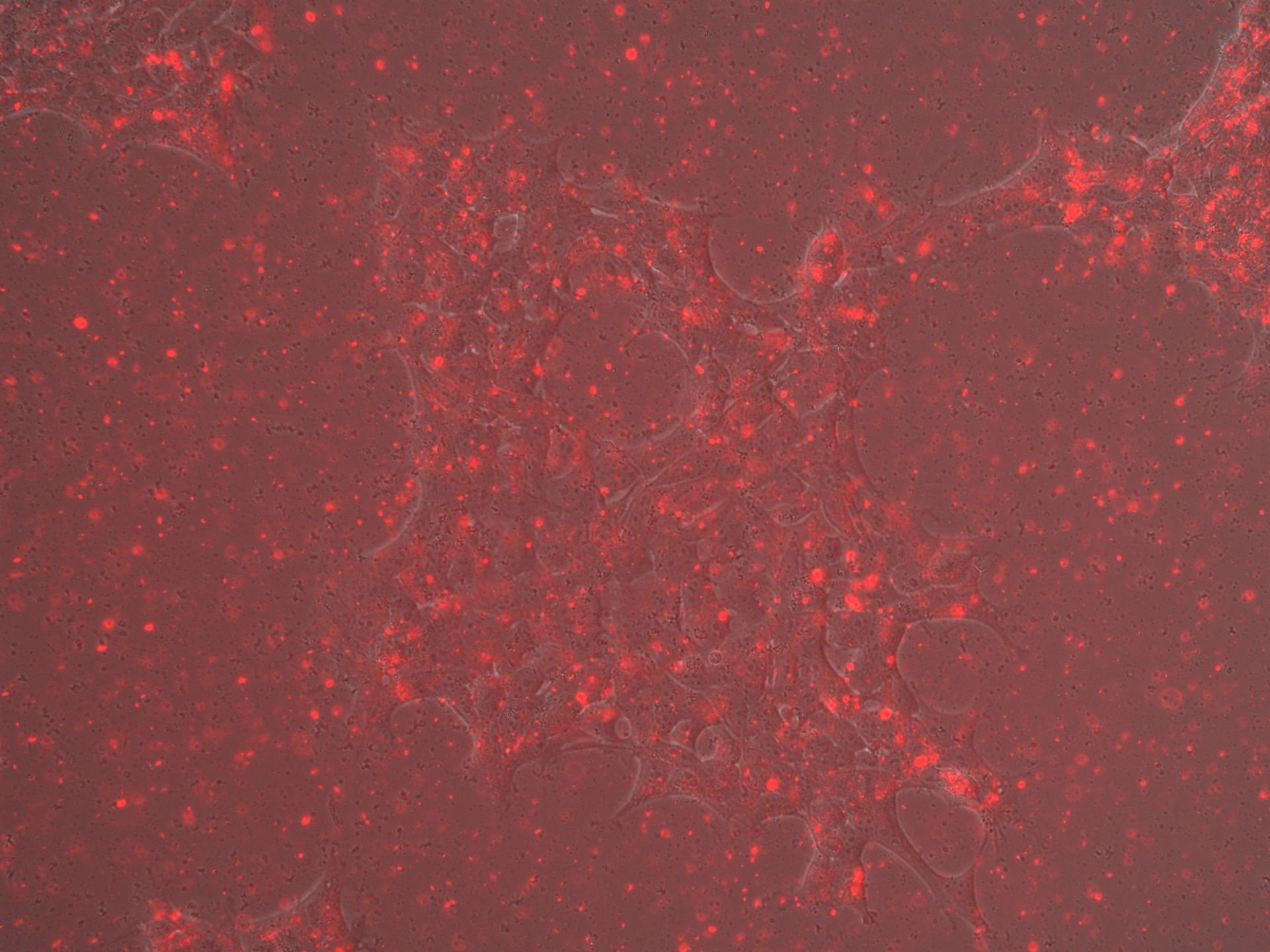

### No Rep-Cap Iodixanol_Overlay_Day5.tif

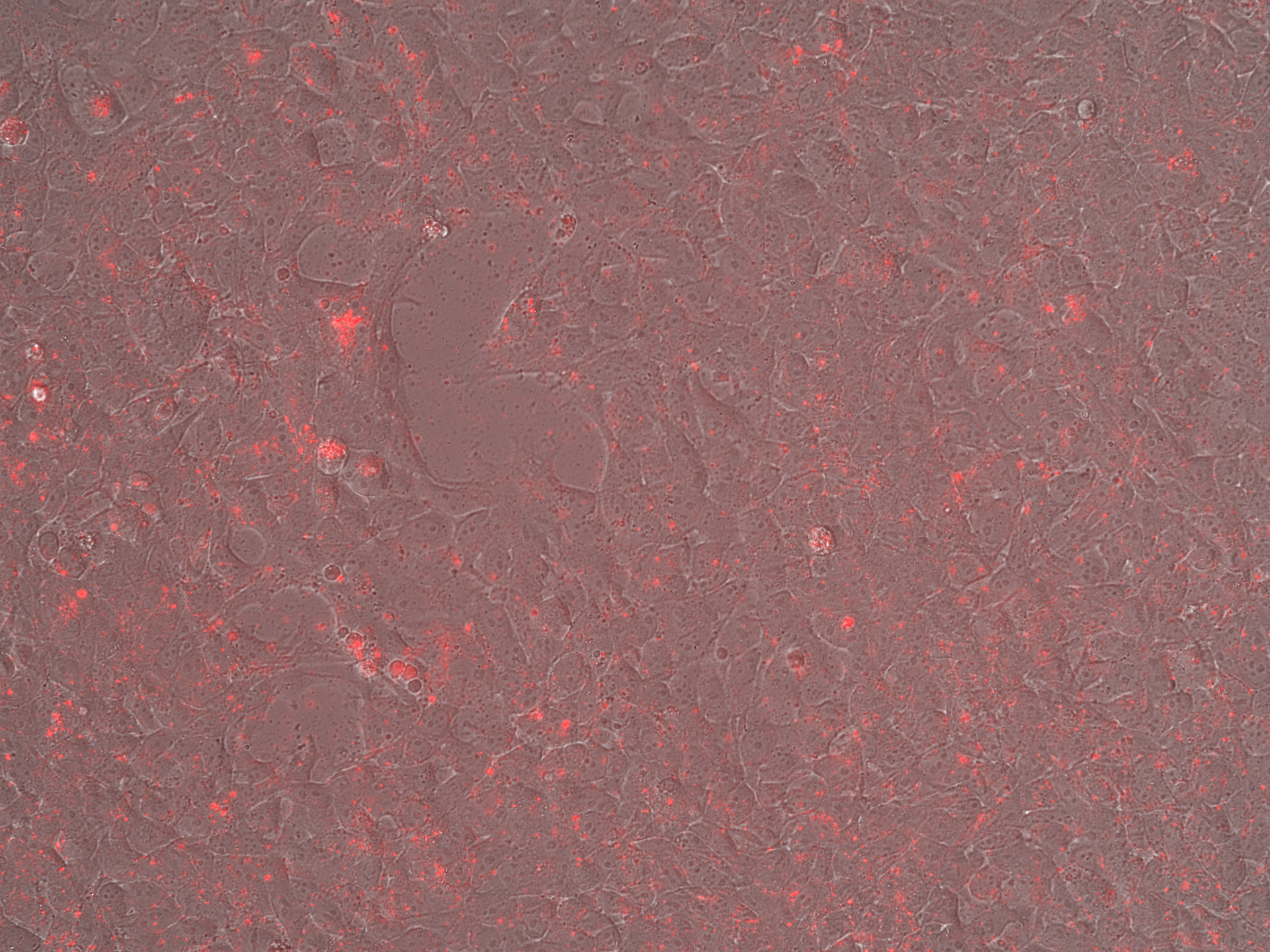

### No Rep-Cap Iodixanol_Overlay_Day7.tif

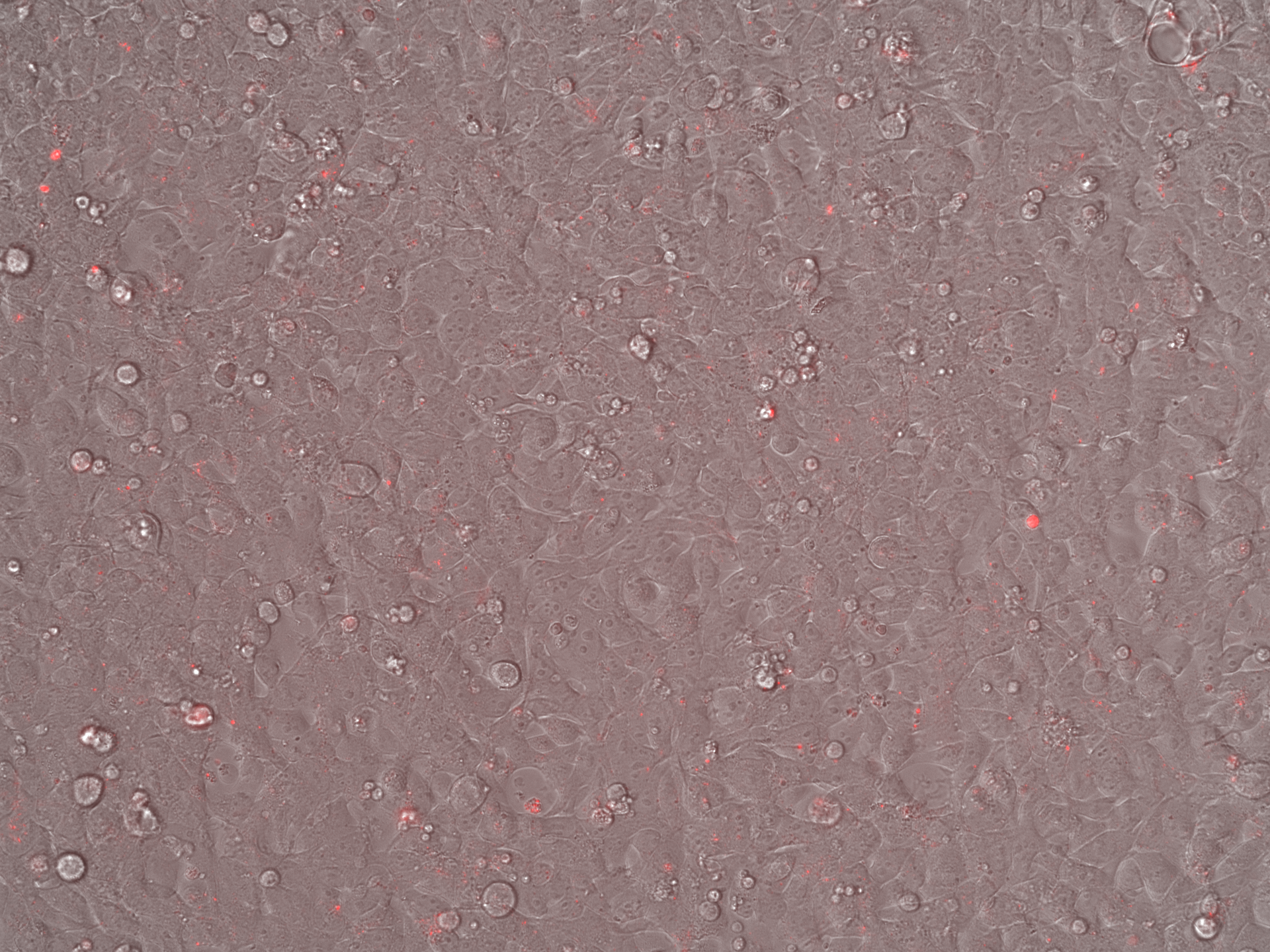

### No Rep-Cap Iodixanol_Overlay_Day7.tif

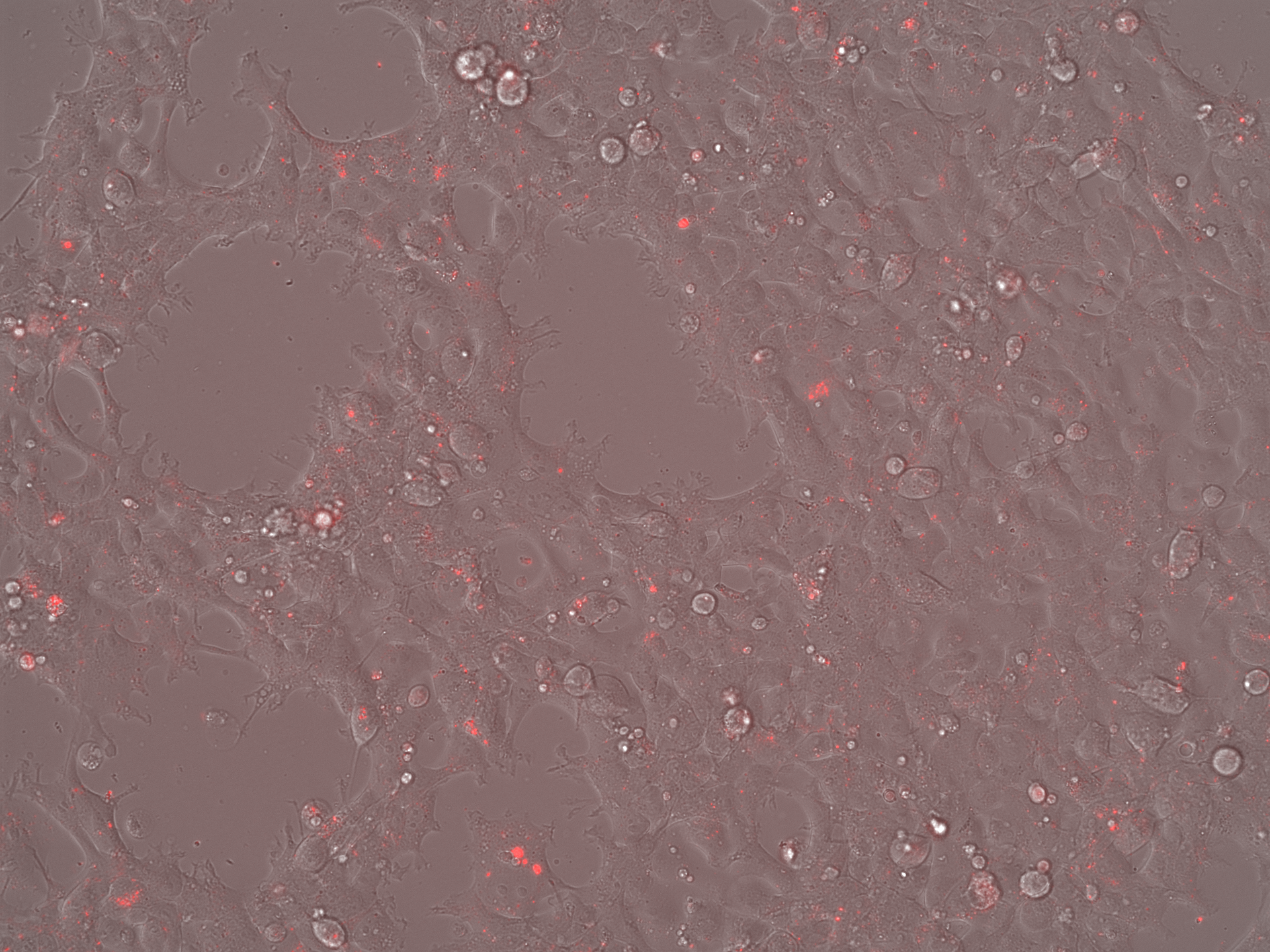

### No Rep-Cap SEC_Brightfield_Day3.tif

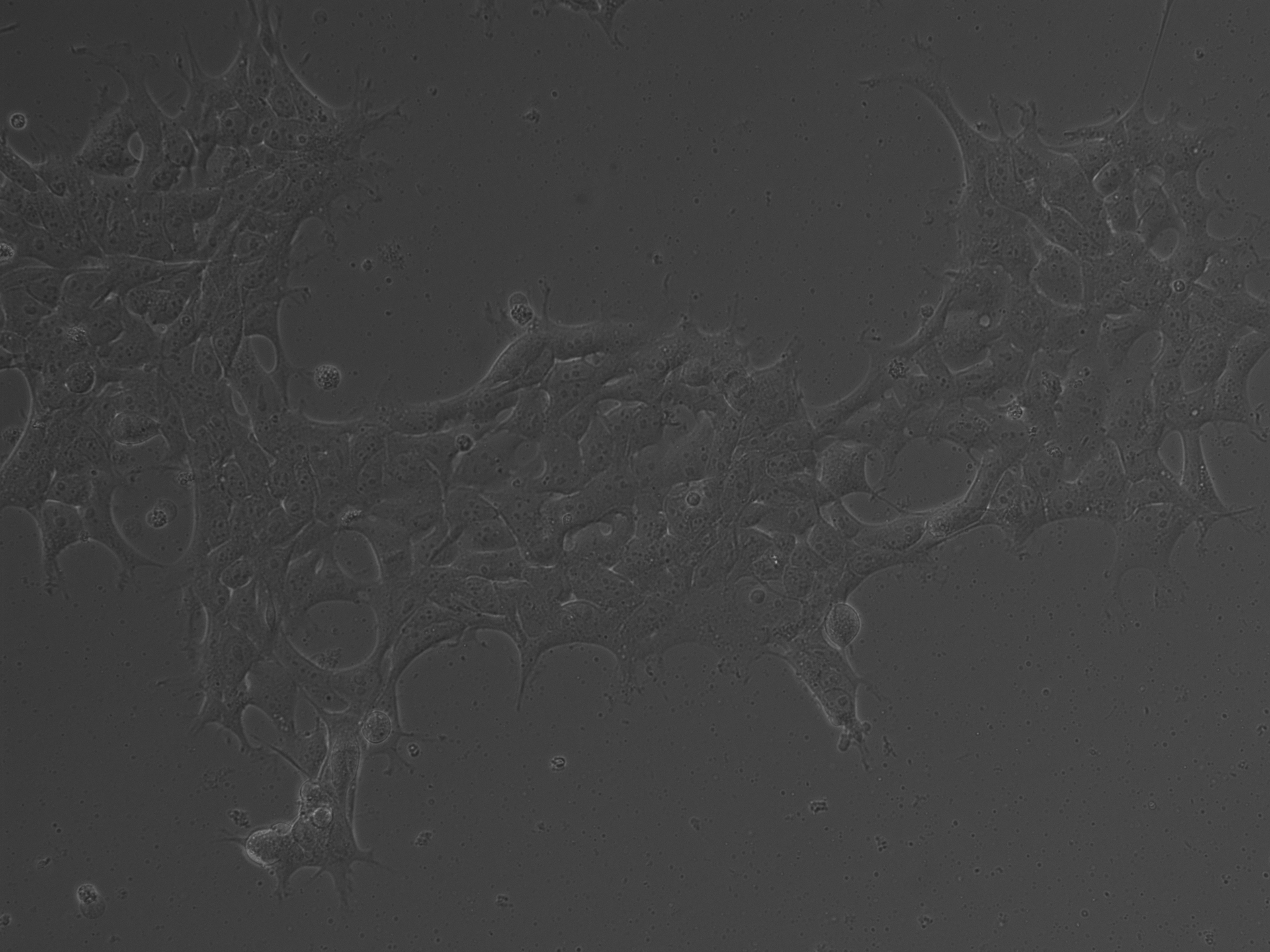

### No Rep-Cap SEC_Brightfield_Day3.tif

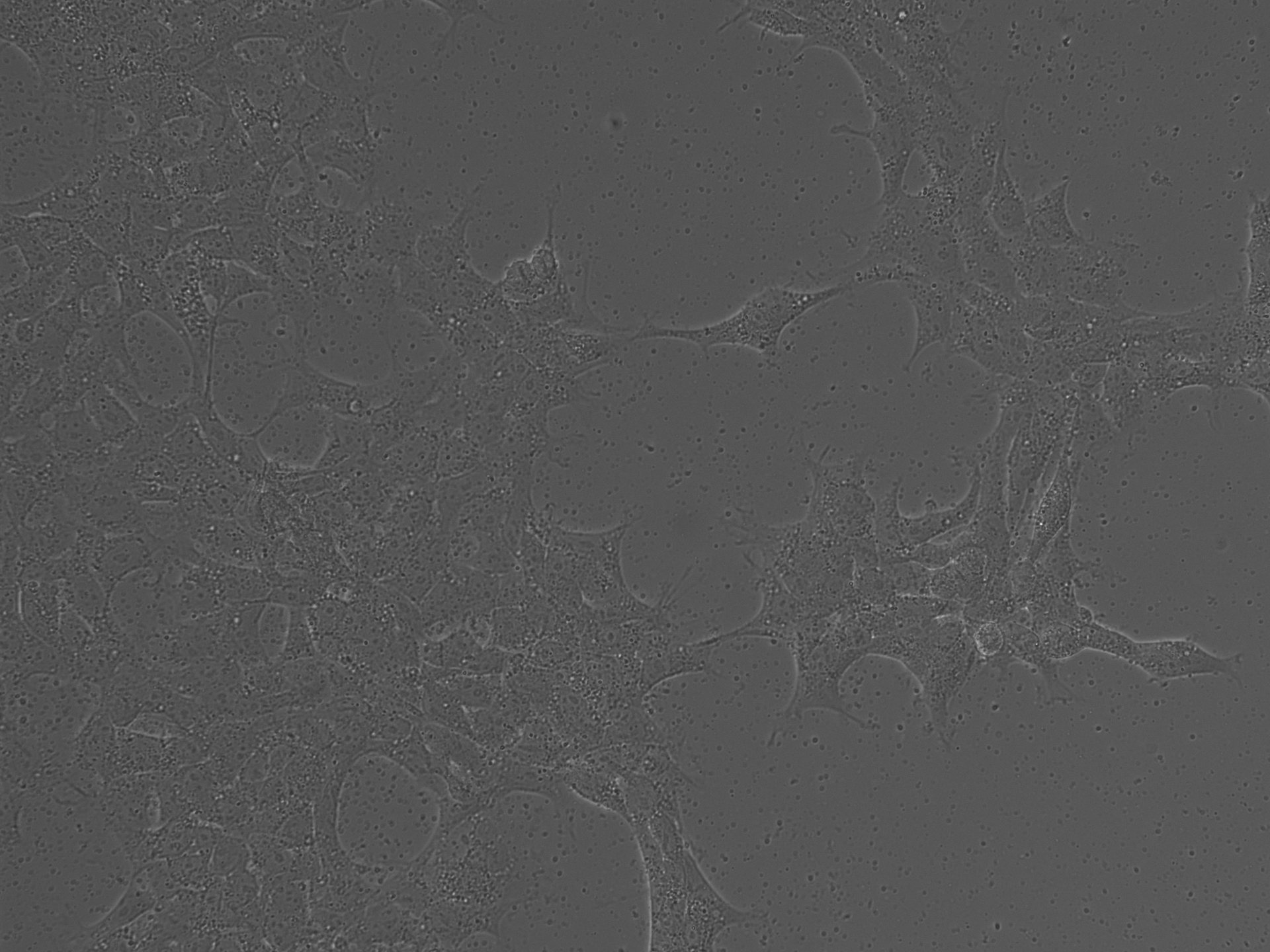

### No Rep-Cap SEC_Brightfield_Day5.tif

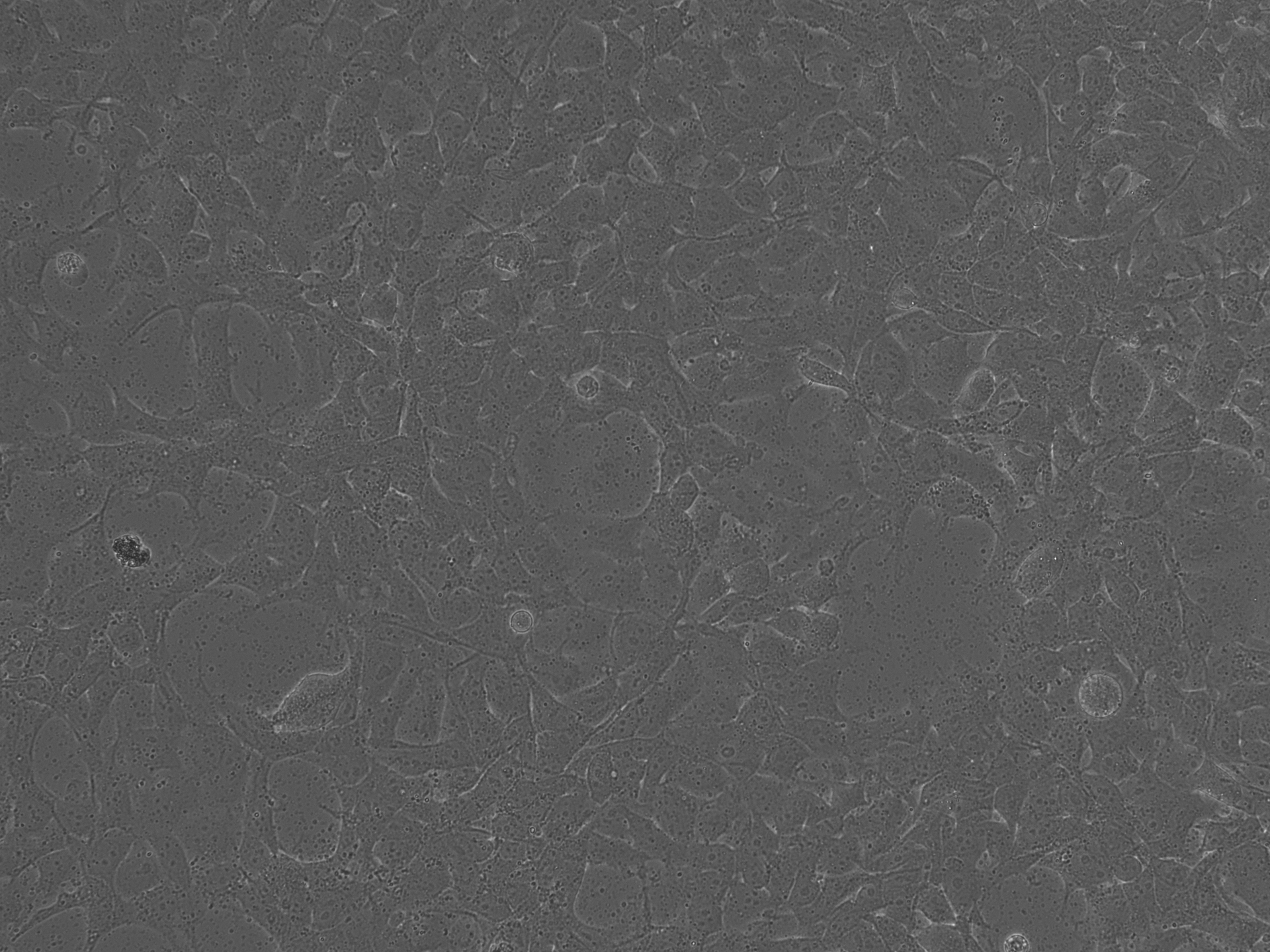

### No Rep-Cap SEC_Brightfield_Day7.tif

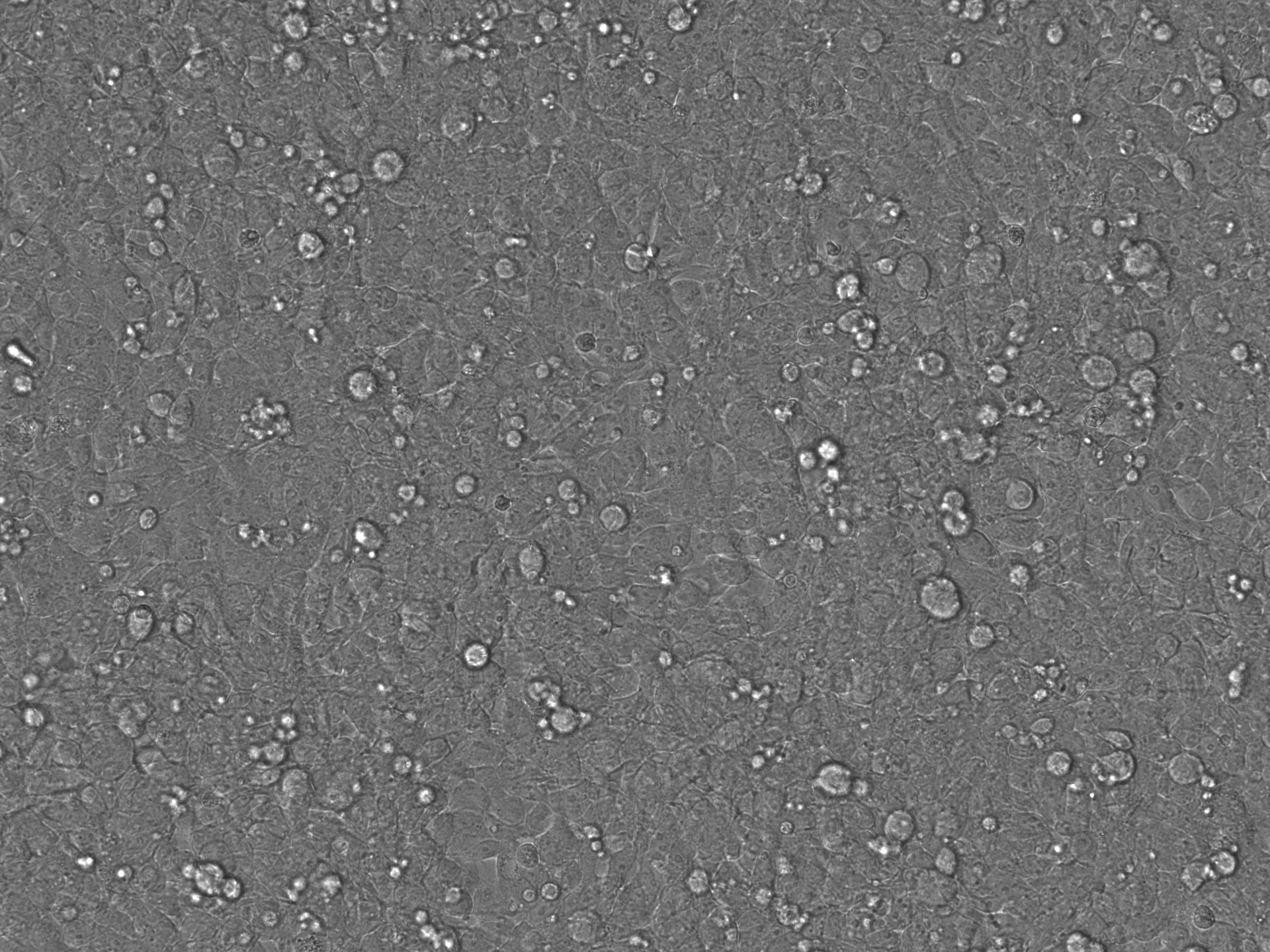

### No Rep-Cap SEC_Brightfield_Day7.tif

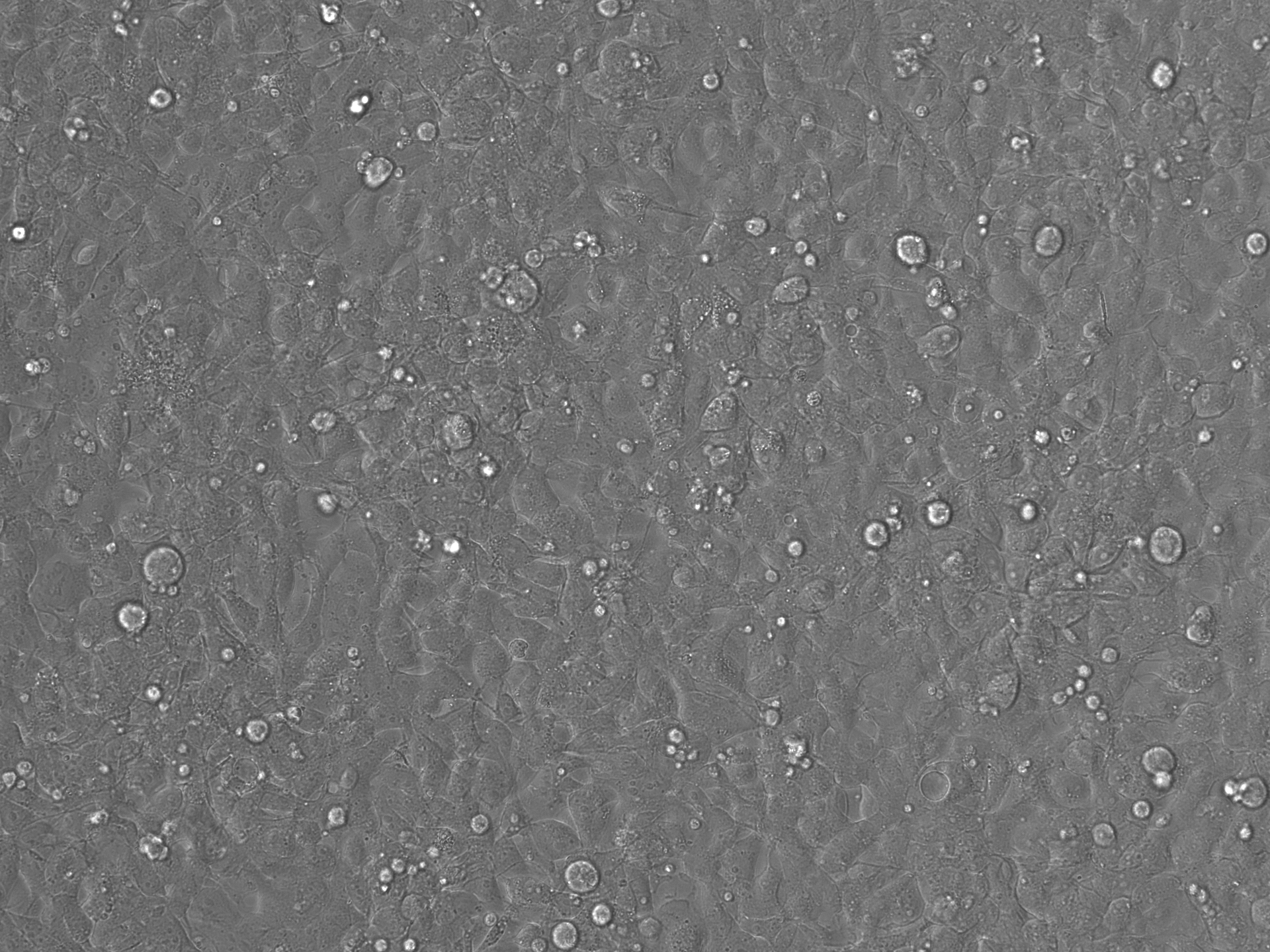

### No Rep-Cap SEC_LumiScarlet_Day3.tif

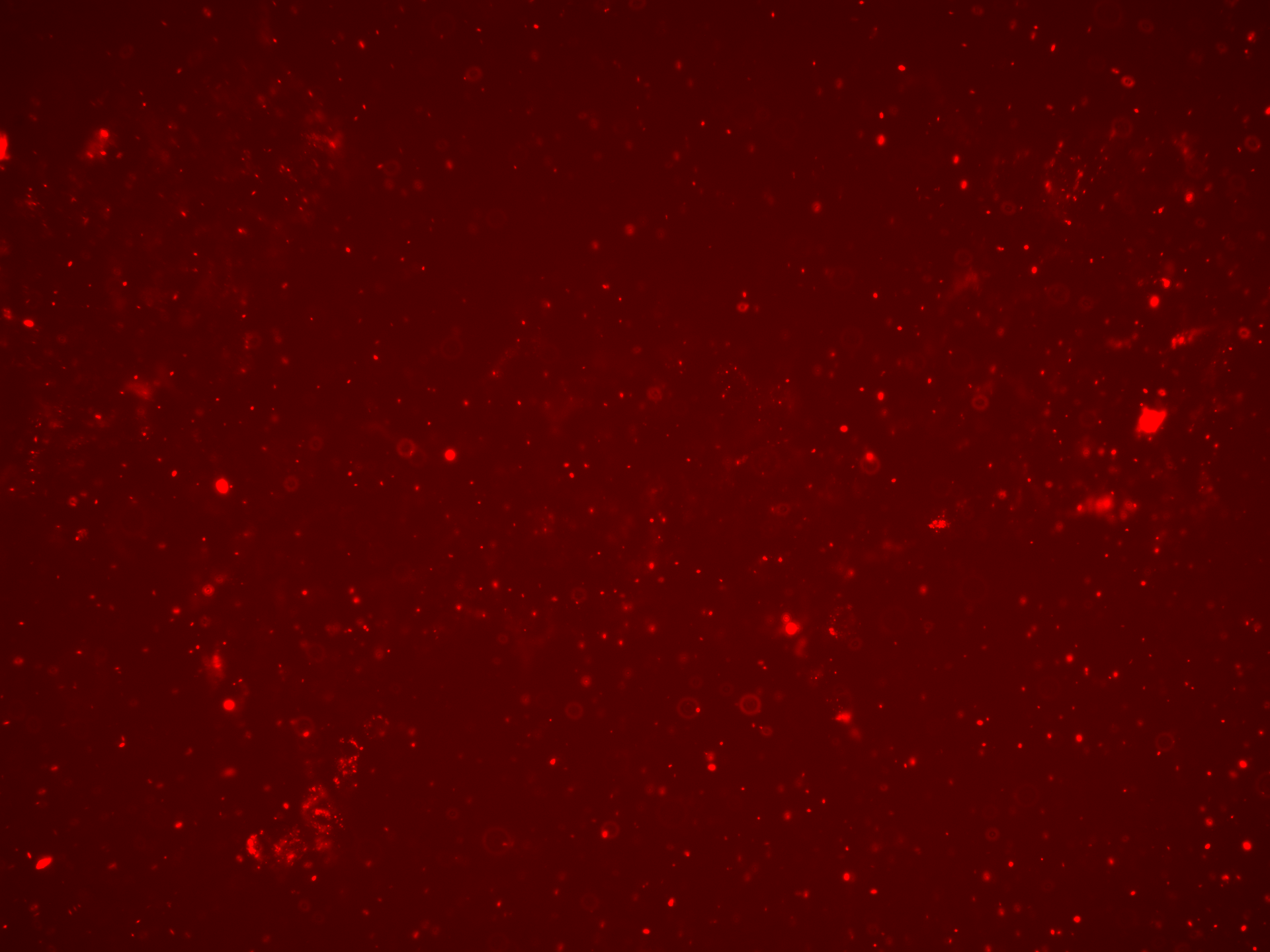

### No Rep-Cap SEC_LumiScarlet_Day3.tif

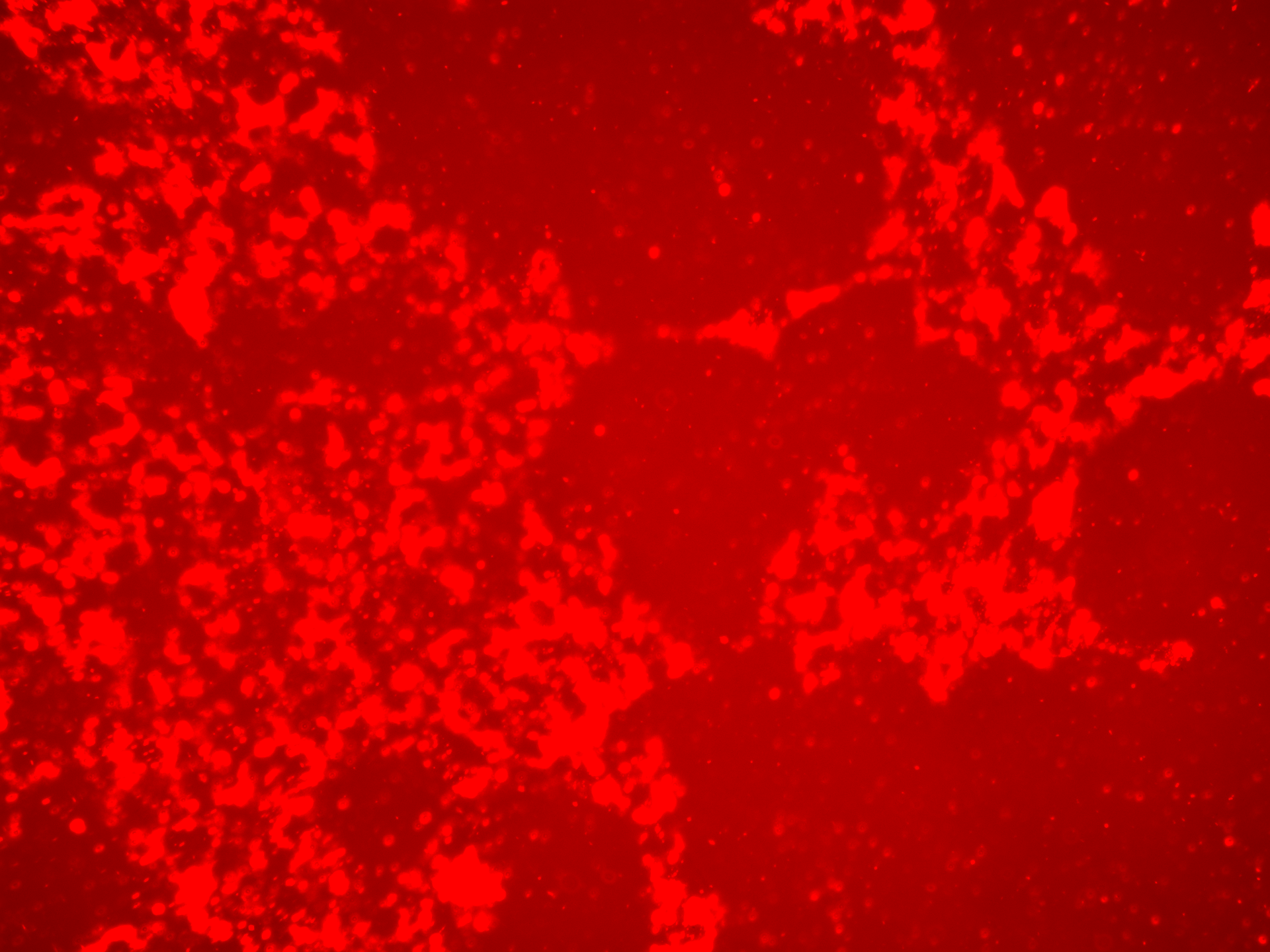

### No Rep-Cap SEC_LumiScarlet_Day5.tif

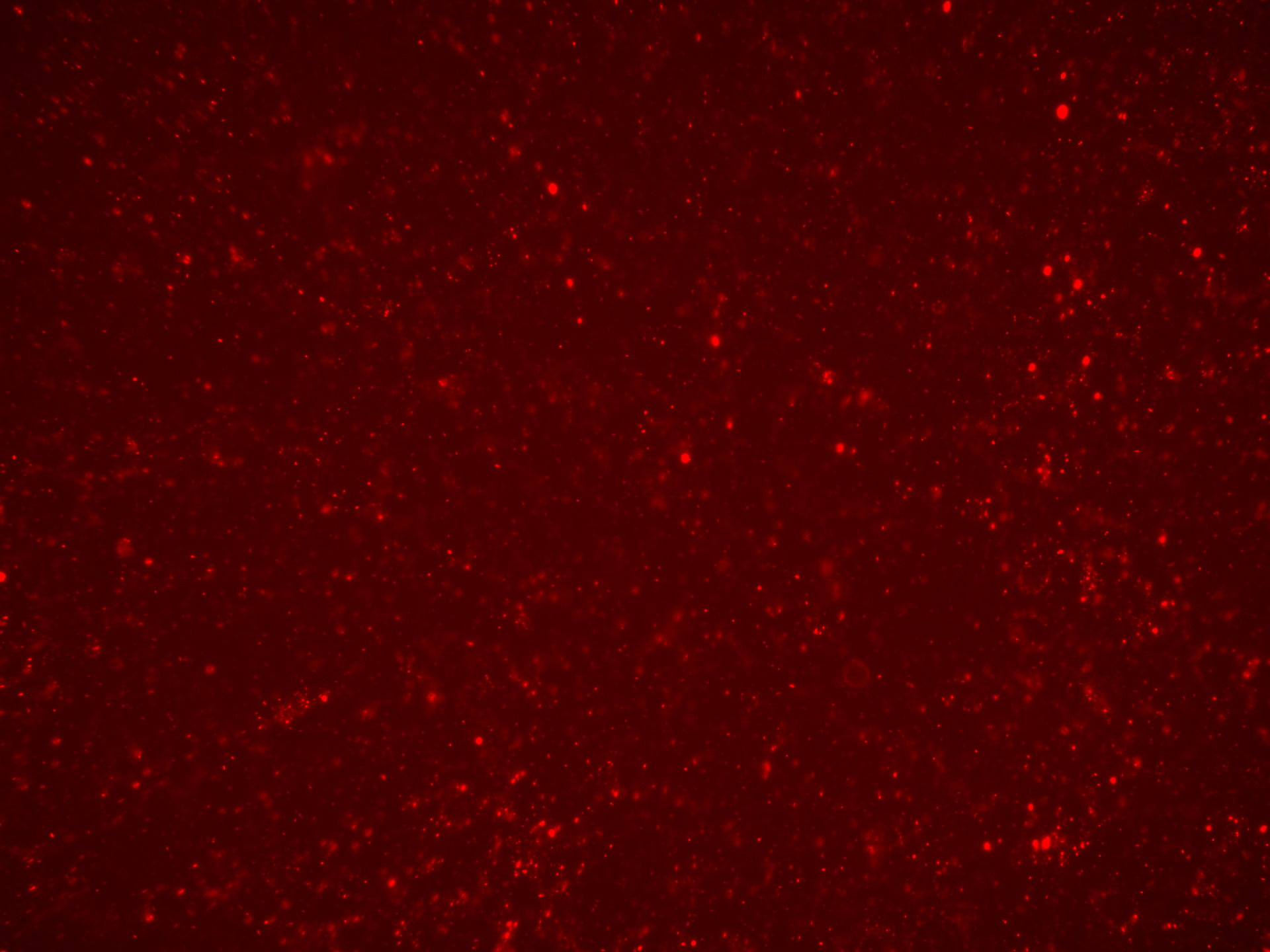

### No Rep-Cap SEC_LumiScarlet_Day7.tif

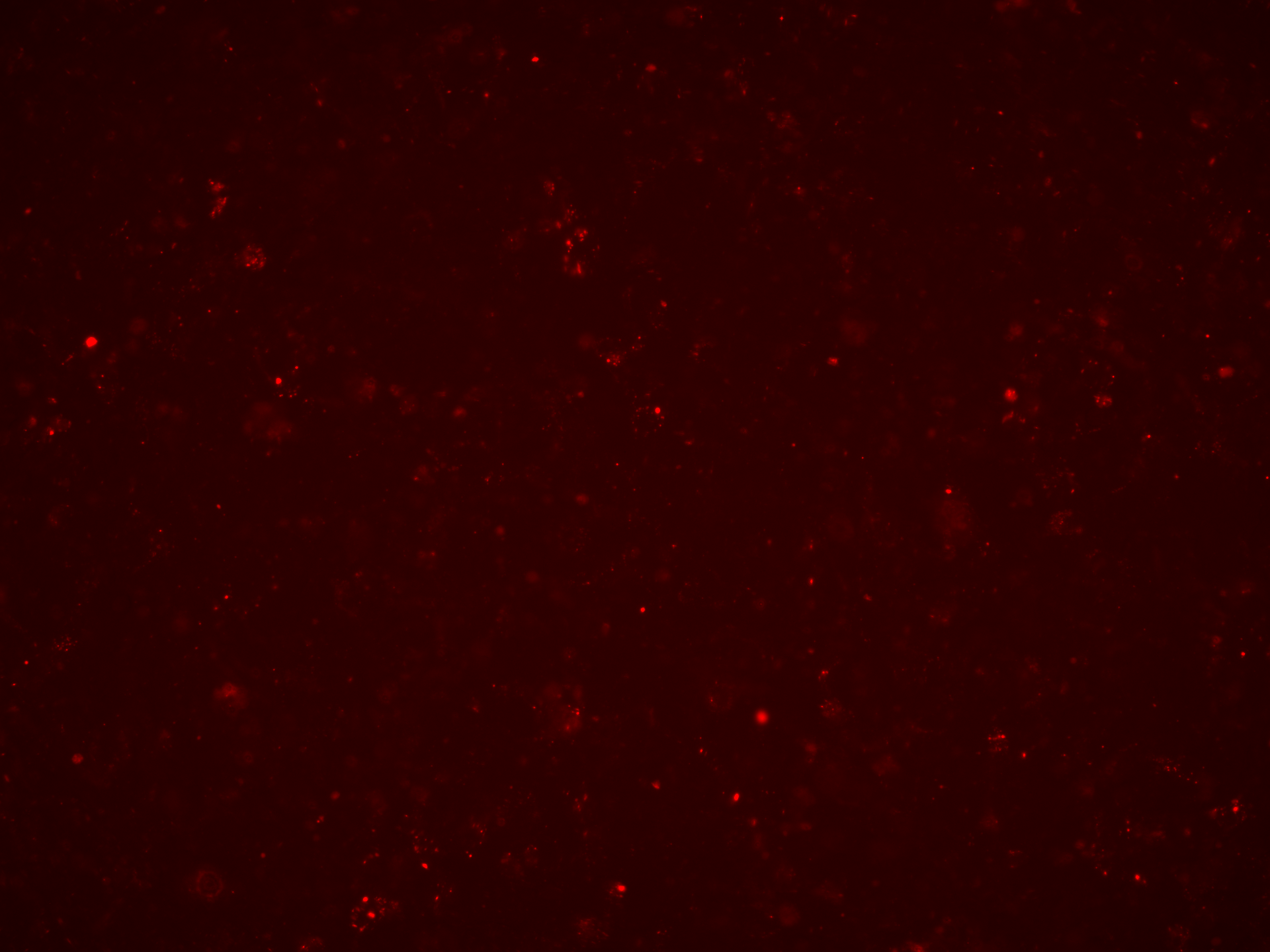

### No Rep-Cap SEC_LumiScarlet_Day7.tif

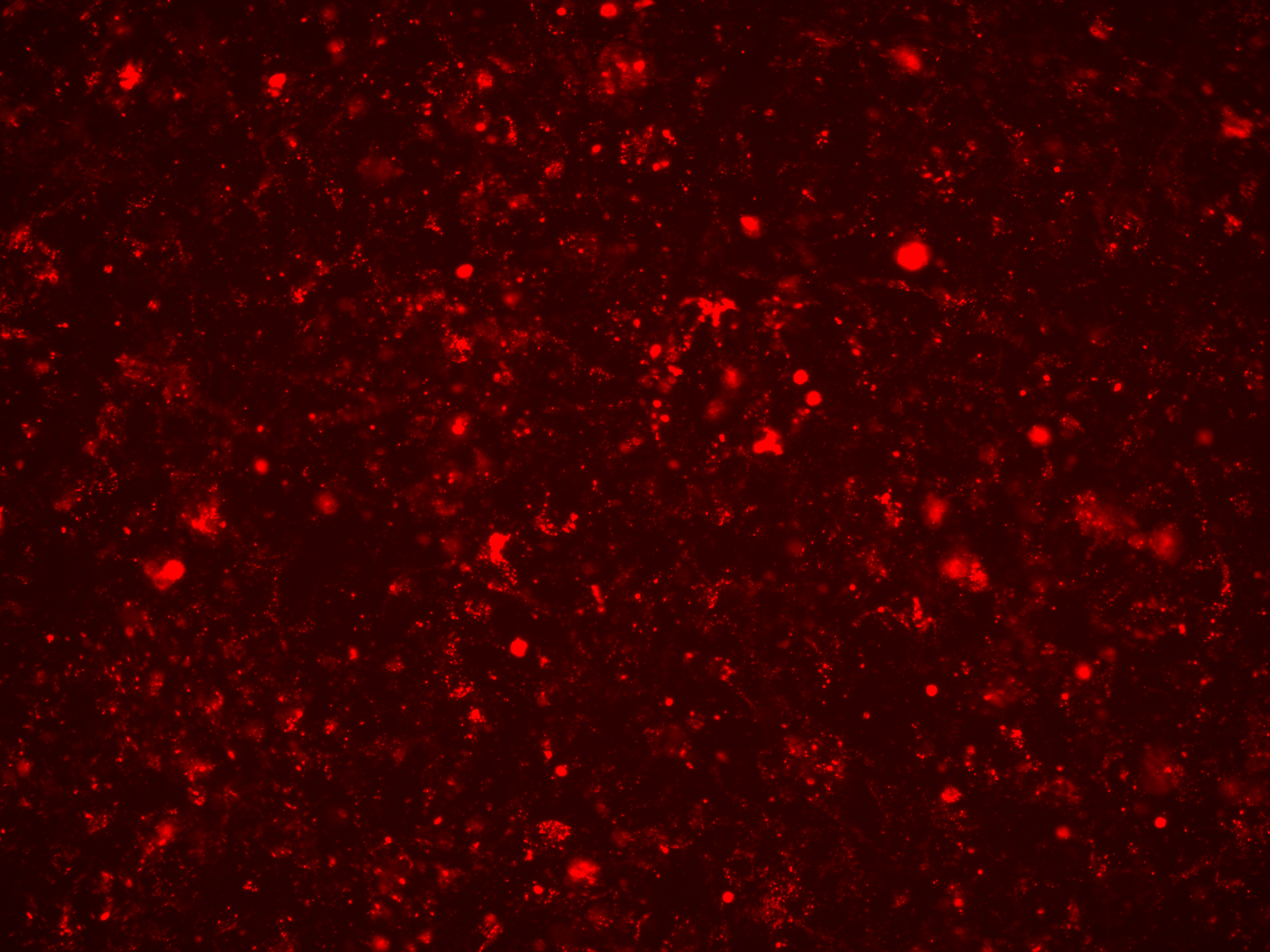

### No Rep-Cap SEC_Overlay_Day3.tif

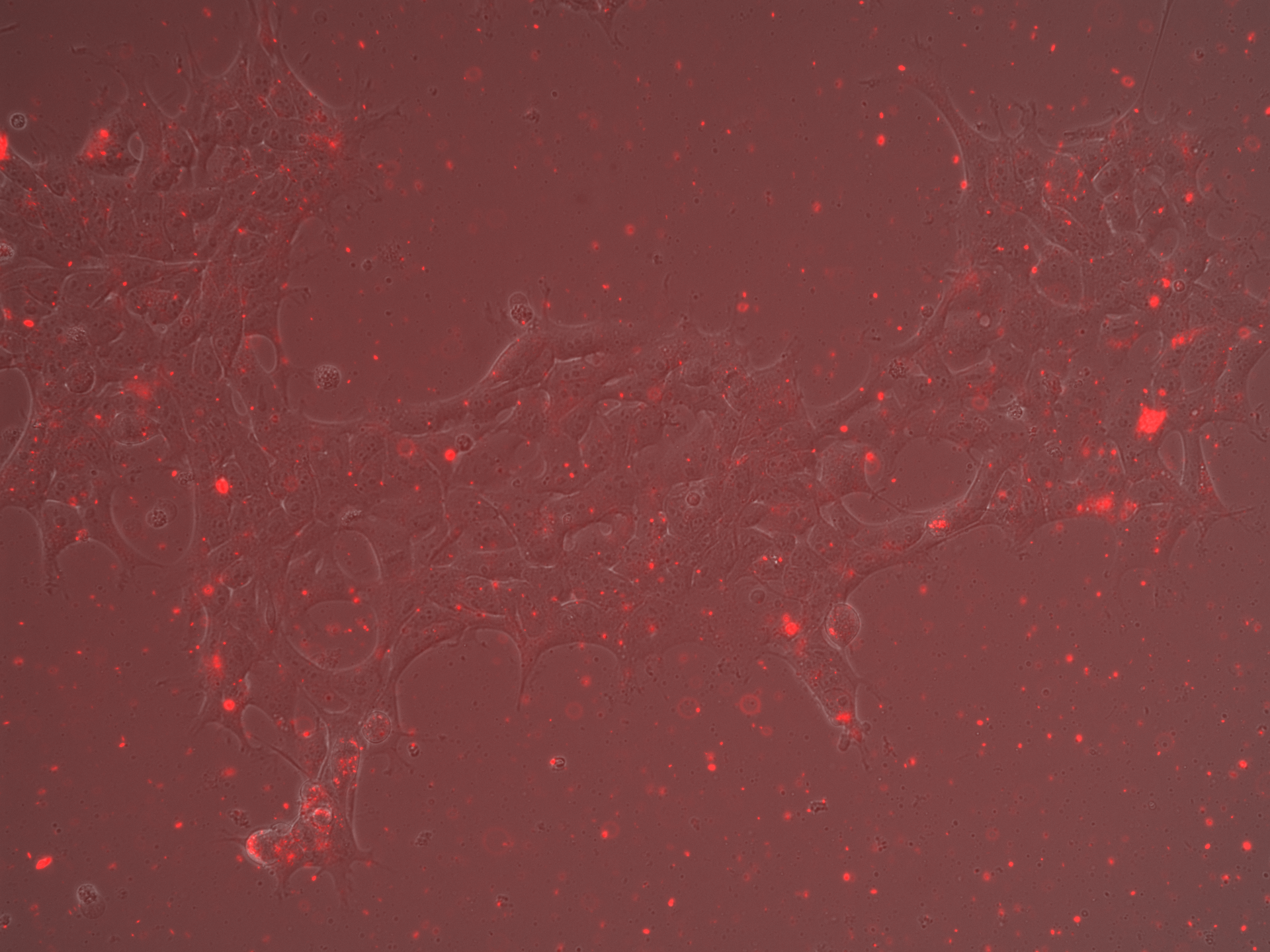

### No Rep-Cap SEC_Overlay_Day3.tif

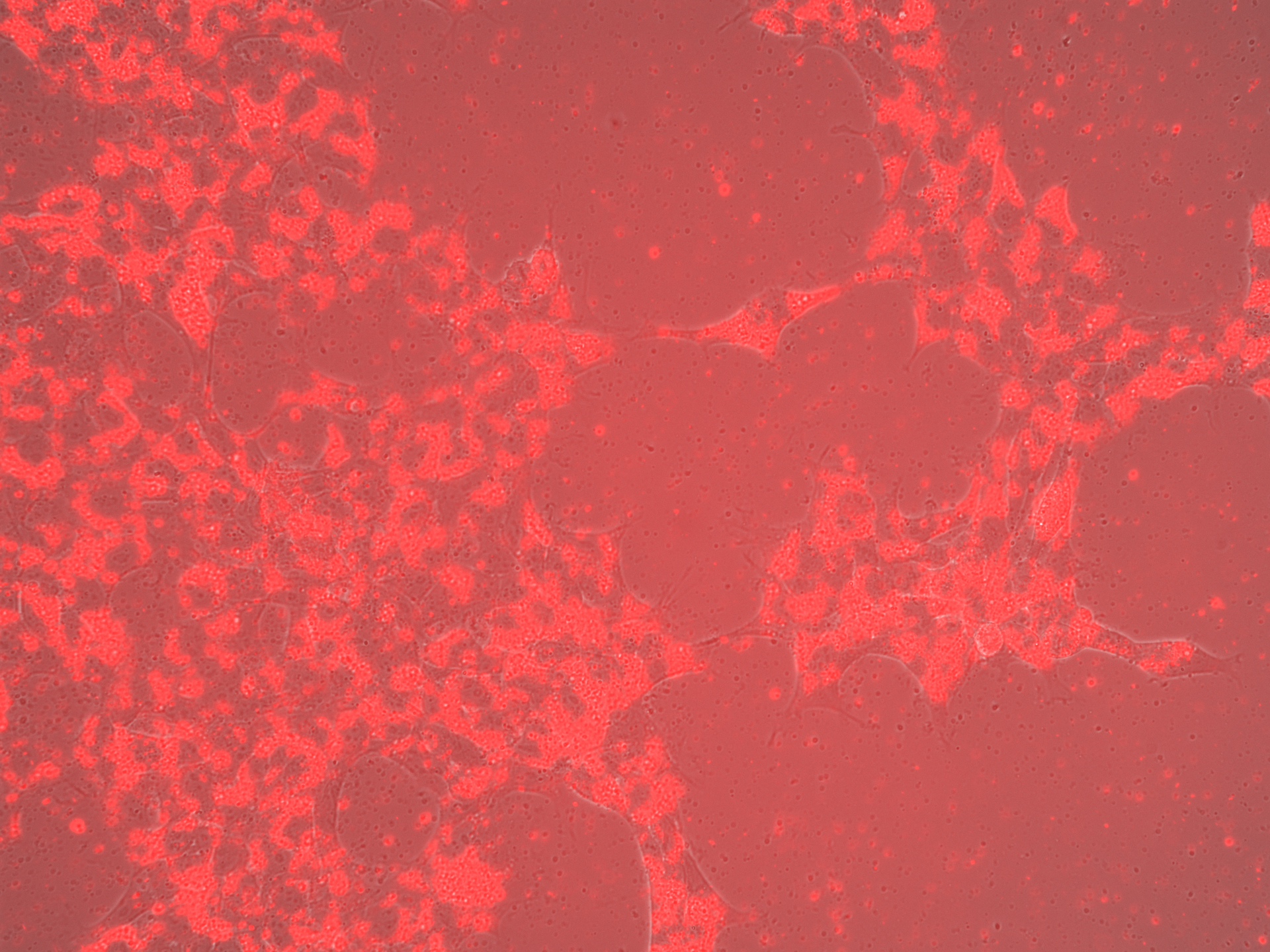

### No Rep-Cap SEC_Overlay_Day5.tif

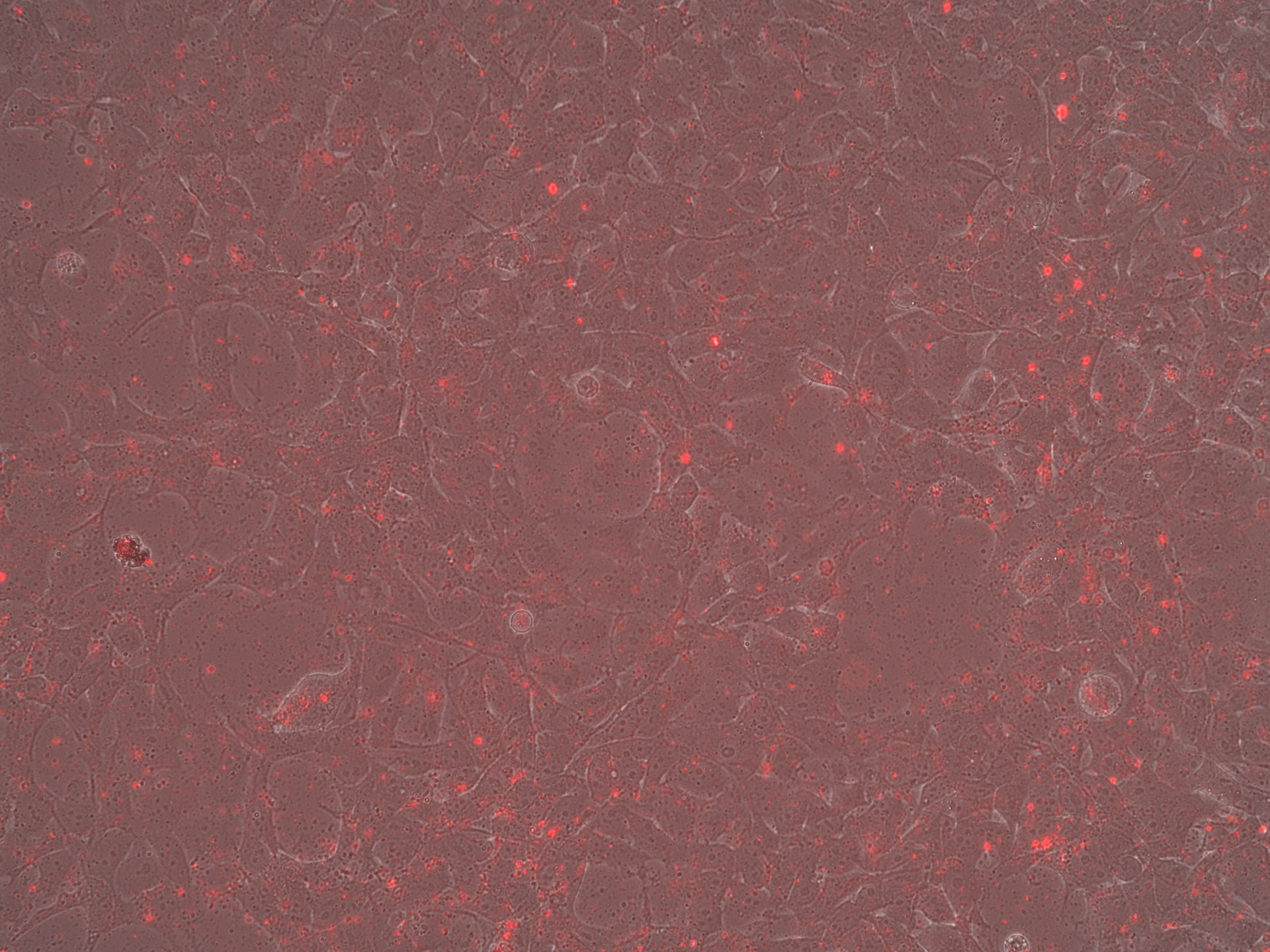

### No Rep-Cap SEC_Overlay_Day7.tif

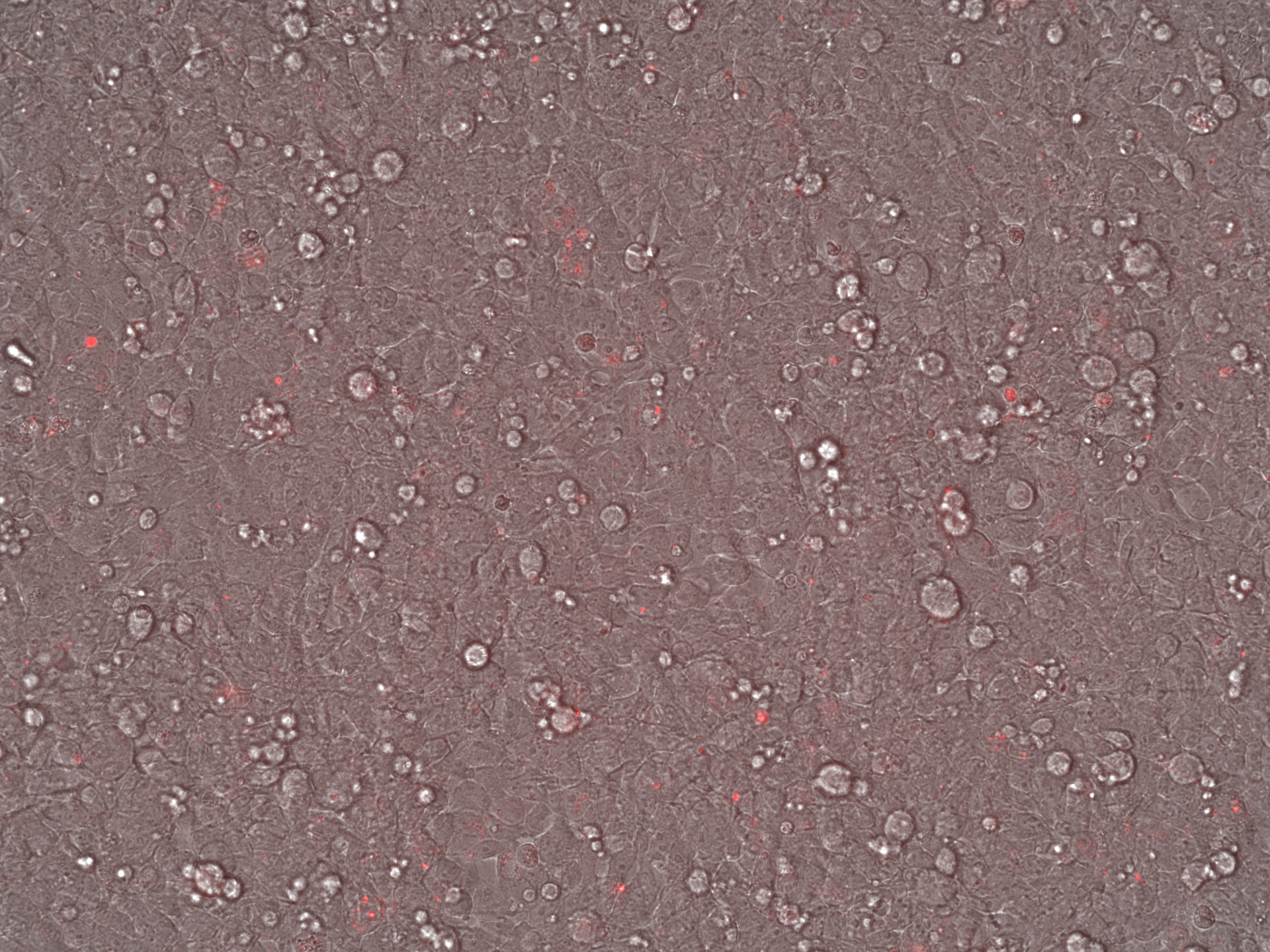

### No Rep-Cap SEC_Overlay_Day7.tif

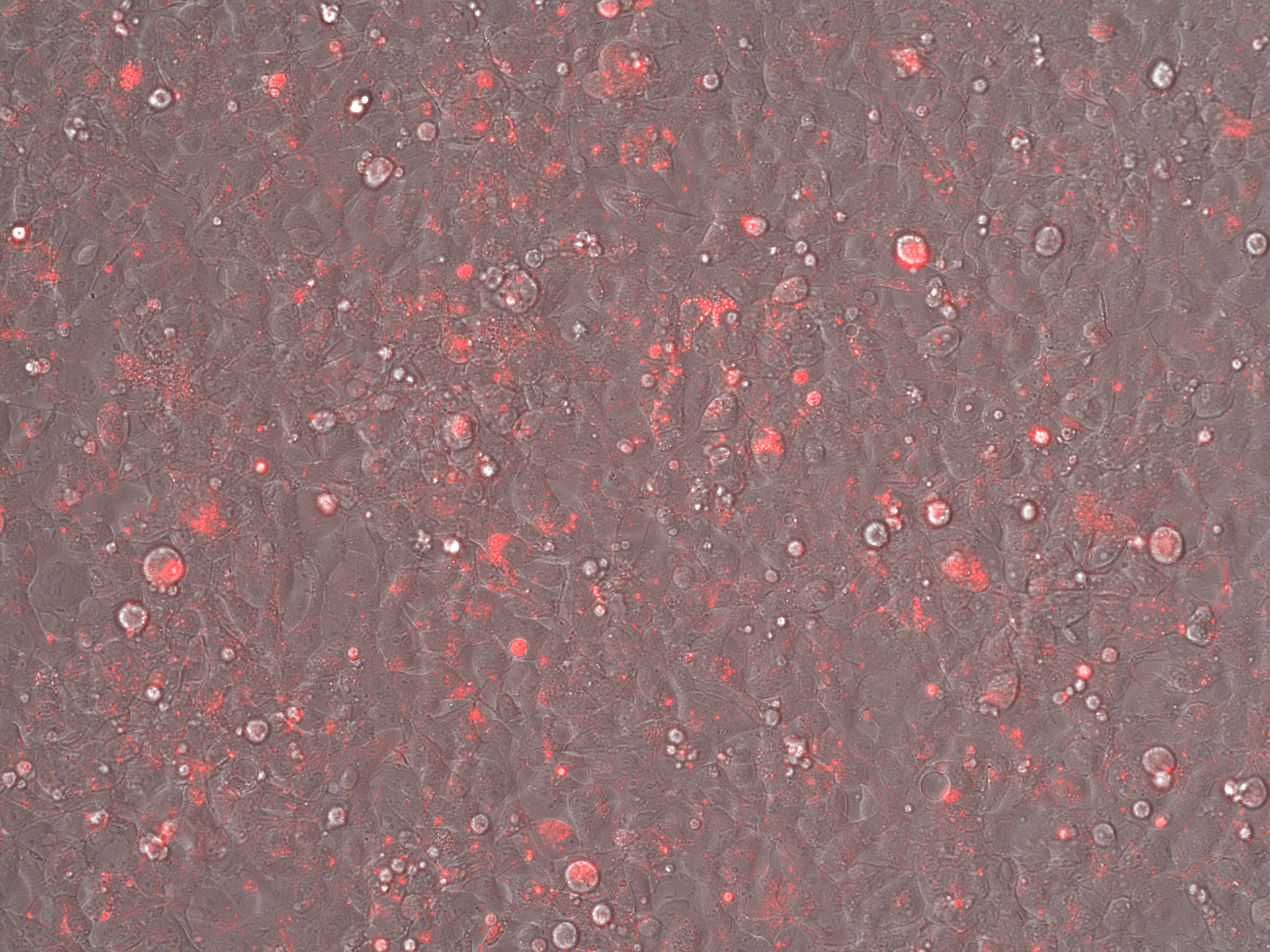

### No RepCap Iodixanol_Brightfield_Day5.tif

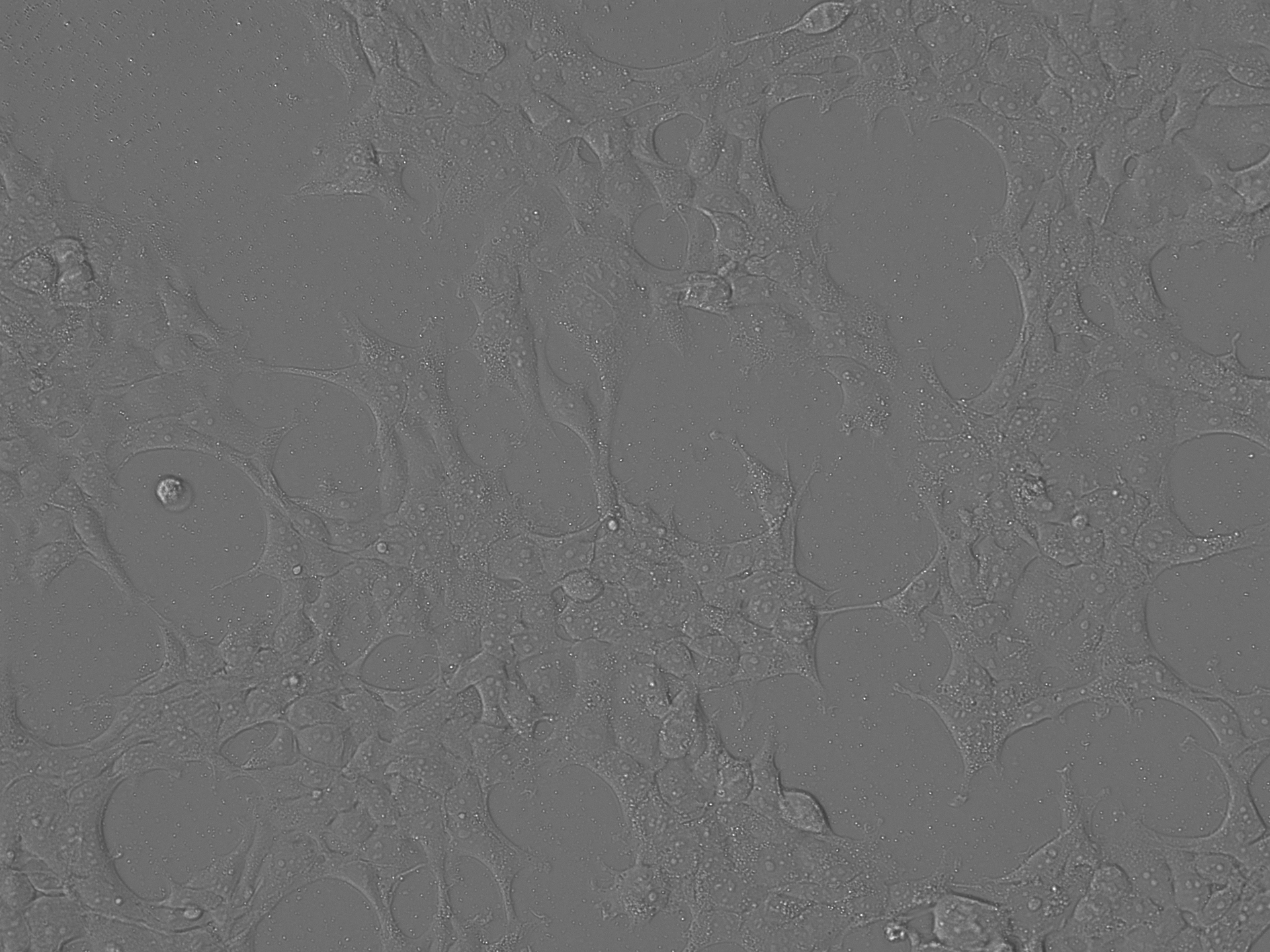

### No RepCap Iodixanol_LumiScarlet_Day5.tif

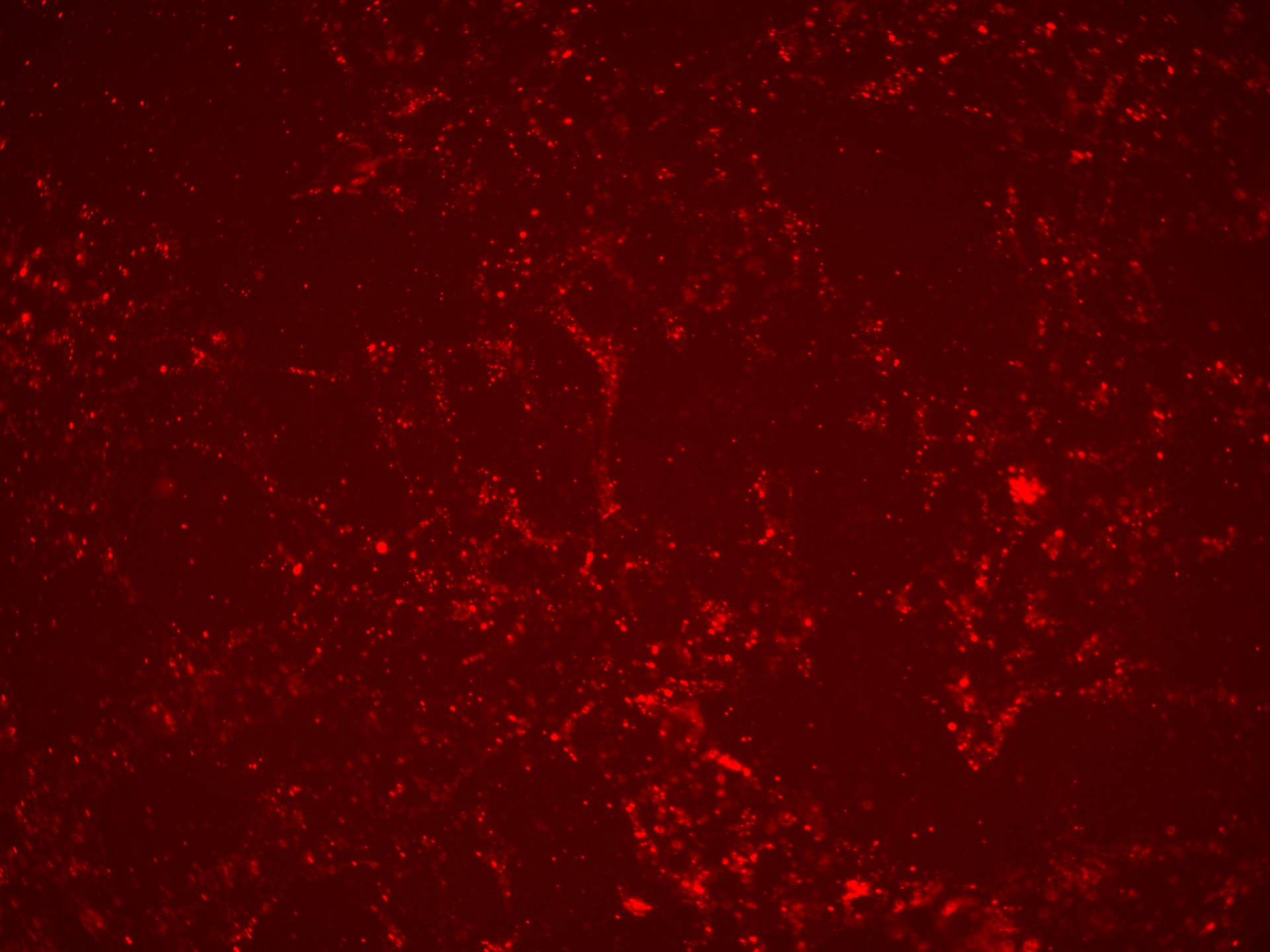

### No RepCap Iodixanol_Overlay_Day5.tif

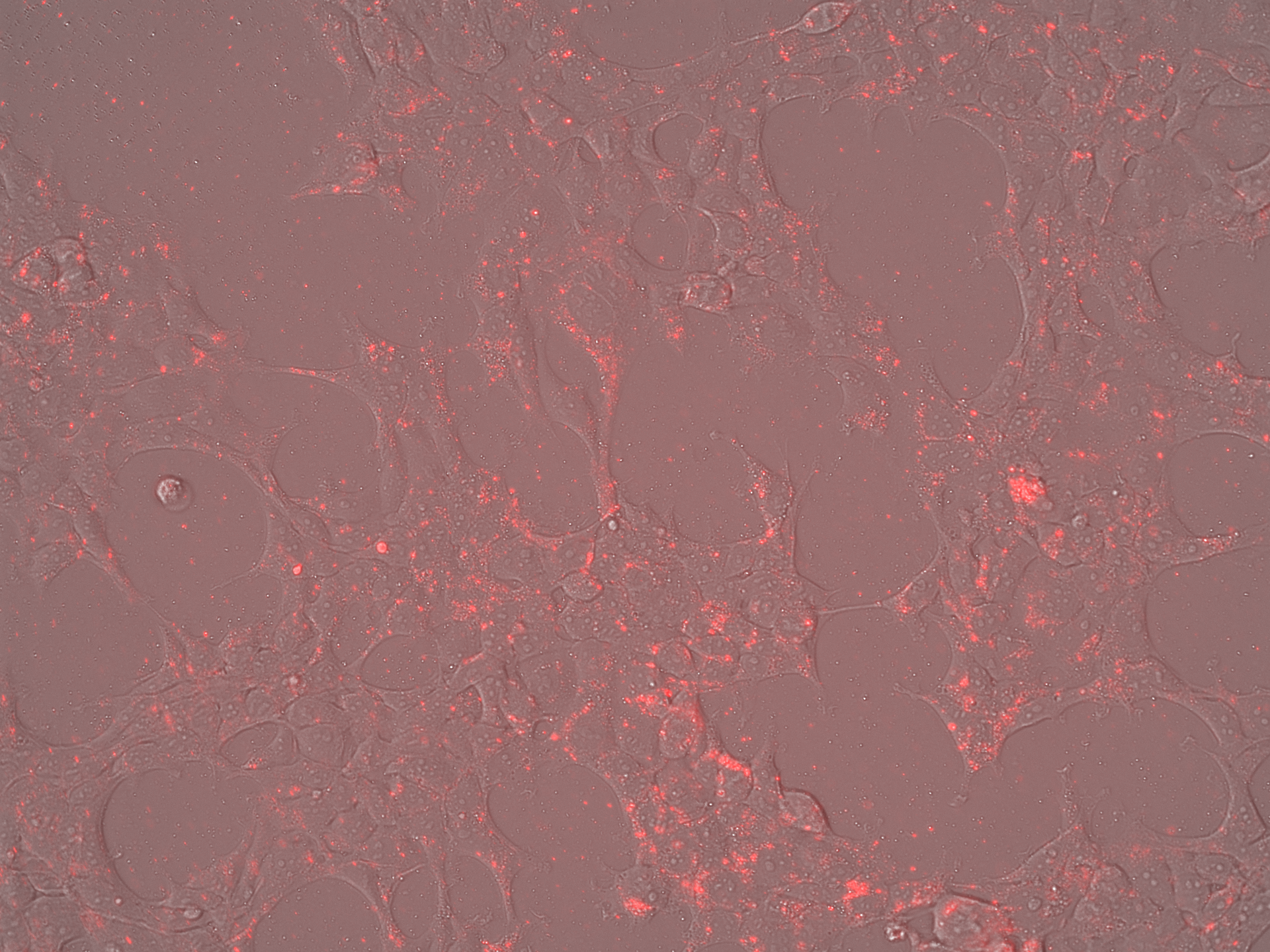

### No RepCap SECl_Brightfield_Day5.tif

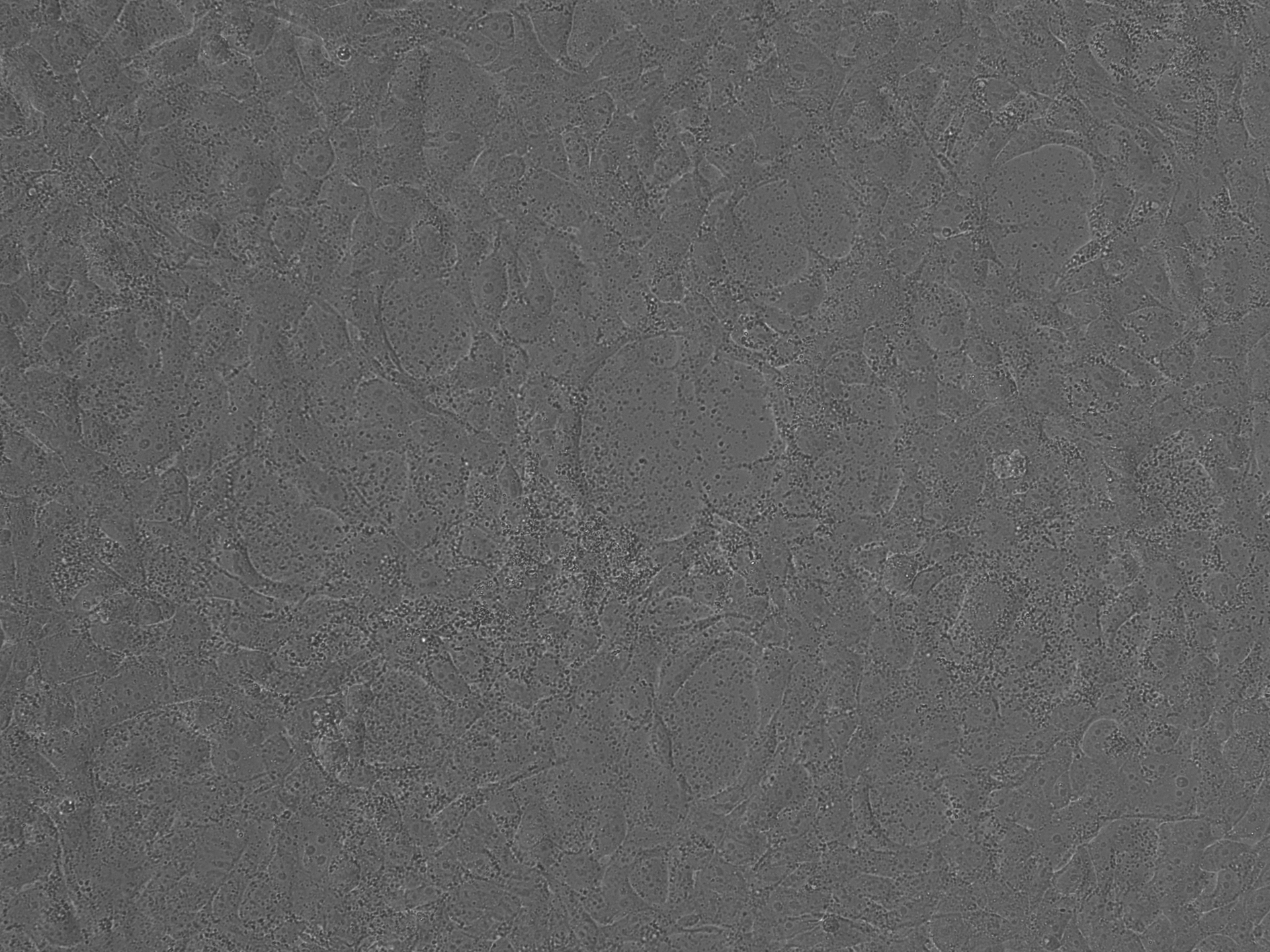

### No RepCap SECl_LumiScarlet_Day5.tif

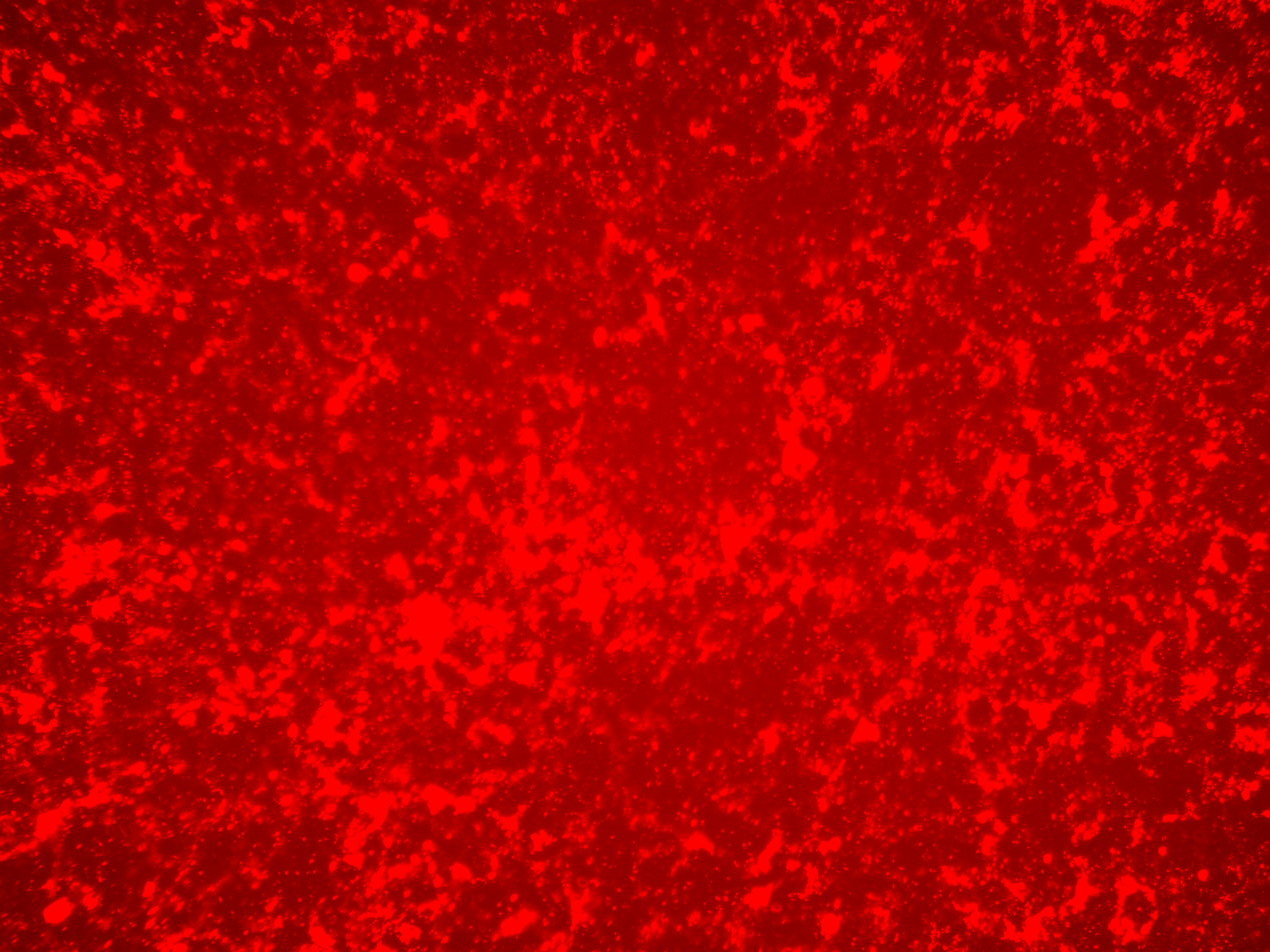

### No RepCap SECl_Overlay_Day5.tif

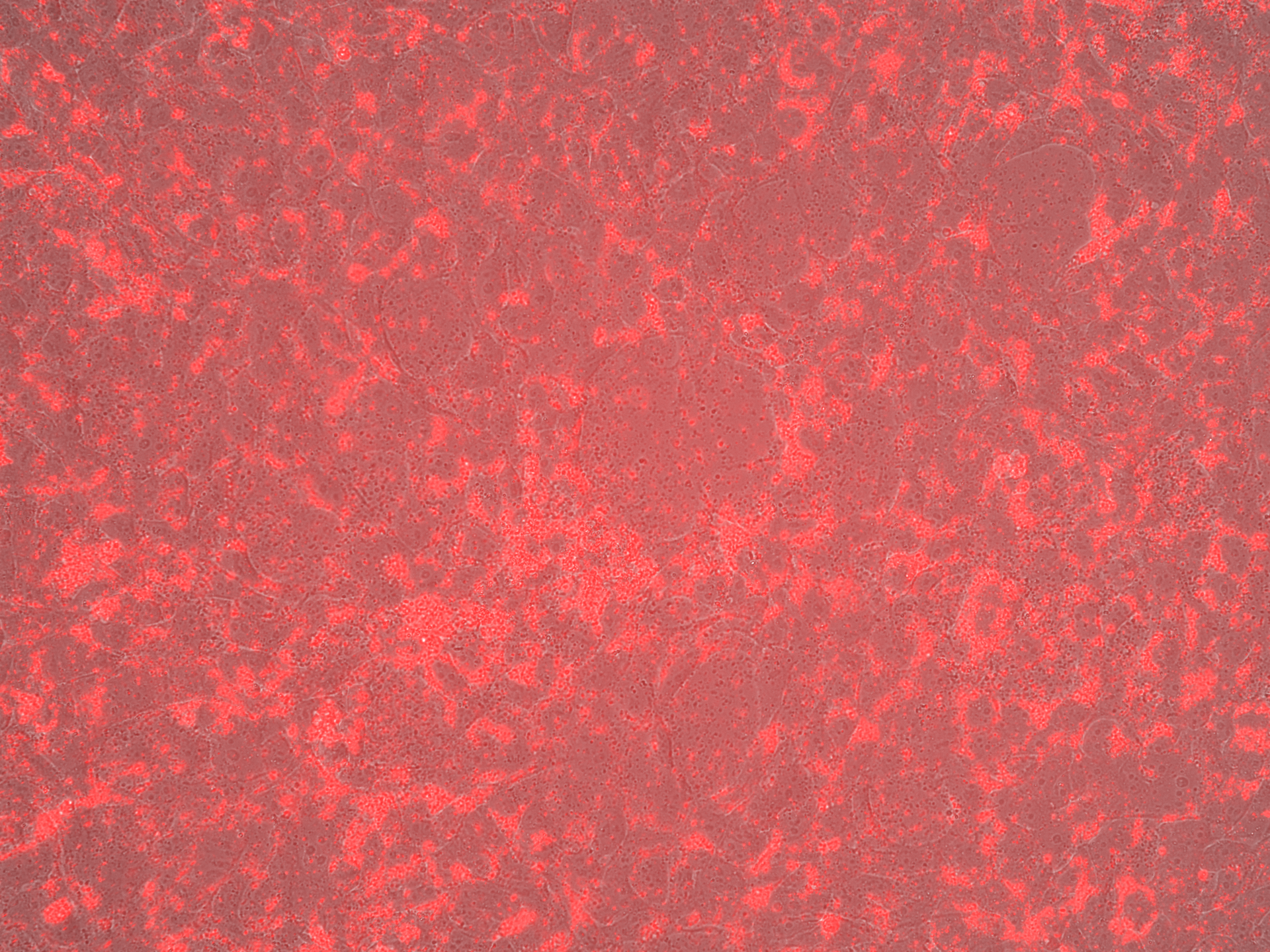
